# SPACE: STRING proteins as complementary embeddings

**DOI:** 10.1101/2024.11.25.625140

**Authors:** Dewei Hu, Damian Szklarczyk, Christian von Mering, Lars Juhl Jensen

**Affiliations:** Novo Nordisk Foundation Center For Protein Research, Faculty of Health and Medical Sciences, University of Copenhagen, Copenhagen, Denmark; Department of Molecular Life Sciences, University of Zurich, Zurich, Switzerland

**Keywords:** Protein embedding, Networks, Function prediction

## Abstract

Representation learning has revolutionized sequence-based prediction of protein function and subcellular localization. Protein networks are an important source of information complementary to sequences, but the use of protein networks has proven to be challenging in the context of machine learning, especially in a cross-species setting. To address this, we leveraged the STRING database of protein networks and orthology relations for 1,322 eukaryotes to generate network-based cross-species protein embeddings. We did this by first creating species-specific network embeddings and subsequently aligning them based on orthology relations to facilitate direct cross-species comparisons. We show that these aligned network embeddings ensure consistency across species without sacrificing quality compared to species-specific network embeddings. We also show that the aligned network embeddings are complementary to sequence embedding techniques, despite the use of seqeuence-based orthology relations in the alignment process. Finally, we demonstrate the utility and quality of the embeddings by using them for two well-established tasks: subcellular localization prediction and protein function prediction. Training logistic regression classifiers on aligned network embeddings and sequence embeddings improved the accuracy over using sequence alone, reaching performance numbers close to state-of-the-art deep-learning methods. A set of precomputed cross-species network embeddings and ProtT5 embeddings for all eukaryotic proteins have been included in STRING version 12.0.

## Introduction

Machine learning has long been used to enhance our understanding of proteins by making predictions about their characteristics [1], including functions, subcellular localizations, and structure. This works because machine learning can identify complex patterns that would be hard to otherwise find, for example in protein sequences. Through the decades, the growth in biological data combined with advances in machine learning techniques have considerably improved the accuracy of these predictions, helped elucidate the functions and localizations of proteins [2, 3], and been applied in the discovery of new drugs [4] and biomarkers [5].

Over the past several years, protein language models trained on vast amounts of protein sequences have revolutionized computational biology [6–9]. These models, such as ProtT5 [6] and ESM2 [9], trained on more than 45 million protein sequences from UniRef50 dataset [10], learn to represent protein sequences as high-dimensional vectors, encapsulating both protein domains and shorter motifs in the sequences. These protein embeddings serve as a foundation for further analysis and are often employed within smaller, labeled datasets to train supervised models. The impact of this approach has been broad; protein language models are now used by all state-of-the-art tools for predicting protein function [11], subcellular localization [12], and post-translational modifications [13].

While sequence embeddings have revolutionized protein sequence analysis by enabling precise predictions across various tasks, protein-protein interaction (PPI) networks [14] serve as a valuable complementary information source that captures the complex interplay between proteins, crucial for understanding biological processes and disease mechanisms [15, 16]. For instance, node2loc [17] demonstrated that PPI-based embeddings can perform better in subcellular localization prediction compared to sequence embeddings, and NetGO [18] shows that incorporating PPIs through nearest-neighbor searches can enhance function prediction accuracy. These studies highlight the potential of PPI networks as a complementary information source for various predictive tasks. An important source of PPI networks is the STRING database, which comprises protein physical interactions and functional associations for 12,535 species, including 1,322 eukaryotes [19]. These interactions are integrated from curated databases, literature mining, computational prediction, and orthology-based knowledge transfer between species.

Network-based protein embeddings can be typically generated using random-walk techniques such as deepwalk [20] and node2vec [21], or graph neural networks (GNNs) [22, 23]. While deepwalk and node2vec employ random walks to learn node representations, node2vec uses biased walks to balance local and global network exploration. Unlike GNNs, node2vec offers several advantages, particularly for unlabeled graphs like STRING. As an unsupervised method, node2vec does not require node features or labeled data. Additionally, node2vec generally exhibits better scalability to large graphs and is often more computationally efficient than many GNN architectures [24]. Moreover, as demonstrated by [25], the weighted version of node2vec, which considers edge weights while performing random walks, outperforms GNNs specifically on weighted, undirected, and unlabeled networks from STRING.

However, when node2vec is applied separately to multiple networks, e.g. PPI networks from different species, the resulting embeddings are not directly comparable. Consequently, proteins with similar characteristics across networks may have completely different embeddings. This hinders the use of network protein embeddings in tasks that span multiple species and prevents knowledge transfer between species [26, 27]. To deal with this challenge, researchers have turned to embedding alignment techniques as a potential solution. Embedding alignment, as defined by [28], is the task of finding a mapping between two vector spaces representing embeddings of different datasets, enabling tasks such as cross-lingual translation [29–31] and knowledge graph integration [32–34].

To create cross-species network protein embeddings, past work has used orthologs as anchors through several approaches: network kernels for pairwise directed alignment [35], autoencoders for pairwise bidirectional alignment [36], and node2vec applied to multi-species networks including ortholog relationships as edges [37]. By aligning the protein networks of diverse species, these approaches enhanced the precision and robustness of cross-species protein predictions, marking a substantial improvement over traditional single-species network embeddings [35–37]. FedCoder [38] is an innovative approach to integrating multiple knowledge graph (KG) embedding spaces into a uniform “Web of Embeddings”. In FedCoder, autoencoders are applied to learn mappings for embeddings from individual KGs that project into a shared latent space. In the latent space of the autoencoders, linked entities from different KGs have similar representations, effectively aligning the original embedding spaces. A key advantage of FedCoder for aligning multiple networks is its scalability, exhibiting linear computational complexity with respect to the number of embedding spaces, in contrast to pairwise methods that scale quadratically. Additionally, the performance of FedCoder improves as more embedding spaces are integrated.

In this study, we introduce SPACE, a comprehensive set of embeddings for all eukaryotic proteins in the STRING database. SPACE includes pre-calculated aligned cross-species network embeddings generated with a modified version of FedCoder (see Methods) [38] as well as protein sequence embeddings from the ProtT5 model. We show that the network embeddings are well-aligned across eukaryotes without sacrificing quality in the individual species and are complementary to sequence embeddings. We further demonstrate the effectiveness of both types of embeddings by applying them to two cross-species prediction tasks, namely subcellular localization prediction and protein function prediction. To facilitate widespread use of SPACE as a foundation for prediction tasks, we make the full set of pre-computed network and sequence embeddings available from the STRING website (https://string-db.org/).

## Results & Discussion

### SPACE: Pre-calculated sequence and cross-species network embeddings

In this study, we present SPACE (STRING Proteins as Complementary Embeddings), a comprehensive set of protein embeddings for all eukaryotic proteins in the STRING database. SPACE includes pre-calculated aligned cross-species network embeddings generated with a modified version of FedCoder (see Methods) as well as 1024-dimensional protein sequence embeddings from the ProtT5 model (Fig. 1a). This approach leverages protein networks and orthology relations for 1,322 eukaryotes from the STRING database to create network-based cross-species protein embeddings. The SPACE workflow begins by creating 128-dimensional species-specific network embeddings using node2vec, which captures the information from protein-protein interaction networks within each species. These embeddings are then aligned across species using a two-step process. First, embeddings from 48 selected seed species are aligned using the FedCoder method. This process employs per-species autoencoders to decrease the distance between orthologs in the latent space while preserving the original network information for each species, resulting in 512-dimensional cross-species embeddings. Subsequently, using a similar encoder–decoder architecture, the embeddings for non-seed species are aligned to the established latent space of their corresponding taxonomic groups (fungi, plants, animals, or protists).

**Figure 1.**
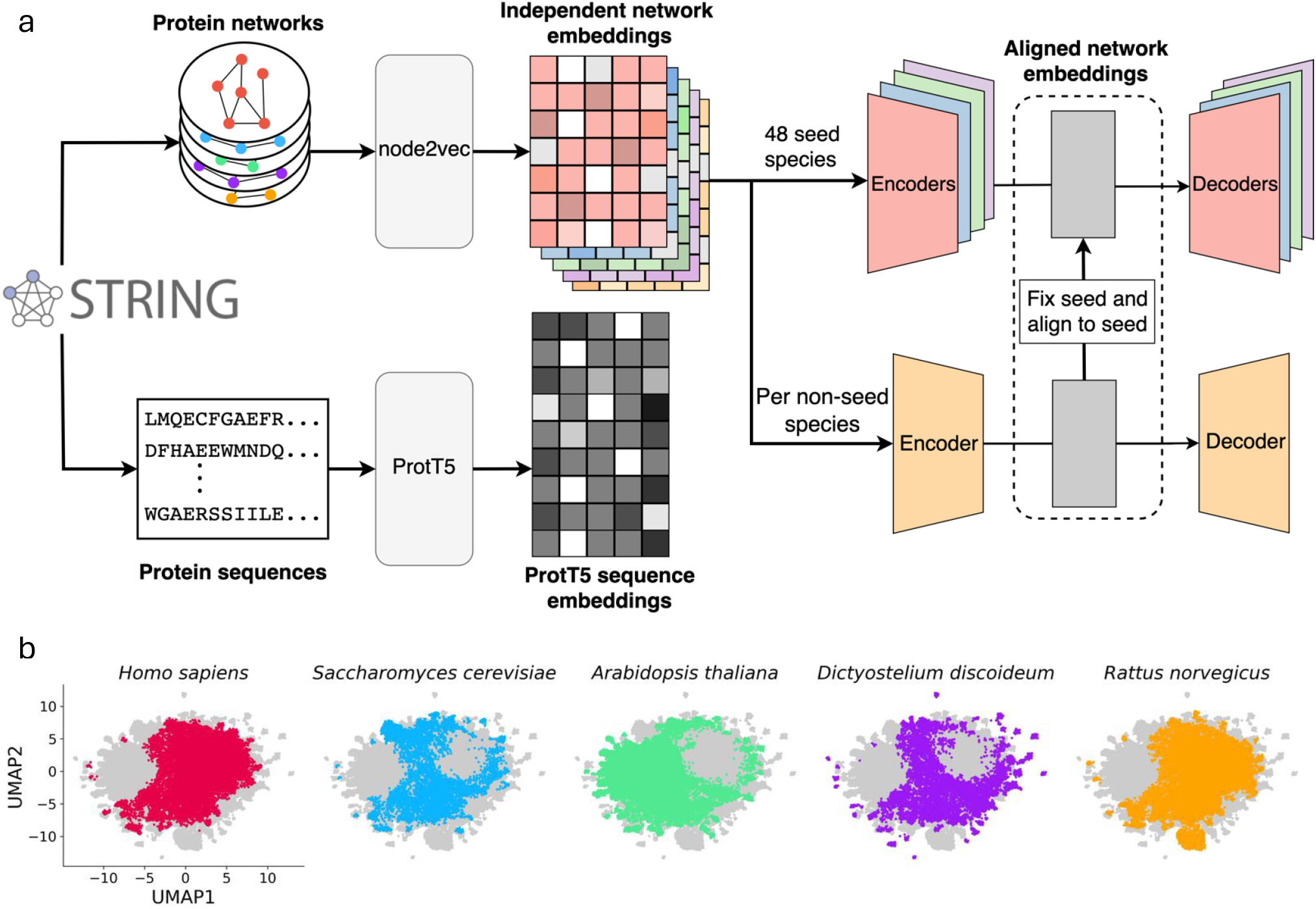
SPACE workflow and demonstration of successful cross-species embedding alignment a: Overview of the SPACE workflow. The pipeline begins with input from the STRING database in two forms: protein-protein interaction networks and protein sequences. The networks are processed through node2vec to generate 128-dimensional species-specific embeddings. The network alignment process first aligns 48 seed species using the FedCoder method to create a 512-dimensional shared latent space, then aligns each remaining non-seed species to their corresponding taxonomic groups (fungi, plants, animals, or protists) in this established latent space using autoencoders. In parallel, sequences are processed through the ProtT5 encoder to generate sequence embeddings. **b**: UMAP visualization demonstrates cross-species embedding alignment’s effectiveness. The plots show aligned network protein embeddings for four evolutionarily diverse seed species (*H. sapiens, S. cerevisiae, A. thaliana*, and *D. discoideum*) and one non-seed species (*R. norvegicus*). Colored points represent proteins from the named species, while gray points show the background distribution of proteins from other species. The overlapping patterns in the embeddings demonstrate successful alignment, with some regions representing functional associations found throughout eukaryotes and others representing functions specific to particular kingdoms. The unmapped cluster from *R. norvegicus* is mainly composed by olfactory proteins.

To assess the quality of our aligned embeddings, we created UMAP [39] visualizations of four evolutionarily distant seed species (*Saccharomyces cerevisiae, Arabidopsis thaliana, Homo sapiens, Dictyostelium discoideum*), and one non-seed species (*Rattus norvegicus*). Fig. 1b shows that the four seed species occupy overlapping parts of the same latent space, with some parts representing biological processes found through-out eukaryotes and others representing processes specific to, e.g. animals or plants. *S. cerevisiae* and *D. discoideum* exhibit remarkably similar embedding patterns. The non-seed species *R. norvegicus* also shows a very similar pattern to the mammalian seed species (*H. sapiens*), with one notable exception: a distinct small cluster unique to *R. norvegicus*. Upon manual examination of this cluster, we discovered that 1,092 of its 1,322 proteins (83%) are olfactory receptors. This finding aligns with previous research by [40], who demonstrated that *R. norvegicus* possesses 1,201 olfactory receptor genes compared to just 388 in *H. sapiens*.

### SPACE embeddings maintain pathway integrity

We benchmarked the SPACE embeddings of 378 eukaryotic species against pathways from the KEGG database [41] to evaluate whether the embeddings capture pathway information and to compare the effectiveness of network-based and sequence embeddings. Due to space constraints, we manually selected 12 seed species covering a wide range in the phylogenetic tree, as shown in Figure 2. Detailed figures for all other species are available in the Supplementary files.

**Figure 2.**
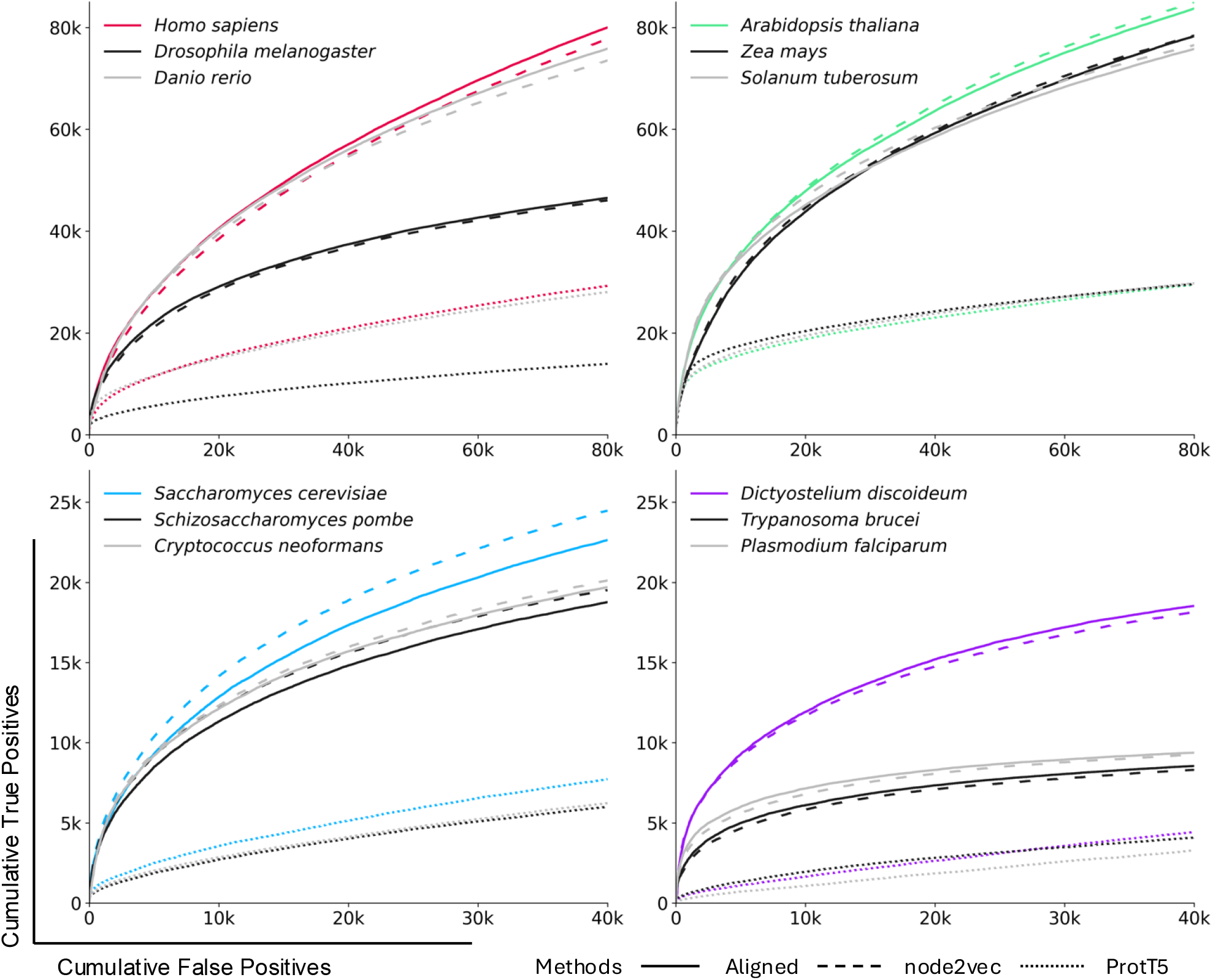
Comparison of protein embedding methods across diverse eukaryotic species using KEGG pathways. The plots show receiver operating characteristic (ROC) curves comparing three different embedding approaches: aligned network embeddings (solid lines), node2vec embeddings (dashed lines), and ProtT5 sequence embeddings (dotted lines). Results are presented for 12 representative species divided into four panels: **(top left)** animal species including *Homo sapiens, Drosophila melanogaster*, and *Danio rerio*; **(top right)** plants including *Arabidopsis thaliana, Zea mays*, and *Solanum tuberosum*; **(bottom left)** fungi including *Saccharomyces cerevisiae, Schizosaccharomyces pombe*, and *Cryptococcus neoformans*; and **(bottom right)** protists including *Dictyostelium discoideum, Trypanosoma brucei*, and *Plasmodium falciparum*. For each species, all possible protein pairs where both proteins are annotated in KEGG pathways were evaluated and ranked by their cosine similarity scores. The x-axis shows cumulative false positives, while the y-axis shows cumulative true positives. True positives are defined as protein pairs sharing at least one KEGG pathway, while false positives are pairs without shared pathways. The curves demonstrate that aligned embeddings maintain pathway information comparable to original node2vec embeddings across most species, while both network-based methods consistently outperform sequence embeddings in capturing pathway relationships.

These curves show that the aligned embeddings (solid lines) capture pathway information as accurately as the node2vec embeddings (dashed lines) that they are based on. The aligned embeddings slightly outper-form node2vec for some species (e.g. *H. sapiens*, and *D. discoideum*), whereas node2vec performs better for certain other species (e.g. *A. thaliana*). The only notable outlier is *S. cerevisiae*, which performs worse after alignment, which is likely explained by this species having one of the highest-quality networks in STRING and thus an unusually good node2vec embedding.

While both aligned and node2vec embeddings show varying agreement with pathways, sequence embeddings (dotted lines) derived from ProtT5 consistently underperform in the KEGG benchmark. This performance gap can be attributed to the nature of the information that sequence embeddings capture. Sequence embeddings are designed to encode protein structure, domains, and local motifs [6]. Conversely, they do not in-corporate information on interactions, which are crucial for understanding biological processes and pathways.

In summary, our alignment process succeeds in preserving the original information contained in the node2vec embeddings while enabling cross-species comparability. This retention of species-specific network characteristics, combined with their support for cross-species comparisons, makes the aligned embeddings a valuable complement to sequence embeddings, with the former capturing the complex interplay of proteins and the latter providing information about protein structure and evolution.

### Combining embeddings improves cross-species subcellular localization prediction

To evaluate the usefulness of ProtT5 embeddings, aligned embeddings, and their concatenation (SPACE), we test them on downstream tasks. The first such task is subcellular localization prediction using the DeepLoc 2.0 [12] datasets (after id-mapping, detailed in Methods). We retained 24,816 out of 28,303 proteins, covering 144 species, from the Swiss-Prot cross-validation set, and 1,646 out of 1,717 proteins from the Human Protein Atlas (HPA) test set, which consists solely of human proteins. We trained logistic regression models to predict each localization from the embeddings. The predictive performance is shown as precision-recall curves (Fig. 3a,b), and standard performance metrics are provided in Sup. Tab. 4 and Sup. Tab. 5.

**Figure 3.**
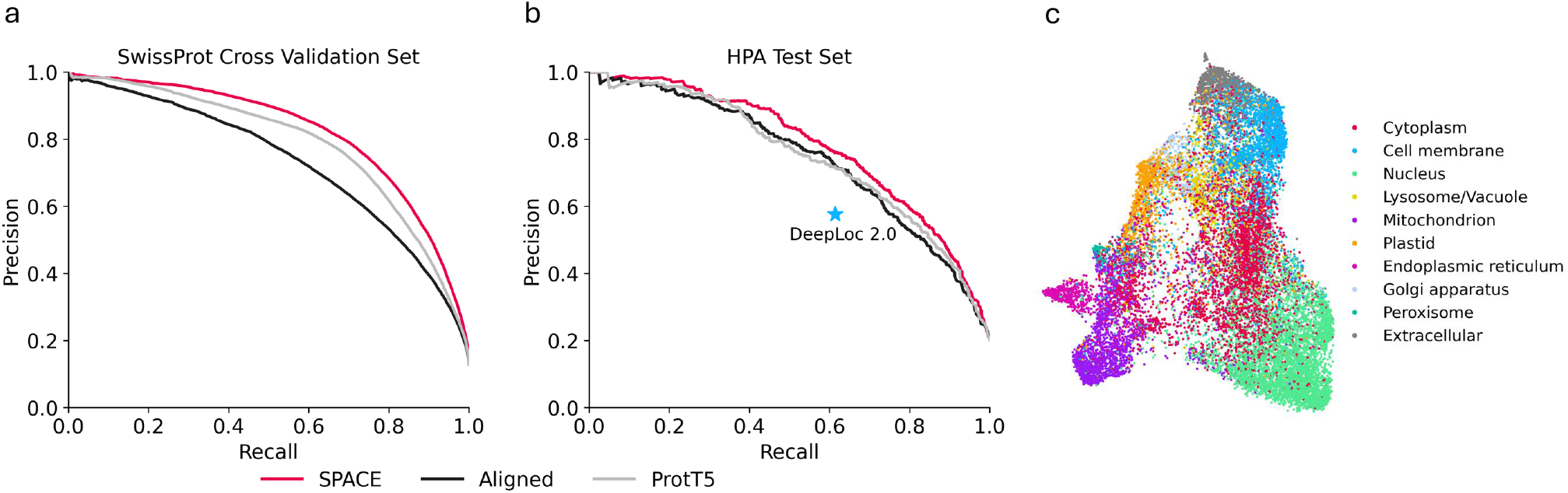
Precision-recall curves of different embeddings and visualization of SPACE embeddings in protein subcellular localization prediction. **a:** Precision-recall curves on SwissProt cross-validation set (24,816 proteins across 144 species) comparing SPACE (concatenation of aligned network and ProtT5 sequence embeddings, red), aligned network embeddings (black), and ProtT5 sequence embeddings (gray). **b:** Precision-recall curves on Human Protein Atlas (HPA) test set (1,646 human proteins), with DeepLoc2 predictions (blue star) included as an additional baseline. The curves demonstrate that SPACE embeddings consistently maintain higher precision across all recall values compared to individual embedding types. **c:** UMAP visualization of aligned network embeddings based on their projections onto logistic regression weight vectors for subcellular localization prediction. The distinct clustering patterns demonstrate that the aligned embeddings successfully capture protein localization information across multiple species, with clear separation observed for major cellular compartments such as nucleus, mitochondrion, and cell membrane. Proteins with multiple localizations were excluded from this visualization to ensure clear compartment separation.

We examined precision-recall curves on both SwissProt cross-validation and HPA test sets (Fig. 3a). On the SwissProt cross-validation set, SPACE embeddings consistently maintain higher precision across all recall values compared to both aligned network embeddings and ProtT5 embeddings (Fig. 3a). This advantage is particularly pronounced from a recall of 0.25 and up, where SPACE can make more predictions with the same precision. The performance difference between SPACE and ProtT5 is further confirmed on the HPA test set (Fig. 3b). Moreover, all the embedding-based predictors outperform DeepLoc 2.0 on the HPA test set (a direct comparison cannot be made for the SwissProt dataset). These results are also reflected in standard performance metrics (Sup. Tab. 4 and Sup. Tab. 5).

To explore the performance of the embedding-based predictors in more detail, we looked at the Matthews Correlation Coefficients (MCCs) for individual localizations (Sup. Tab. 4 and Sup. Tab. 5). Aligned and SPACE embeddings allow for better predictions for some localizations that are challenging for sequence embeddings. For example, the targeting mechanisms for lysosome/vacuole, Golgi apparatus, and peroxisome are complex and make sequence-based prediction challenging. In lysosomes, proteins require both signal sequences and subsequent M6P modifications in the Golgi apparatus but share trafficking machinery and hydrolase pathways[42]; Golgi resident proteins have specific targeting/retention mechanisms [43], and proteins passing through the Golgi apparatus have various targeting signals for final destinations, but they participate in conserved glycosylation and sorting complexes [44]; peroxisomal proteins use two different targeting signals (PTS1/PTS2) but engage in metabolic pathways with shared import machinery (PEX proteins) [45]. Despite this complexity, these organelles have distinct functions, which are captured by the STRING network. This explains why network-based embeddings perform better than sequence embeddings alone for these organelles, highlighting how functional context can be more informative than sequence features for predicting localization.

We further visualized the aligned embeddings of the SwissProt dataset, excluding proteins with multiple locations, with UMAP to demonstrate the biological relevance of our aligned embeddings (Fig. 3c). To obtain a UMAP that best shows the information relevant to subcellular localization, we used the projection of the aligned embeddings on the set of weight vectors from the logistic regressions as input. This representation shows that proteins cluster according to their cellular compartments. The distinct grouping of proteins from certain subcellular locations, such as nucleus, mitochondrion, and cell membrane, shows that SPACE embeddings successfully capture protein localization across multiple species.

### Combining embeddings enhances cross-species protein function prediction

Our second downstream task is protein function prediction using the NetGO 2.0 [46] datasets. We mapped 74,838 proteins across 152 eukaryotic species from the training set of 120,856 prokaryotic and eukaryotic proteins, with an additional 15,827 proteins from 12 species serving from the test set (detailed in Methods). We trained logistic regression models to predict each Gene Ontology [47, 48] (GO) term from the embeddings. The predictive performance over the test set is shown as precision-recall curves (Fig. 4a,b,c), and standard performance metrics are provided in Sup. Tab. 6.

**Figure 4.**
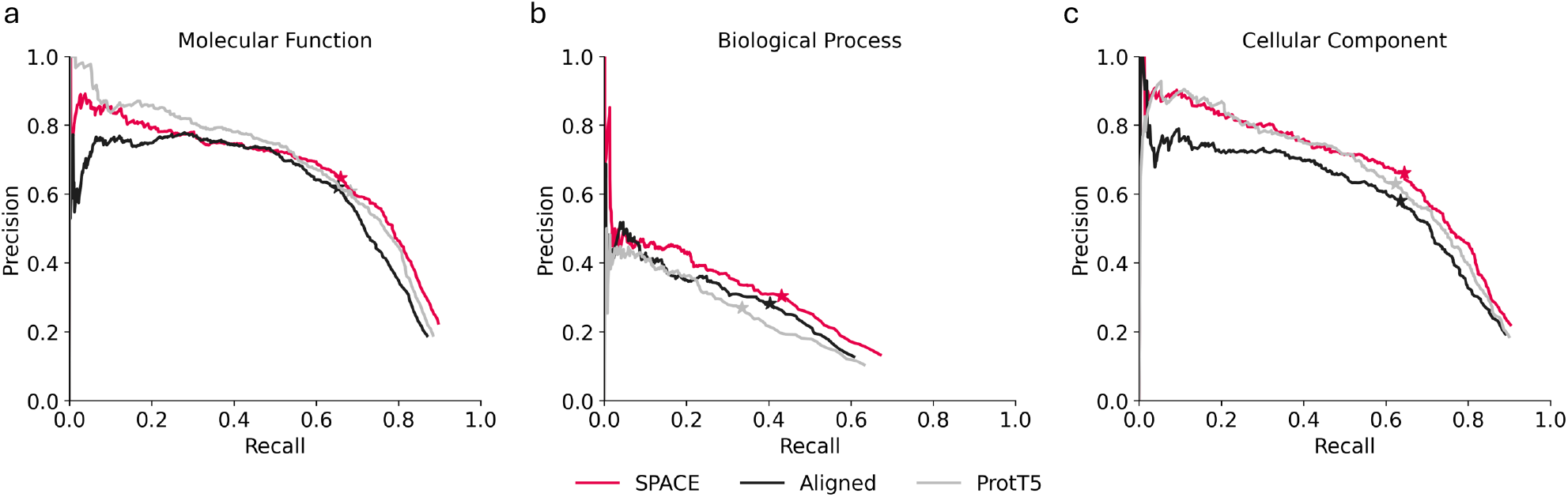
Precision-recall curves of different embeddings in protein function prediction. **a:** Molecular Function, **b:** Biological Process, and **c:** Cellular Component. Each panel compares SPACE (concatenation of aligned network and ProtT5 sequence embeddings, red), aligned network embeddings (black), and ProtT5 sequence embeddings (gray). Stars indicate precision and recall values that yield maximum MicroF1 scores. The curves reveal that SPACE embeddings show particular strength in biological process prediction, while maintaining competitive performance in molecular function and cellular component prediction, highlighting the complementary nature of sequence and network information in capturing different aspects of protein function.

**Table 1.** node2vec Hyperparameter Tuning Space.

| Hyperparameter | Search Space | Selection |
| --- | --- | --- |
| Dimensionality | 32, 64, 128, 256, 512 | 128 |
| Length of walk per source | 20, 50, 100 | 50 |
| Number of walks per source | 10, 20, 30 | 10 |
| Window size | 5, 10 | 5 |
| Epochs | 5, 10, 20 | 5 |
| Return and in-out hyperparameters | (0.1, 0.9), (0.3, 0.7), (0.7, 0.3), (0.9, 0.1) | (0.3,0.7) |

**Table 2.**
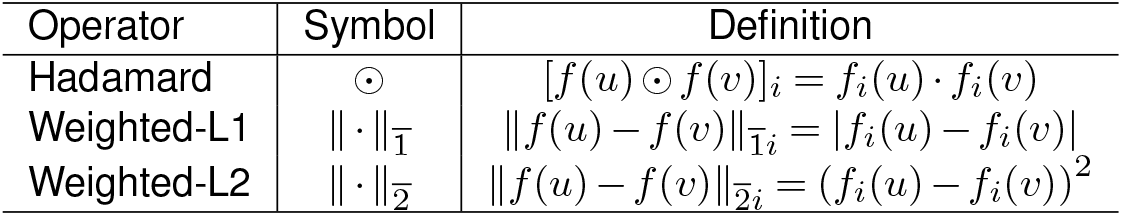
Operators for generating edge features in link prediction tasks. The table lists the operators used, their corresponding symbols, and their definitions. The Hadamard operator computes the element-wise product of two feature vectors. The Weighted-L1 and Weighted-L2 operators compute the L1 and L2 norms, respectively, weighted by the differences between corresponding feature vector elements.

**Table 3.** Seed Species Alignment Hyperparameter Tuning Space.

| Hyperparameter | Search Space | Selection |
| --- | --- | --- |
| Distance function | Cosine distance, Euclidean distance | Euclidean distance |
| Margin | 1, 0.1, 0.01 | 0.1 |
| Balance hyperparameter | 0.1, 0.2, 0.3, 0.4, 0.5 | 0.5 |
| Latent dimensionality | 128, 256, 512, 1024 | 512 |

**Table 4.** Protein subcellular localization prediction on DeepLoc 2.0 SwissProt cross-validation dataset. The best scores for each metric are shown in bold. SPACE: the concatenation of ProtT5 and aligned network embeddings. A logistic regression model was trained on each location with the corresponding embeddings. The scores are reported in the mean score with standard deviation on 5 partitions.

|  |  | ProtT5 | Aligned | SPACE |
| --- | --- | --- | --- | --- |
| Accuracy | | $0.53 \pm 0.01$ | $0.44 \pm 0.02$ | **$0.56 \pm 0.01$** |
| Jaccard | | $0.55 \pm 0.02$ | $0.47 \pm 0.02$ | **$0.58 \pm 0.02$** |
| MicroF1 | | $0.71 \pm 0.02$ | $0.64 \pm 0.02$ | **$0.74 \pm 0.01$** |
| MacroF1 | | $0.52 \pm 0.00$ | $0.54 \pm 0.01$ | **$0.65 \pm 0.01$** |
| MCC per location | count |  |  |  |
| Nucleus | 9135 | $0.70 \pm 0.01$ | $0.66 \pm 0.01$ | **$0.72 \pm 0.01$** |
| Cytoplasm | 9108 | $0.59 \pm 0.02$ | $0.43 \pm 0.02$ | **$0.60 \pm 0.01$** |
| Cell membrane | 3930 | $0.65 \pm 0.02$ | $0.54 \pm 0.02$ | **$0.66 \pm 0.01$** |
| Mitochondrion | 2451 | $0.73 \pm 0.02$ | $0.67 \pm 0.02$ | **$0.77 \pm 0.02$** |
| Endoplasmic reticulum | 2021 | $0.45 \pm 0.03$ | $0.47 \pm 0.04$ | **$0.56 \pm 0.04$** |
| Extracellular | 1563 | $0.74 \pm 0.05$ | $0.53 \pm 0.03$ | **$0.75 \pm 0.03$** |
| Lysosome/Vacuole | 1424 | $0.07 \pm 0.06$ | $0.37 \pm 0.03$ | **$0.38 \pm 0.03$** |
| Golgi apparatus | 1209 | $0.27 \pm 0.06$ | $0.36 \pm 0.06$ | **$0.41 \pm 0.07$** |
| Plastid | 926 | $0.87 \pm 0.01$ | $0.67 \pm 0.03$ | **$0.88 \pm 0.01$** |
| Peroxisome | 272 | $0.06 \pm 0.09$ | $0.46 \pm 0.08$ | **$0.52 \pm 0.04$** |

**Table 5.** Protein subcellular localization prediction on DeepLoc 2.0 HPA test set. The best scores for each metric are shown in bold. SPACE is the concatenation of ProtT5 and aligned network embeddings. A logistic regression model was trained on each location with the corresponding embeddings and cross-validation dataset.

|  |  | DeepLoc 2.0 | ProtT5 | Aligned | SPACE |
| --- | --- | --- | --- | --- | --- |
| Accuracy |  | 0.38 | 0.51 | 0.48 | **0.53** |
| Jaccard |  | 0.42 | 0.50 | 0.50 | **0.53** |
| Micro F1 |  | 0.59 | 0.66 | 0.66 | **0.70** |
| Macro F1 |  | 0.45 | 0.48 | 0.56 | **0.61** |
| MCC per location | count |  |  |  |  |
| Nucleus | 865 | 0.44 | 0.58 | 0.54 | **0.60** |
| Cytoplasm | 518 | 0.34 | **0.47** | 0.46 | 0.46 |
| Cell membrane | 271 | 0.35 | 0.46 | 0.53 | **0.57** |
| Mitochondrion | 183 | 0.47 | 0.71 | 0.72 | **0.77** |
| Golgi apparatus | 85 | 0.24 | 0.16 | 0.31 | **0.44** |
| Endoplasmic reticulum | 74 | 0.20 | 0.19 | 0.41 | **0.45** |

**Table 6.** Protein function prediction metrics on NetGO 2.0 test set. The best scores for each metric are shown in bold. SPACE is the concatenation of ProtT5 and aligned network embeddings. A logistic regression model was trained on each GO term with the corresponding embeddings.

| | $F_{max}$ | | | AUPRC | | | $S_{min}$ | | |
| --- | --- | --- | --- | --- | --- | --- | --- | --- | --- |
|  | MF | BP | CC | MF | BP | CC | MF | BP | CC |
| ProtT5 | 0.64 | 0.30 | 0.63 | **0.63** | 0.17 | 0.61 | 5.78 | **33.08** | 6.72 |
| Aligned | 0.64 | 0.33 | 0.61 | 0.56 | 0.19 | 0.55 | 5.37 | 36.81 | 7.44 |
| SPACE | **0.65** | **0.36** | **0.65** | 0.62 | **0.23** | **0.63** | **4.99** | 33.36 | **6.36** |

We examined precision-recall curves on each GO aspect (Fig. 4a,b,c). For molecular functions, ProtT5 sequence embeddings show slightly better performance in the mid-recall range, but SPACE embeddings are marginally better than them at the higher recall range. SPACE consistently outperforms aligned network embeddings and ProtT5 sequence embeddings across most recall values for biological processes. Finally, SPACE shows modest advantages at higher recall values (*>* 0.6) for cellular components.

The standard evaluation metrics (Sup. Tab. 6) confirm the patterns observed in the precision-recall curves. Across all three GO aspects, SPACE demonstrates robust performance, with similar performance to ProtT5 in both molecular function and cellular component prediction. SPACE particularly shows its strength in predicting biological processes.

The precision-recall curves and the standard metrics show that the molecular function predictions rely heavily on the sequence embeddings, presumably because they encode protein domain information. Meanwhile, biological process prediction benefits from integrating network embeddings that capture protein-protein interactions and pathway relationships.

## Conclusions

This research presents SPACE, a collection of complementary embeddings for all proteins from the 1,322 eukaryotic species in the STRING database. This consists of pre-calculated protein sequence embeddings using the ProtT5-XL-UniRef50 model and aligned, cross-species network embeddings created using node2vec and a modified version of FedCoder. The aligned network embeddings are designed to complement sequence-based embeddings. SPACE provides these pre-calculated sequence and aligned network embeddings with the aim to support a broad range of prediction tasks across species.

Our results show that the embeddings are directly comparable between proteins from different species, that the aligned network embeddings retain the information from species-specific embeddings, and that the sequence and network embeddings are complementary. We demonstrate the latter on two downstream tasks, namely subcellular localization and protein function prediction. The performance, which can be achieved on these tasks using simple logistic regression, demonstrates the power of the embeddings.

The practical applications of SPACE embeddings are vast, extending beyond basic research to potential clinical implications. For example, cross-species embeddings can be the starting point for training a general predictor of protein interactions among eukaryotic parasites and their hosts. The comparability across species can also improve large-scale functional annotations, especially for less-studied organisms.

In conclusion, our research highlights the potential of combining sequence and cross-species network-based embeddings. We believe that SPACE provides easy access to embeddings that can serve as a foundation for protein-related prediction tasks.

## Methods

### Dataset

The datasets used to generate SPACE include protein sequences, protein functional association networks, and orthologs. The protein sequences and functional association networks are sourced from STRING version 12.0 [19], while the orthologs are from eggNOG 6.0 [49]. The STRING database provides functional association networks for 1,322 eukaryotic species, with undirected networks and edge weights ranging from 0.15 to 1.00. We used the combined score channel of the full functional association networks. Orthologs, which serve as alignment anchors, are crucial for making network embeddings comparable across different species. In the eggNOG database, orthologs are organized based on the taxonomy levels and orthologous groups (OGs), with proteins within an OG having evolved from the same ancestral protein at the taxonomic level in question. We construct pairs of orthologs for any two species based on the OGs at their latest common ancestor taxonomy level. For instance, if within an OG species *A* and *B* have *i* and *j* unique proteins, respectively, they will result in *i × j* number of orthologous pairs.

In the network alignment process, we start with 48 seed species carefully selected to include a diverse range of well-studied organisms. Species were evaluated by comparing the STRING functional association networks, excluding evidence from the curated database channel and evidence transferred by orthology, with pathways from the KEGG database [41]. All the other eukaryotic species are defined as non-seed species.

The 48 seed species were selected for hyperparameter tuning of both the individual network embedding (using node2vec) and the network embedding alignment. For node2vec, the edges of each network were randomly divided into 6 partitions. Self-loops were added to singleton proteins that were not connected to any other proteins in the training set. The first five partitions were used for cross-validation, while the sixth served as a test set. For the network embedding alignment, proteins from the networks were partitioned into 6 groups, ensuring that all proteins from the same OGs were assigned to the same partition. Similarly, the first five partitions were utilized for cross-validation, with the sixth partition reserved as a test set.

### ProtT5 for protein sequence embeddings

The encoder-only ProtT5-XL-UniRef50 [6], Rostlab/prot_t5_xl_half_uniref50-enc (https://huggingface.co/Rostlab/prot_t5_xl_half_uniref50-enc), generated the 1024-dimensional protein sequence embeddings. ProtT5-XL-UniRef50 is a self-supervised protein language model built upon the T5-3B [50] model, trained on the UniRef50 [10] dataset, which contains more than 45 million protein sequences. We selected the half-precision and encoder-only version model, based on the original publication [6] and other downstream tasks [51–53]. The ProtT5 model produces embeddings of each amino acid in a protein sequence, and we average the amino acid embeddings to get the embedding of a protein. The full-length sequences of all proteins were fed into the model, using CPU mode for the less than 1000 proteins that could not fit in GPU memory.

### node2vec for single species network embedding

The node2vec algorithm [21] was used to create network-based protein embeddings for each species. node2vec is a random-walk-based algorithm [54, 55] that efficiently maps nodes in a graph to a low-dimensional space of features. We used a weighted version of node2vec, which considers the edge weights in the networks to create 128-dimensional embeddings that capture the network’s topological information, encoding both local and global structural properties. A grid search of predefined hyperparameter search spaces (Sup. Tab. 1) was carried out, and the results were expressed as mean and standard deviation link prediction (Sup. Tab. 2) scores over the cross-validation set.

The hyperparameters producing the best link prediction scores across all seed species were selected and used for all the species. The node2vec implementation of PecanPy (https://pypi.org/project/pecanpy/) was used throughout our work.

### FedCoder for multiple networks embedding alignment

To make FedCoder[38] suitable for our task, we made two changes to the method. As many-to-many orthology relations are common, we introduced a weighting mechanism (Eq. 1). For each pair of orthologous proteins *i* and *j*, we defines the weighting factor *W*_*ij*_ used in the alignment process as:

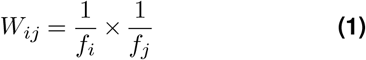

where *f*_*i*_ and *f*_*j*_ represent the occurrence frequencies of protein *i* and protein *j* among the orthologous groups of two species.

To reduce the complexity of training and improve the data quality, we excluded pairs with weights smaller than a threshold value, which we treated as a hyperparameter. Additionally, unlike the original loss function in FedCoder, we added another hyperparameter *α*, ranging from 0 to 1, to balance the alignment and reconstruction. Eq. 2 shows the overall loss function after this modification. It consists of two parts: the alignment loss ℒ^*I*^ and the reconstruction loss ℒ^*A*^, summing up the reconstruction loss of overall *N* species.

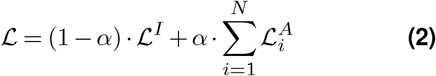

### Seed species alignment

We started the alignment with 48 seed species for two main reasons: (1) computational scalability, as it is computationally expensive to load all 1,322 eukaryotic species and their orthologous relationships into a single alignment process; and (2) study bias, as aligning understudied species with well-studied species can degrade information and add noise, given that many model organisms are well-studied with dense networks, while many other species in STRING are understudied.

We tuned the hyperparameters using grid search (Sup. Tab. 3). The best hyperparameters were selected based on the average alignment loss with standard deviation. Consistent with the original FedCoder publication, an autoencoder with one layer and without activation functions performed the best. However, our experiments showed that omitting negative sampling during alignment further improved alignment quality. We obtained aligned seed species embeddings with 512 dimensions. The autoencoders were implemented with PyTorch (https://pytorch.org/), and the same for non-seed species.

### Incorporating non-seed species

To broaden our analysis, we incorporated non-seed species by aligning their embeddings to those of the previously aligned seed species. We found that aligning non-seed species to their corresponding taxonomic groups (fungi, plants, animals, and protists) yielded better results compared to aligning them with all seed species collectively. Despite not utilizing all seed species in the alignment process, the non-seed species effectively align with the entire seed species set due to the pre-established alignment among seed species. During the integration of non-seed species, we fixed embeddings of seed species and employed a separate autoencoder for each non-seed species. This approach aims to simultaneously (1) minimize the distance between orthologous pairs of seed and non-seed species in the latent space, and (2) reconstruct the input embeddings of non-seed species from the latent space using a decoder. This independent alignment strategy enabled parallel processing of multiple non-seed species and enhanced scalability. To ensure consistency, we retained the same hyperparameters used in the seed species alignment process.

### Singletons in the networks

Some proteins, known as singletons, do not interact with other proteins in our networks. However, their sequence information is still valuable and can be used to generate sequence-based embeddings. To ensure all proteins in STRING are represented in both sequence and network embeddings for downstream tasks, we provide network embeddings for singletons as well. We first scale the network embeddings of all species to the range [−0.99, 0.99]. We then assign network embeddings to singletons based on three categories: (1) Singletons in interaction orthologous groups, where at least one protein is part of any network: these are assigned the average of their interaction orthologs’ embeddings, with small random noise added to ensure uniqueness; (2) Singletons in singleton orthologous groups, where no proteins are part of any network, but they are orthologs to each other: random embeddings in the ranges of [−1.0, −0.99] and [0.99, 1.0] are generated for these groups. This ensures their embeddings are distinct from the scaled network embeddings. Singletons in the same singleton OGs receive these embeddings with added random noise; (3) Singletons without orthologs: individual embeddings are generated using the same method, ensuring each is unique and appropriately positioned in the embedding space.

### Benchmark datasets and methods

KEGG [41] pathways comprise a detailed collection of pathway maps, and it is crucial for understanding biological functions and interactions. A benchmark dataset that spans 378 eukaryotic species in STRING was assembled from KEGG. We generated the Receiver Operating Characteristic (ROC) curves for each species by ranking protein pairs based on their cosine similarities in different types of embeddings (node2vec, aligned, and ProtT5) to compare their ability to capture the pathway information. Protein pairs were classified as false positives (FPs) if they did not share any pathway affiliation, or as true positives (TPs) if they co-located in at least one KEGG pathway. Protein pairs where at least one member was absent from the KEGG database were excluded from TP/FP designation. These ROC curves plotted cumulative FPs on the x-axis against cumulative TPs on the y-axis.

Subcellular localization [56, 57] refers to identifying a given protein’s specific location(s) within a cell. This task is essential in cell biology and proteomics because the functionality of proteins is inherently tied to their locations within the cell. To evaluate the performance of our embeddings in predicting subcellular localization, we used the SwissProt cross-validation set and the Human Protein Atlas (HPA) test set from DeepLoc 2.0 [12], mapped to STRING identifiers using the UniProt ID mapping (https://www.uniprot.org/id-mapping) and the STRING human aliases file, respectively. The SwissProt cross-validation set contains multiple species, providing a comprehensive evaluation of the embeddings’ performance across different organisms, while the HPA test set focuses on human proteins. To measure the effectiveness of the embeddings, we compared ProtT5 embeddings, aligned embeddings, and the concatenation of aligned and sequence embeddings. We trained a simple logistic regression model for each localization task. Additionally, we compared the performance of DeepLoc 2.0 with our logistic regression models over the HPA subset by running the original DeepLoc 2.0 implementation. The performance was measured by precision-recall curves, accuracy, Jaccard score, MicroF1, MacroF1, and per-location Matthews Correlation Coefficient (MCC). All the metrics were calculated with scikit-learn (https://pypi.org/project/scikit-learn/).

Furthermore, we explored the capability of SPACE embeddings to predict Gene Ontology [47, 48] (GO) terms, a crucial benchmark for assessing the functional annotation accuracy of protein embeddings. GO term prediction encompasses three aspects: biological processes, cellular components, and molecular functions, offering a comprehensive lens to evaluate our embeddings’ biological relevance and applicability. We built subsets of the training and test data used in NetGO 2.0 [46], filtered through the UniProt ID mapping. The description of the processed datasets can be found in the Supplementary Material. We trained logistic regression models per GO term on the training set and evaluated the performance on the test set using three types of embeddings: ProtT5, aligned embeddings, and the concatenation of aligned embeddings and ProtT5 embeddings. We show precision–recall curves for our methods. We also used the same evaluation metrics mentioned in NetGO 2.0 [46]: maximum MicroF1 (Fmax), Area under the Precision-Recall curve (AUPRC), and minimum semantic distance (Smin). All the metrics were calculated with CAFA-evaluator (https://pypi.org/project/cafaeval/).

## Availability and implementation

The source code and scripts for generating the networkbased cross-species protein embeddings are available at https://github.com/deweihu96/SPACE. Precomputed network embeddings and sequence embeddings for all eukaryotic proteins are included in STRING version 12.0 (https://string-db.org).

## Acknowledgements

We acknowledge Deic, Denmark, for awarding this project access to the LUMI supercomputer, owned by the EuroHPC Joint Undertaking, hosted by CSC (Finland) and the LUMI consortium through Deic, Denmark, Deic-KU-L5-2023-004.

## Author contributions

L.J.J. and D.H. jointly developed the conceptual framework and scientific direction. D.H. performed all computational analyses. D.S. selected seed species and made all the embeddings available on the STRING database.

D.H. and L.J.J. drafted the manuscript, and all authors provided input and feedback and approved the final manuscript.

## Hyperparameter Tuning

### node2vec

We tuned the hyperparameters of node2vec in the following search spaces:

### Link Prediction

Link prediction is a binary classification task that predicts whether a given unseen link exists in a network. In our networks, edges with scores higher than 0.7 were sampled as positive edges, while negative edges (not in the networks) were sampled based only on the training set. We maintained a positive-to-negative edge ratio of 1:10. To convert the node embeddings into edge vectors, we used several operators, and a logistic regression model was trained for the link prediction of each species.

### Network Embedding Alignment

## Evalutation Metrics

