## Supplementary figures and images for "SPACE: STRING proteins as complementary embeddings"

### 3218.png

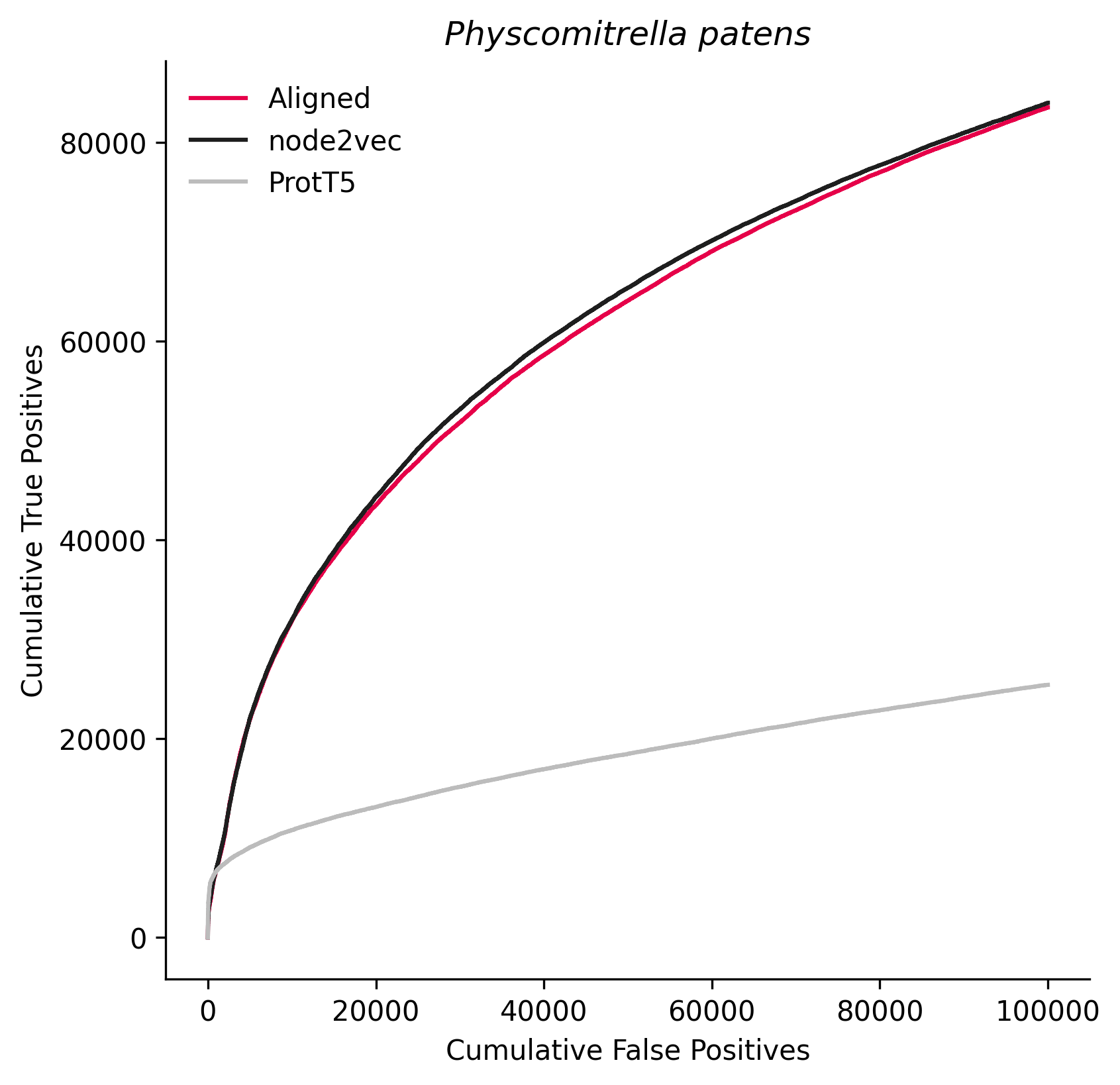

### 3641.png

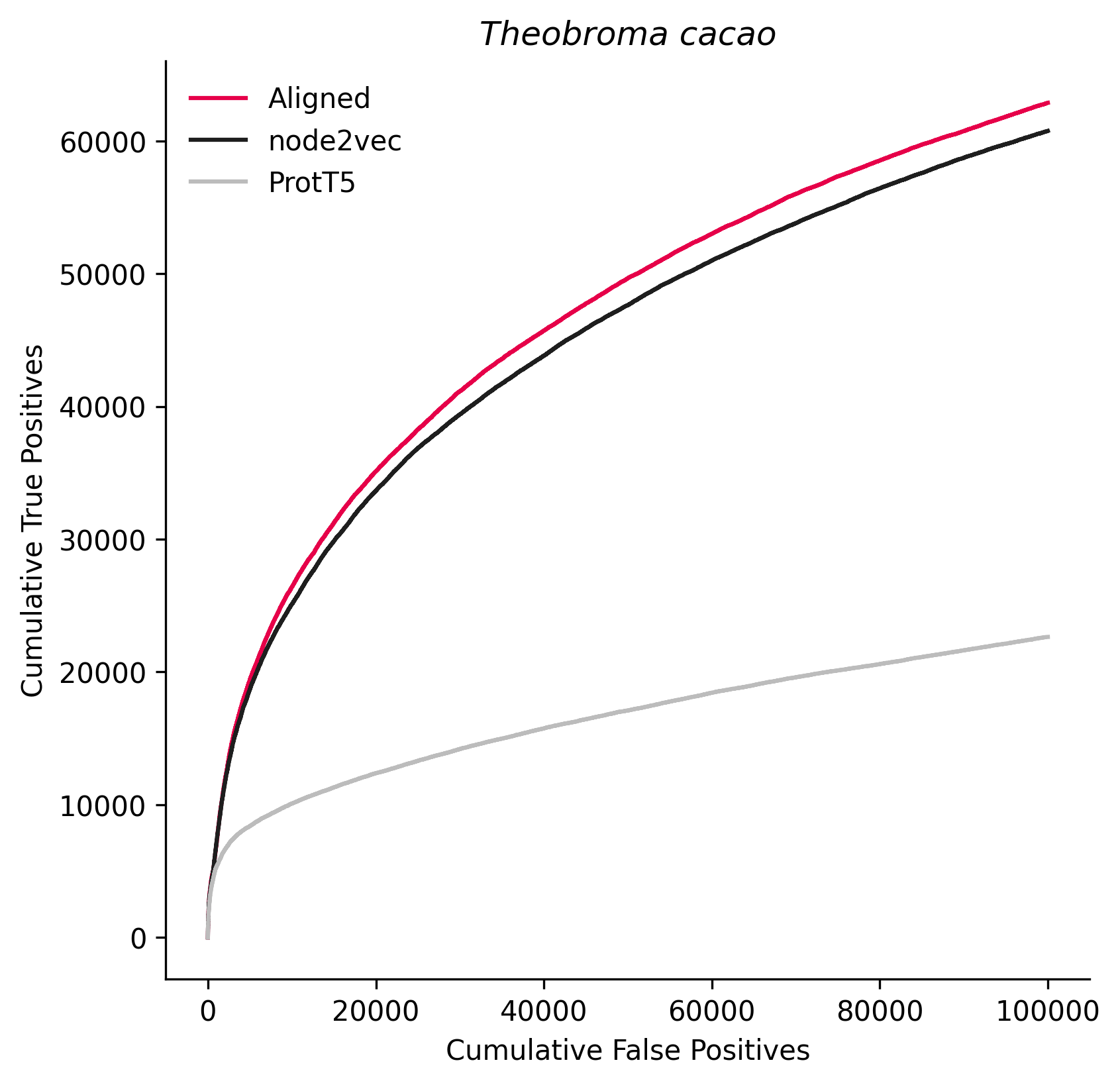

### 3659.png

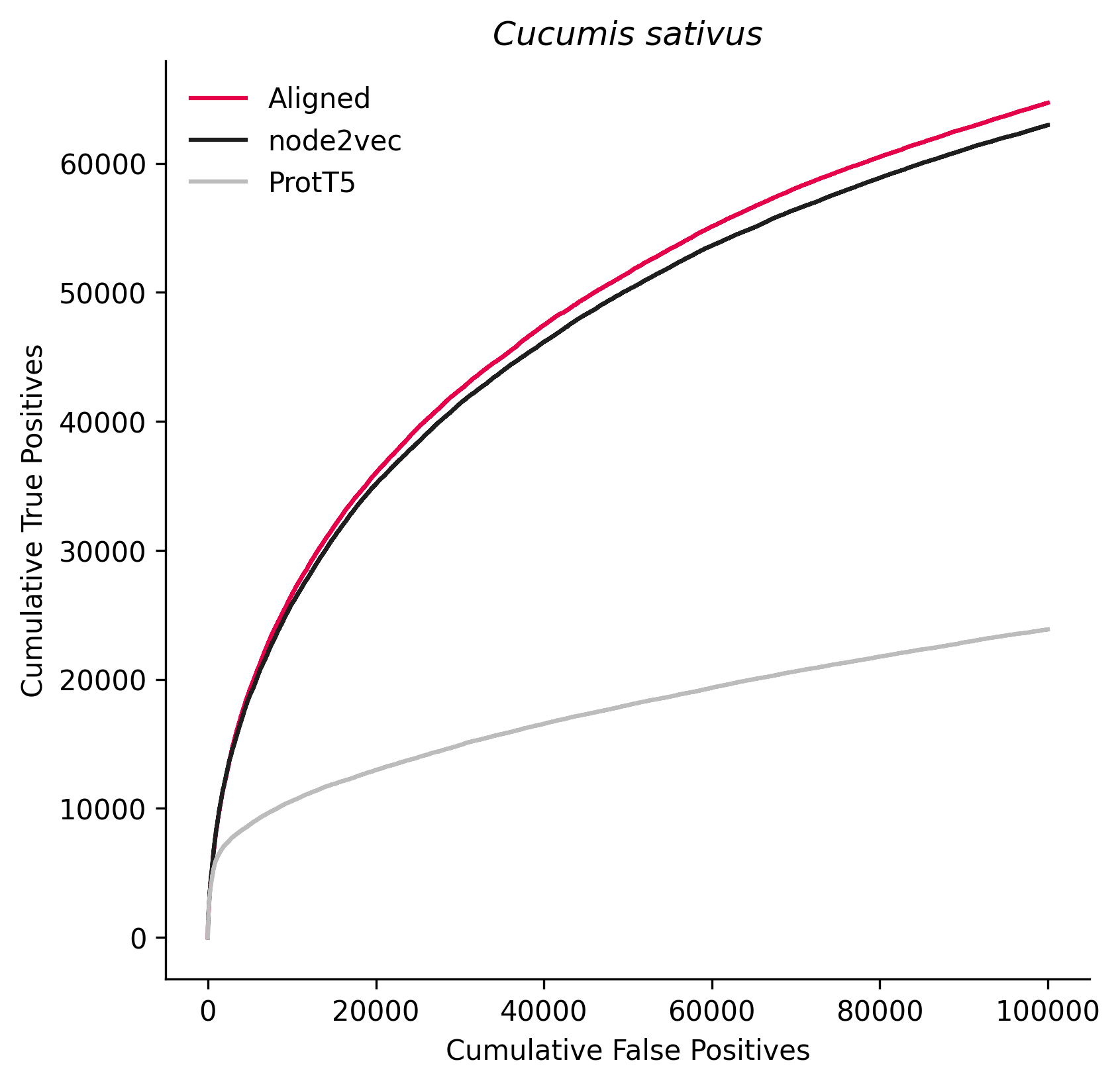

### 3750.png

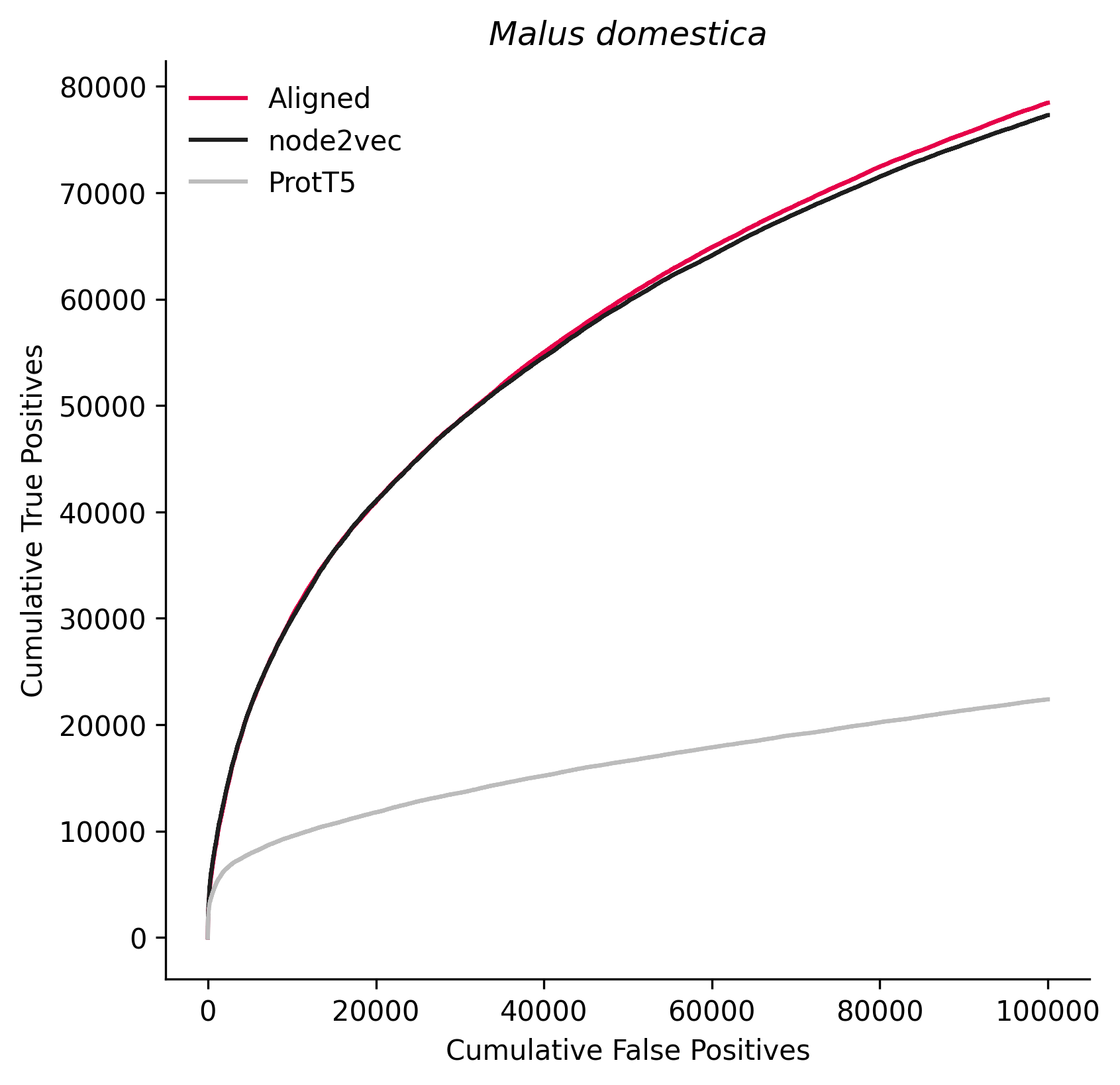

### 3821.png

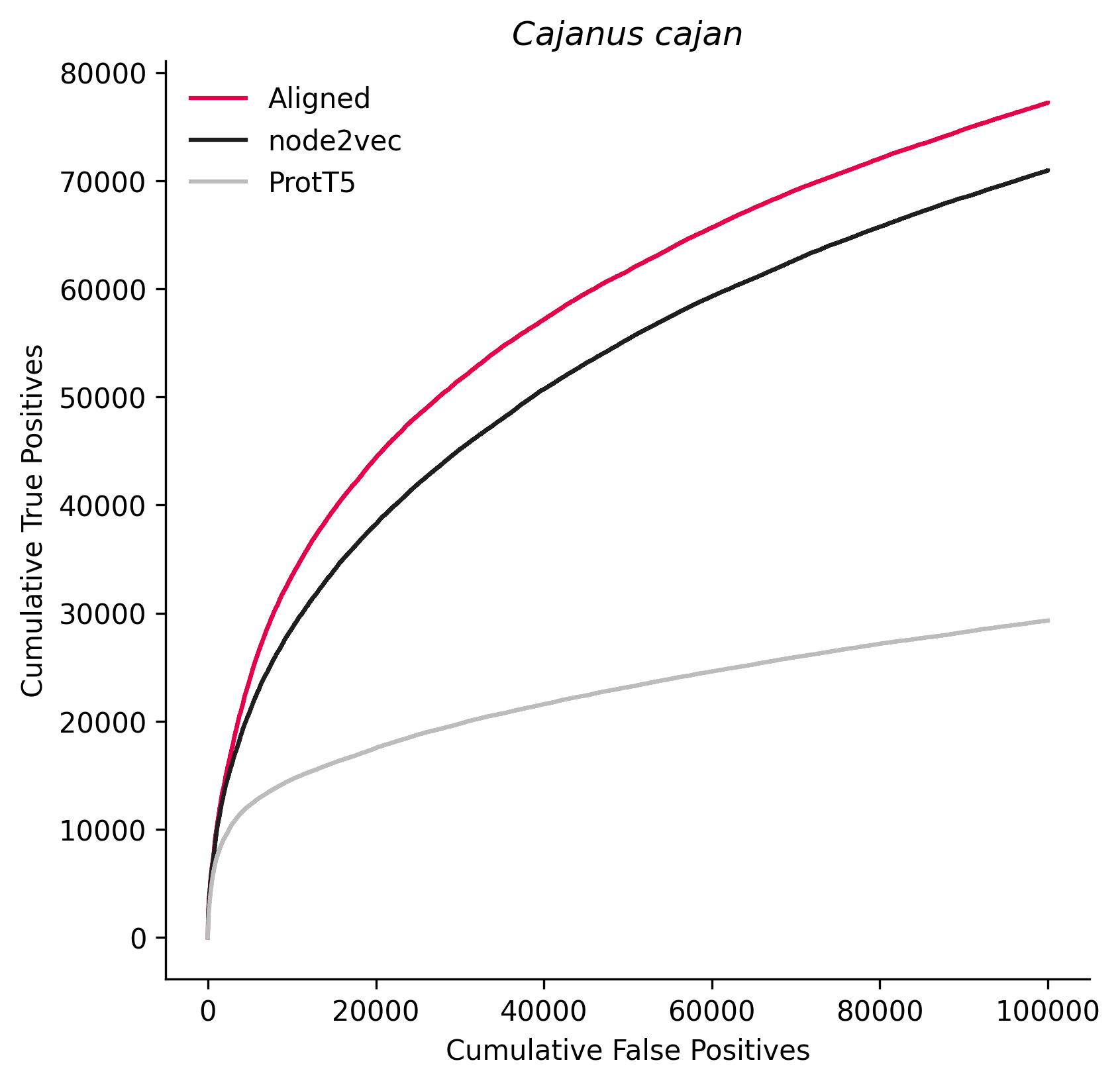

### 3880.png

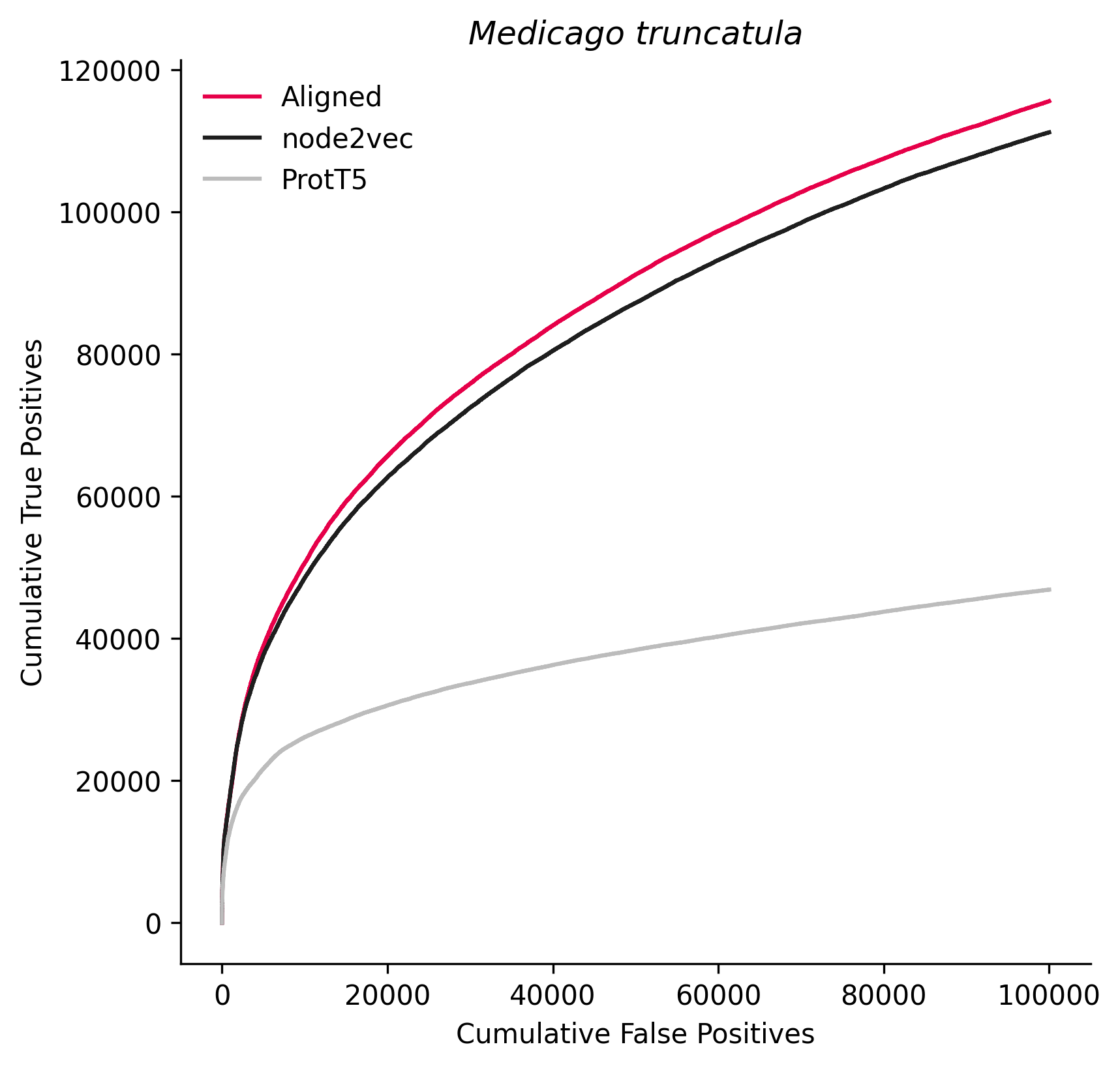

### 3914.png

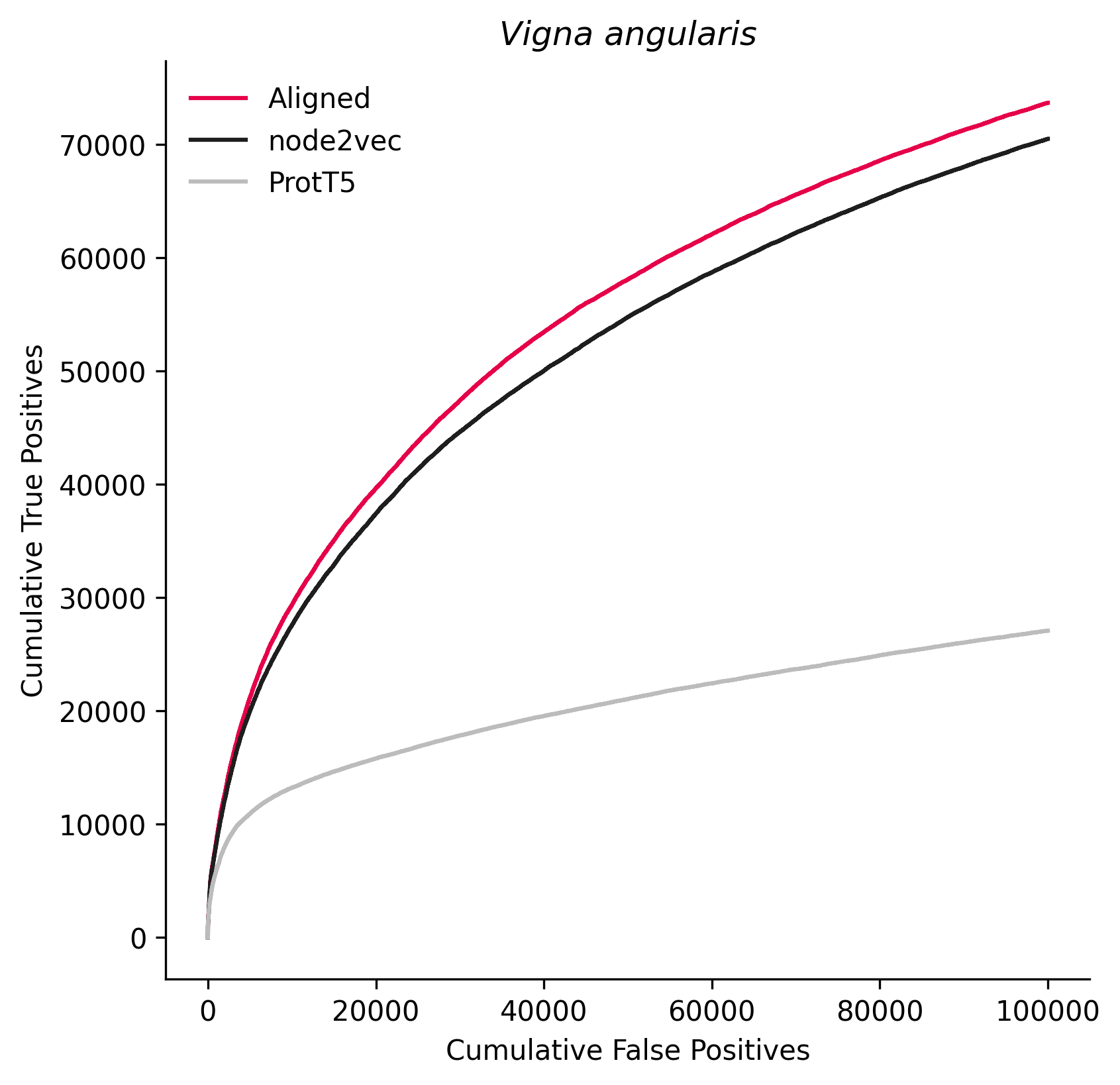

### 3916.png

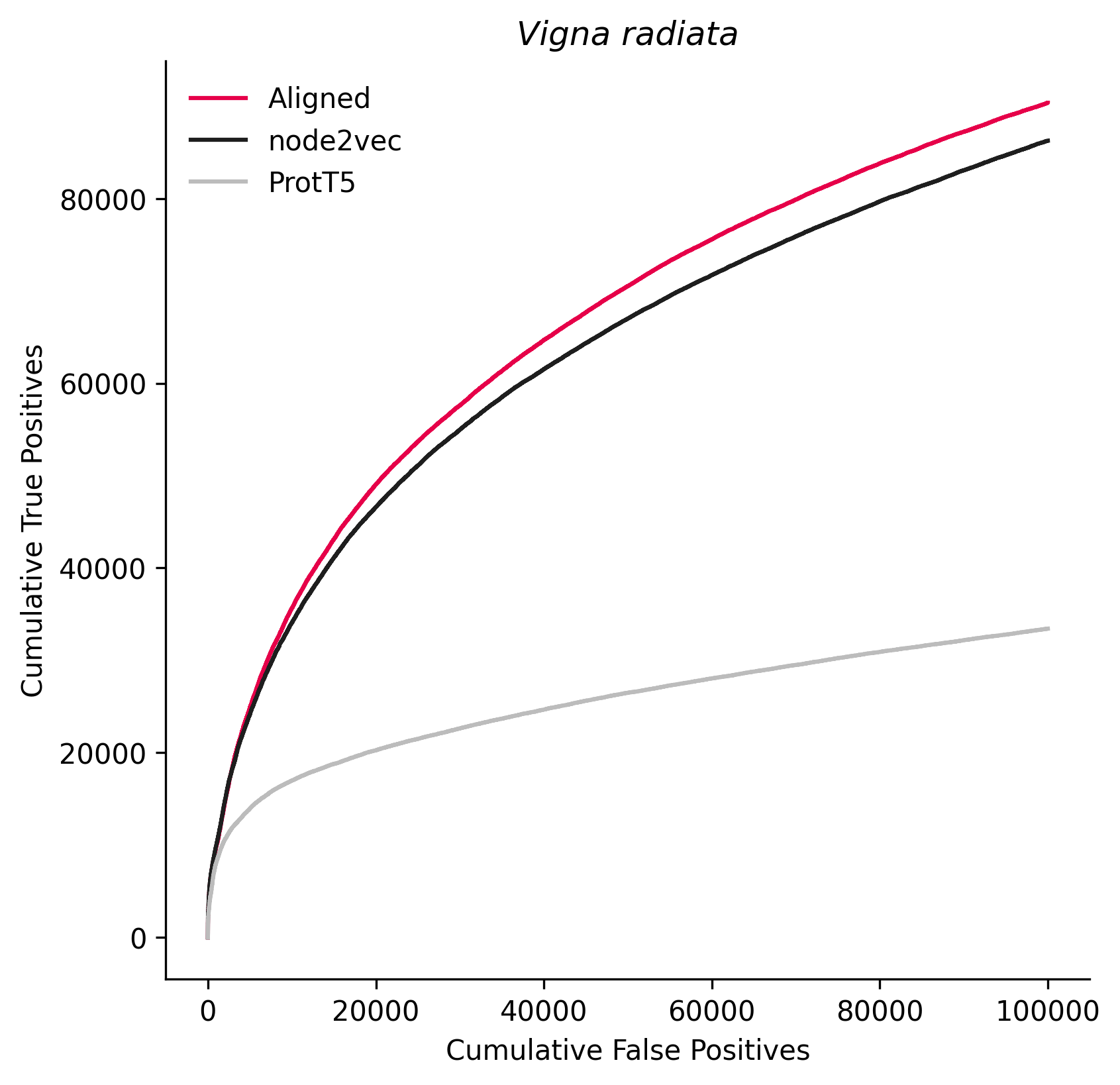

### 4113.png

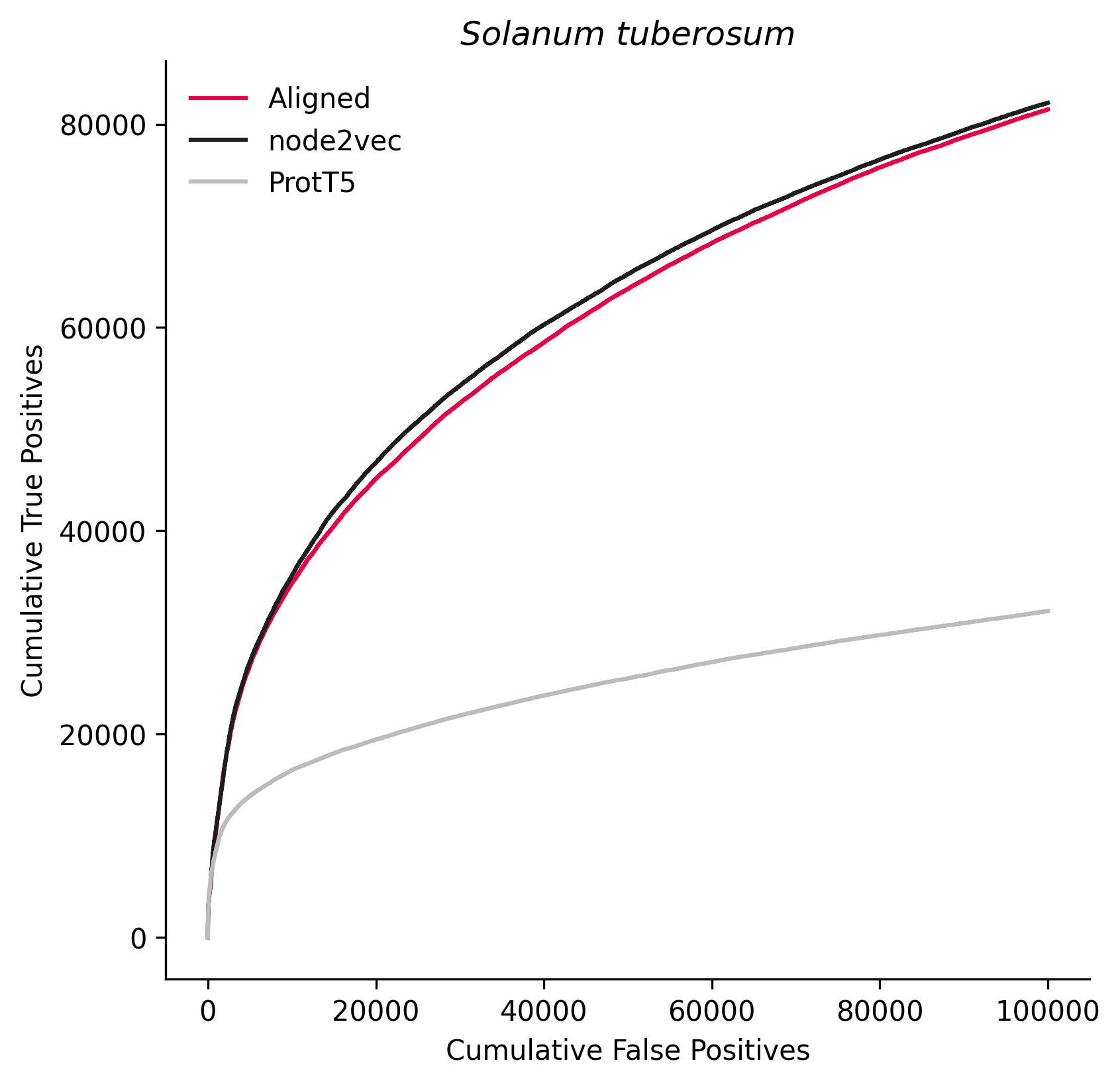

### 4932.png

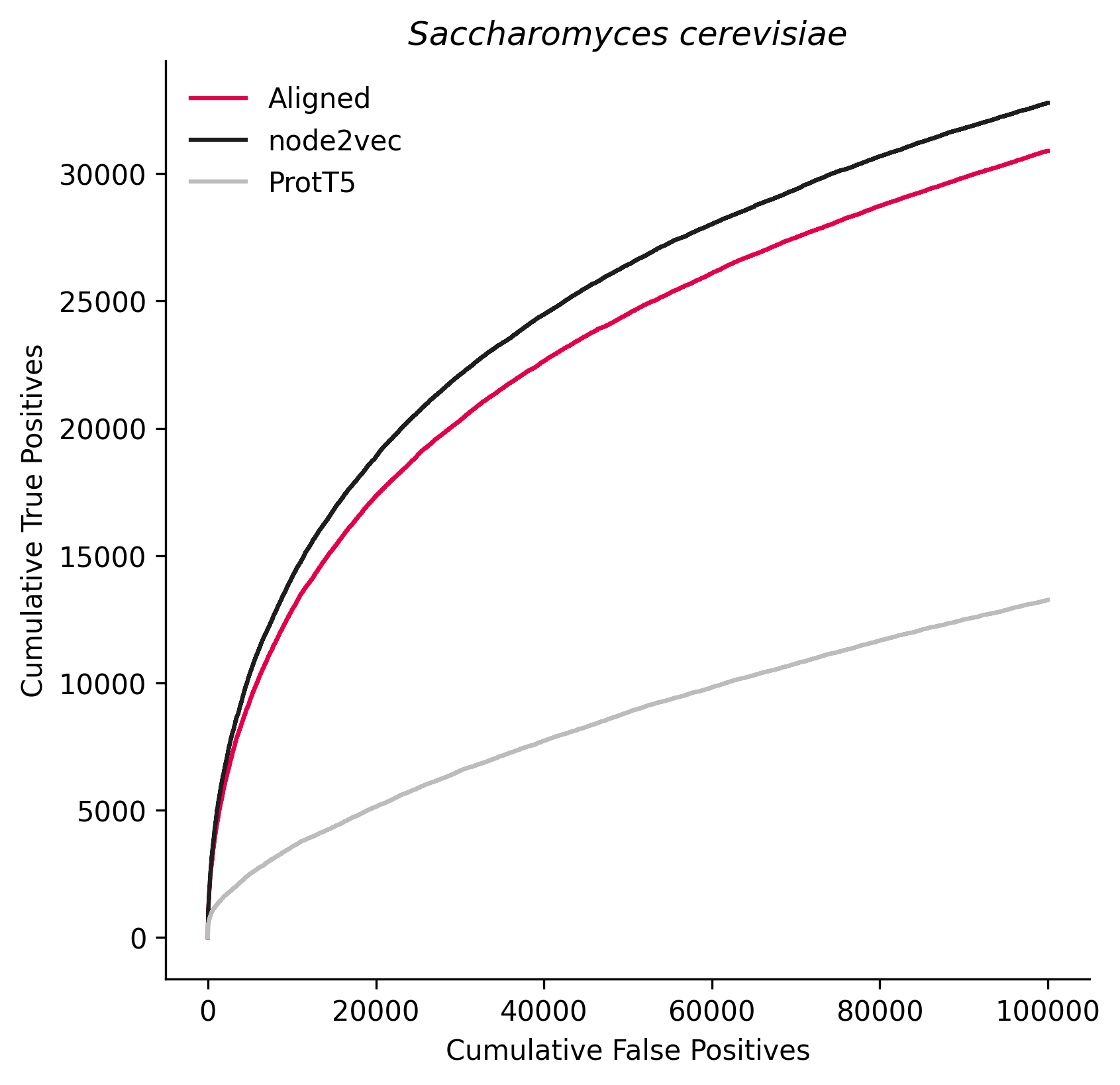

### 5664.png

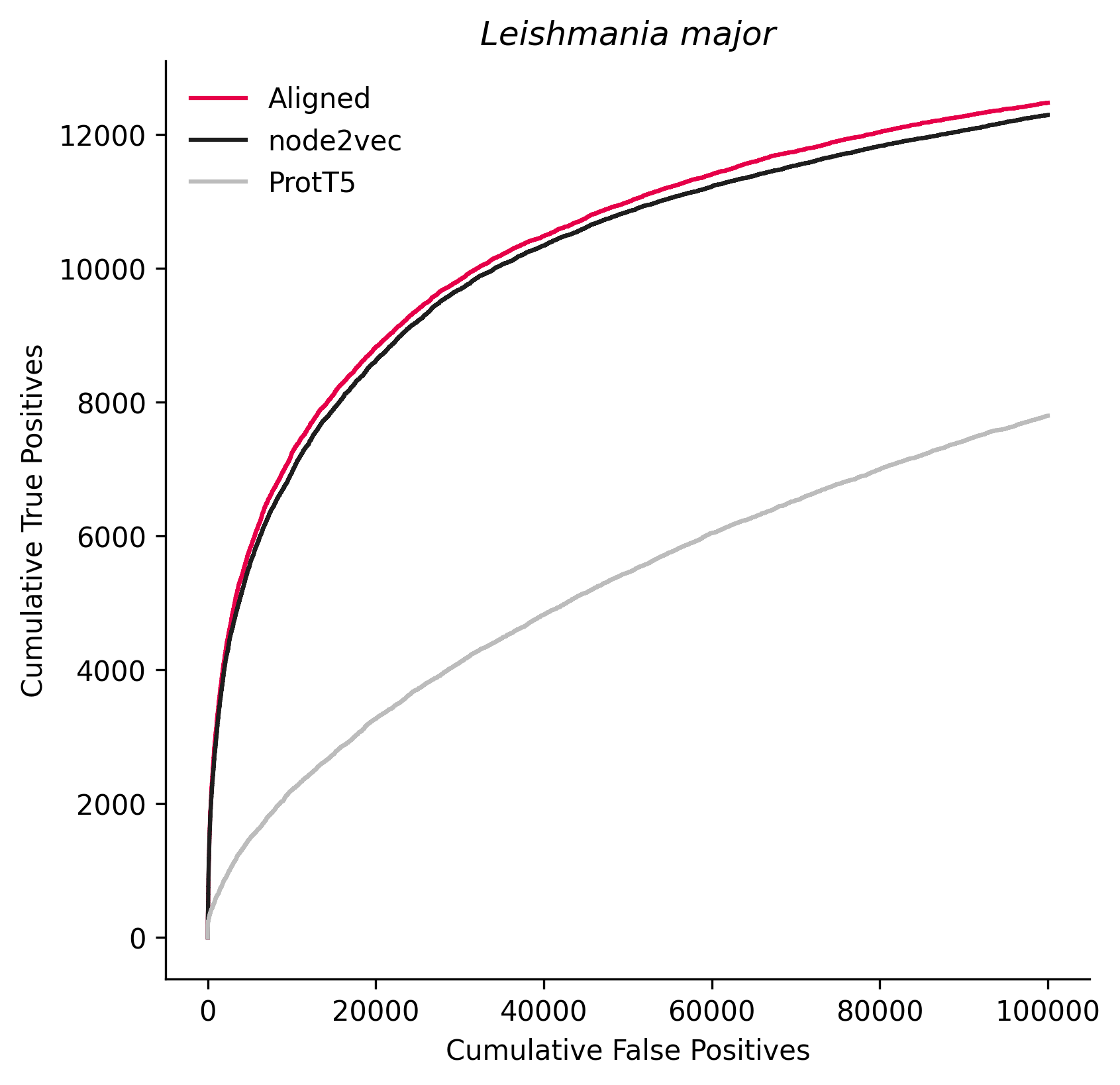

### 5671.png

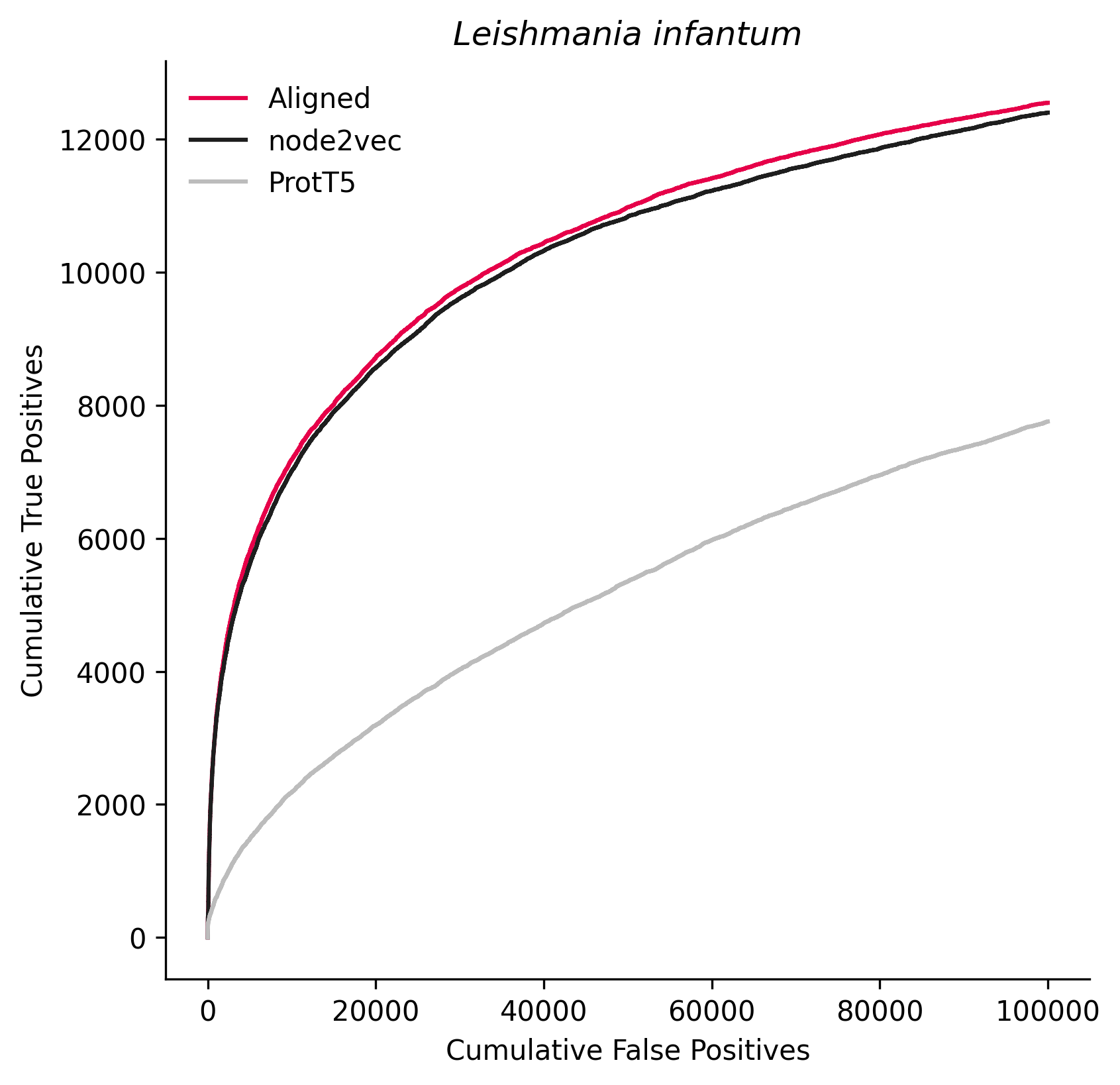

### 5722.png

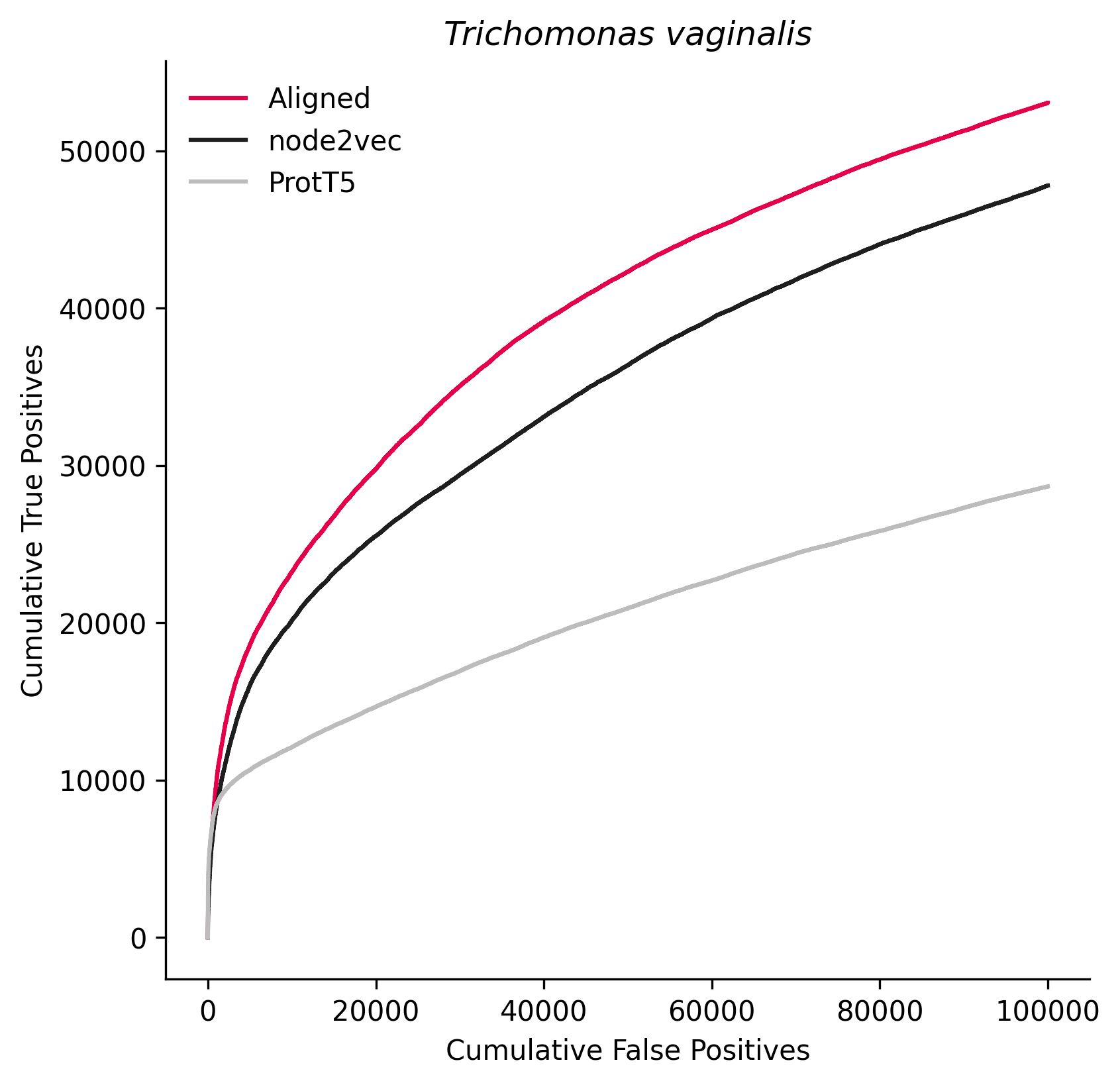

### 5786.png

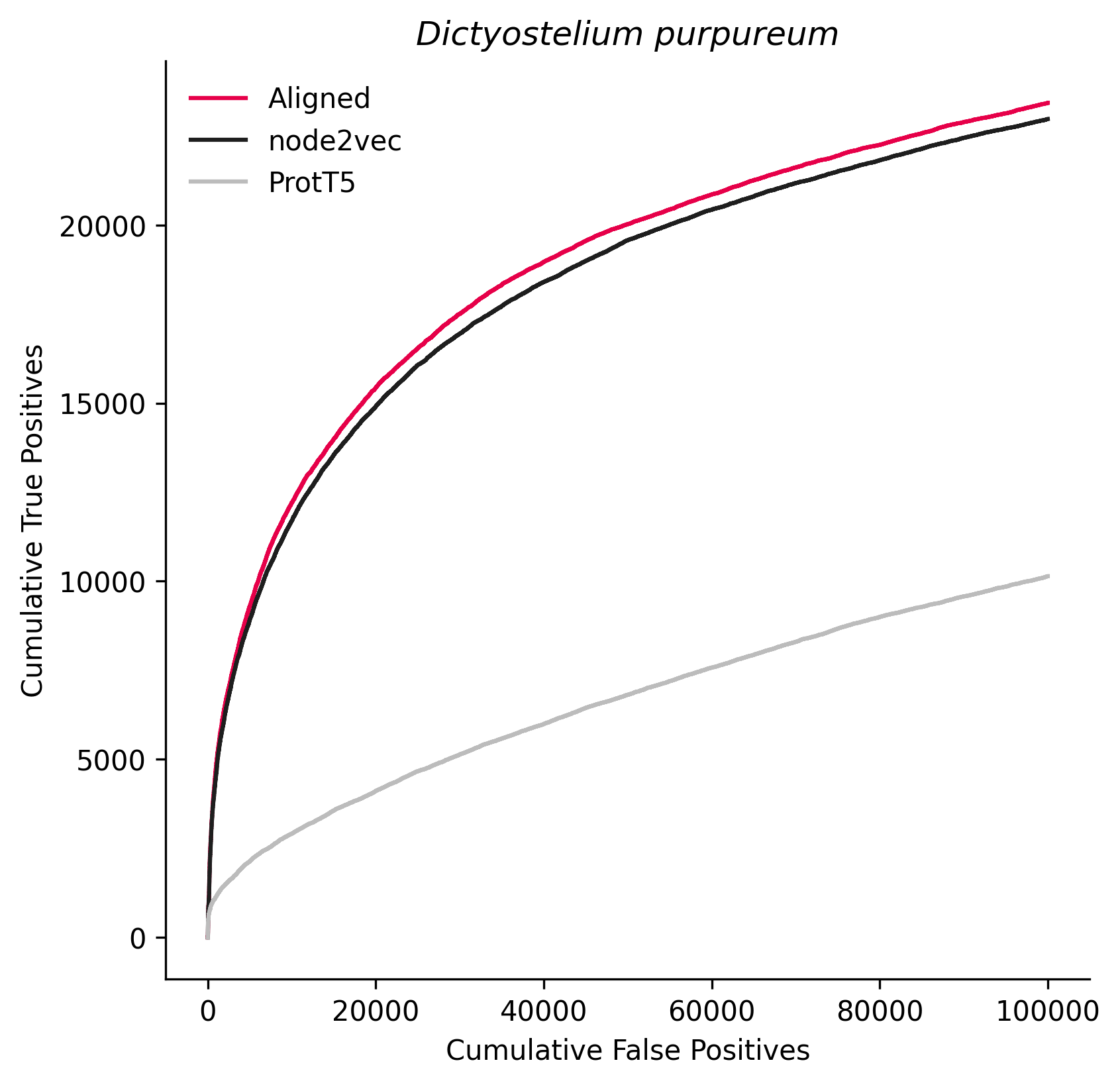

### 5888.png

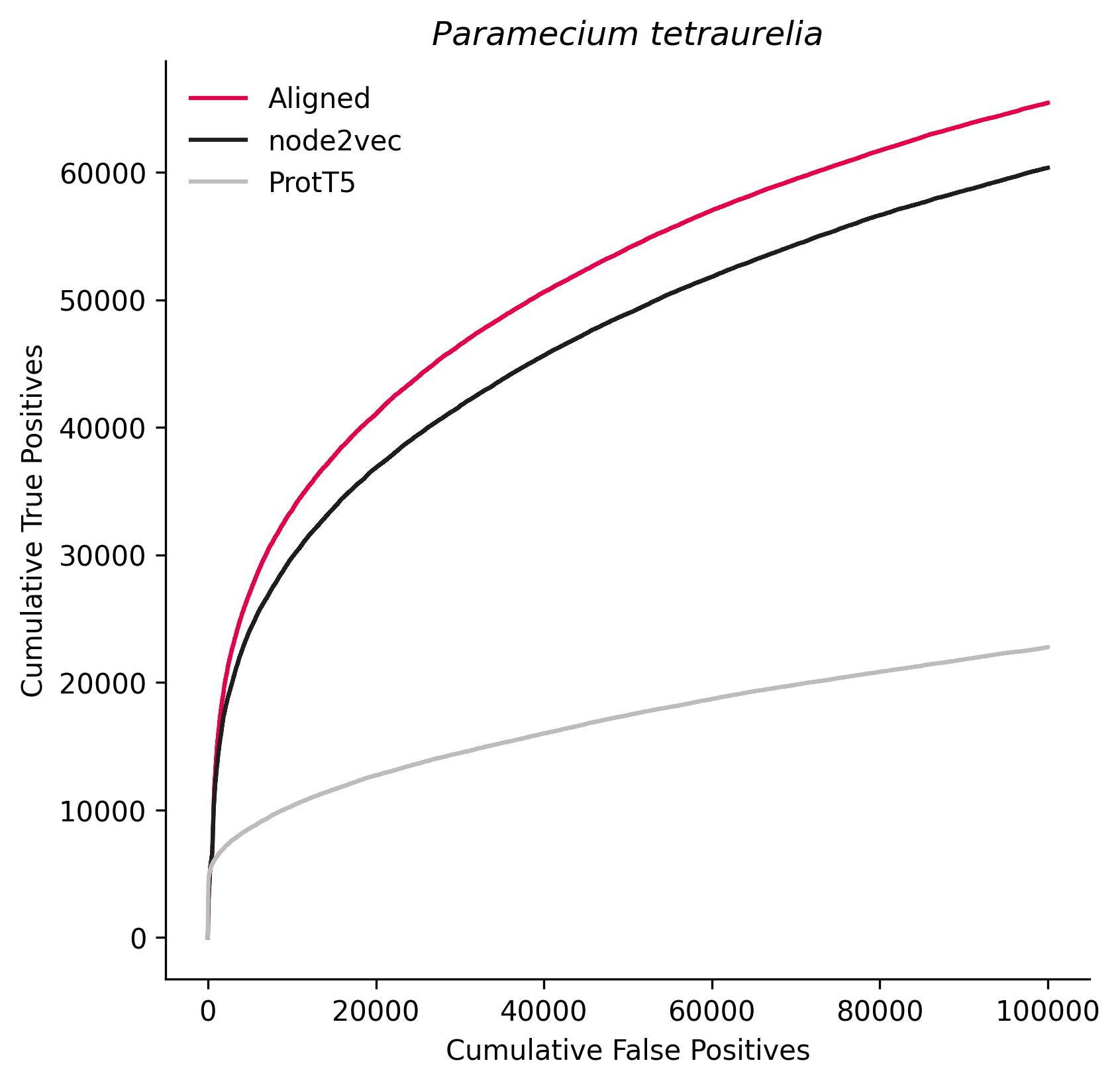

### 6334.png

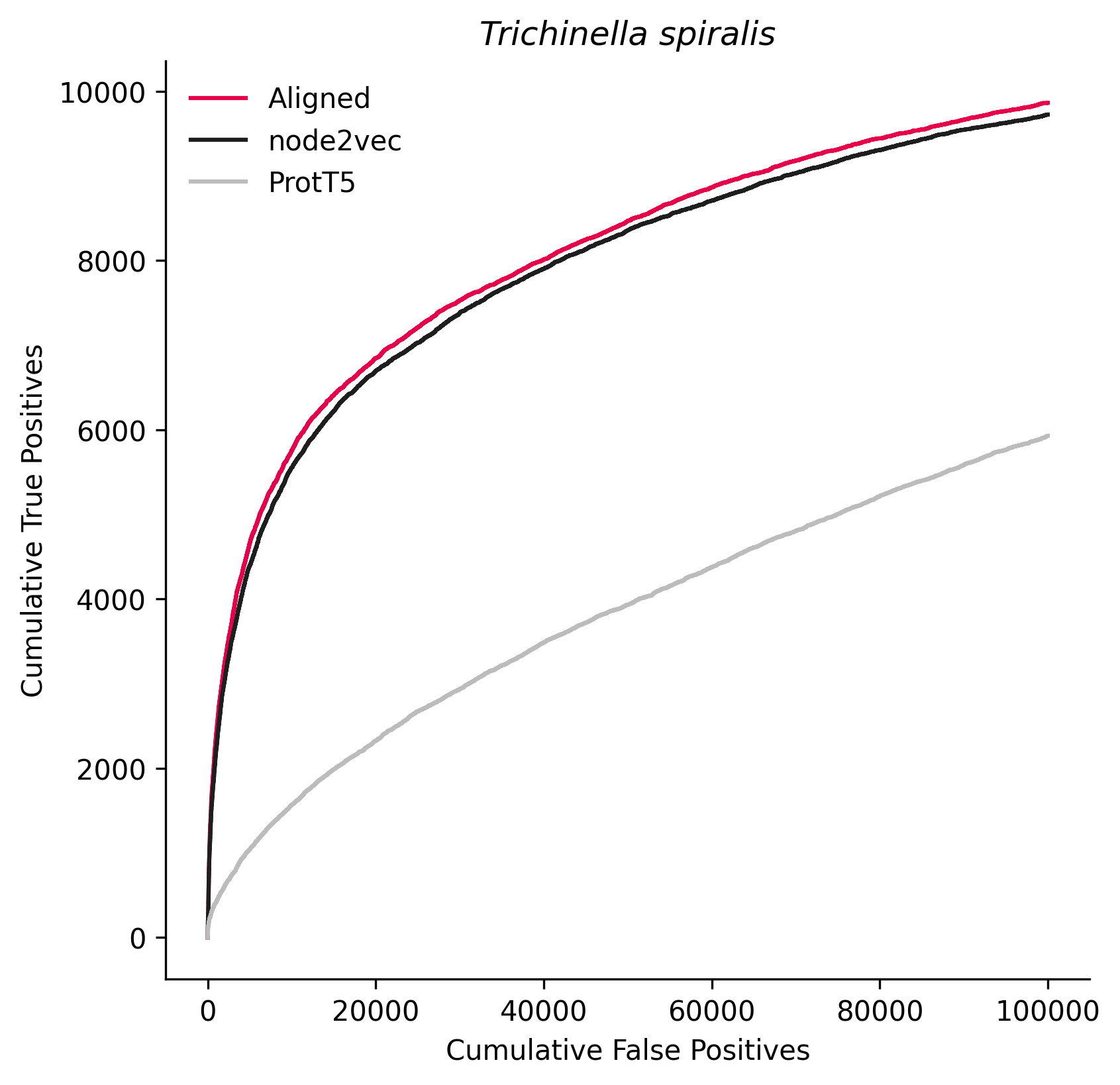

### 6573.png

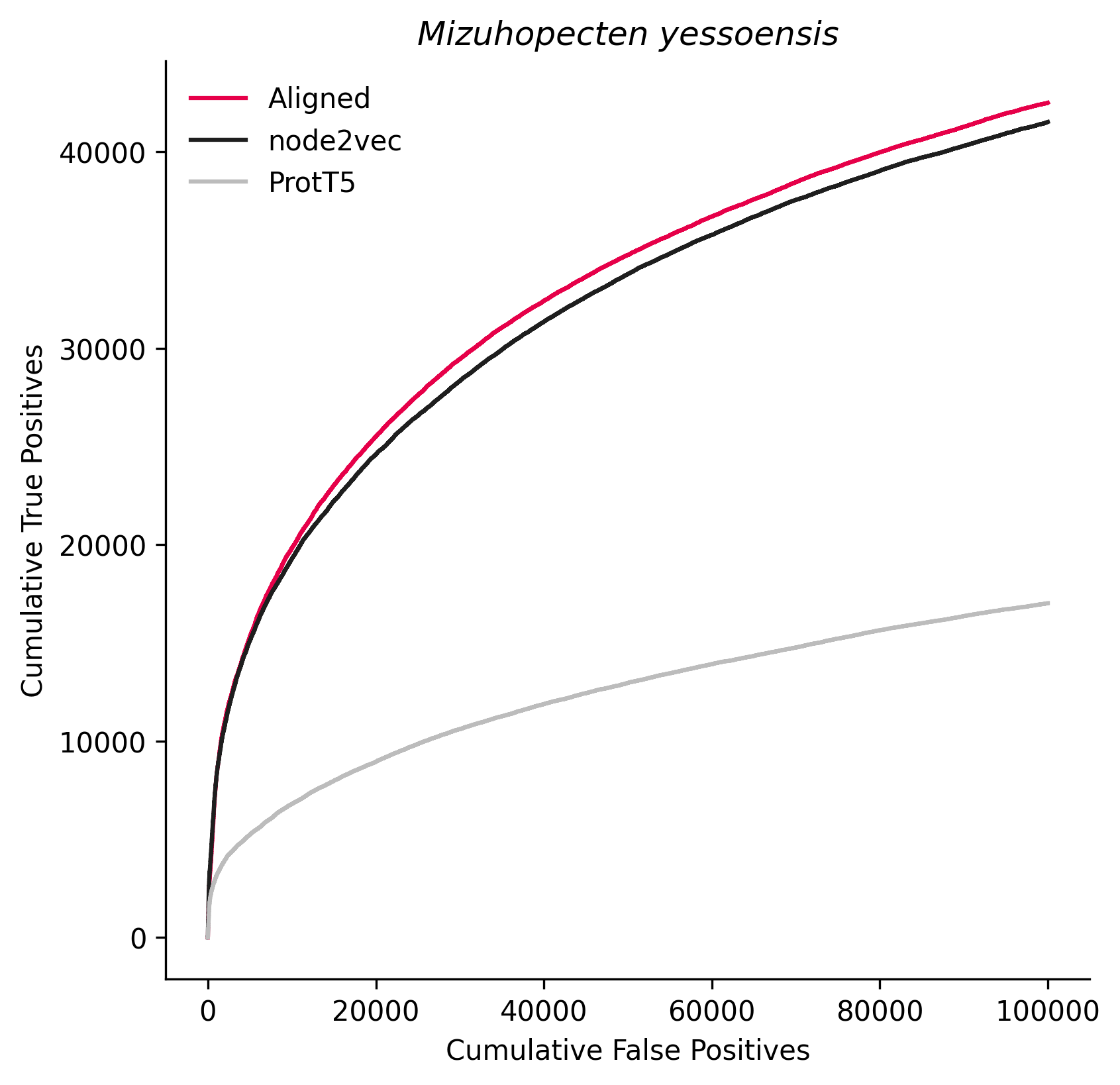

### 6669.png

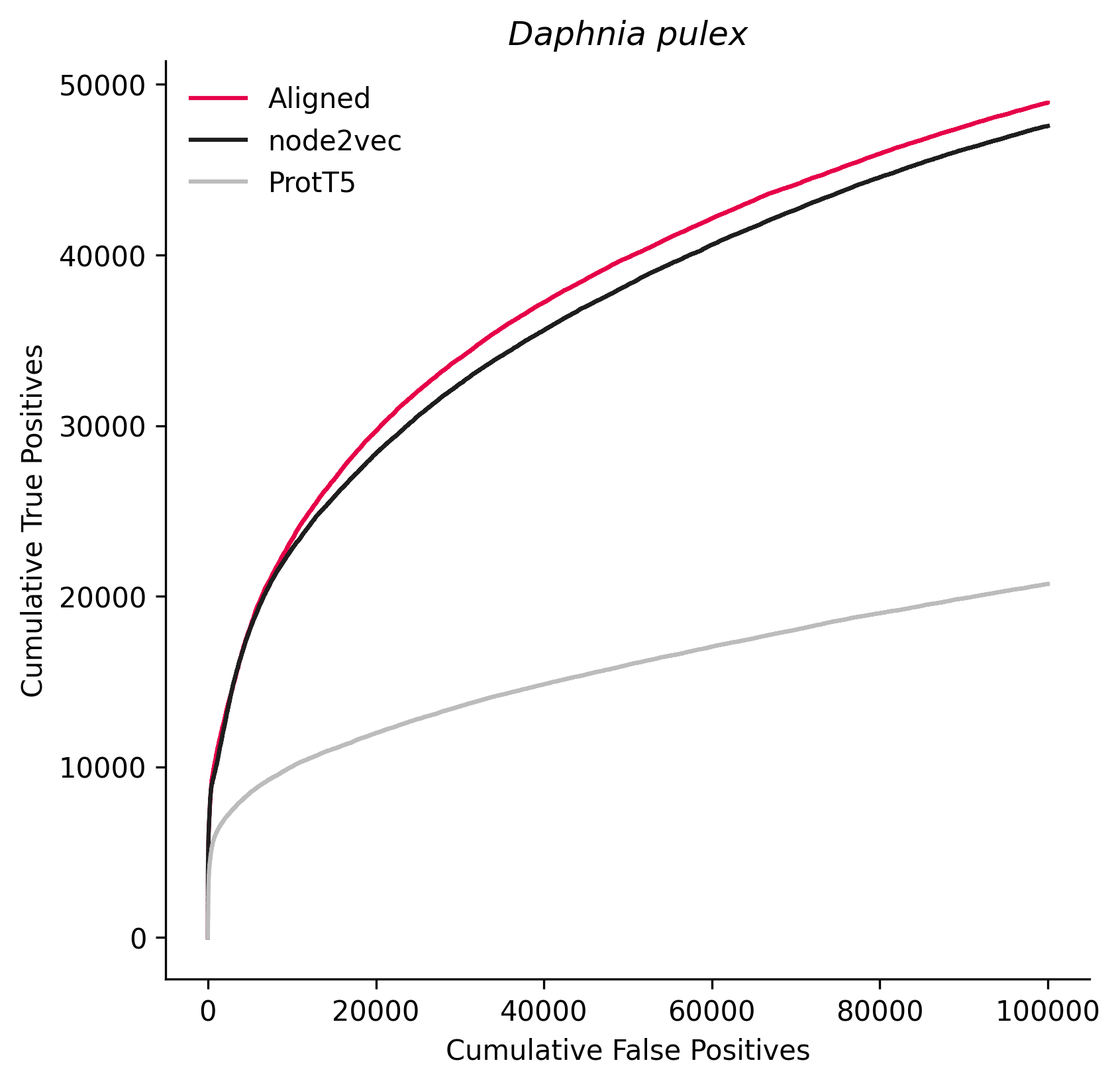

### 6689.png

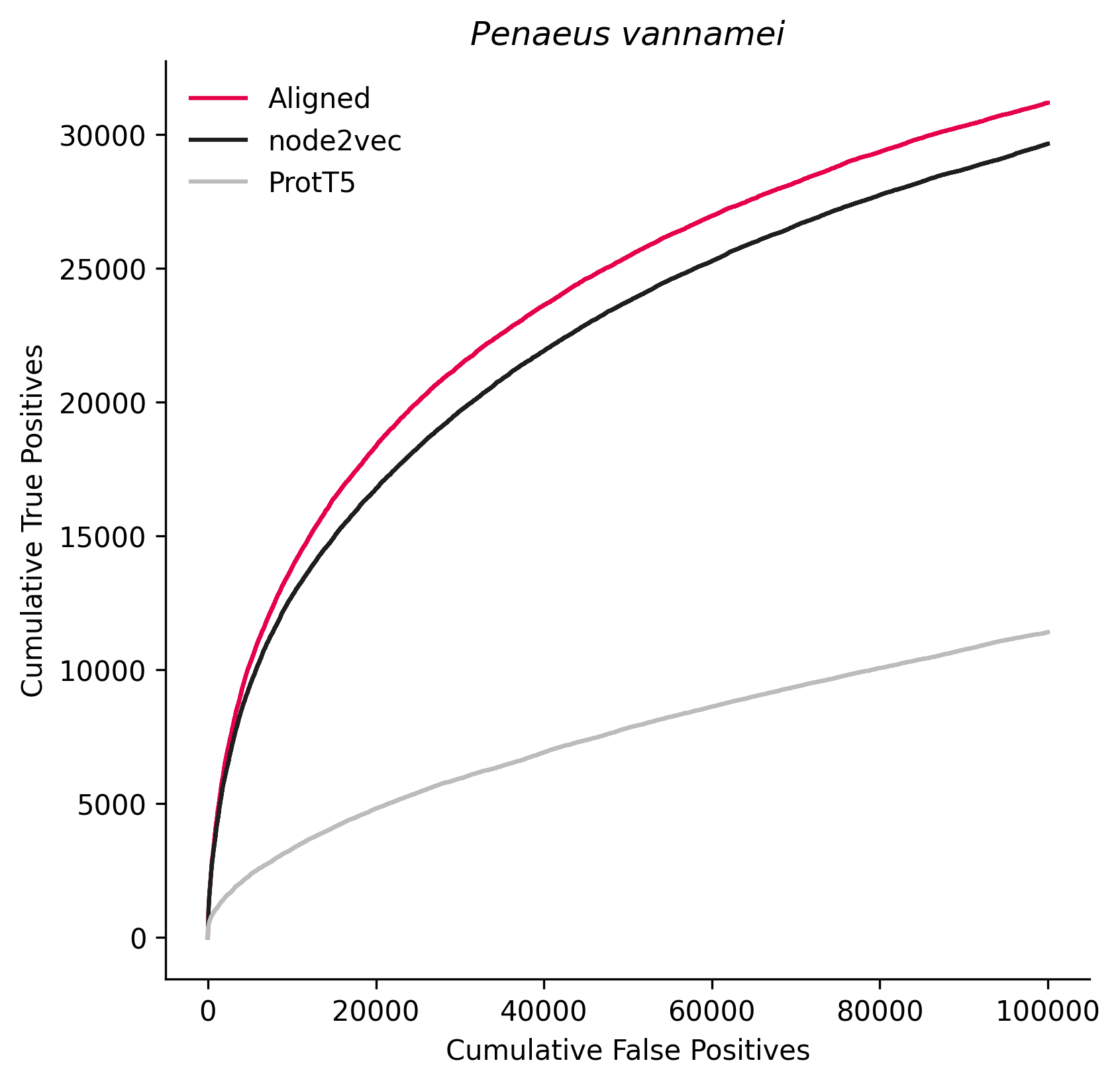

### 7091.png

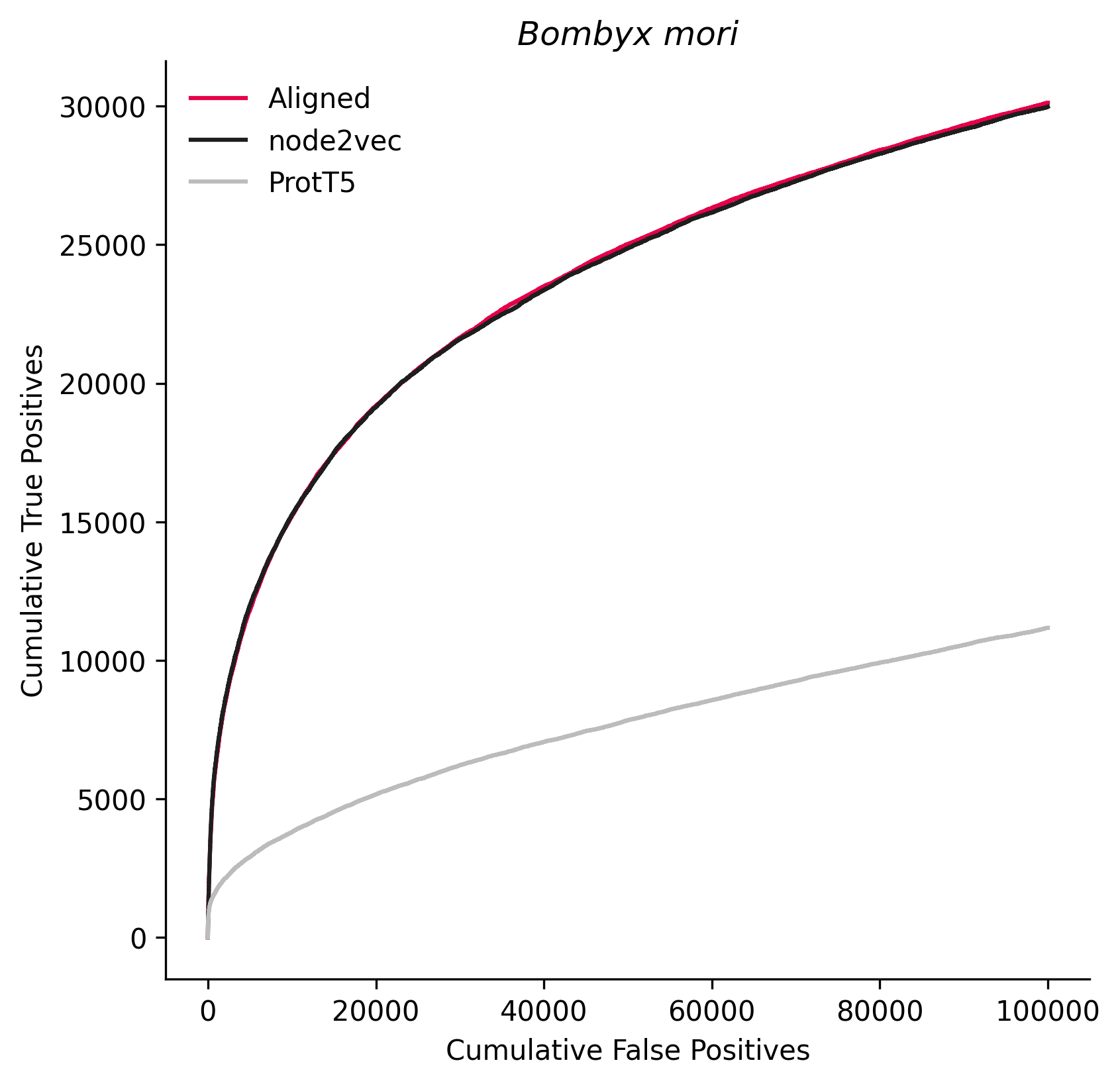

### 7160.png

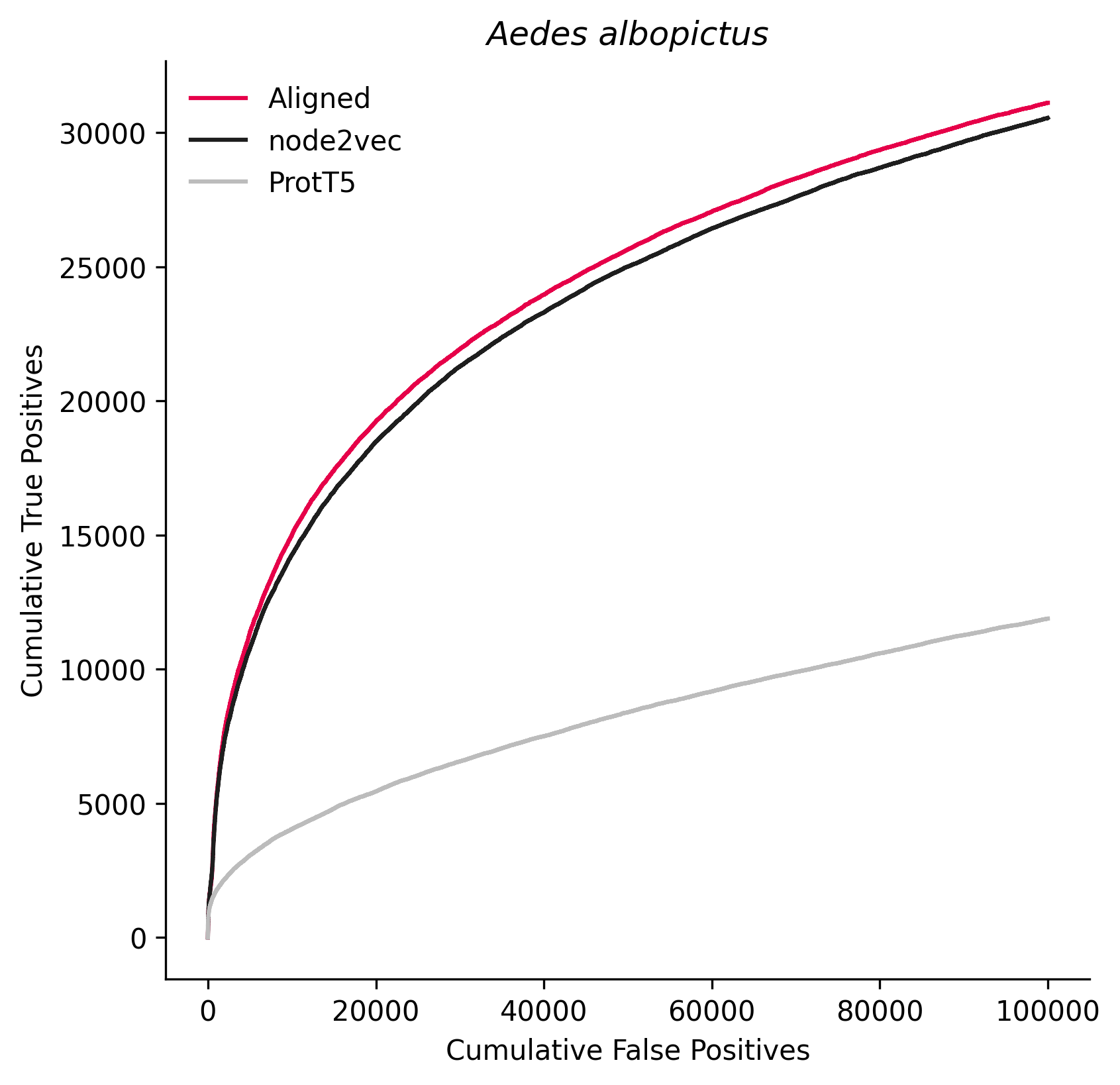

### 7209.png

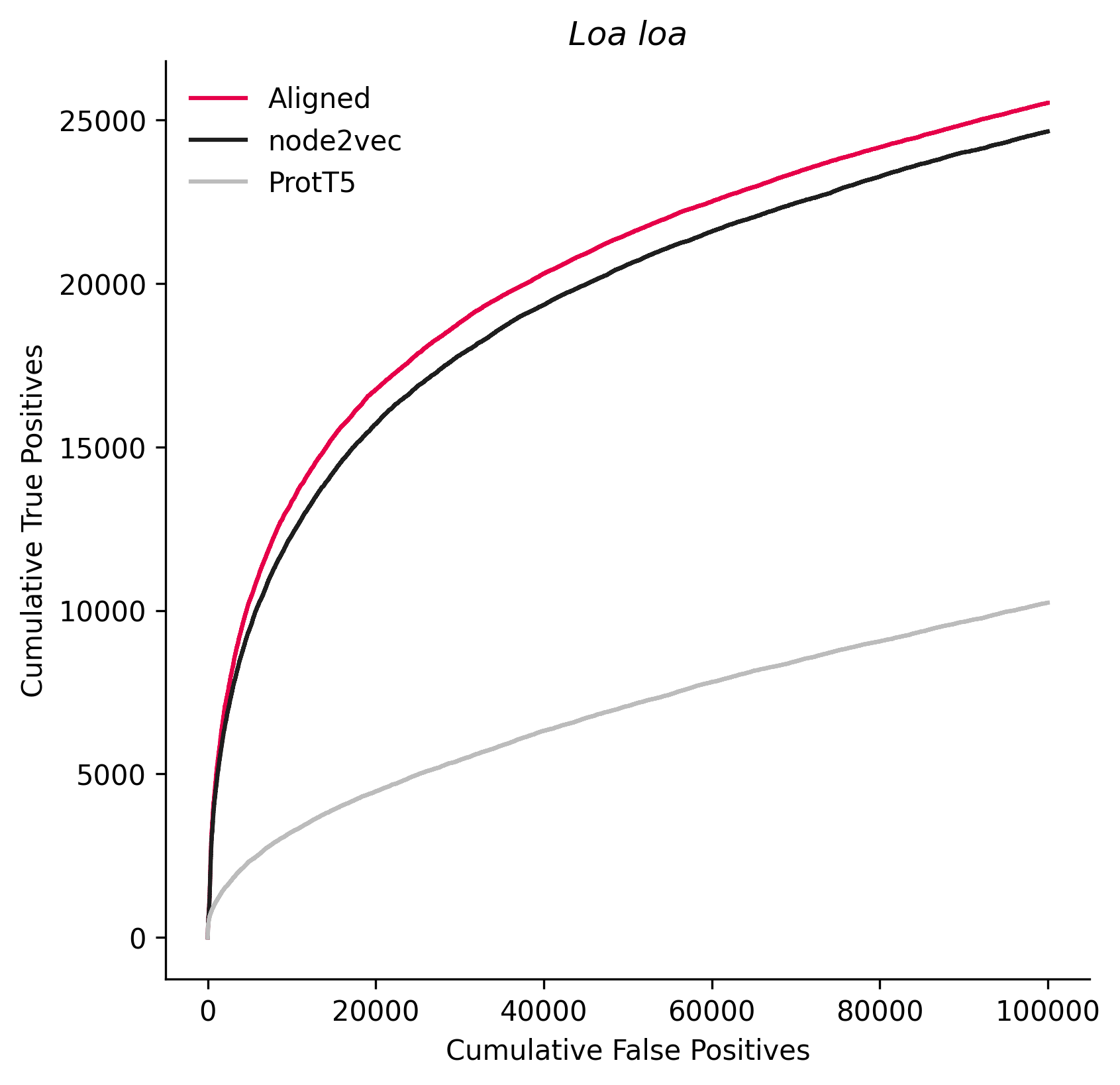

### 7222.png

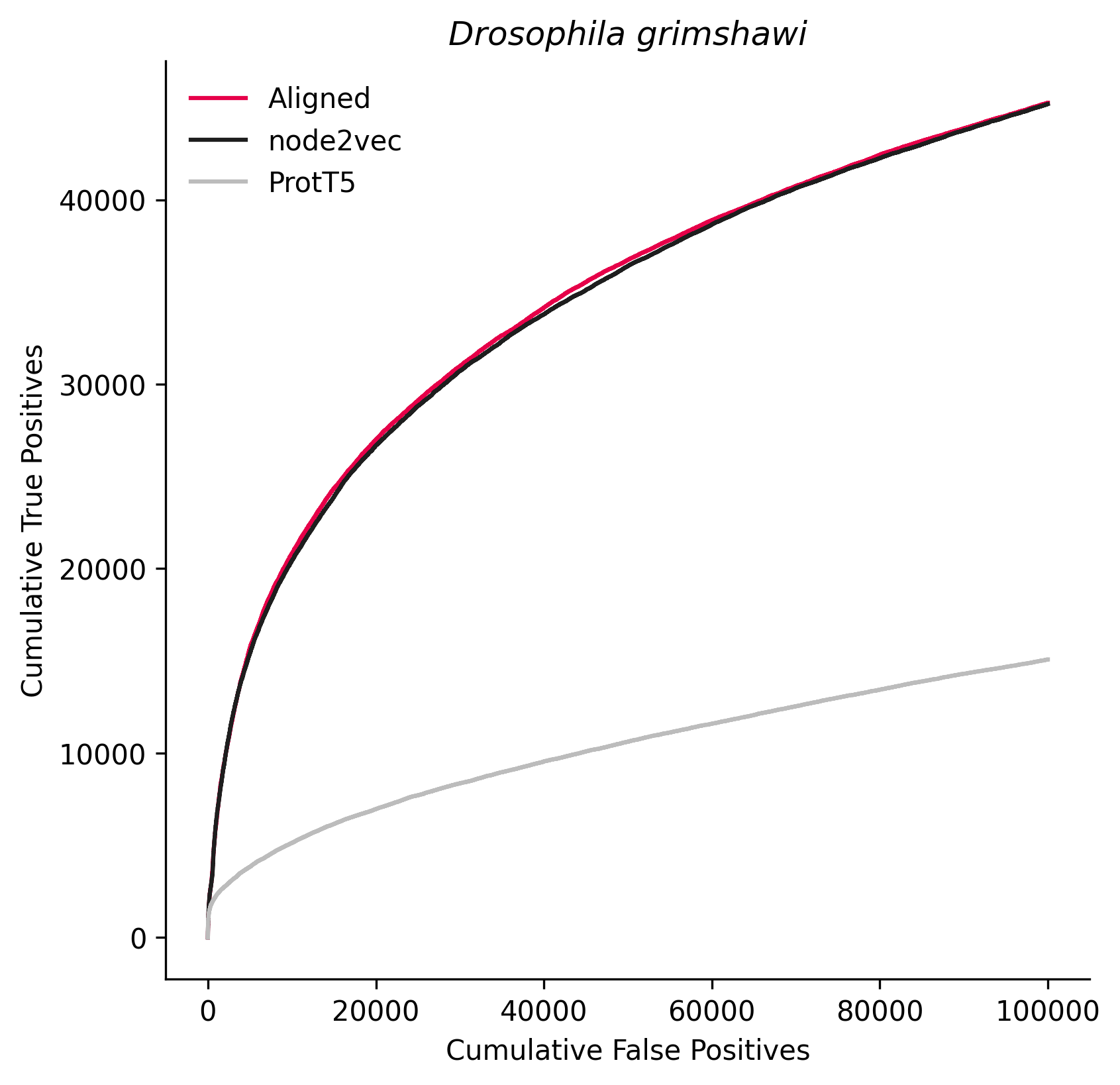

### 7232.png

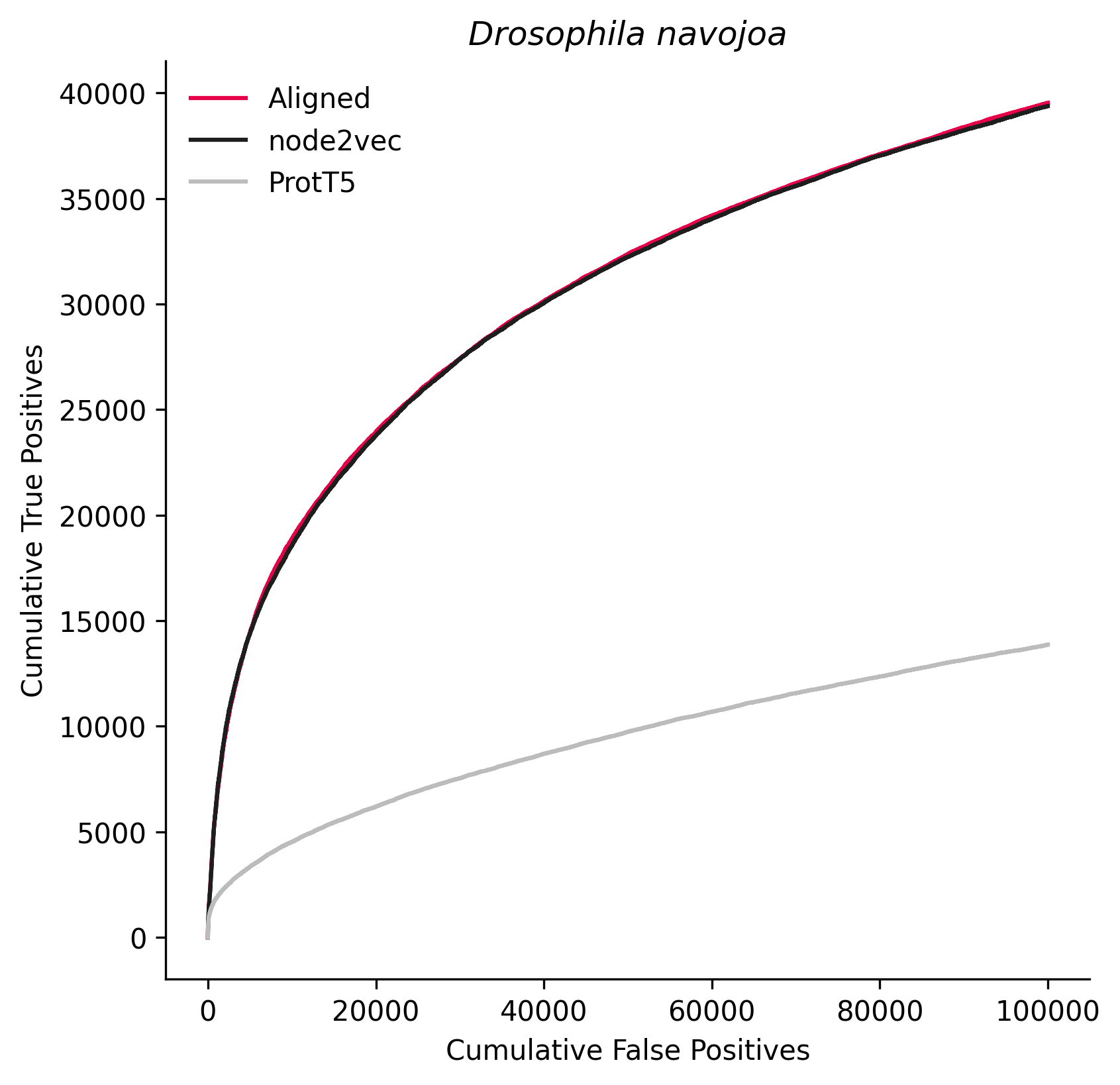

### 7370.png

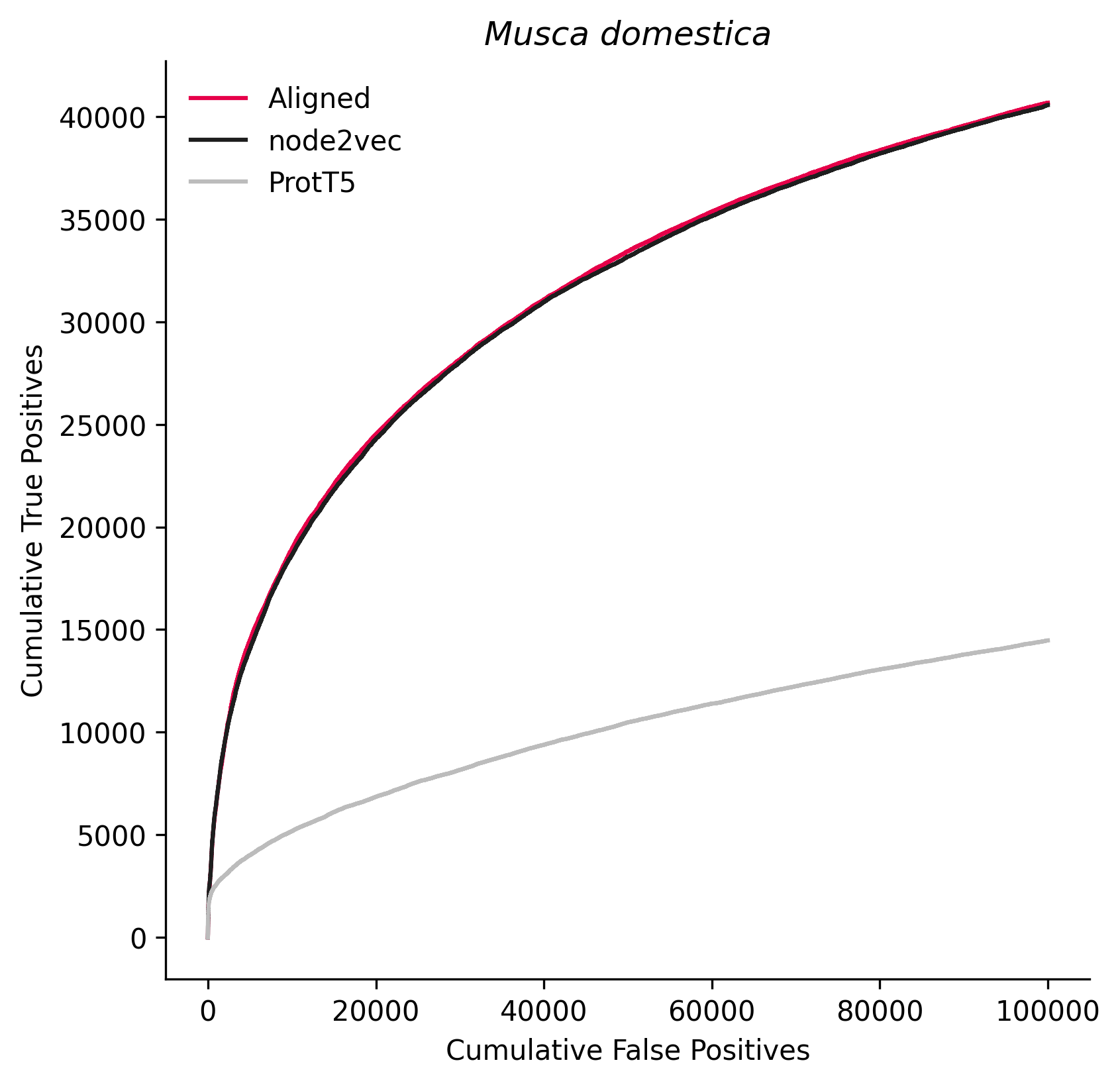

### 7574.png

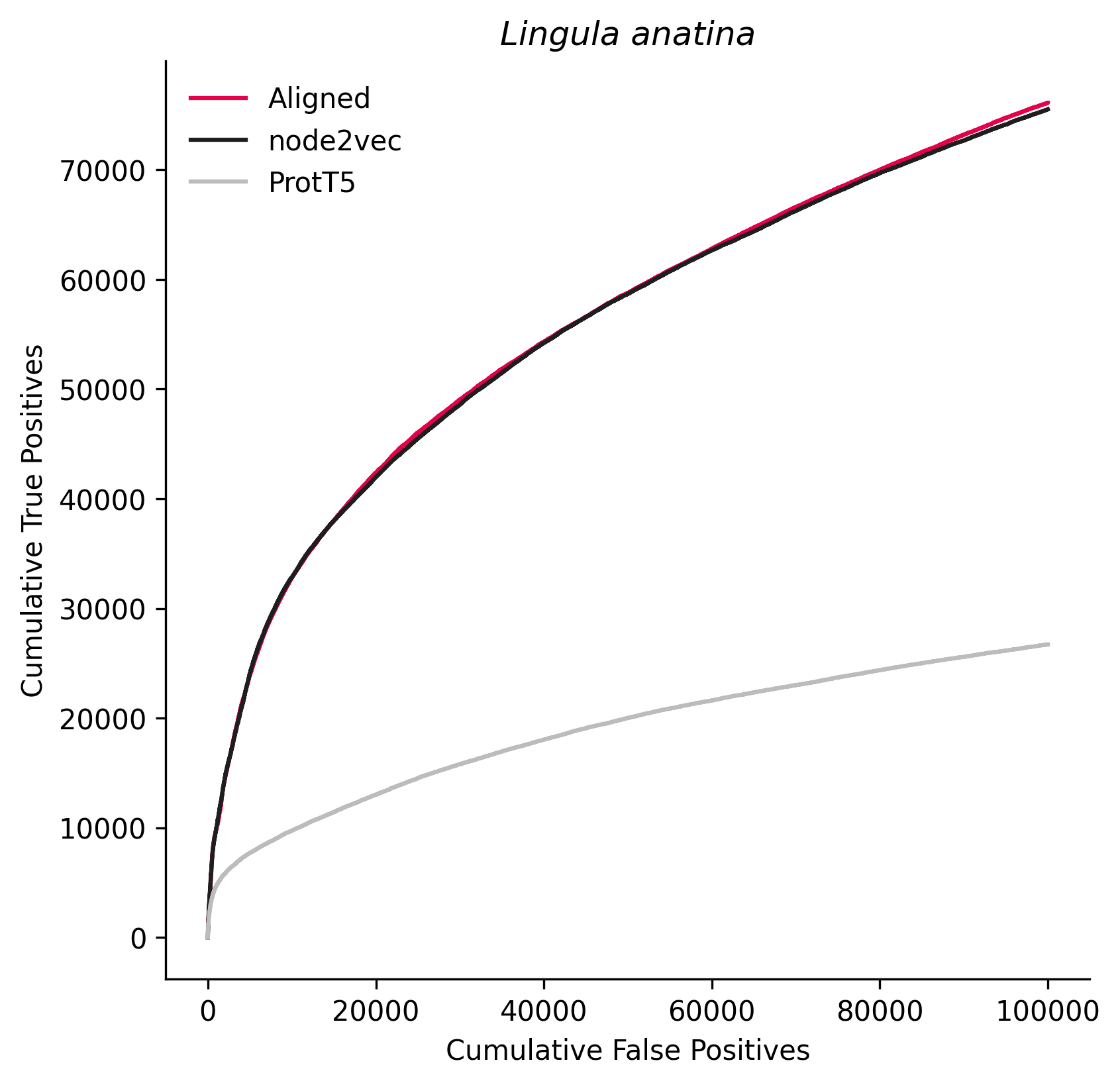

### 7668.png

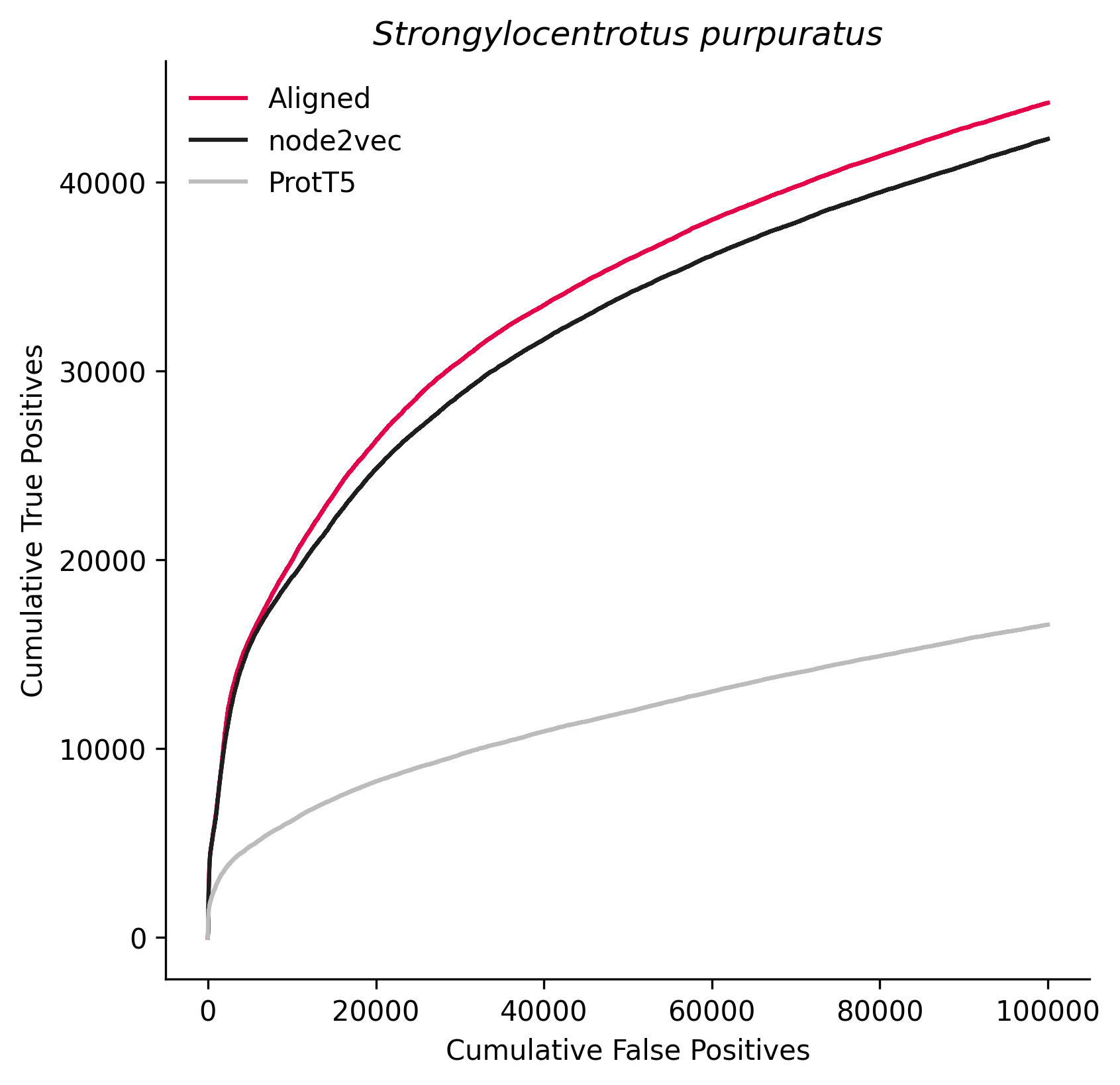

### 7719.png

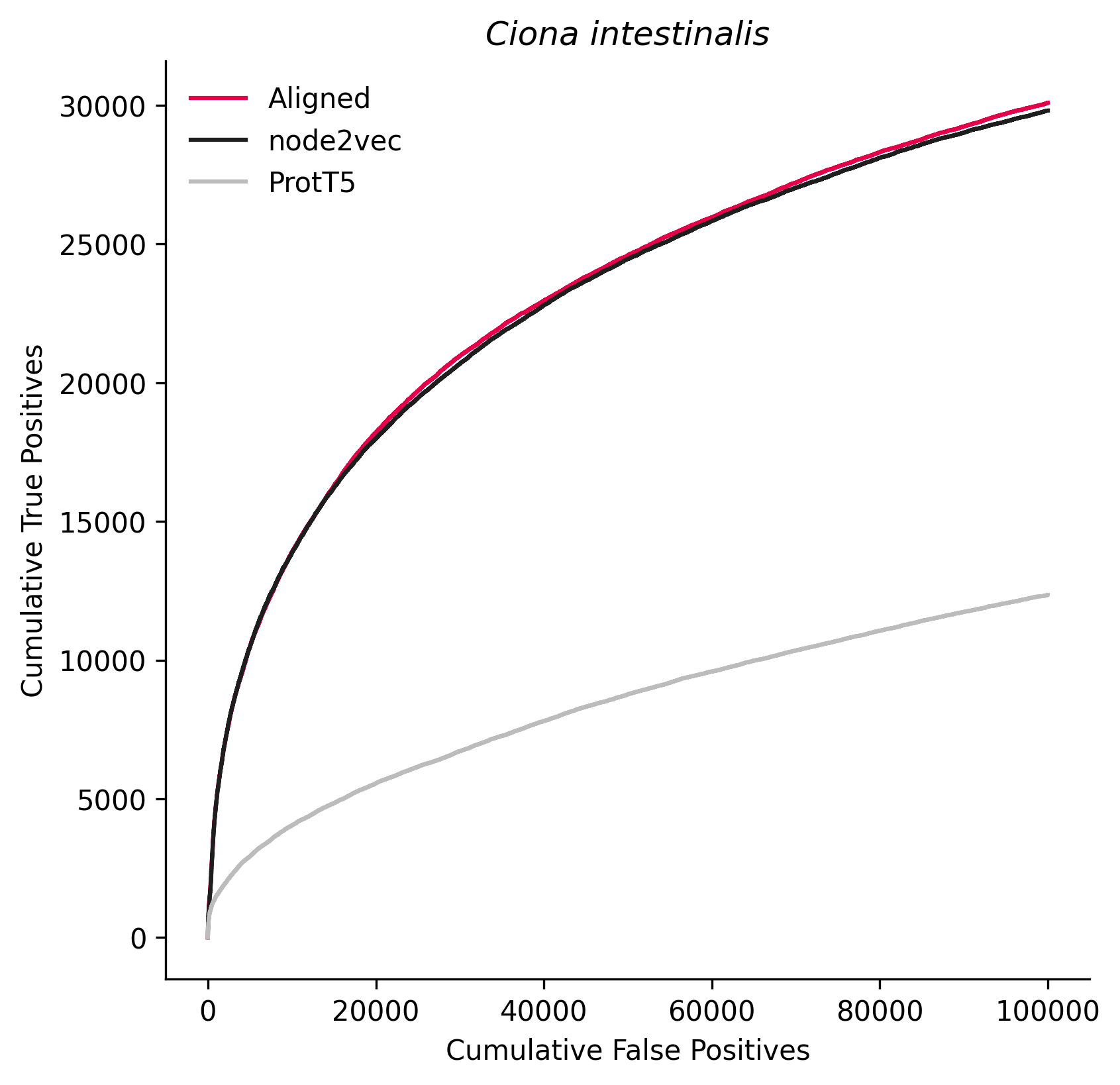

### 7868.png

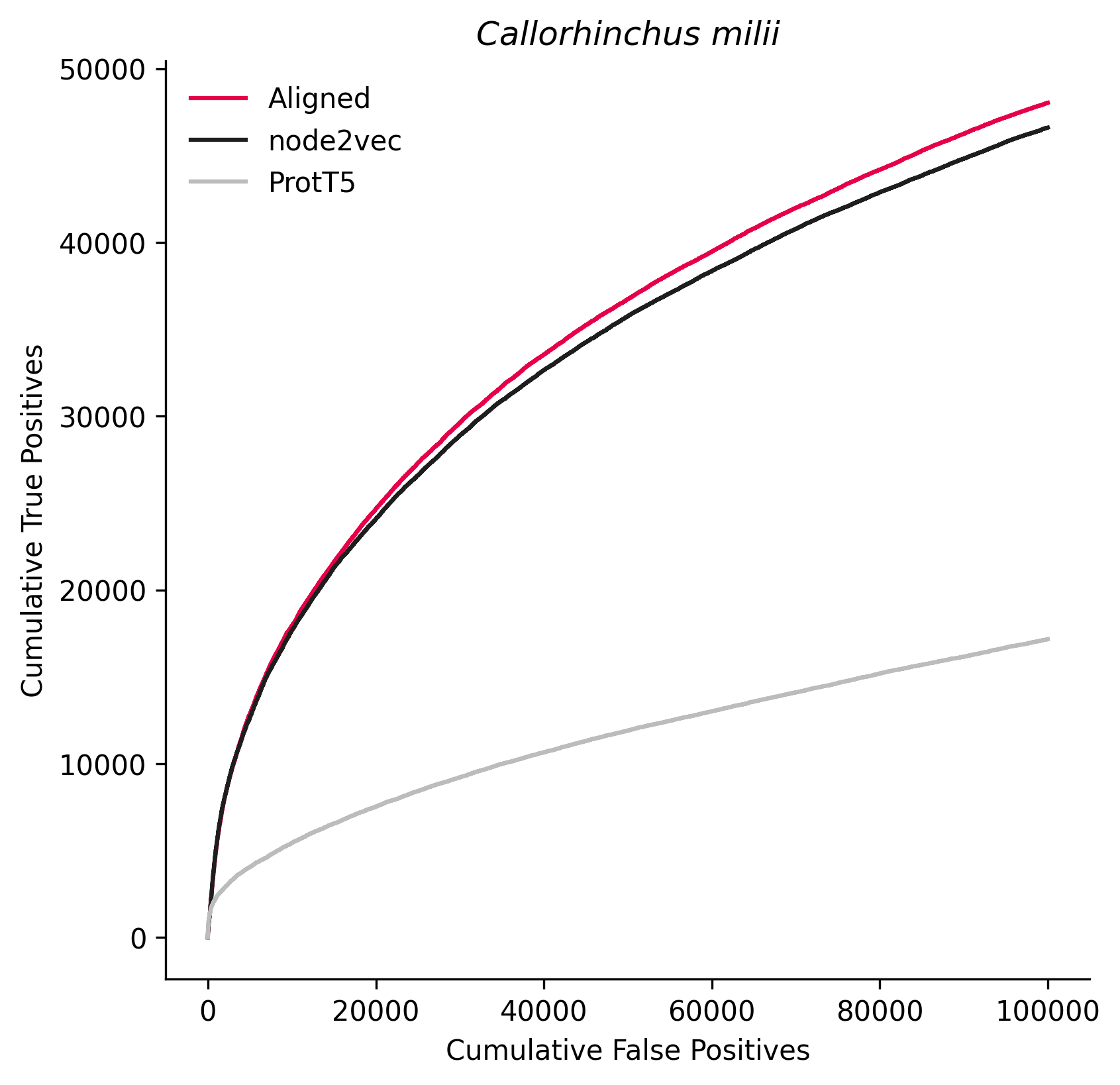

### 7994.png

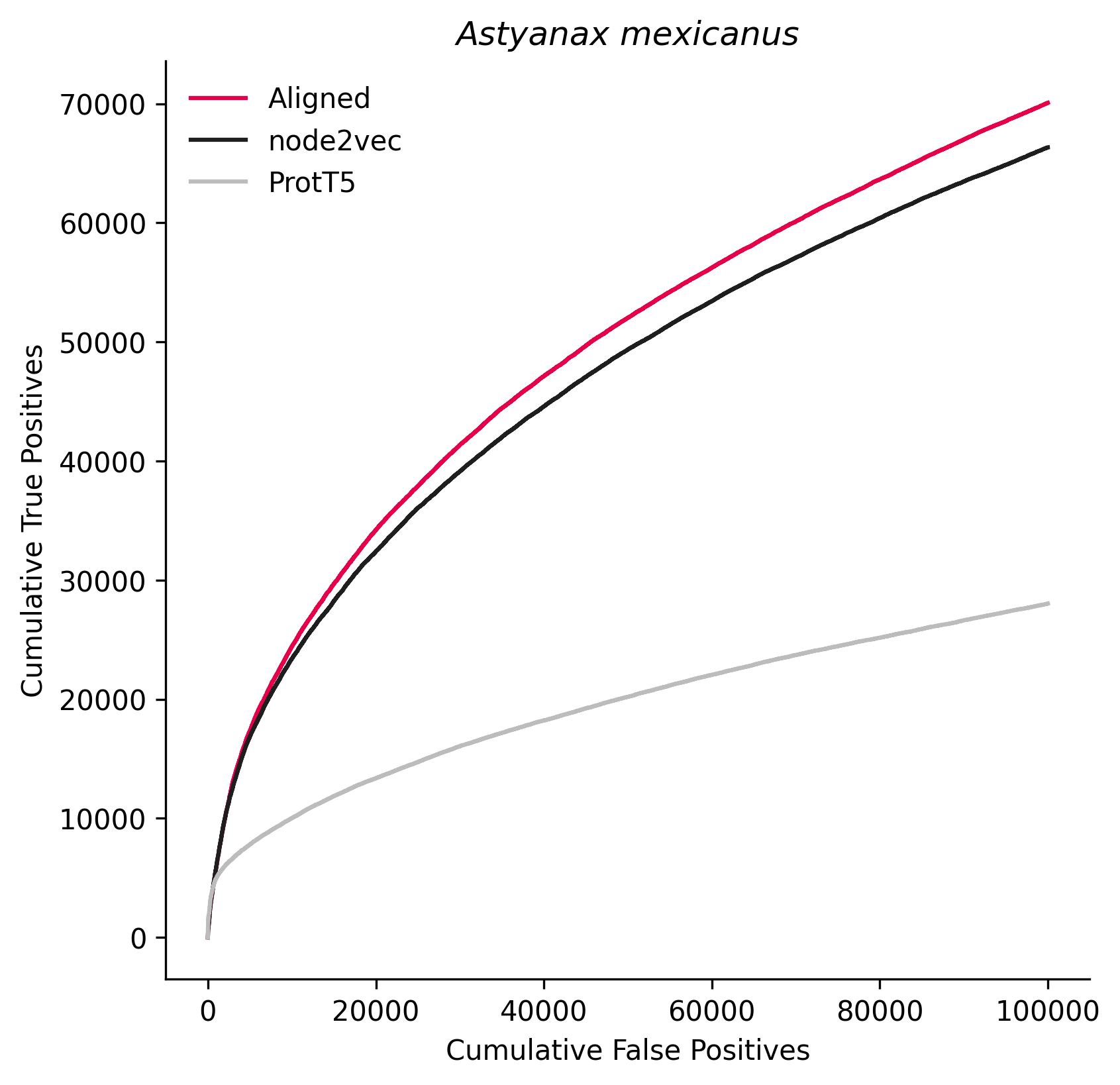

### 8128.png

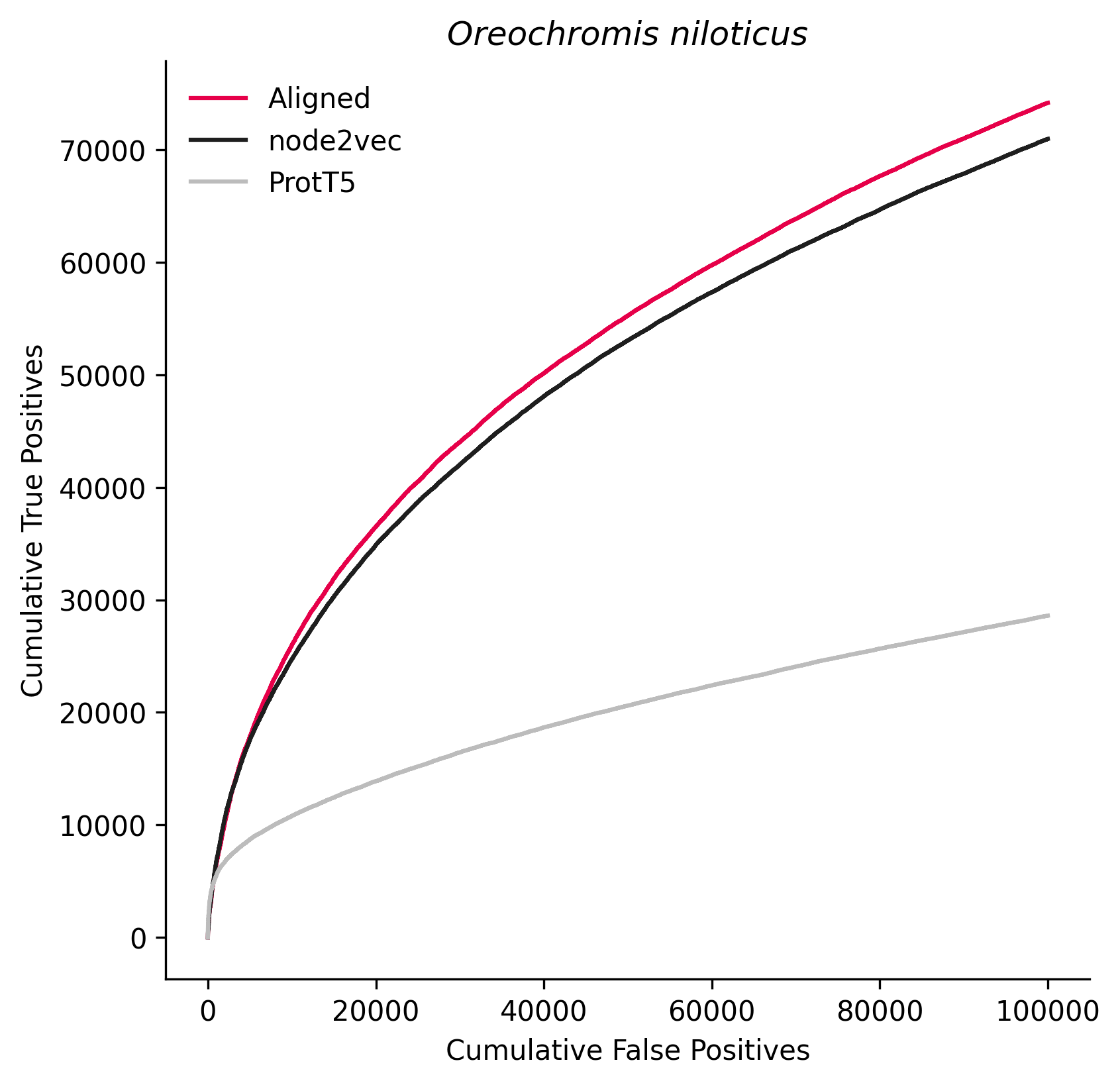

### 8364.png

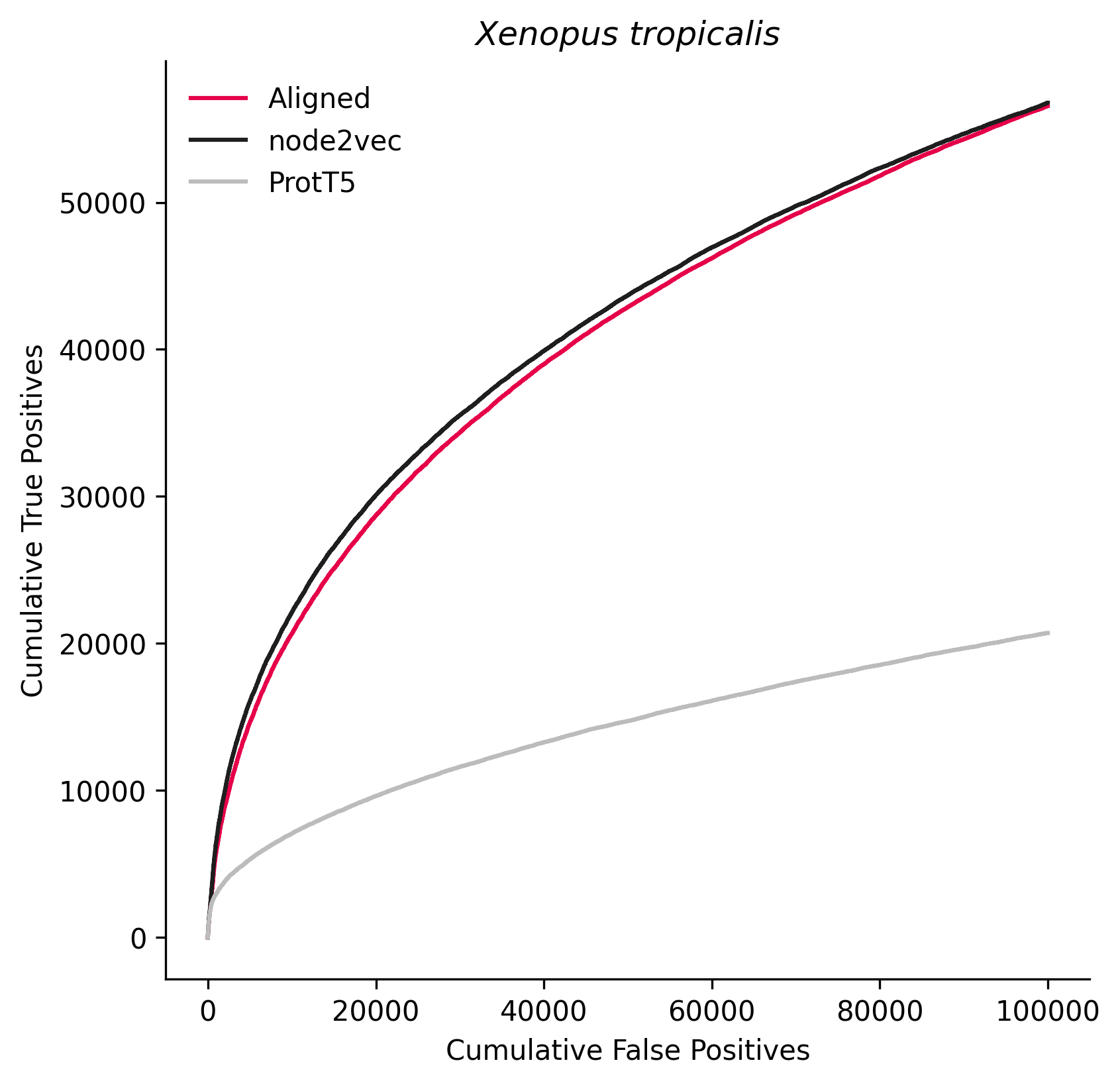

### 8478.png

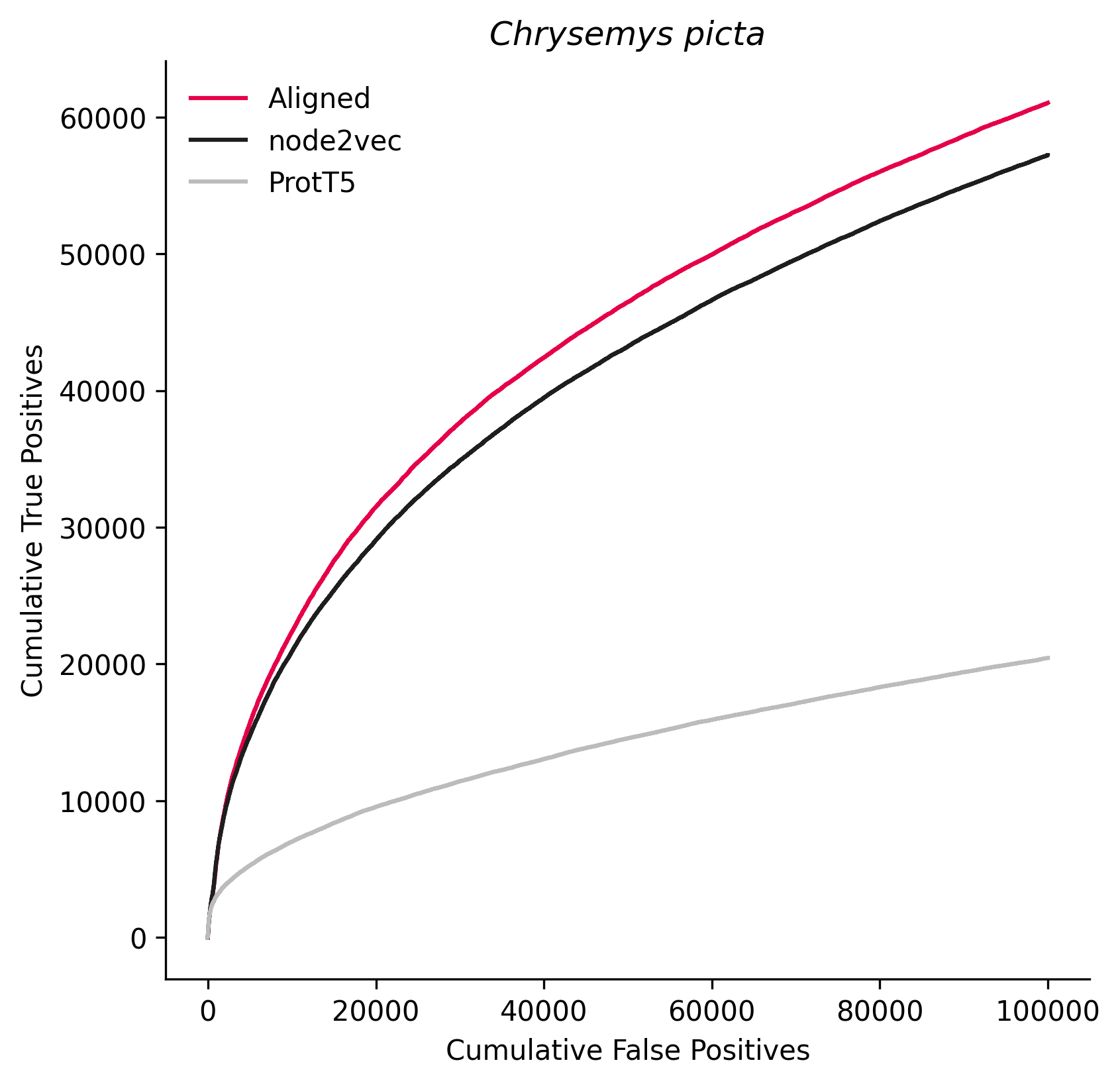

### 8996.png

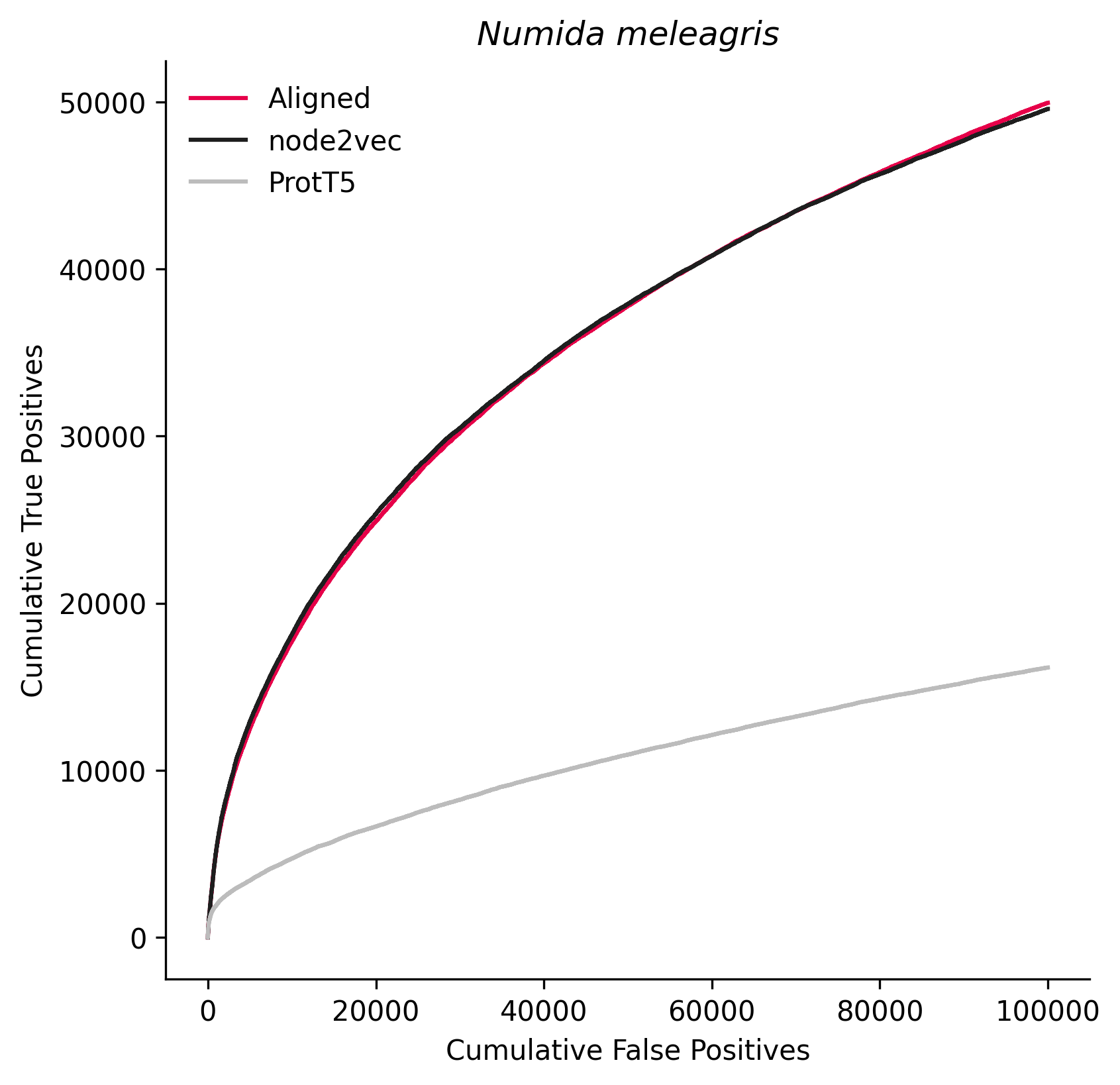

### 9031.png

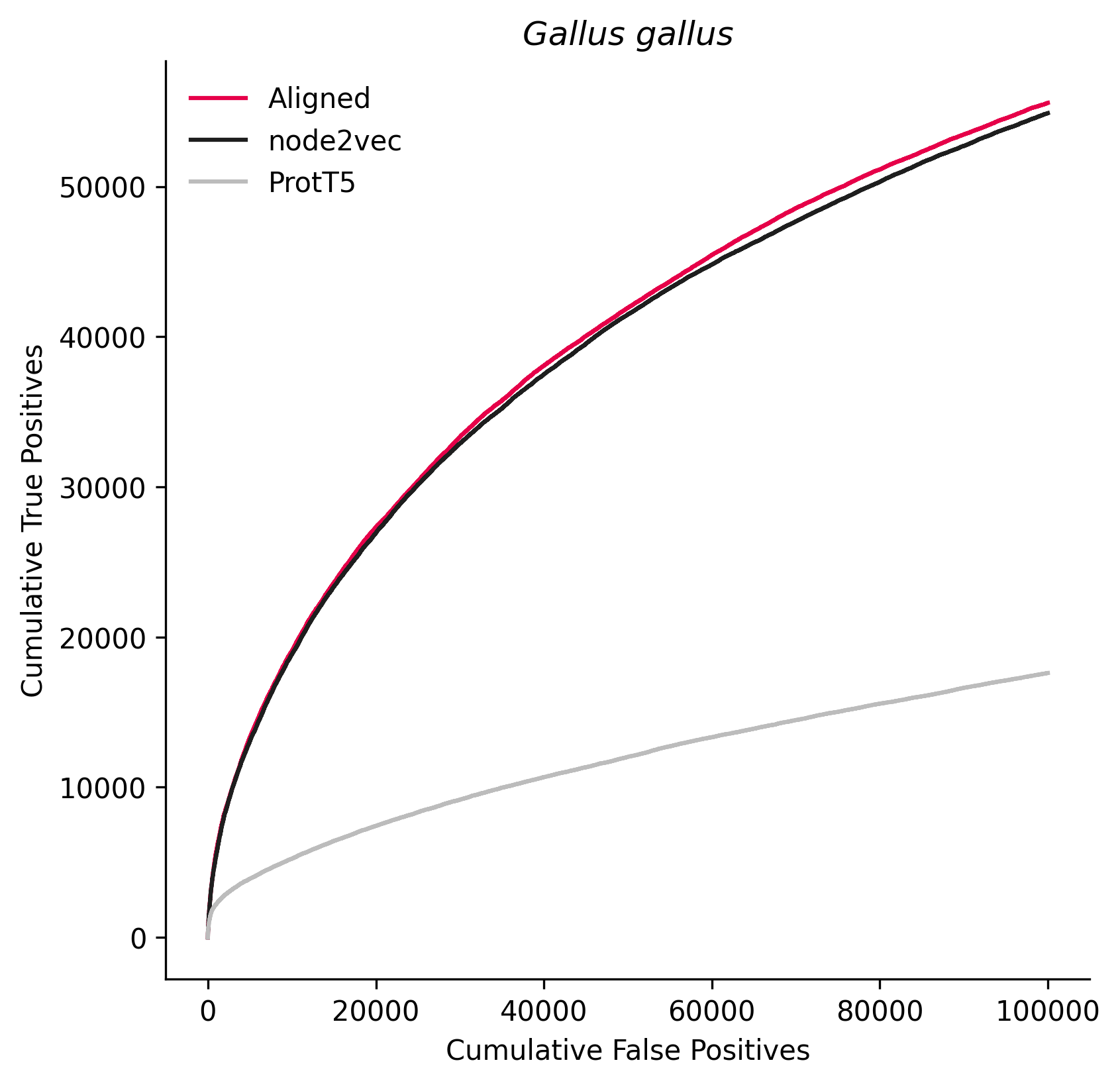

### 9103.png

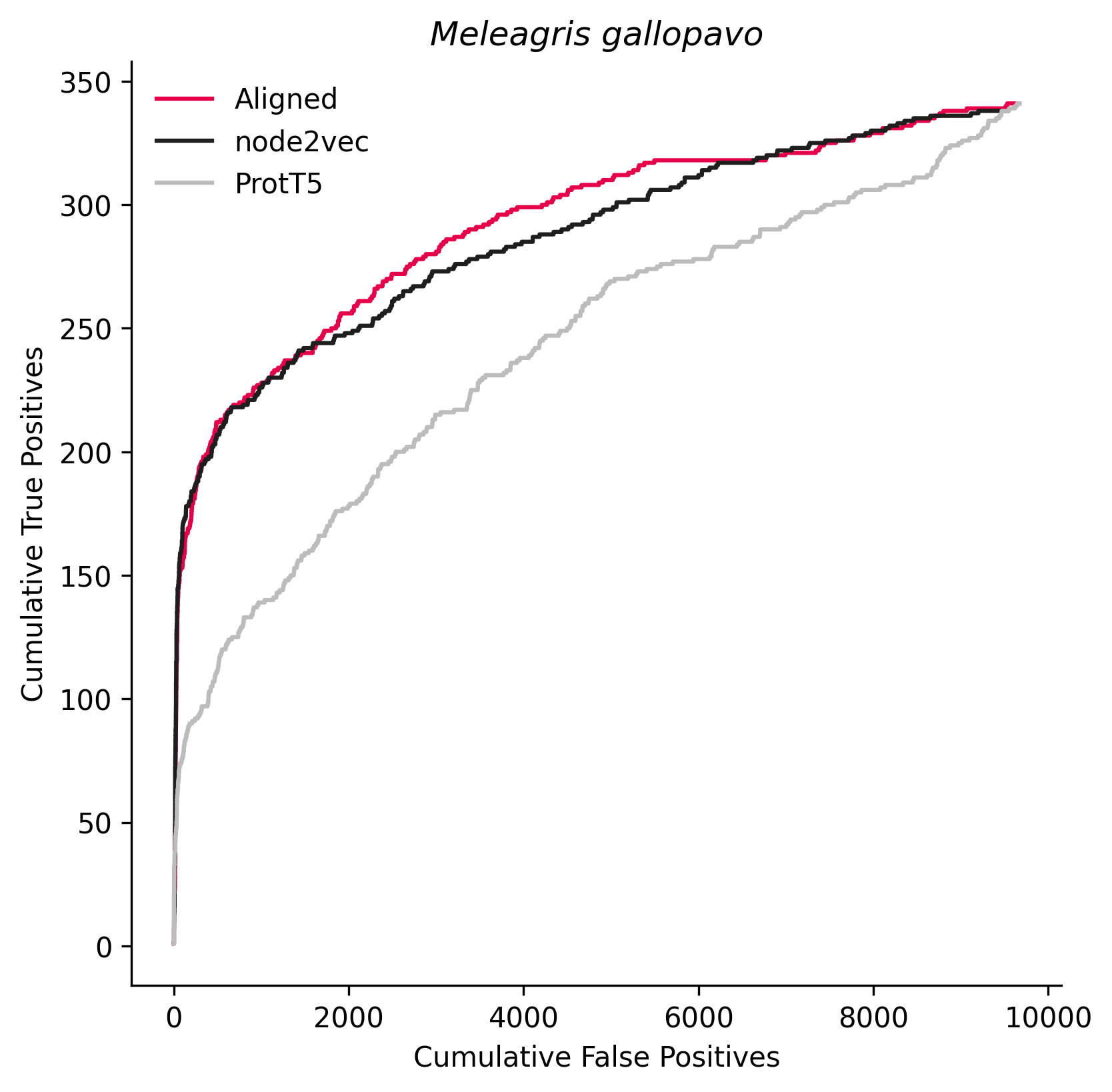

### 9135.png

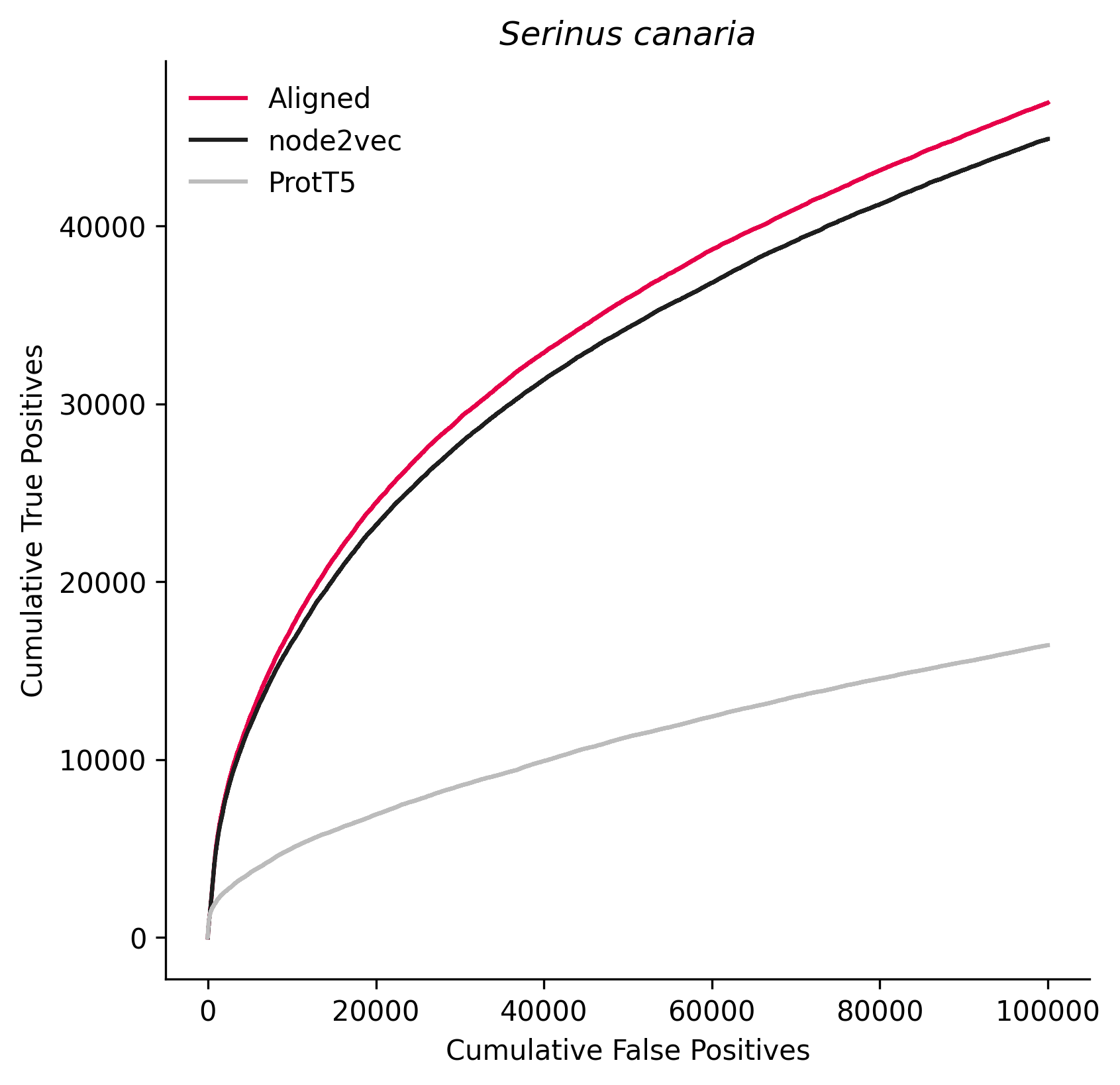

### 9541.png

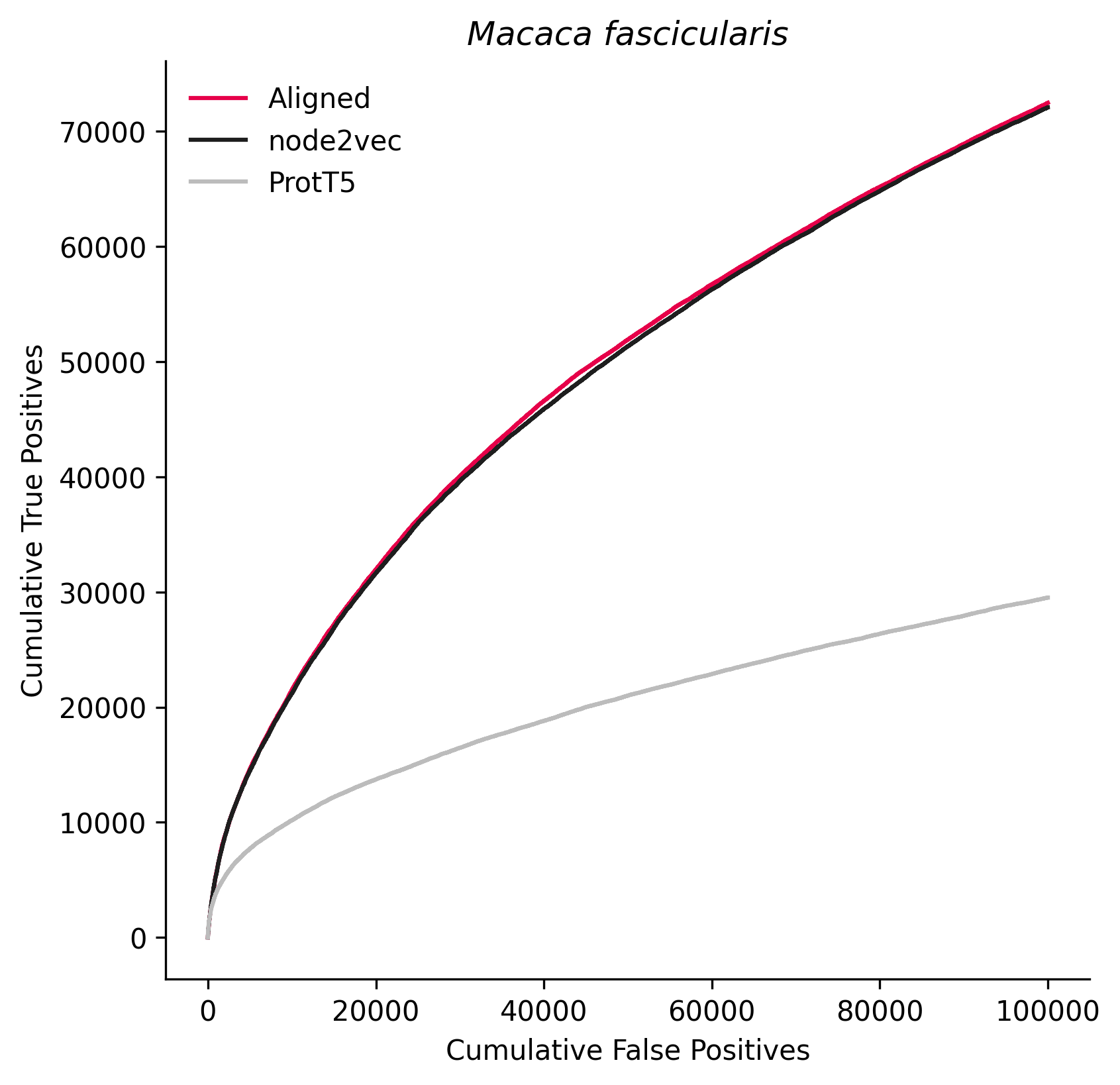

### 9601.png

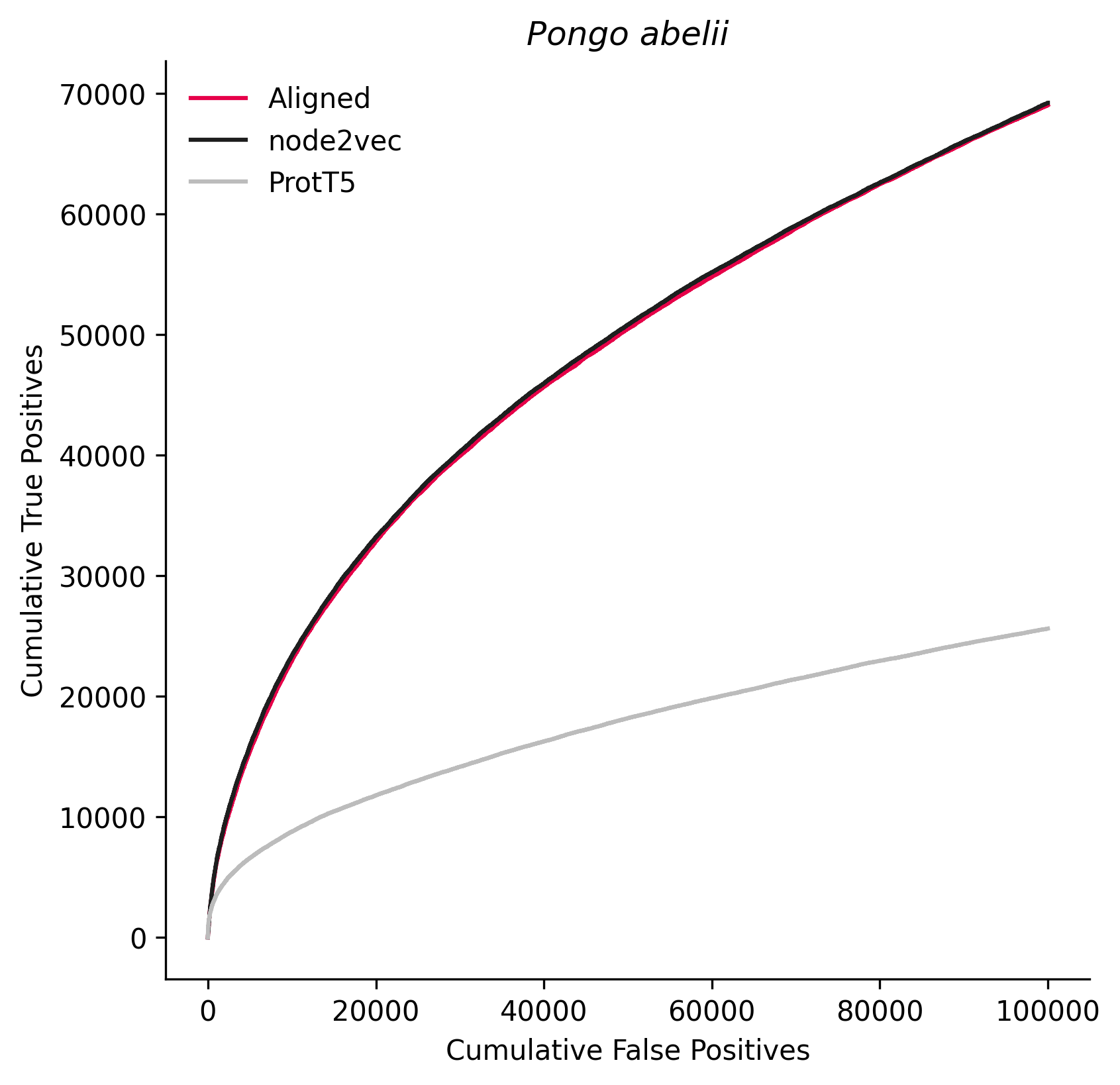

### 9755.png

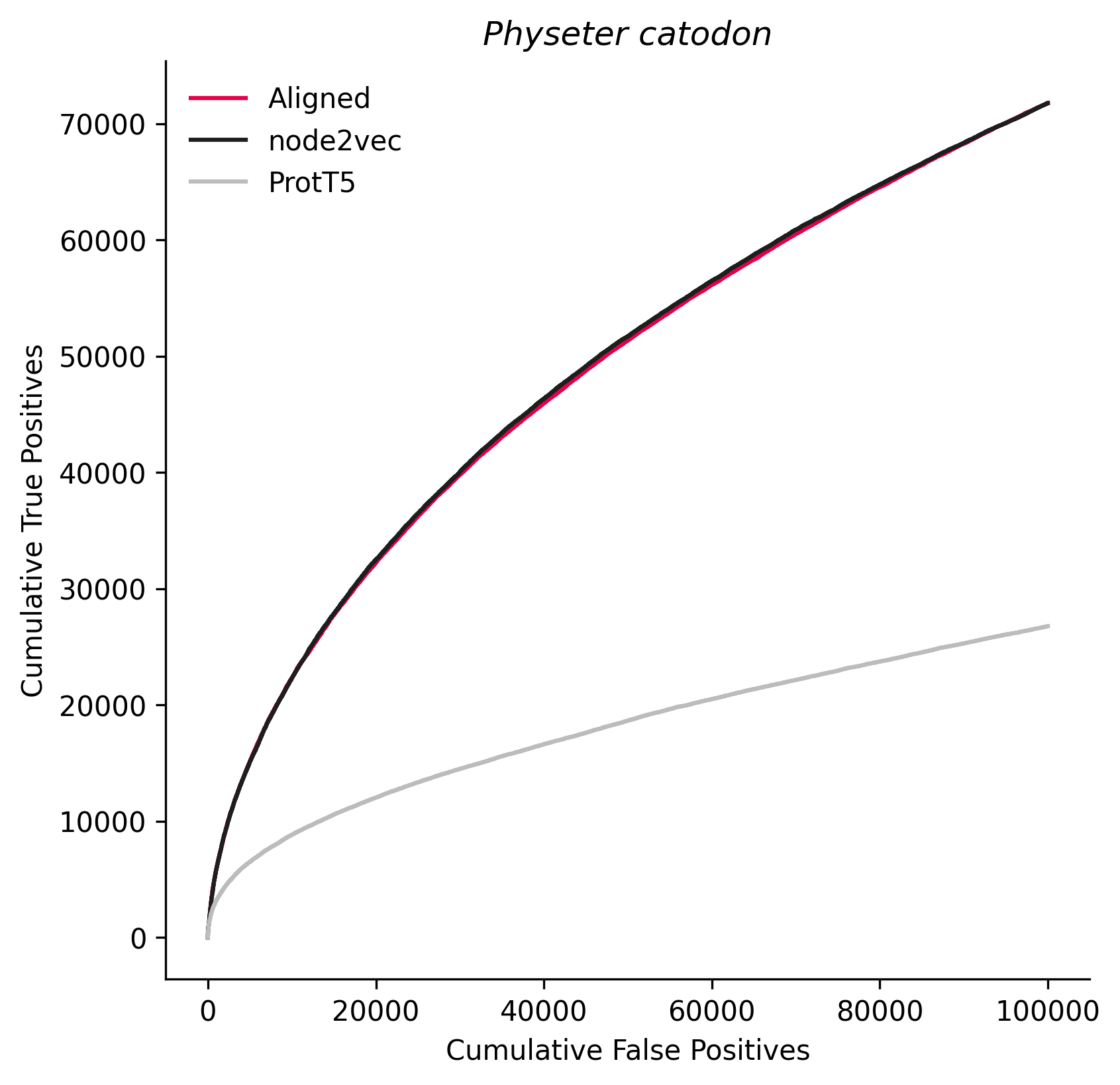

### 10089.png

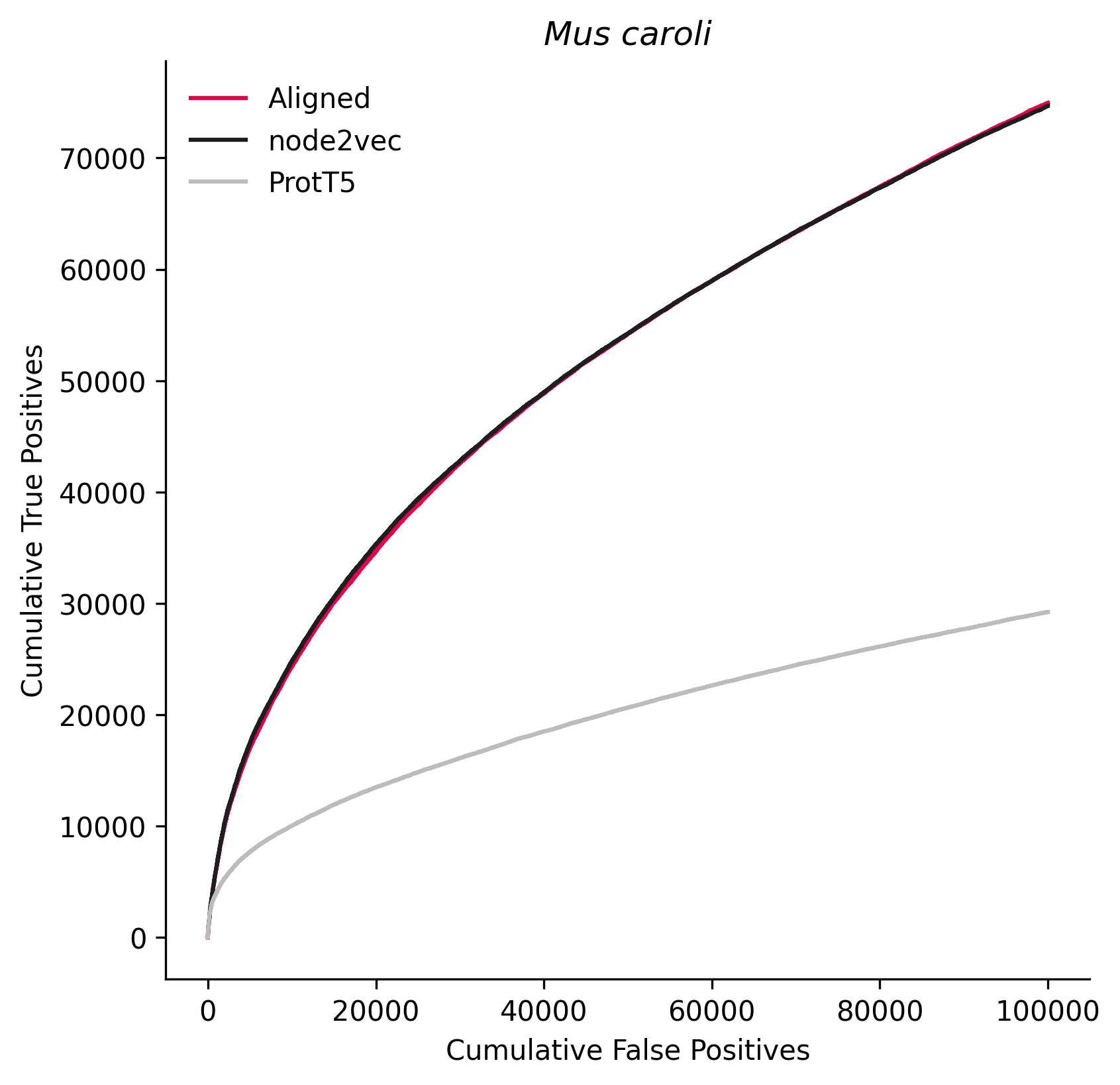

### 10181.png

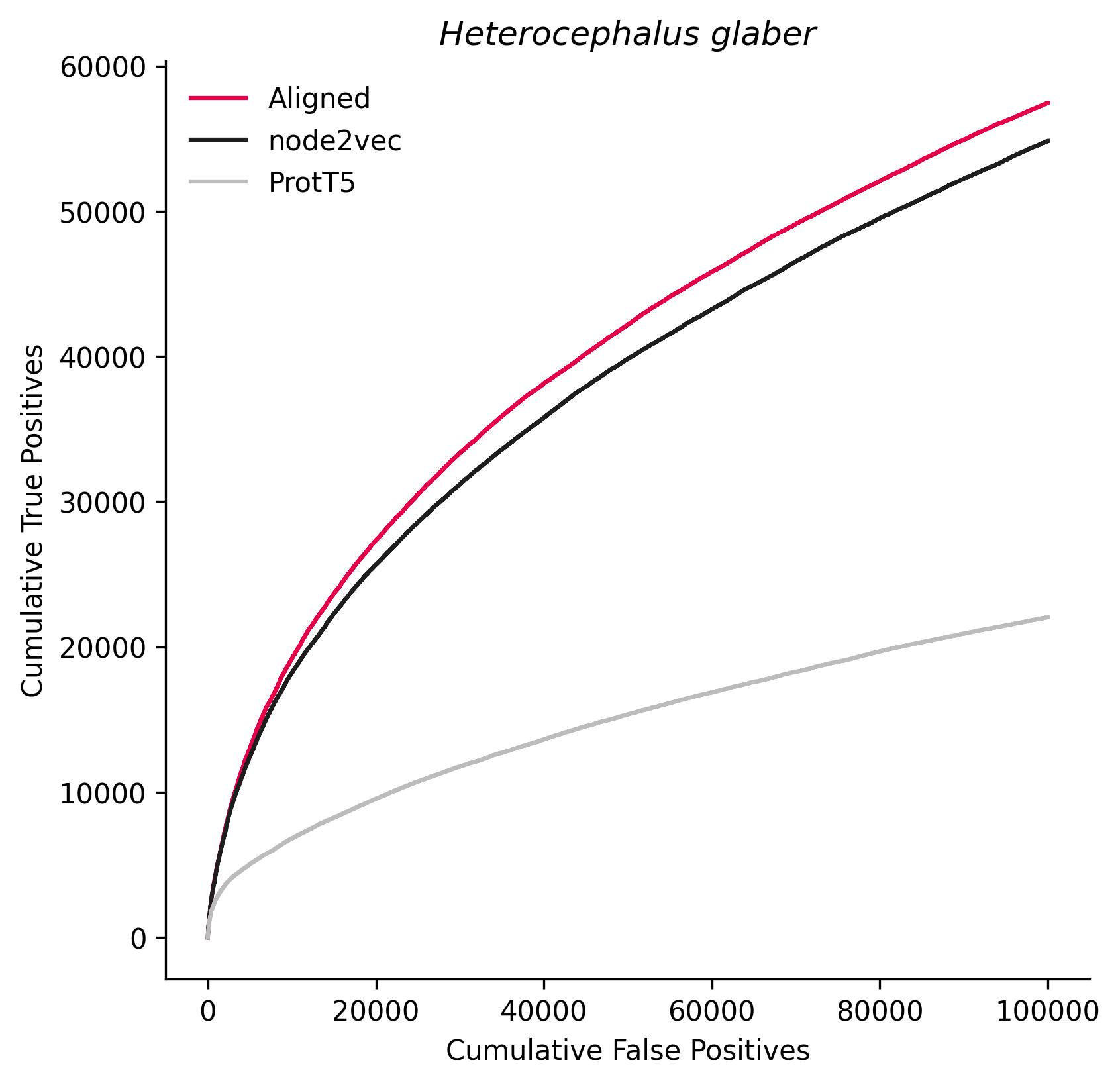

### 28377.png

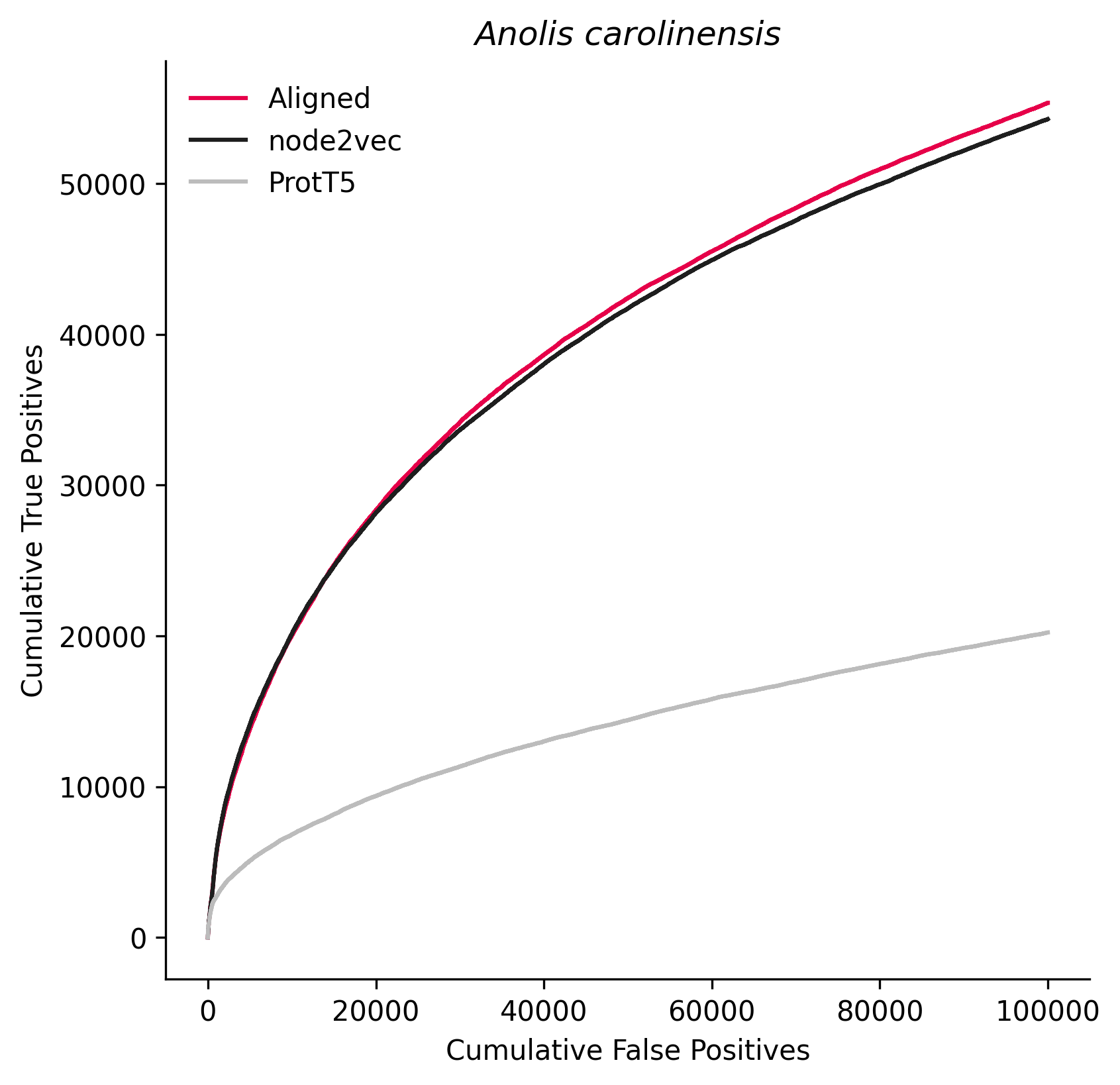

### 32264.png

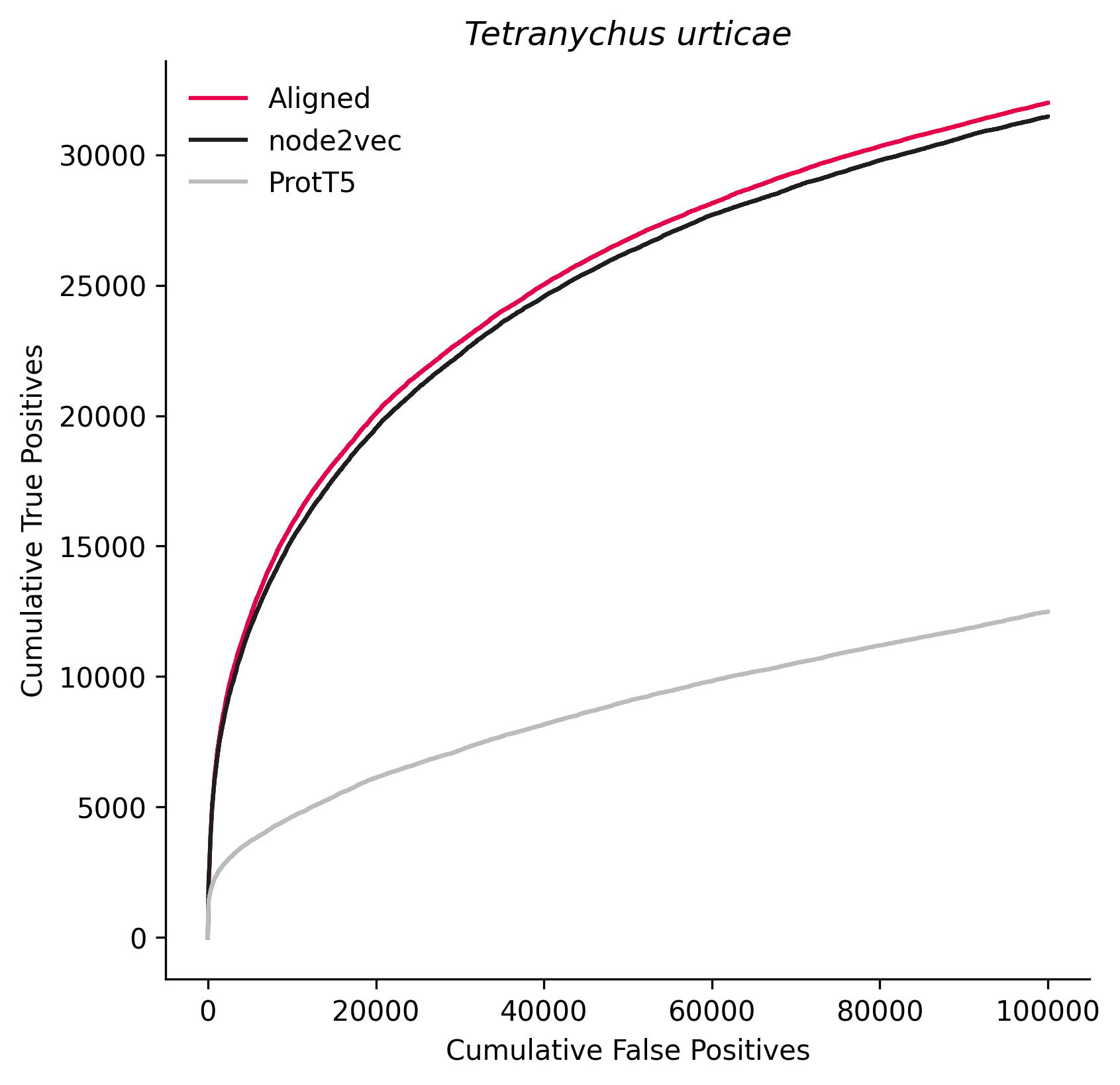

### 37003.png

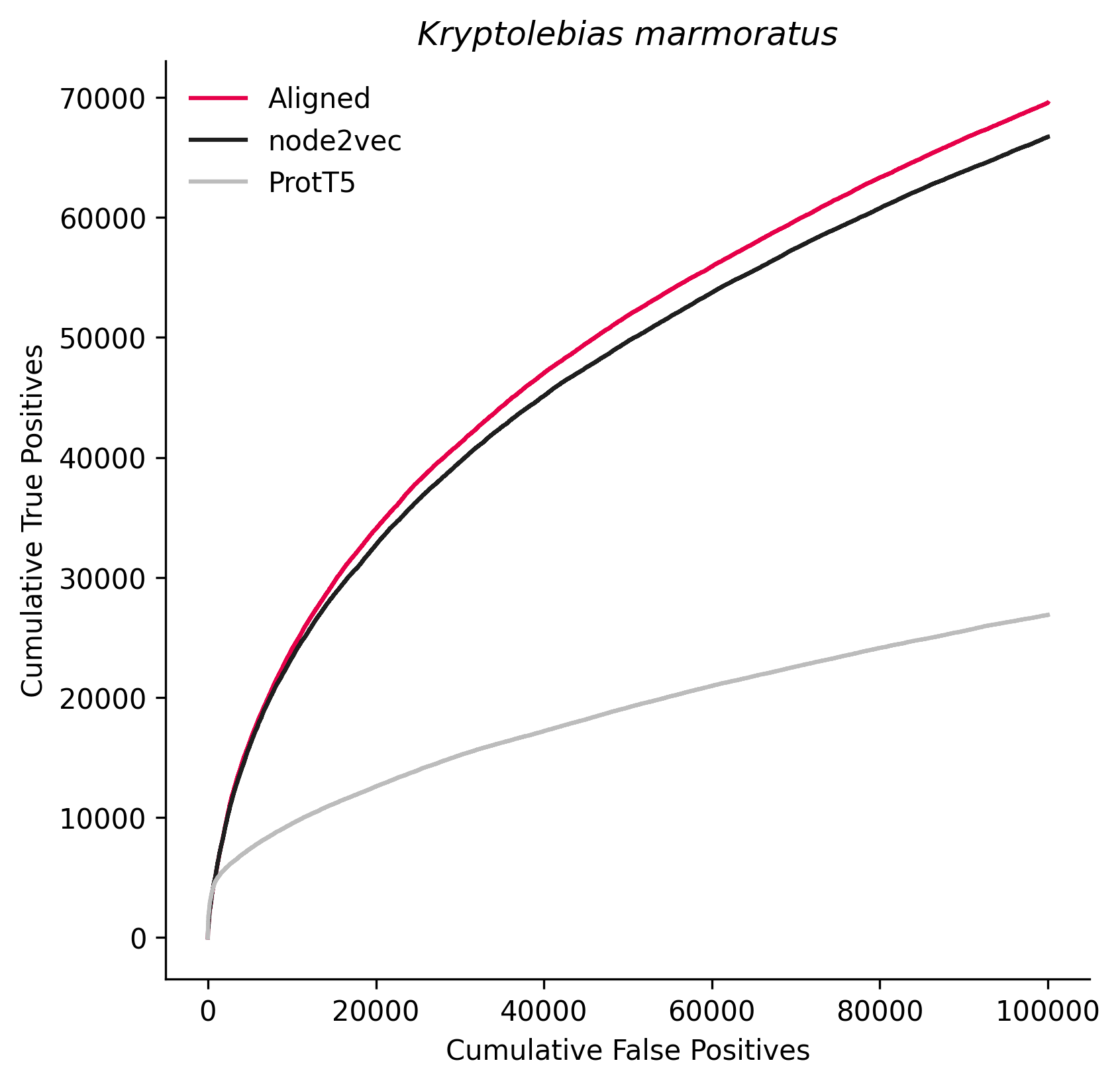

### 44689.png

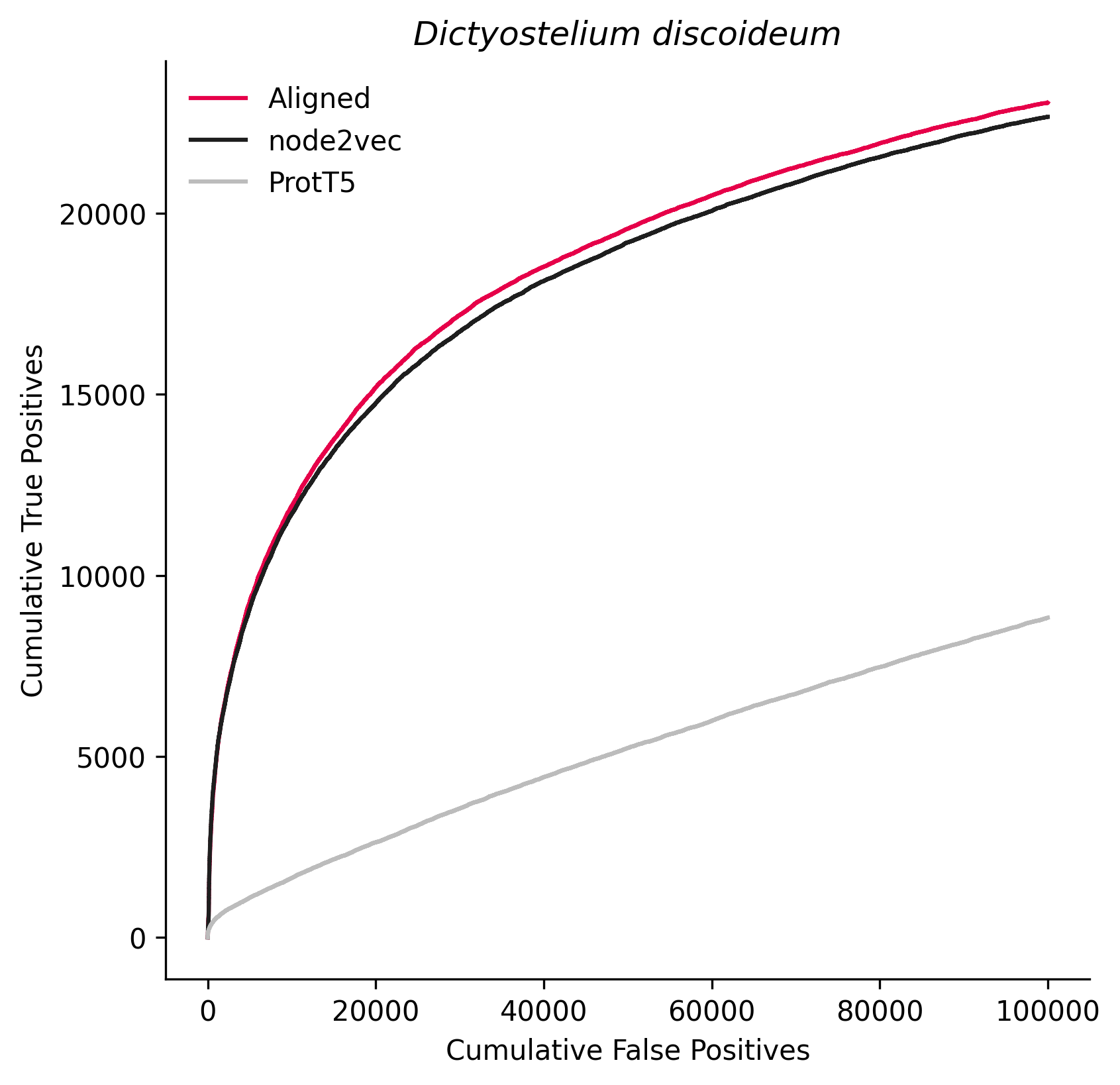

### 46245.png

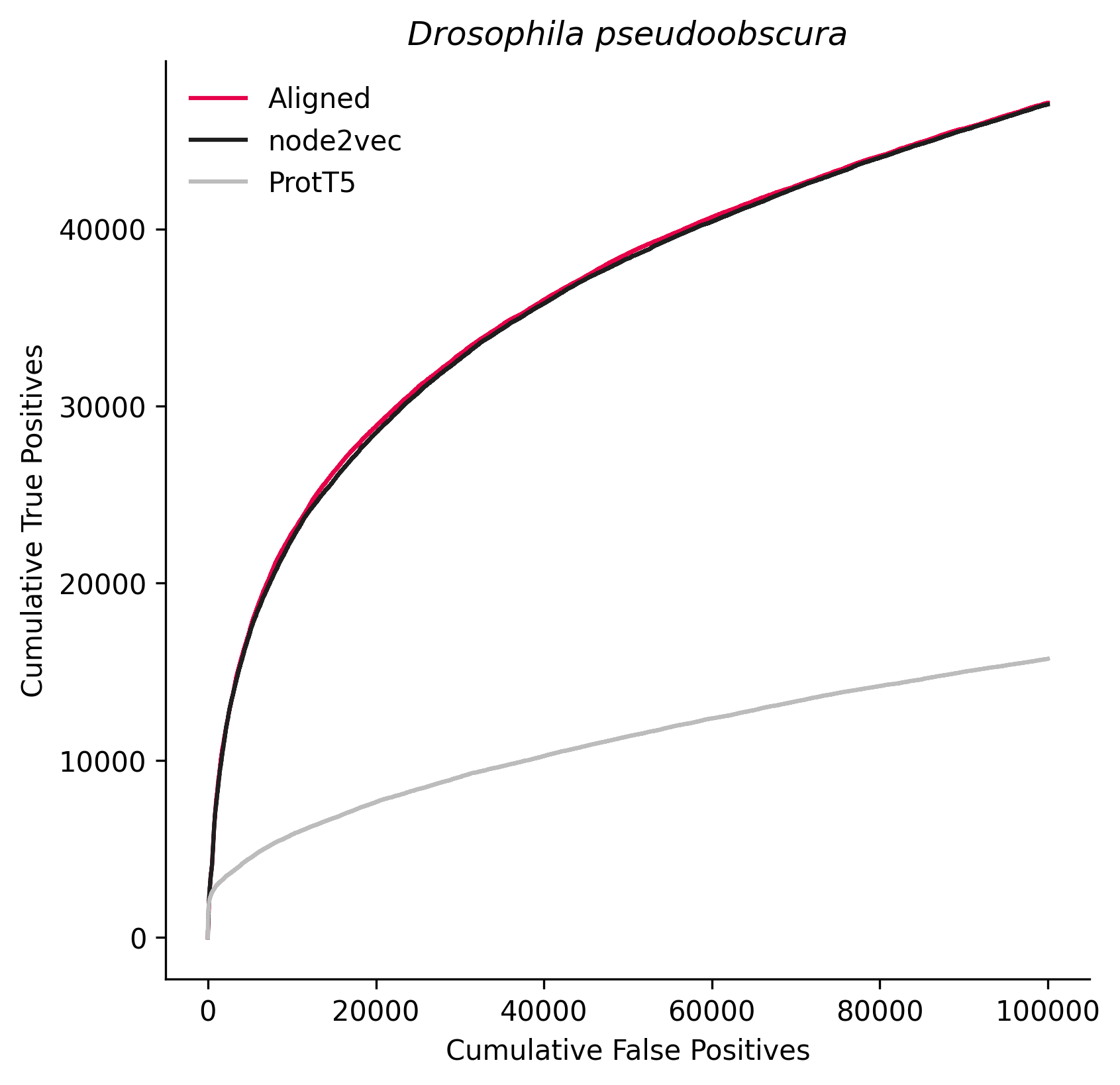

### 49451.png

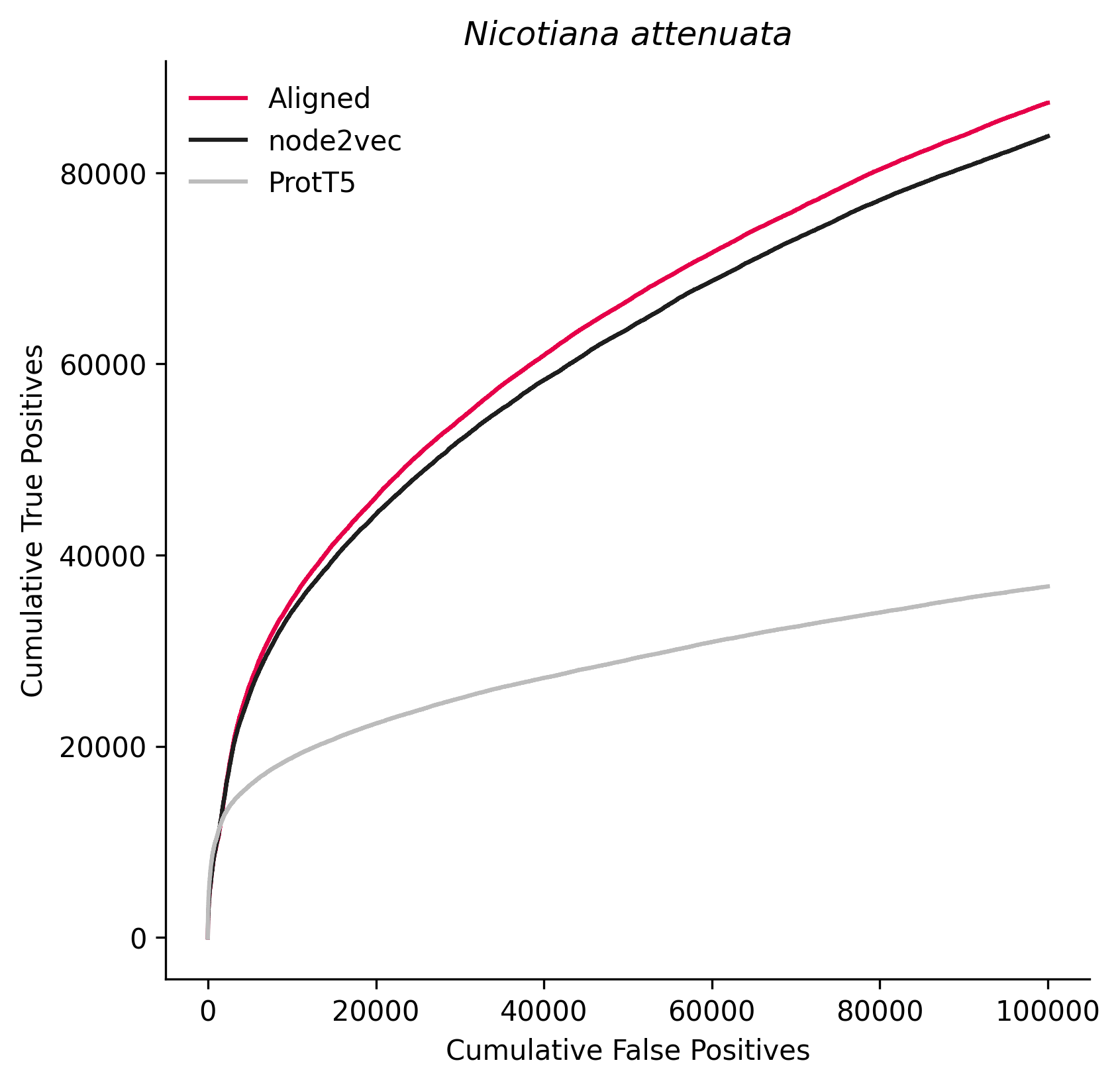

### 50429.png

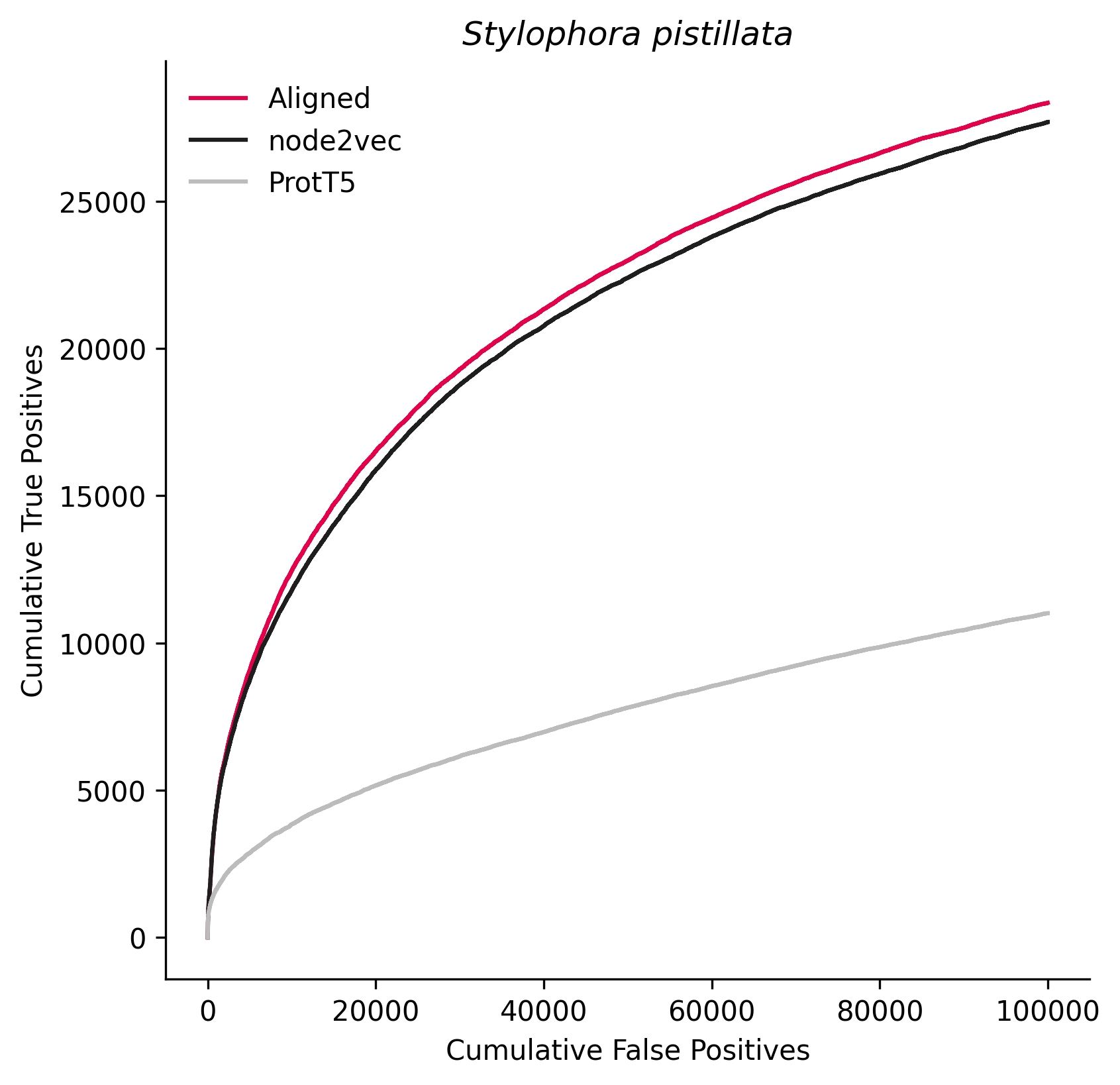

### 51240.png

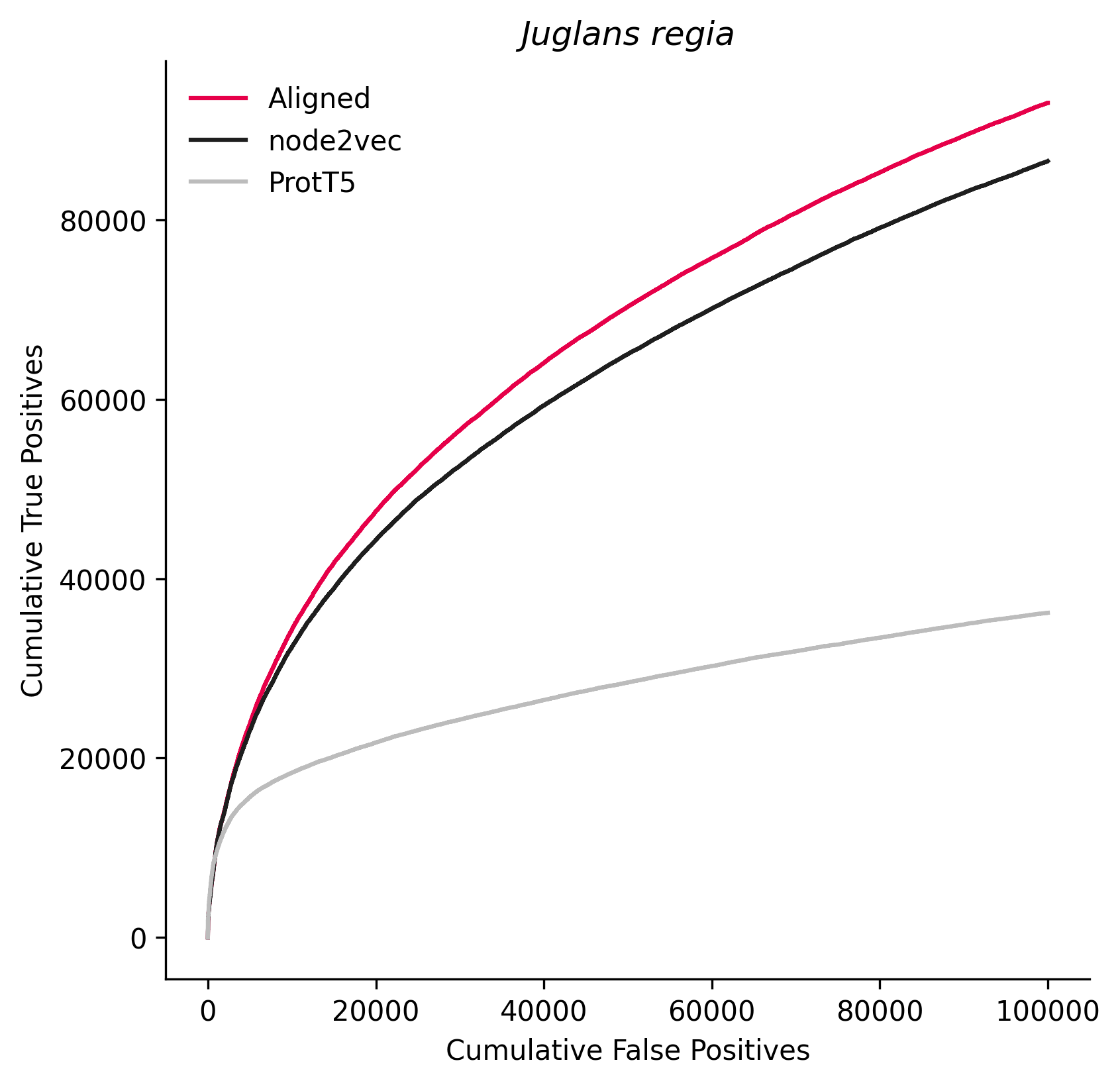

### 61621.png

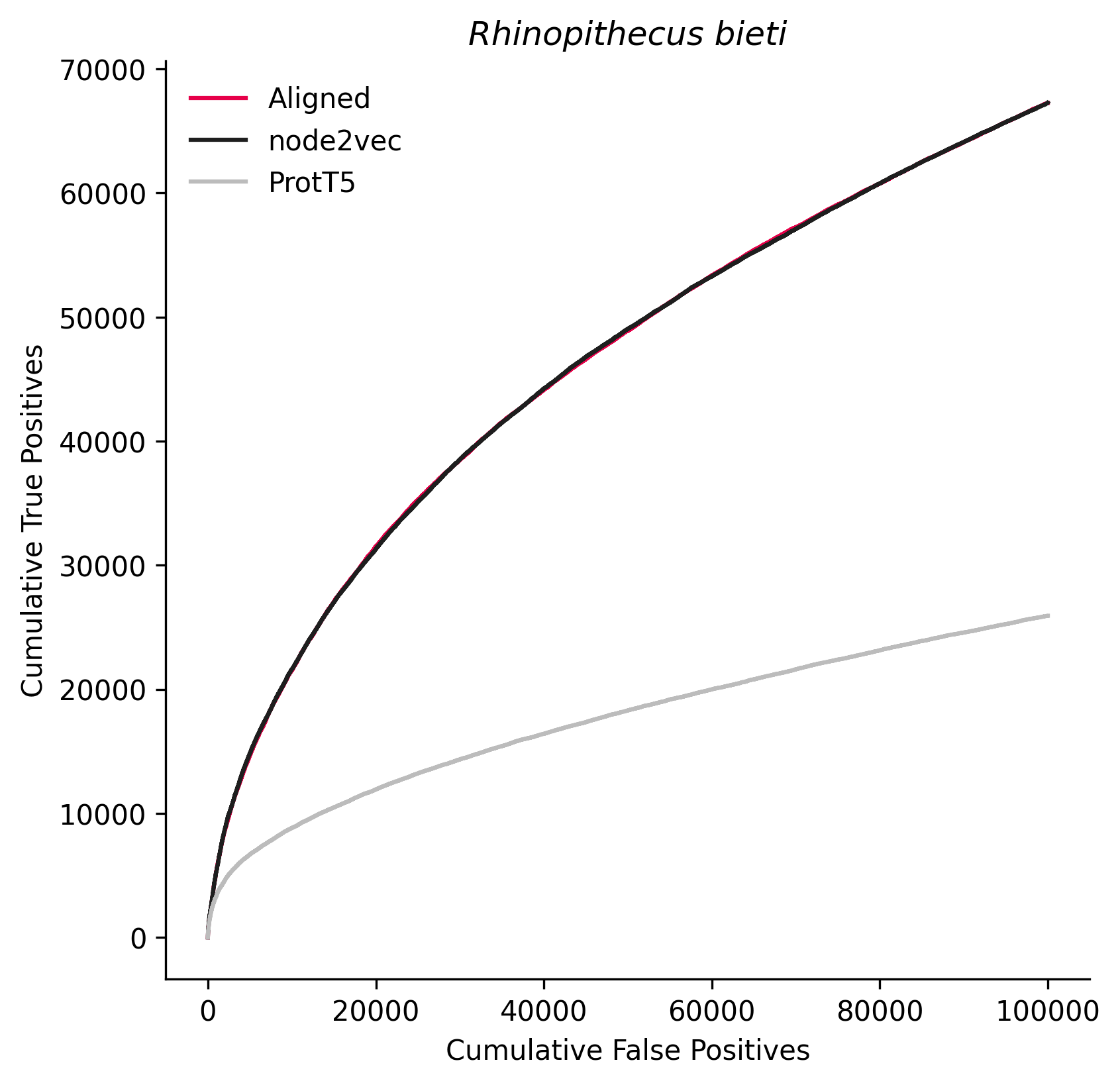

### 71139.png

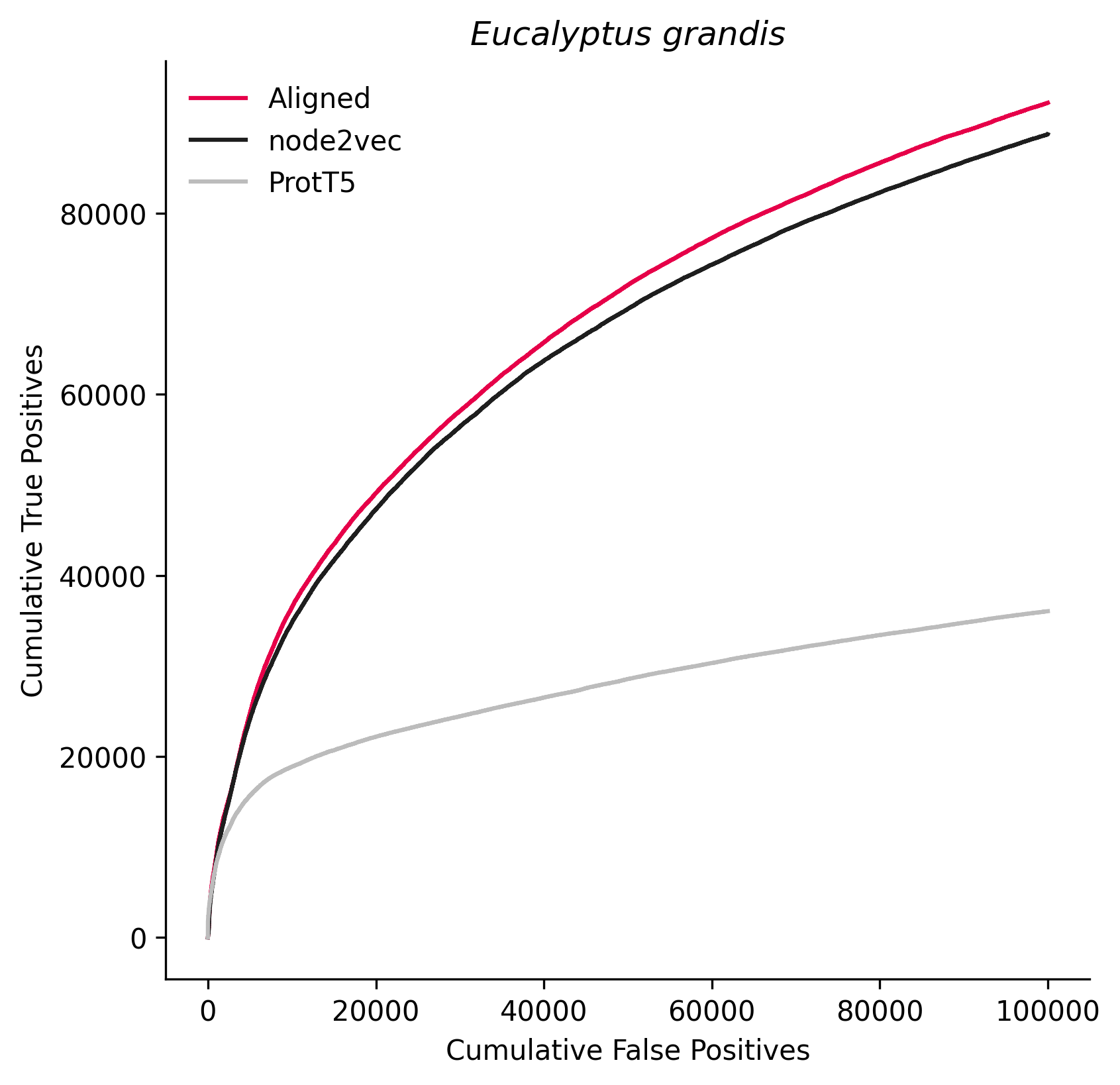

### 72004.png

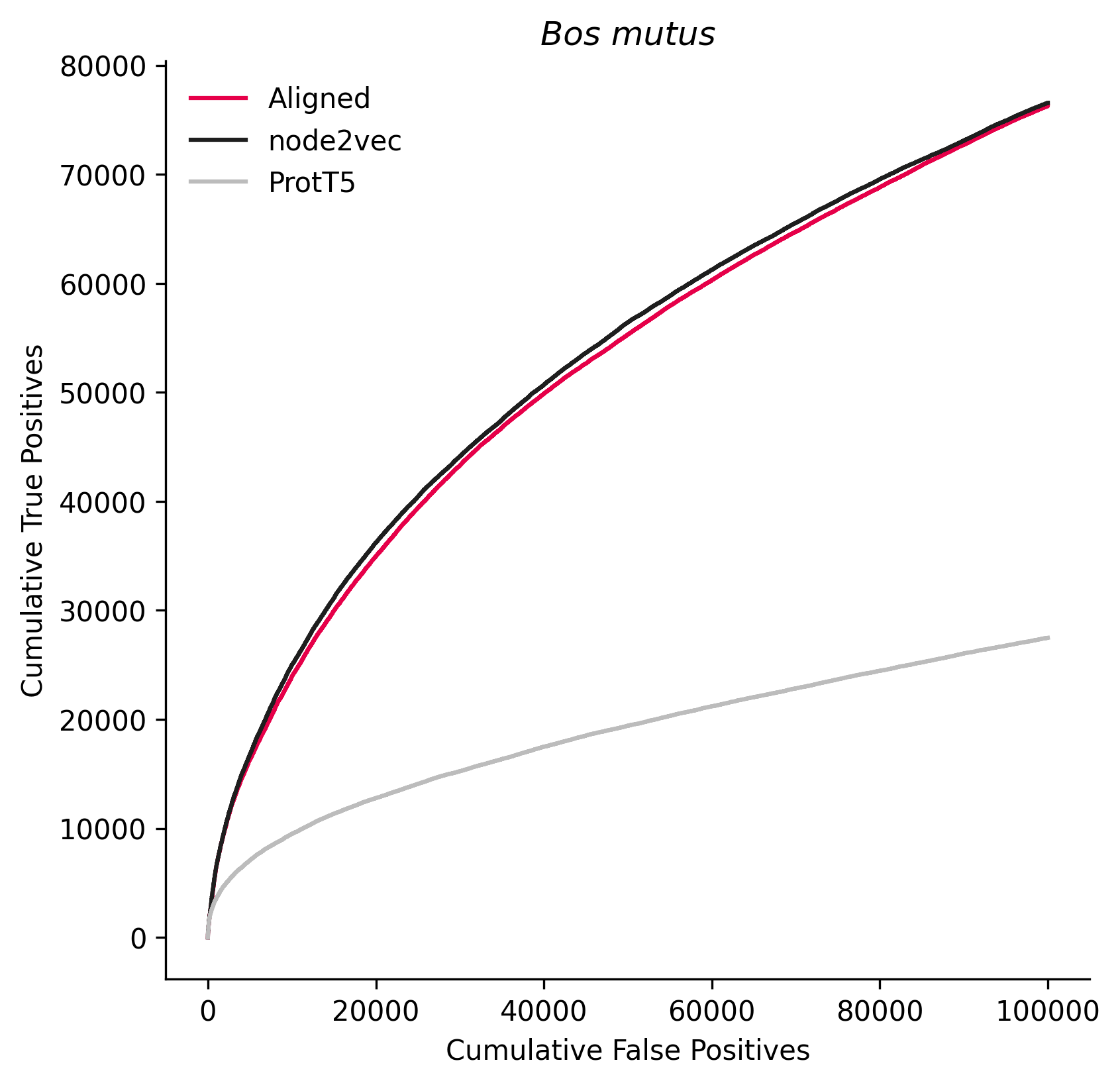

### 73239.png

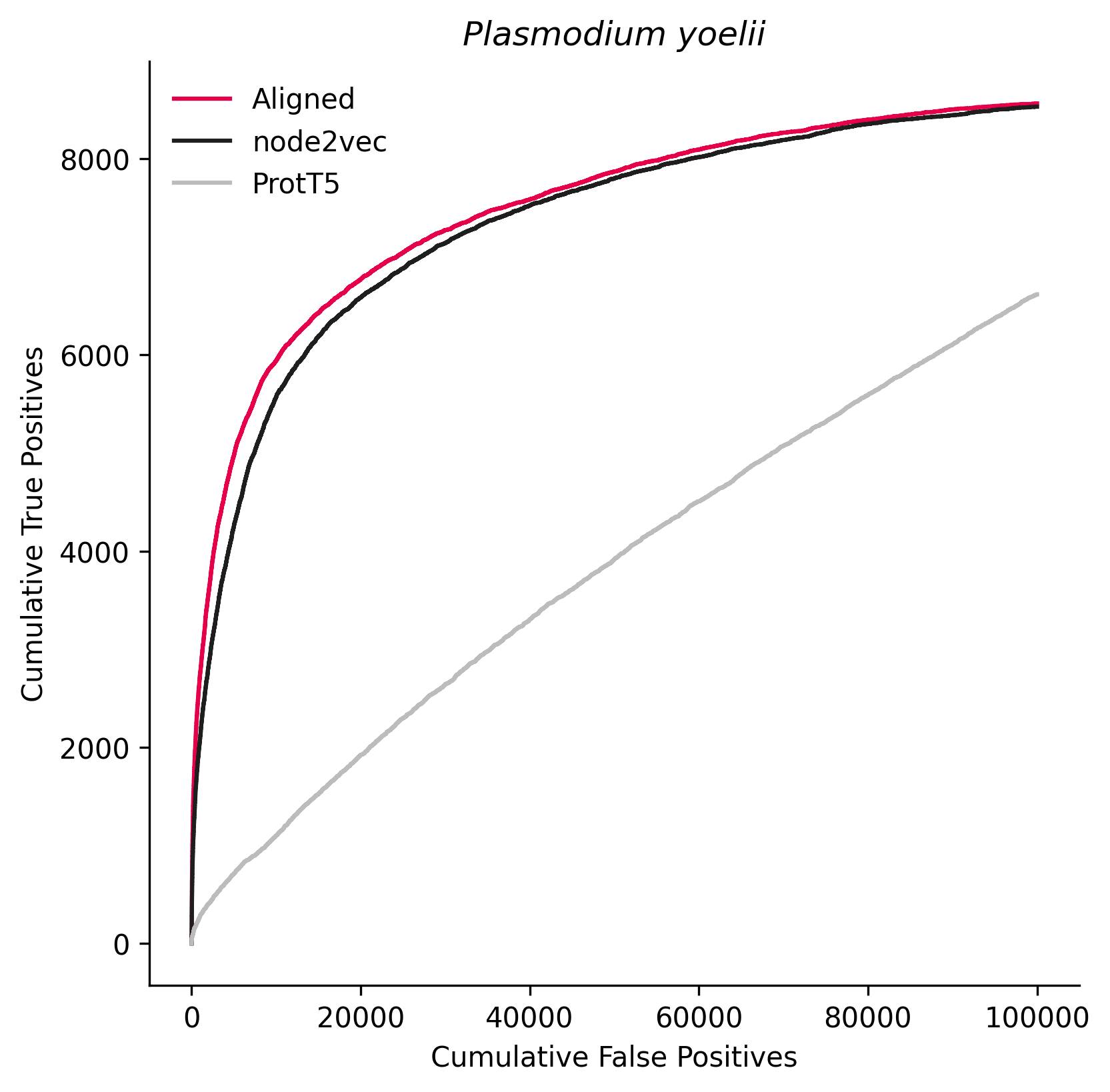

### 75366.png

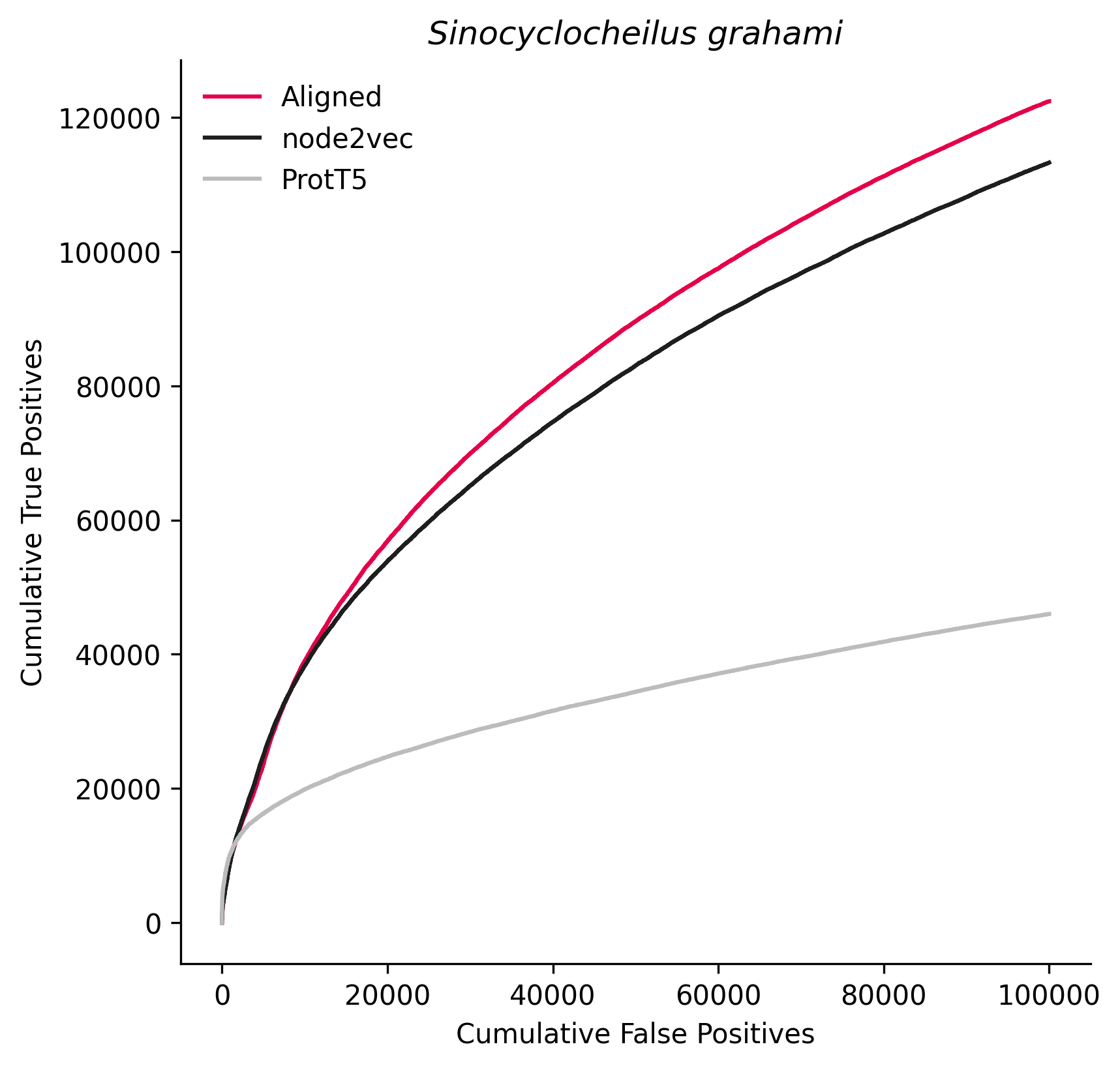

### 81972.png

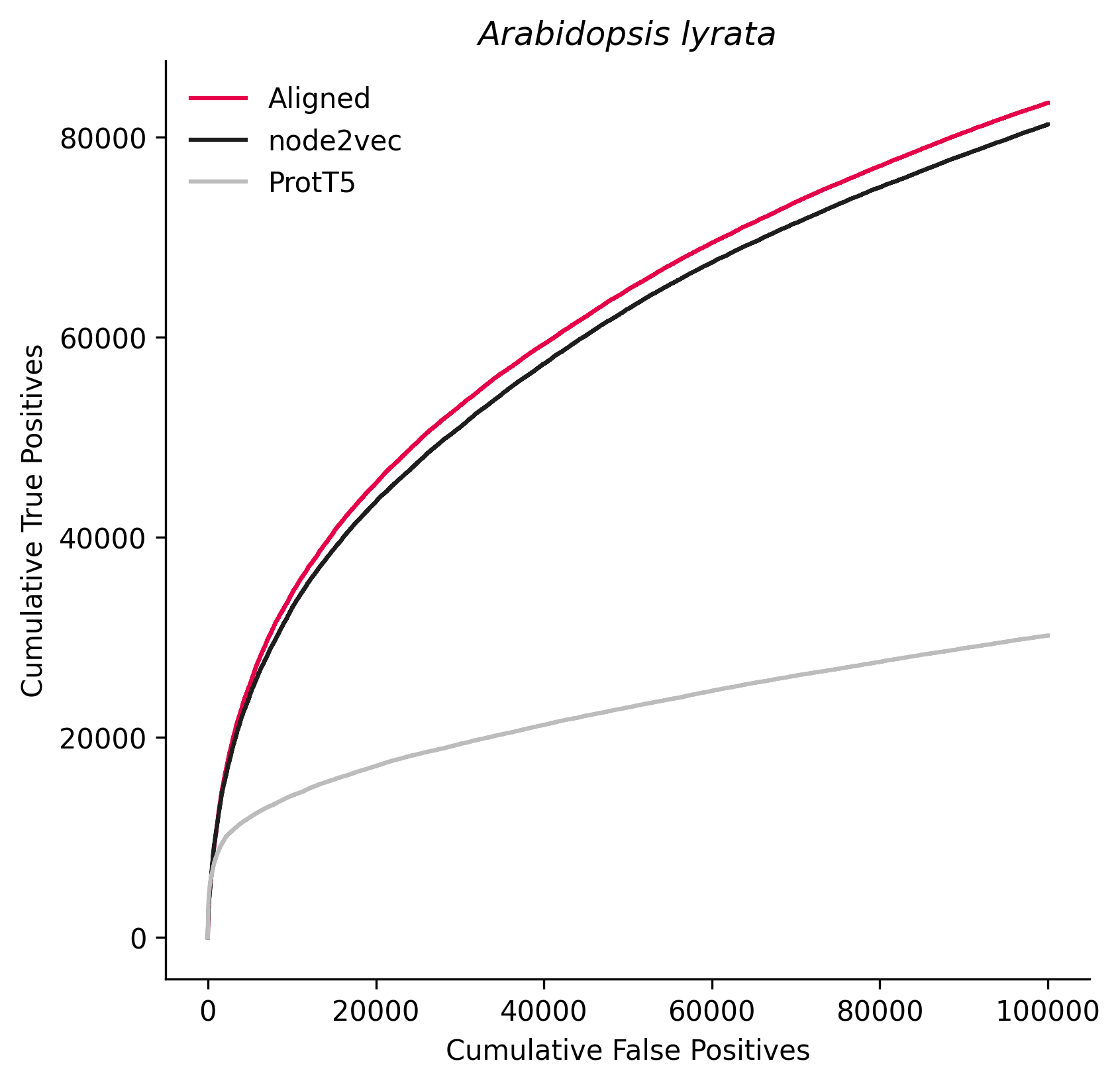

### 83772.png

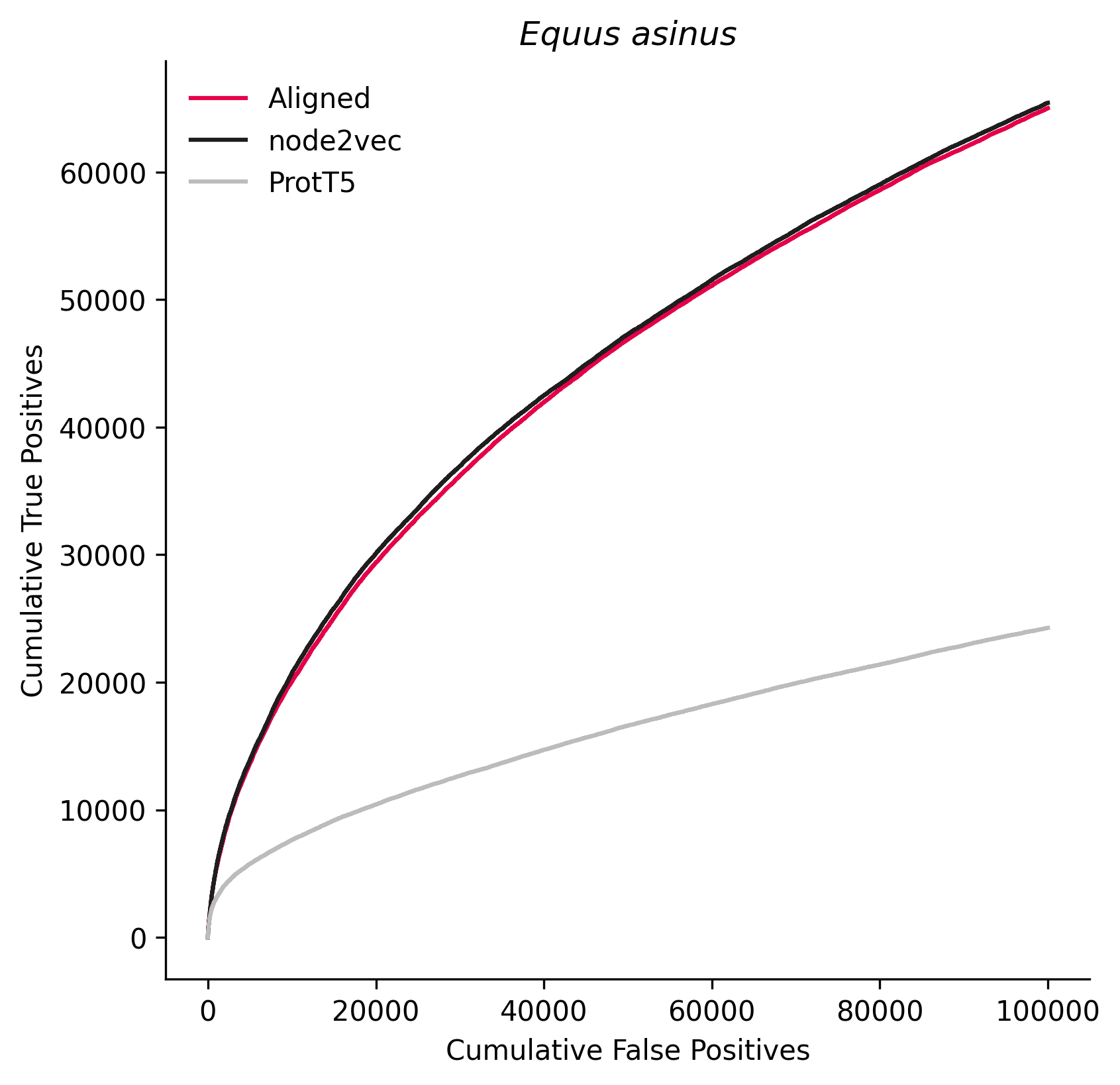

### 85681.png

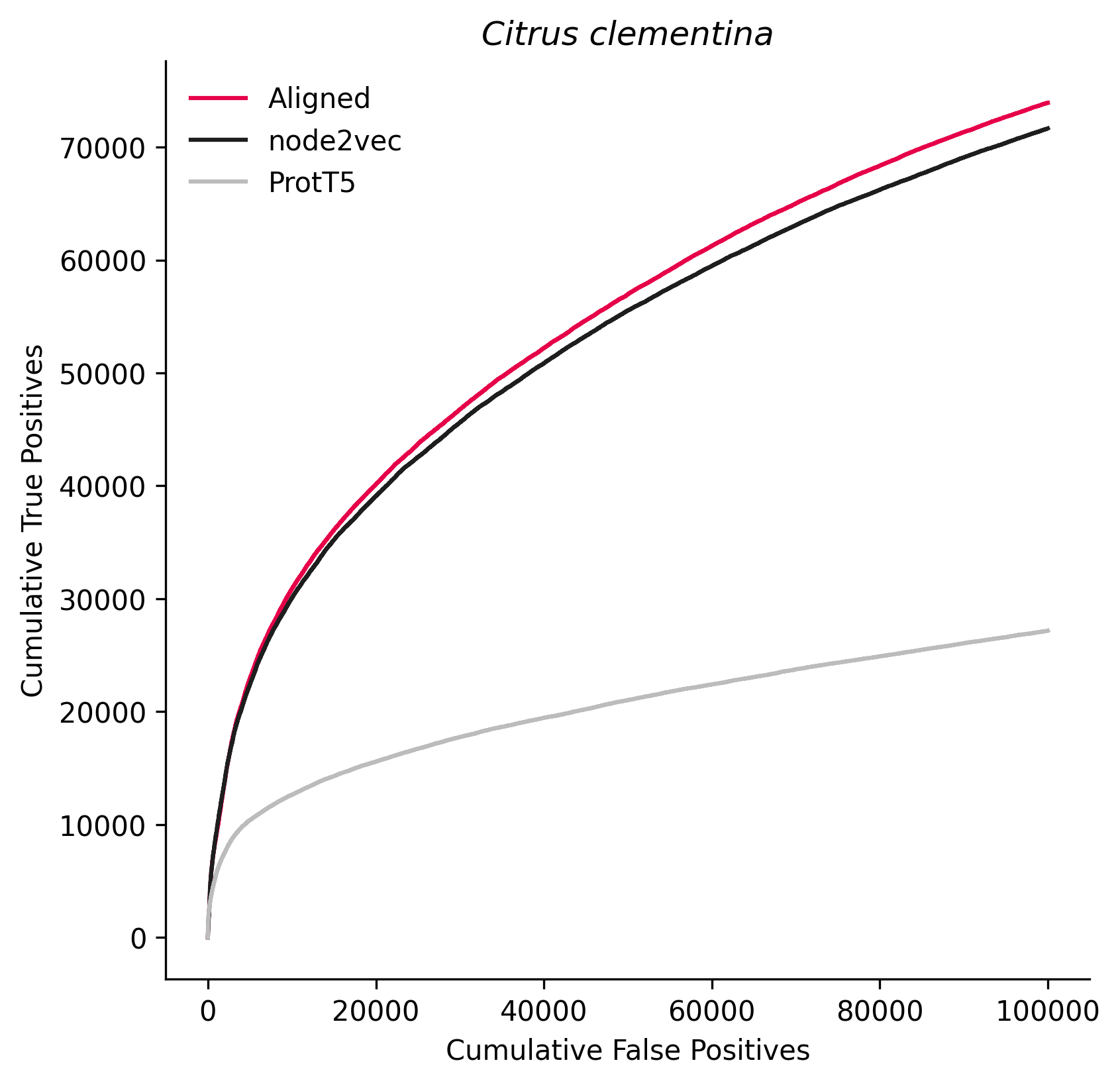

### 88036.png

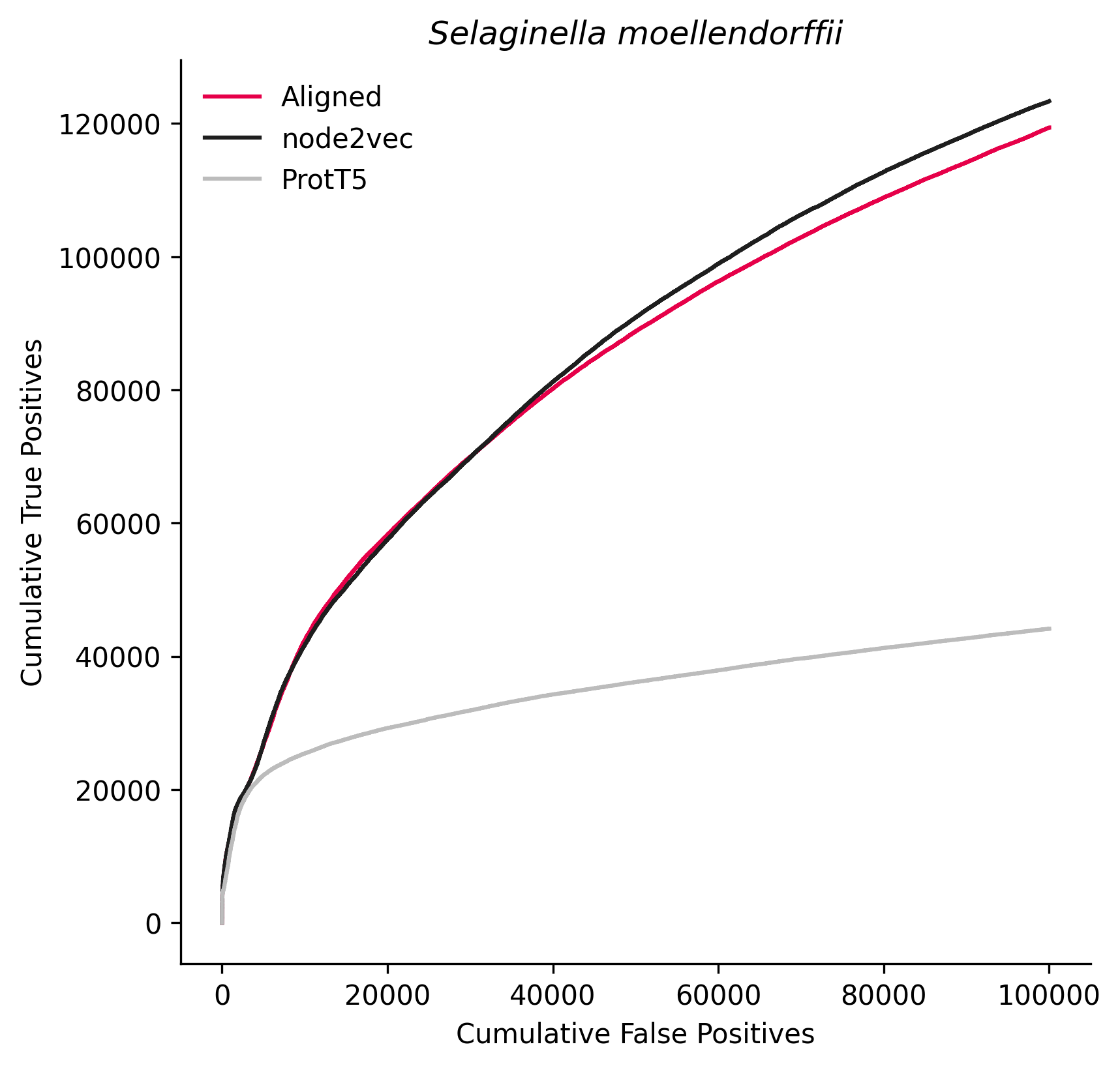

### 103695.png

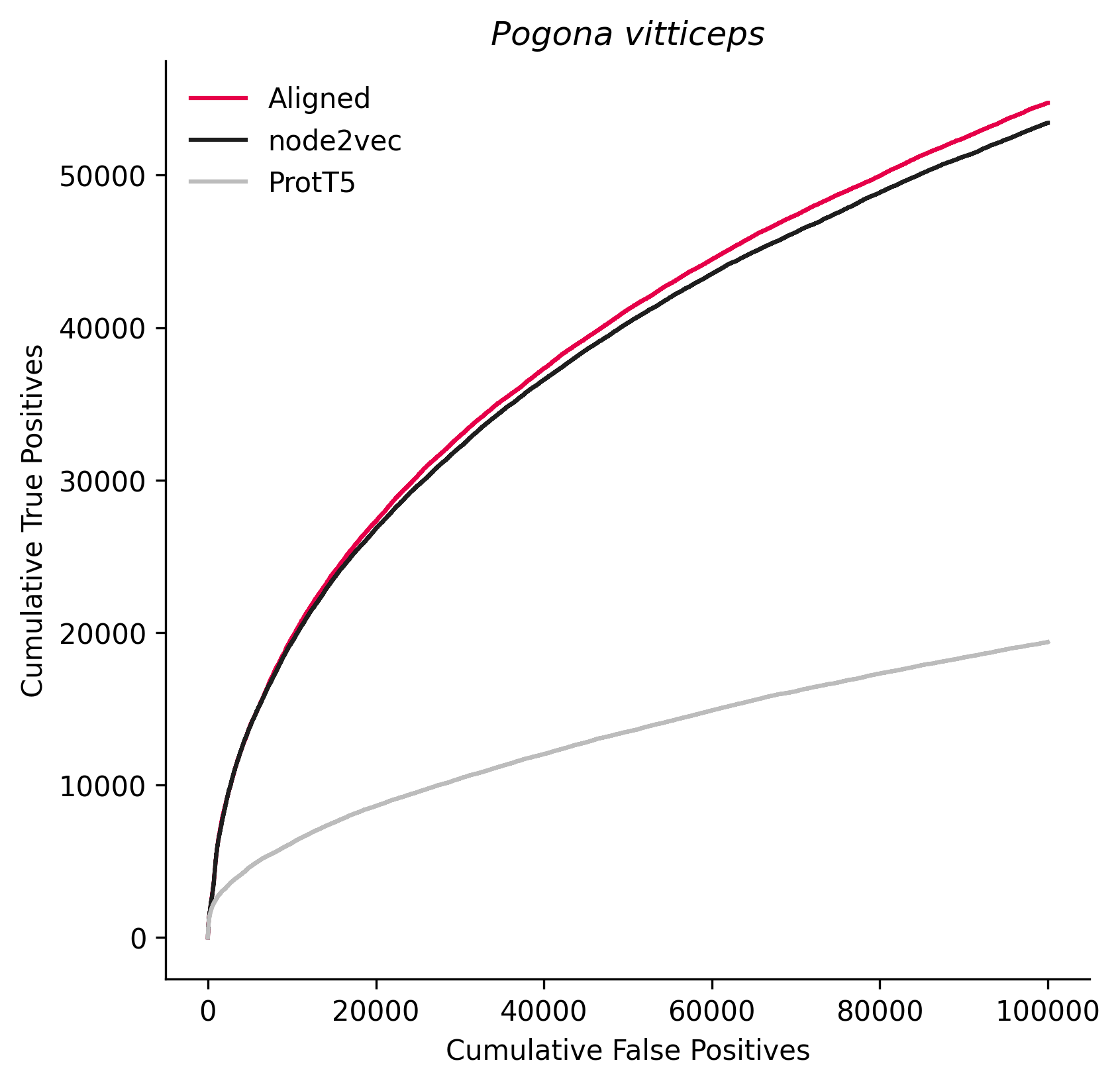

### 104421.png

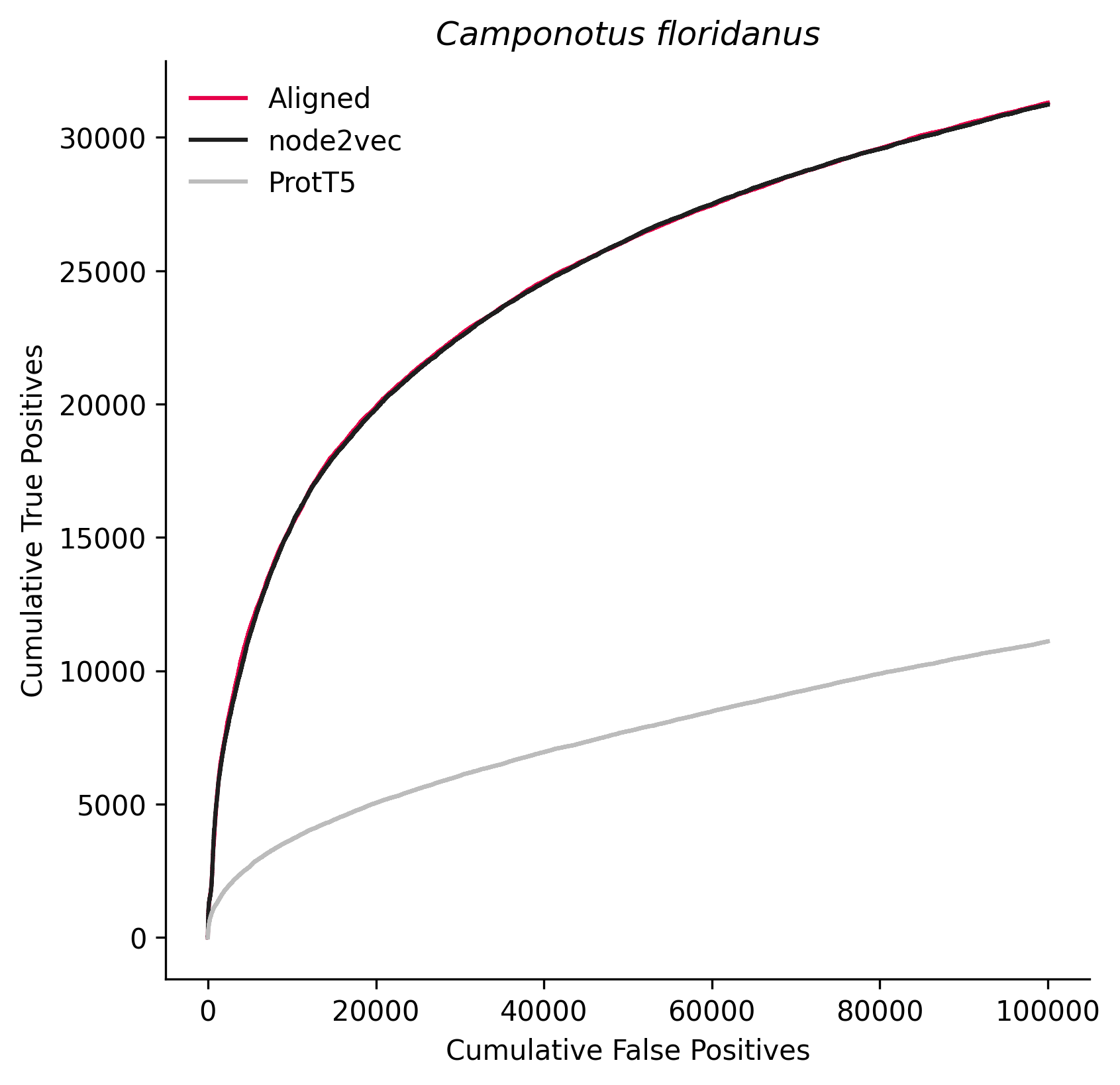

### 127582.png

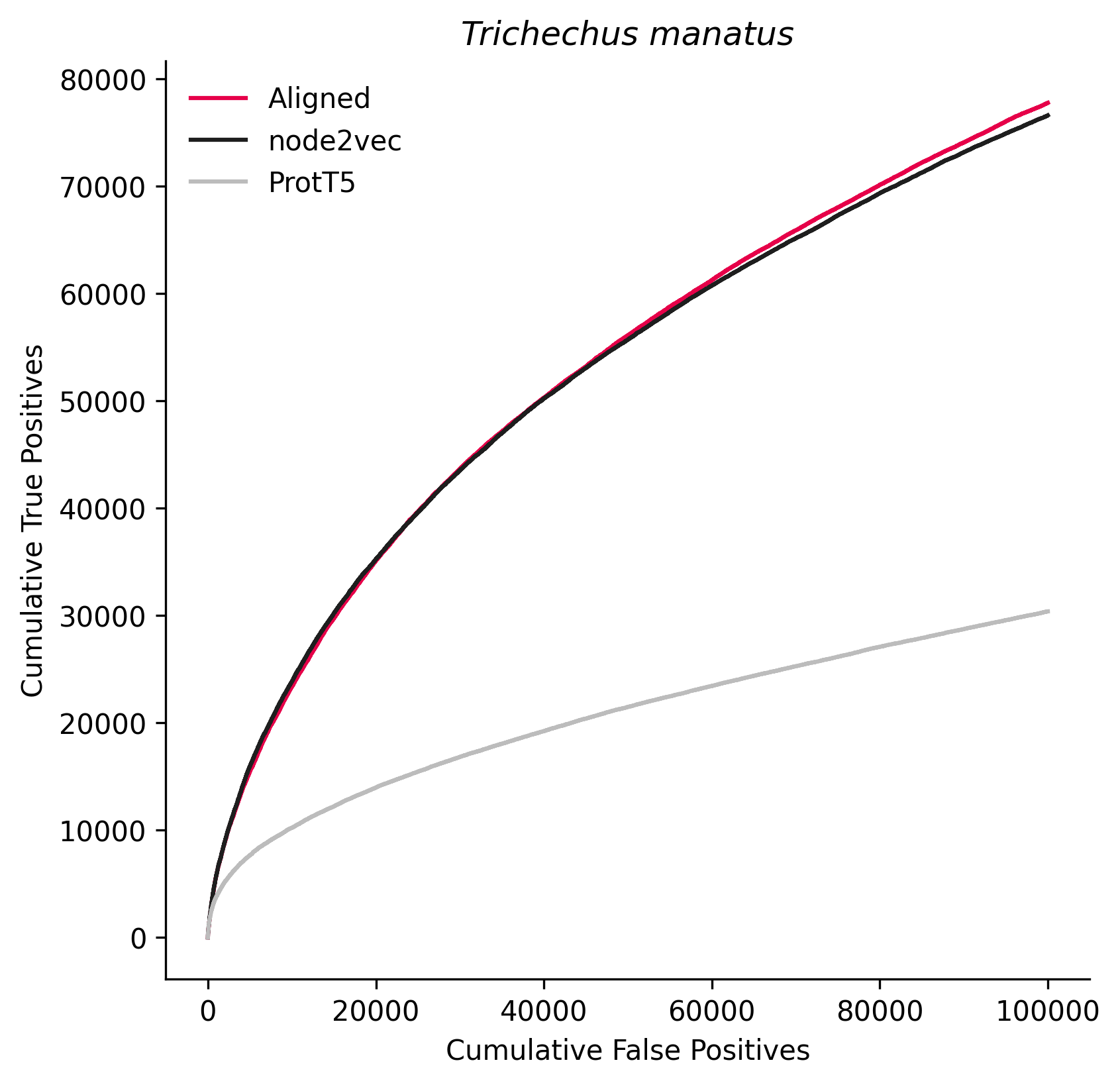

### 128390.png

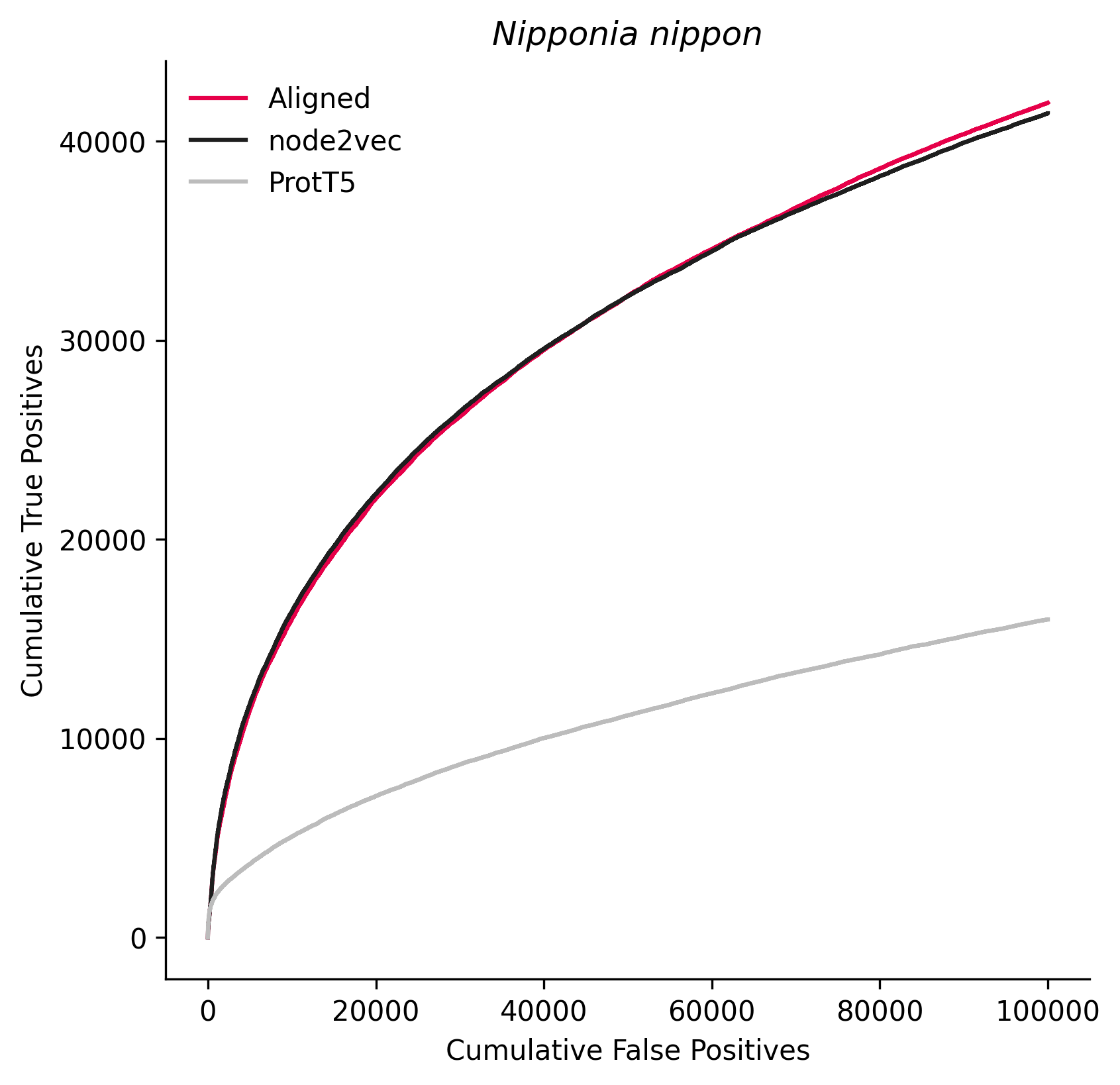

### 180498.png

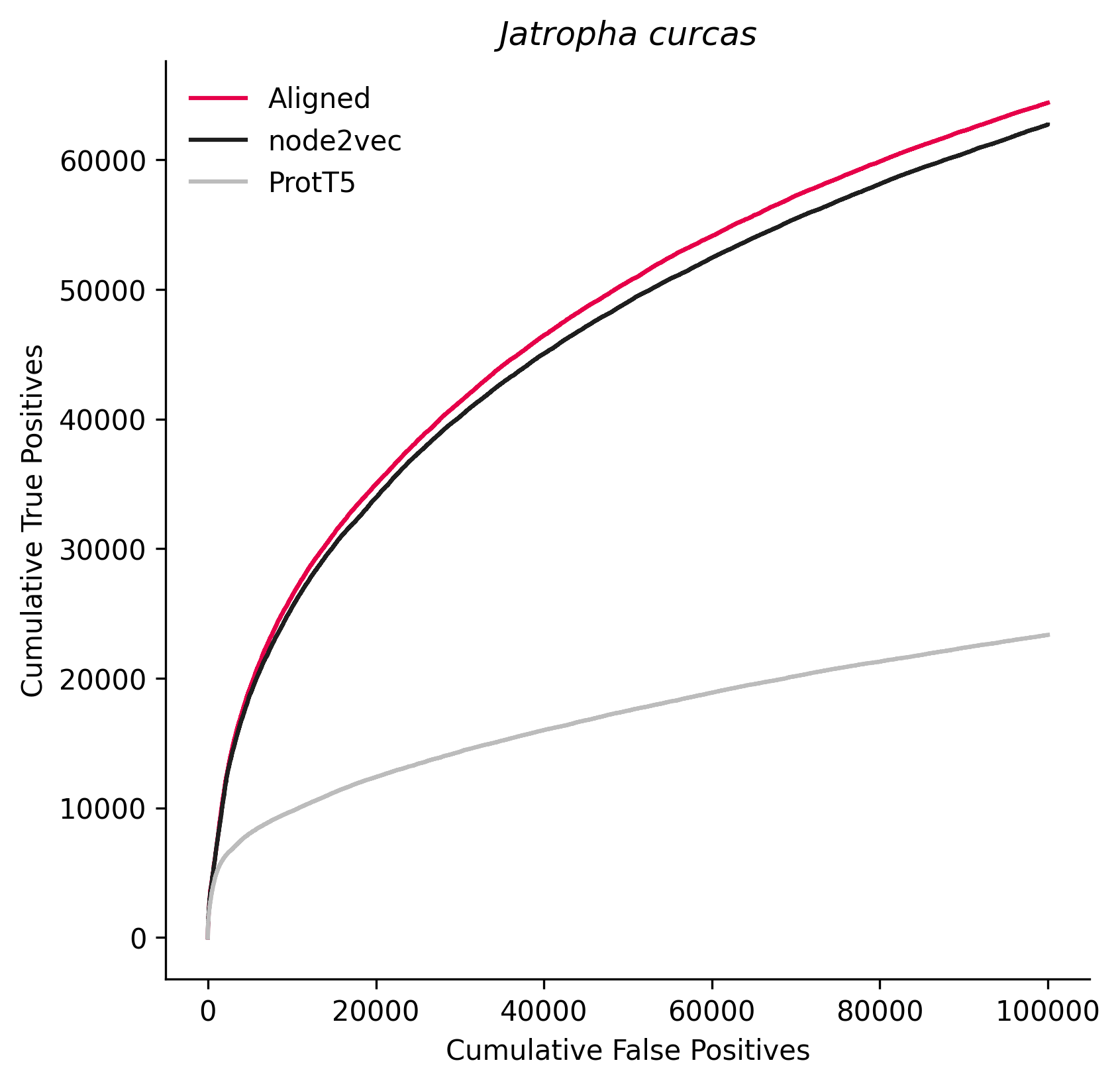

### 194338.png

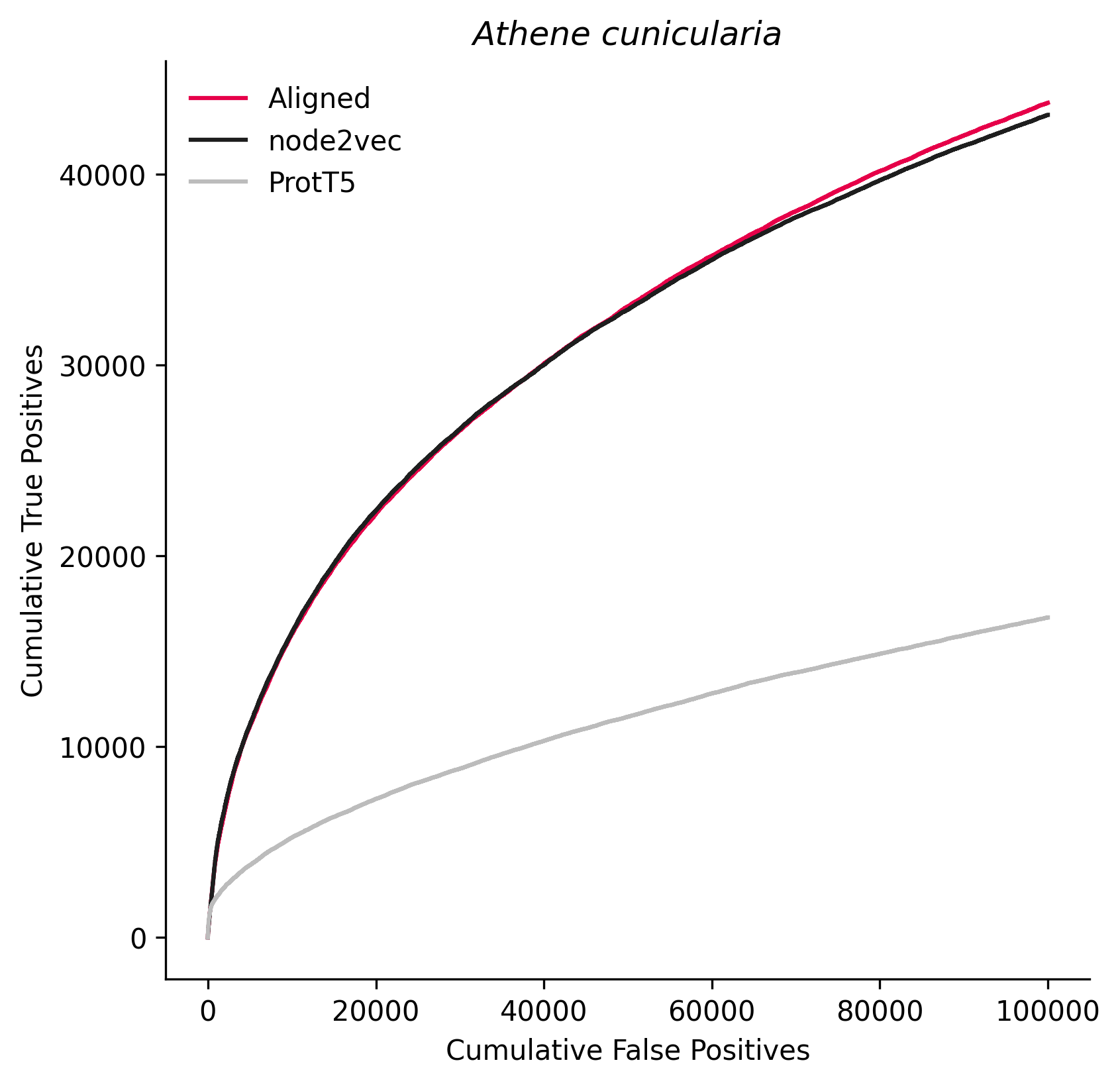

### 227321.png

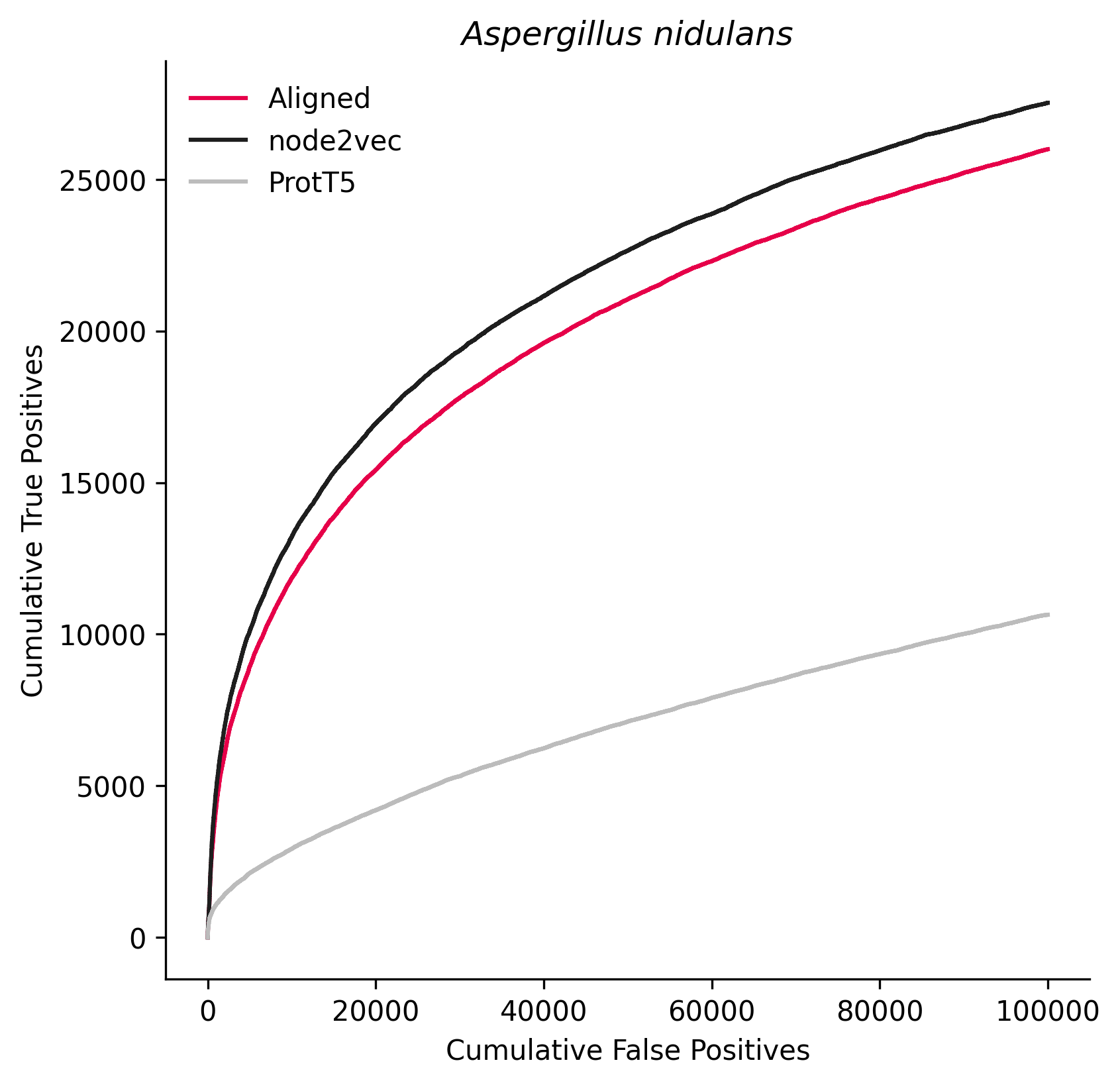

### 237561.png

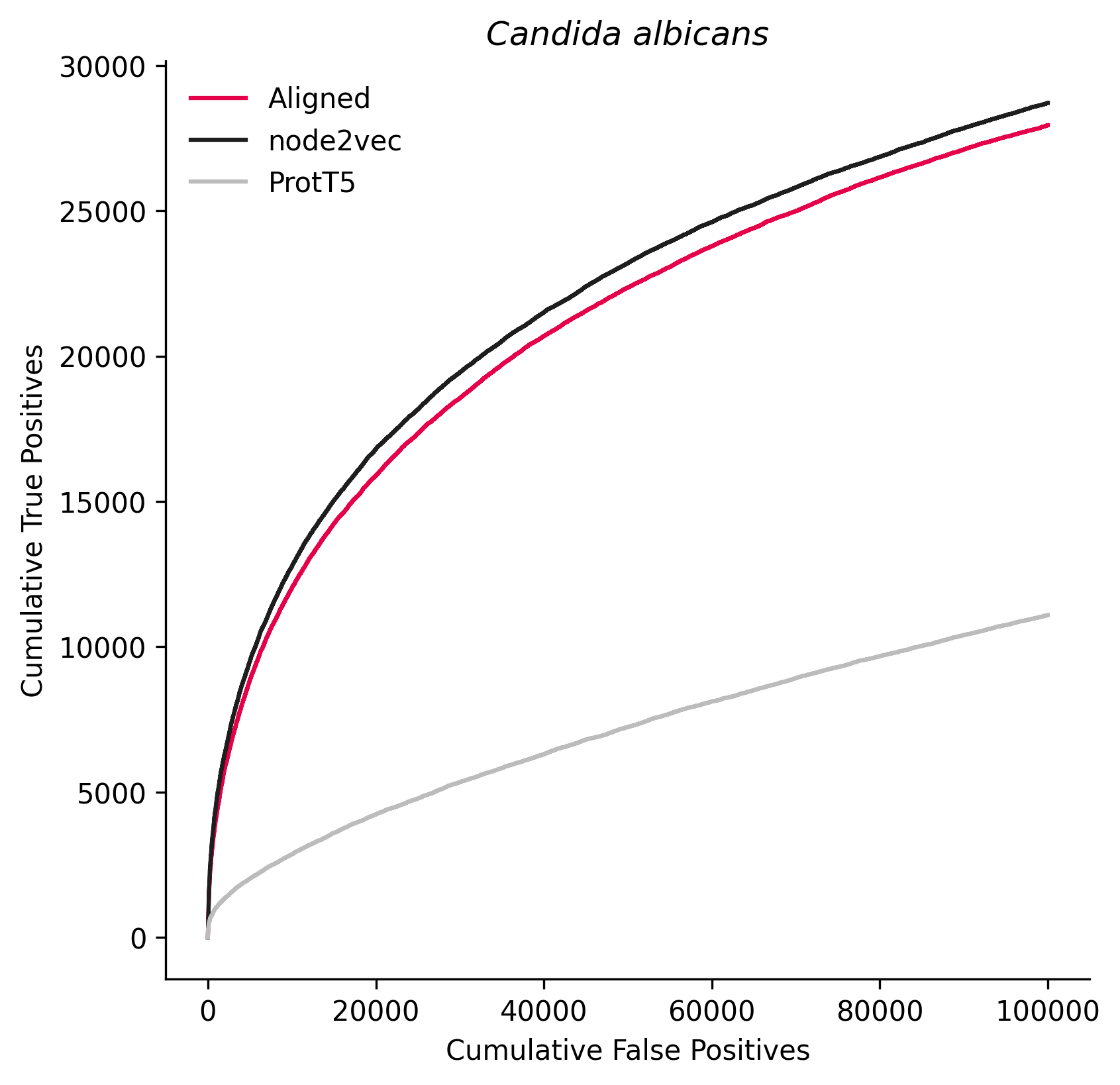

### 244447.png

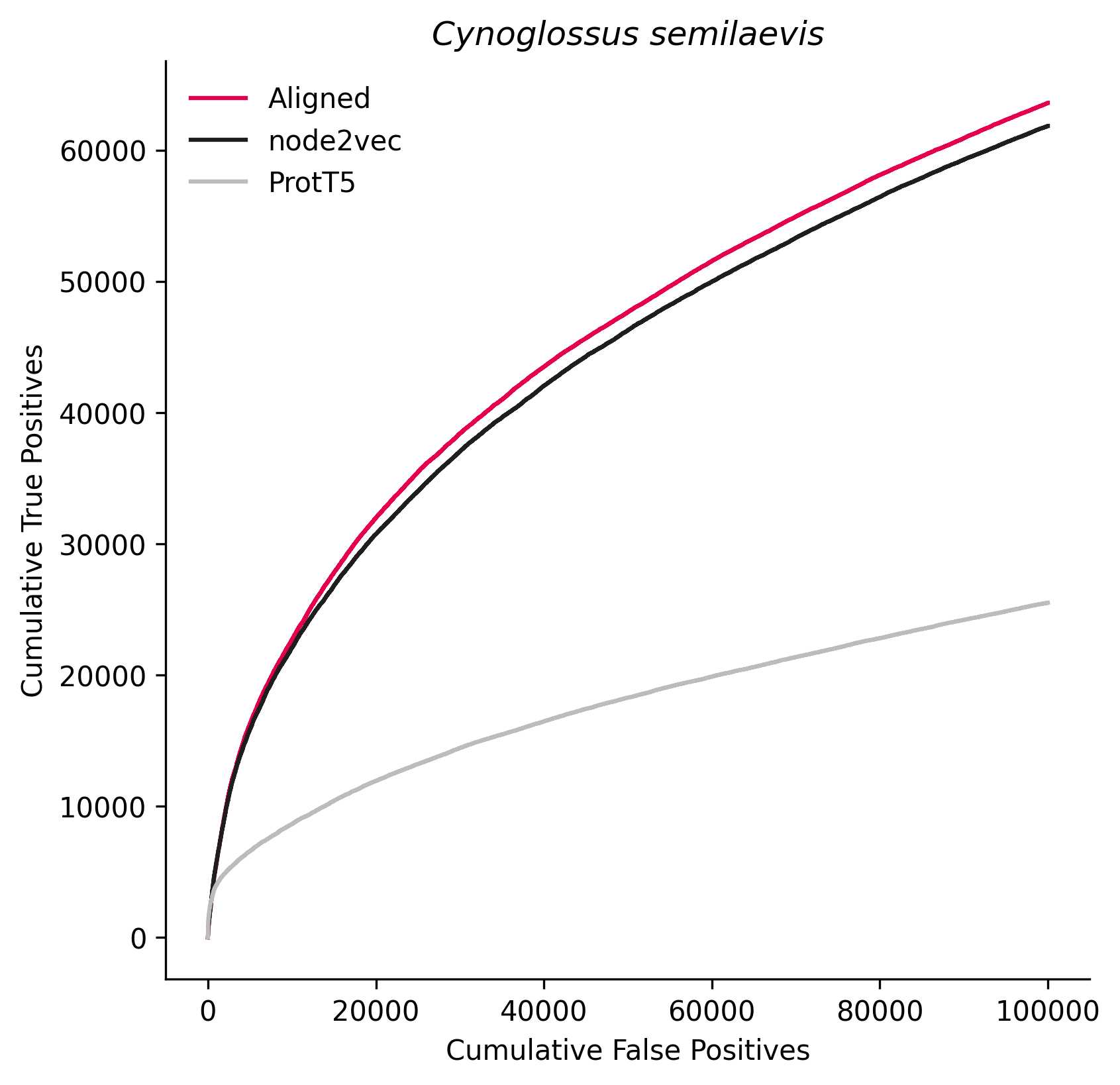

### 246437.png

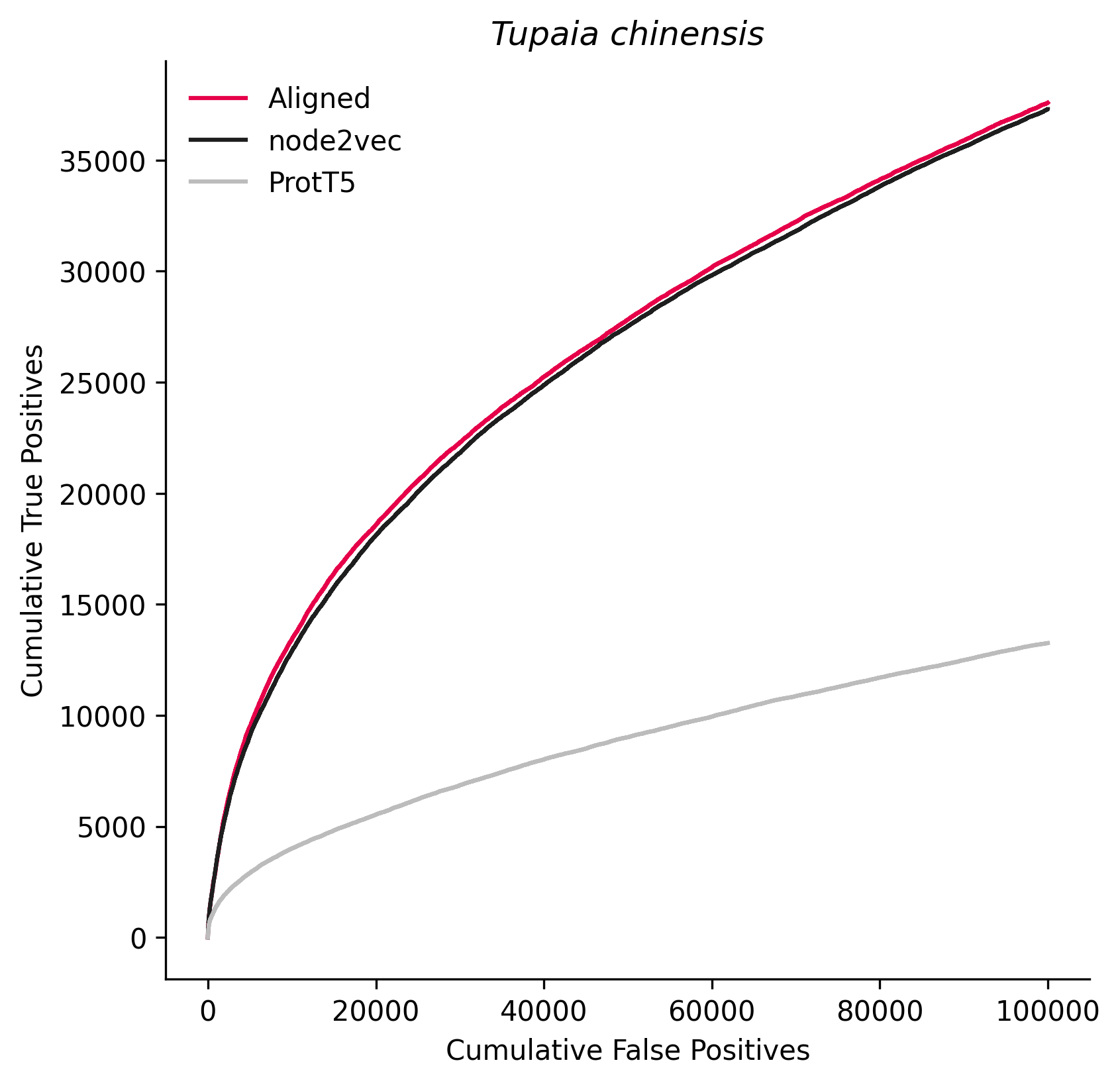

### 280699.png

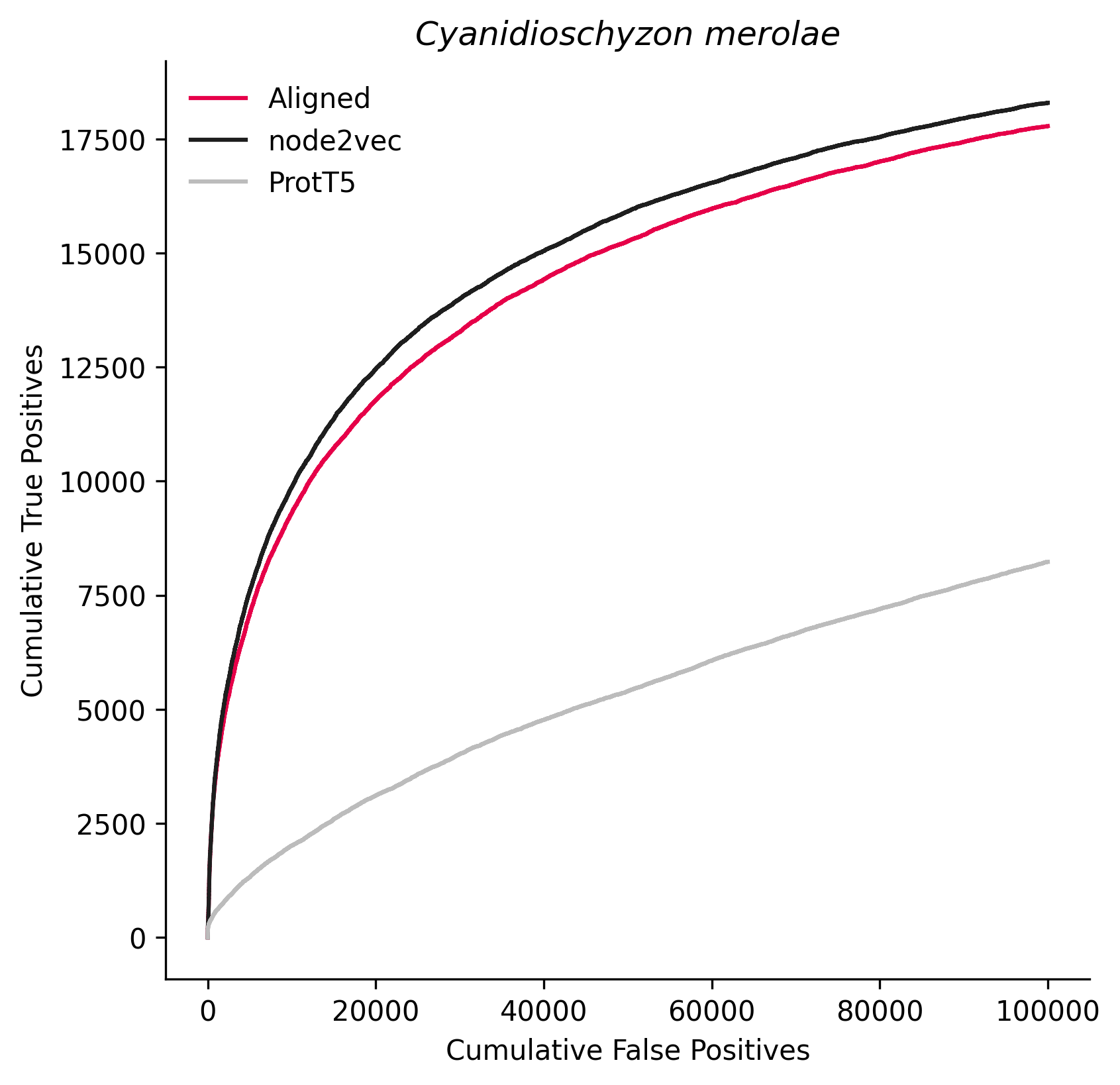

### 299123.png

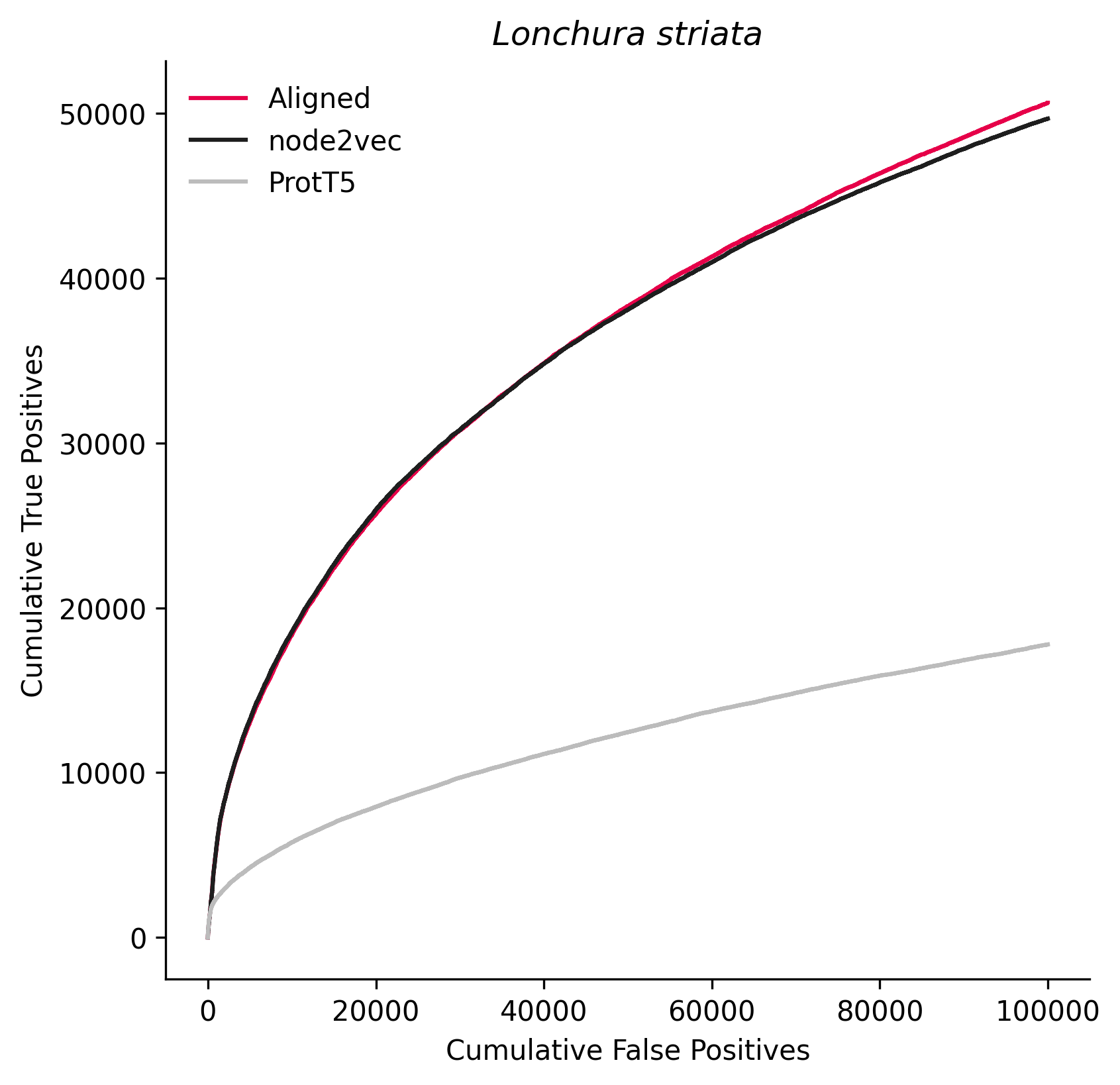

### 312017.png

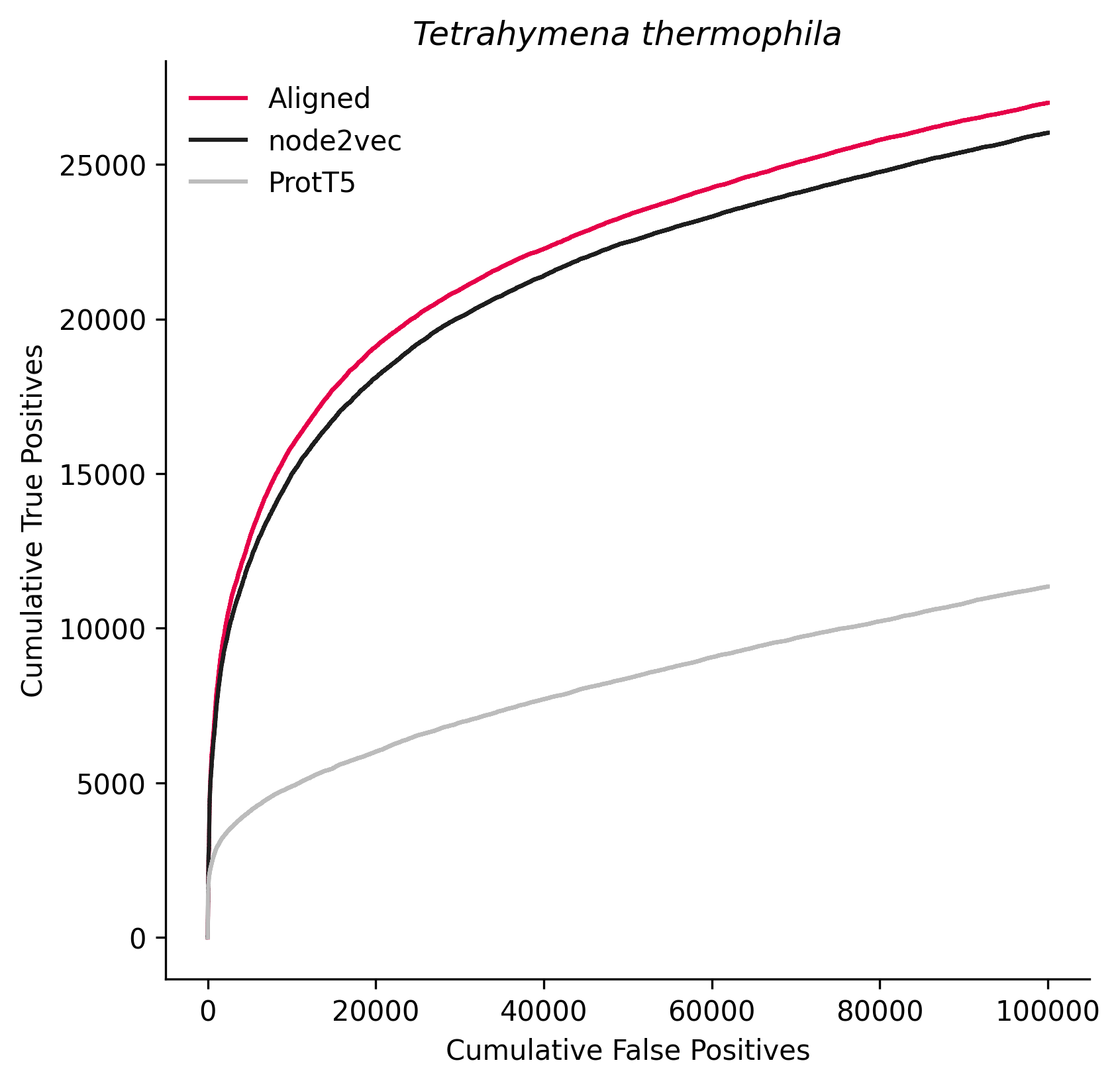

### 321614.png

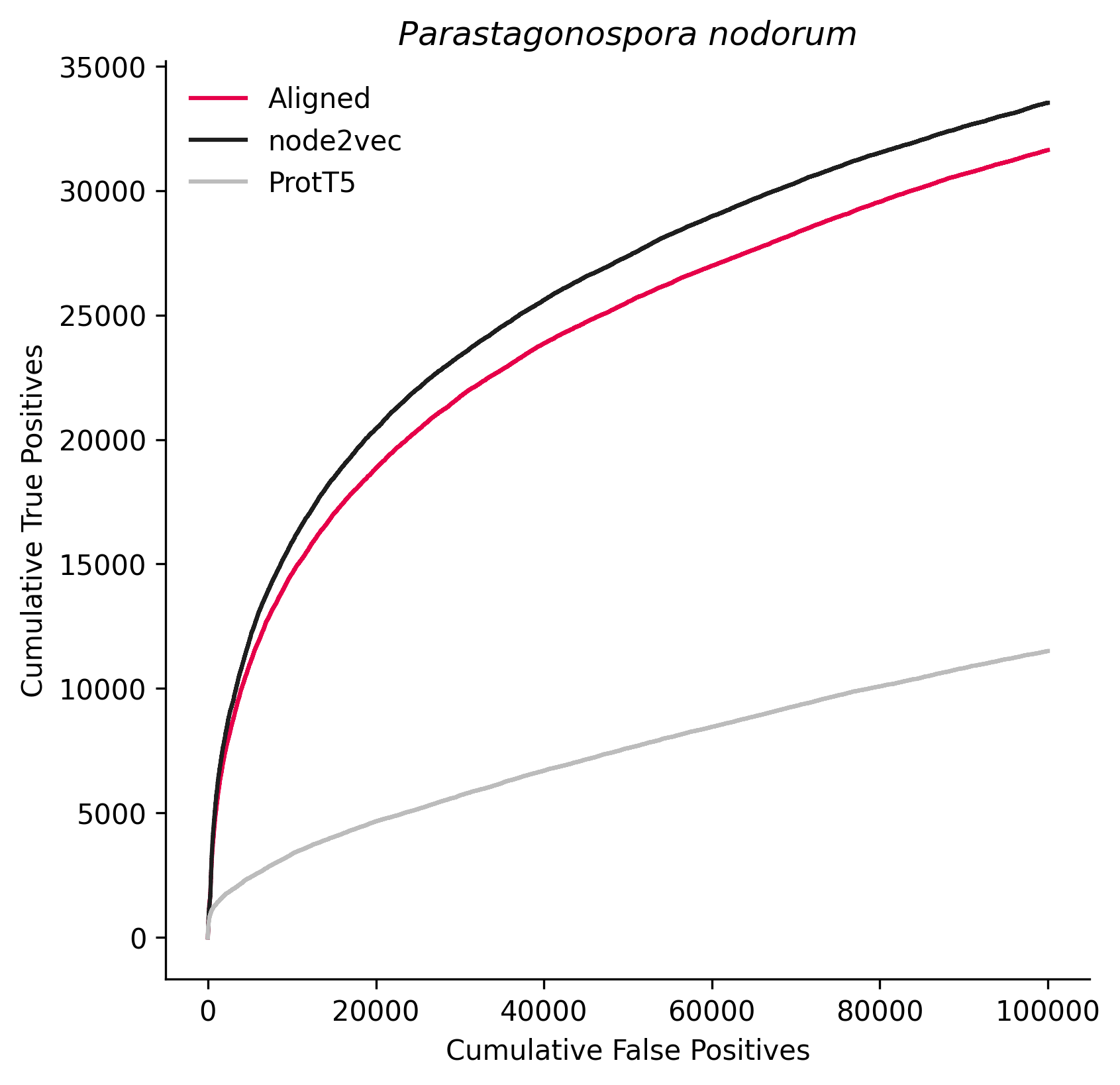

### 331117.png

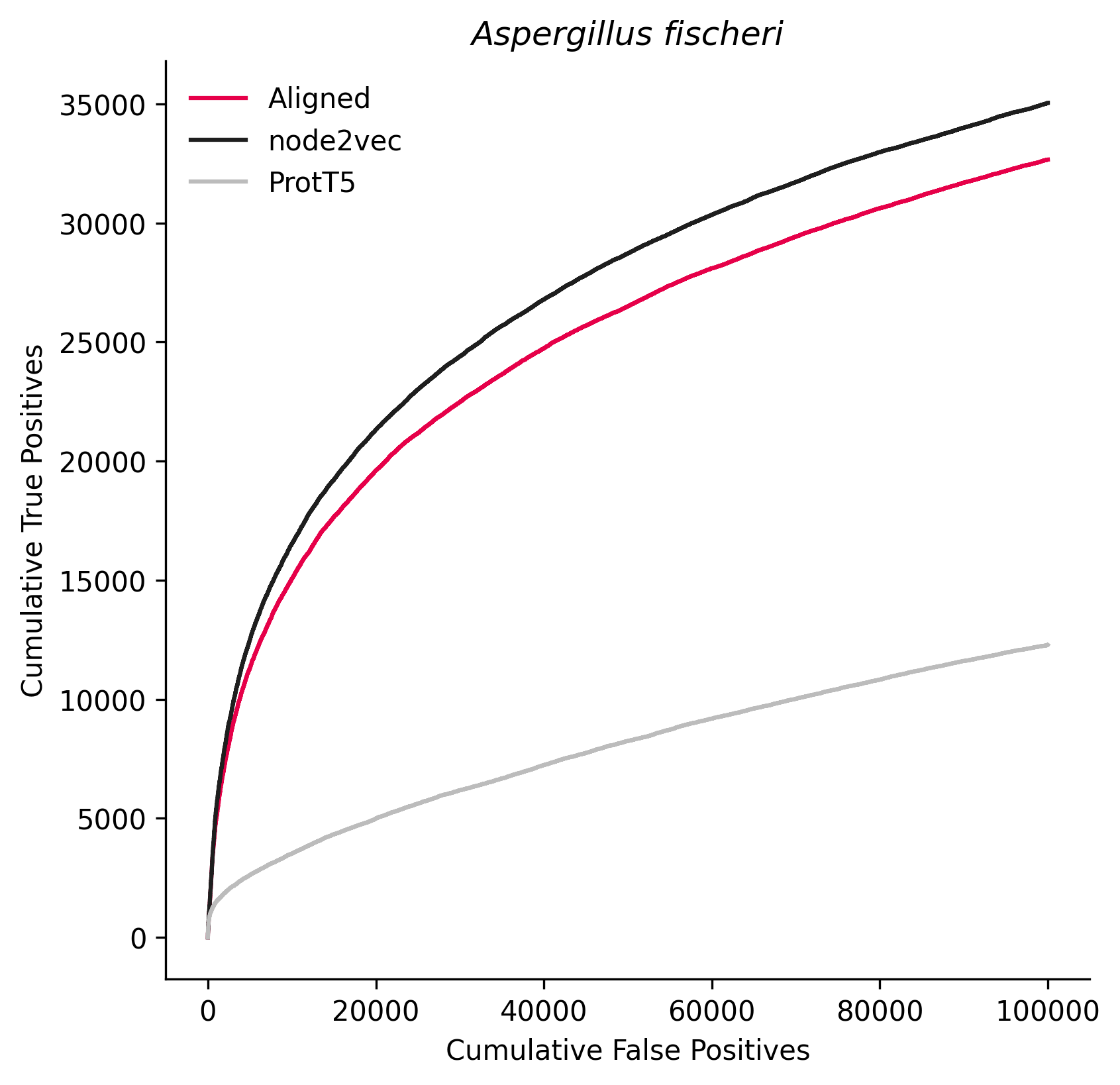

### 336722.png

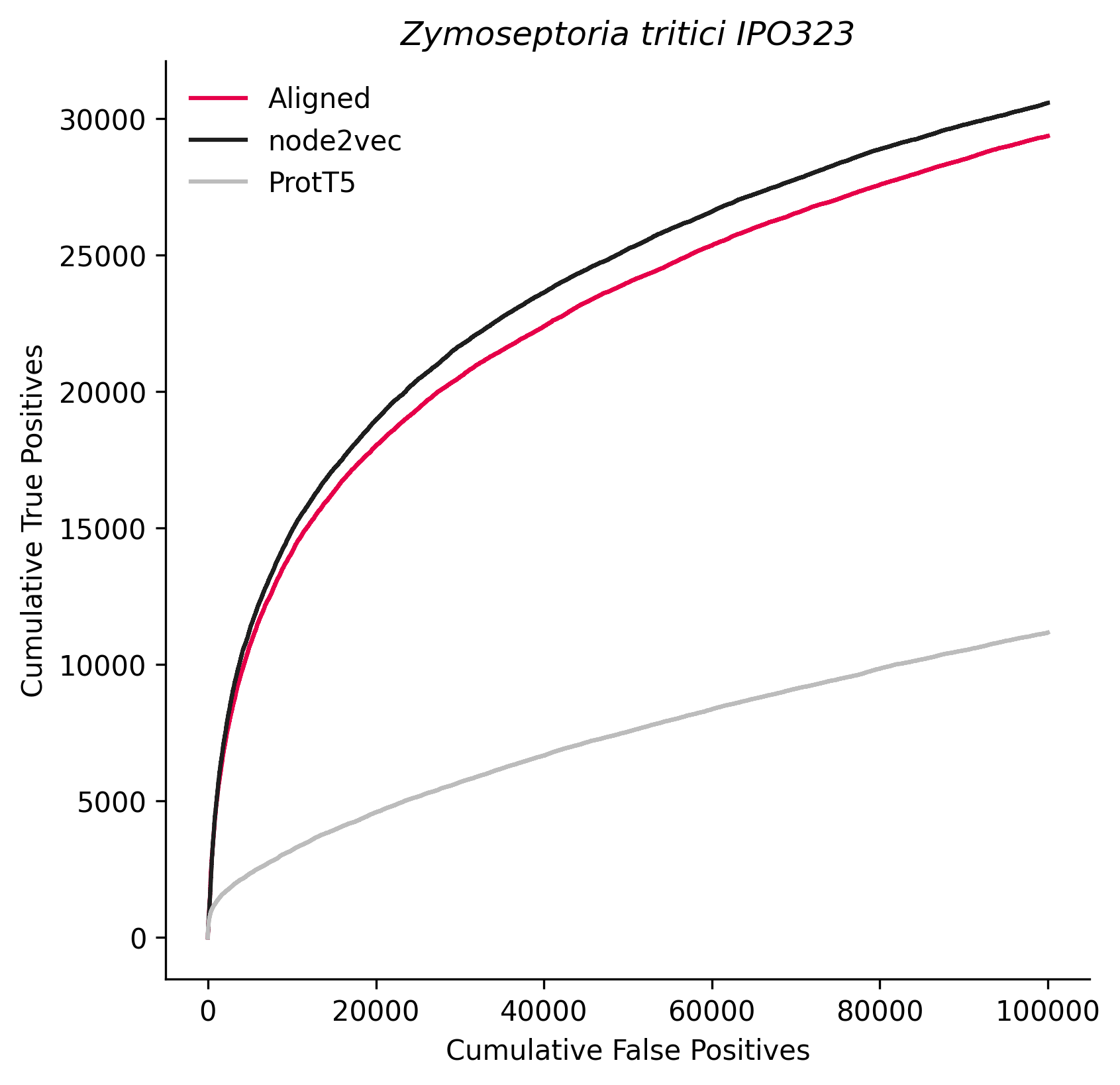

### 353152.png

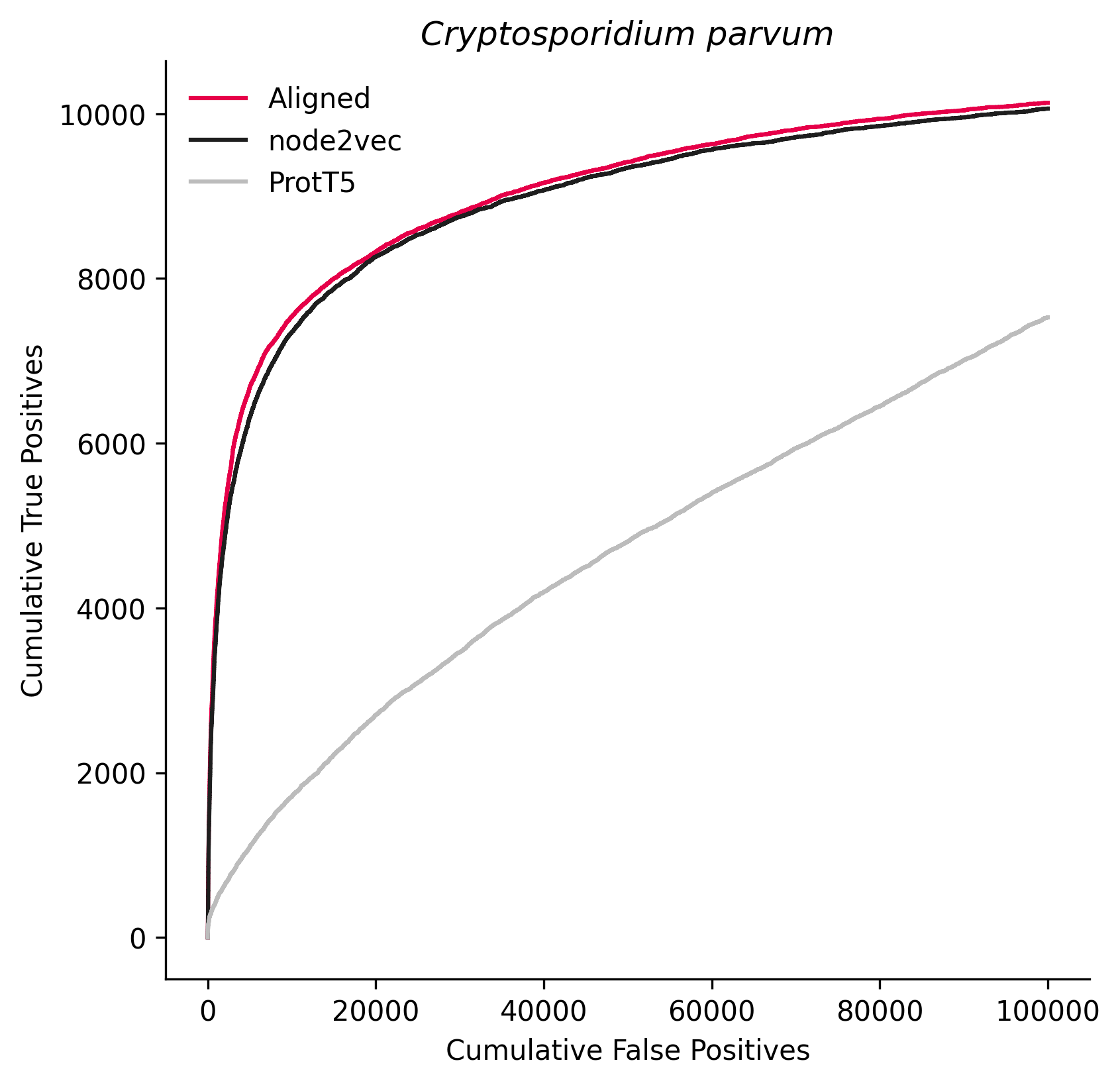

### 403677.png

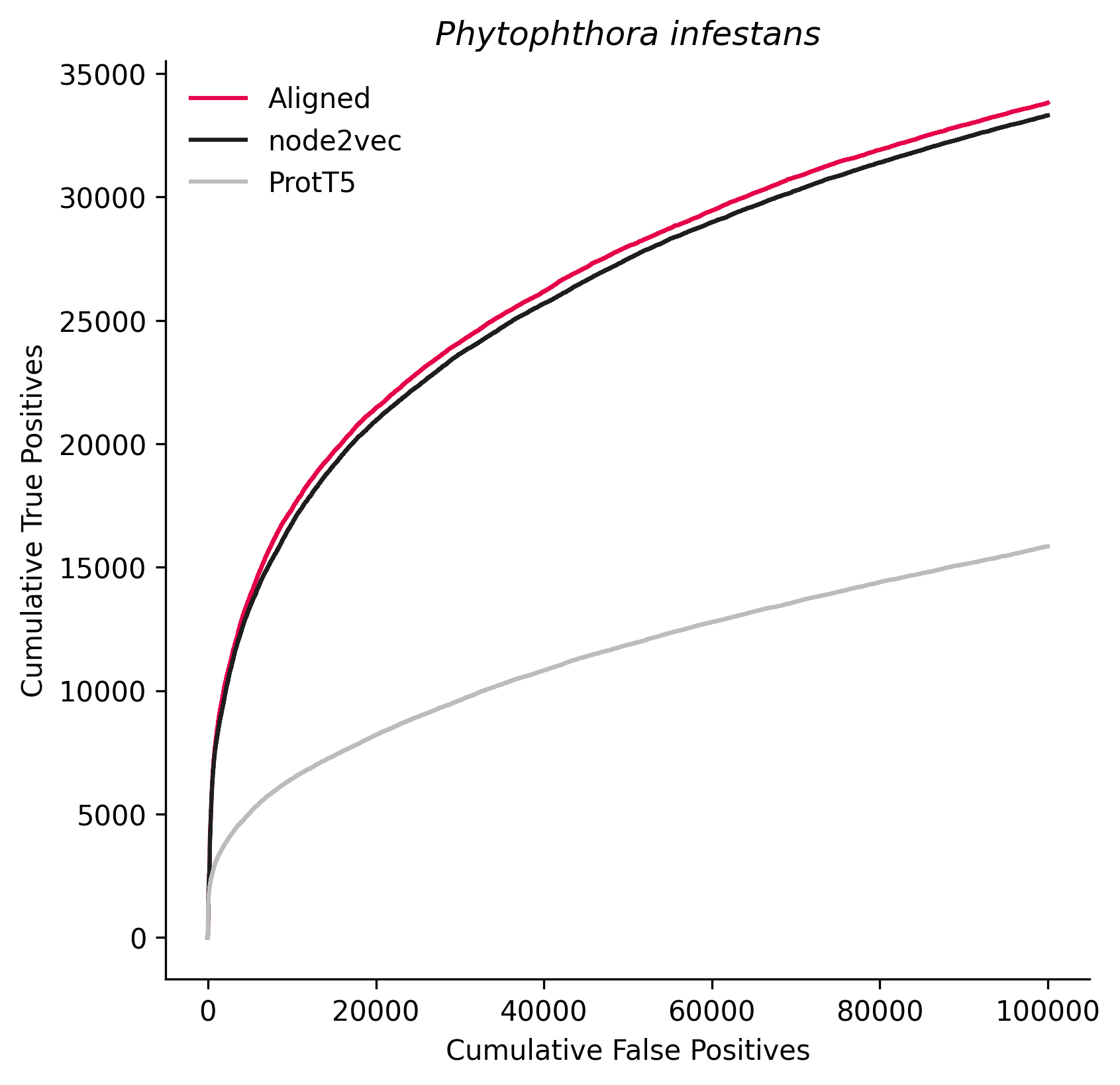

### 498019.png

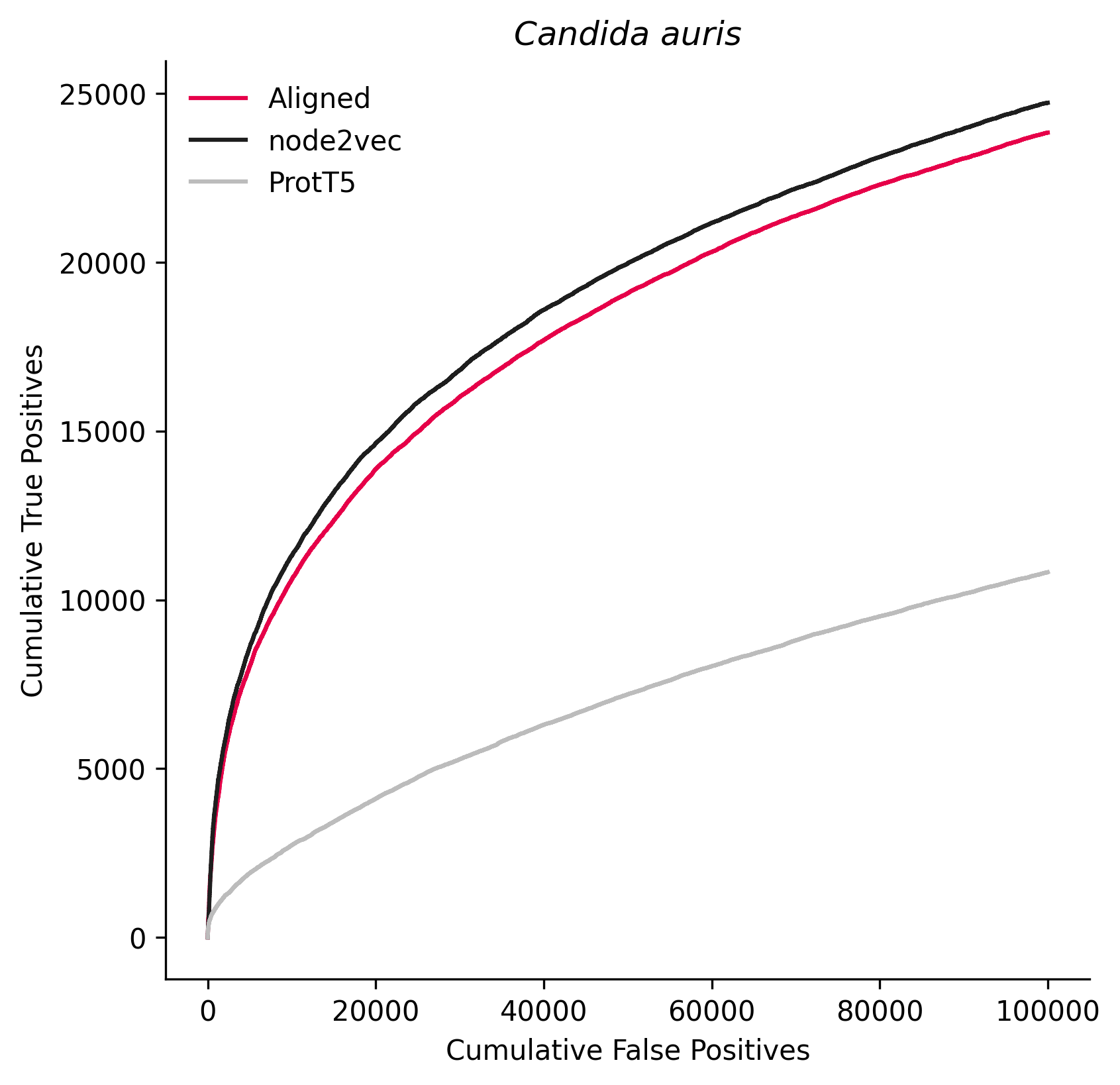

### 510516.png

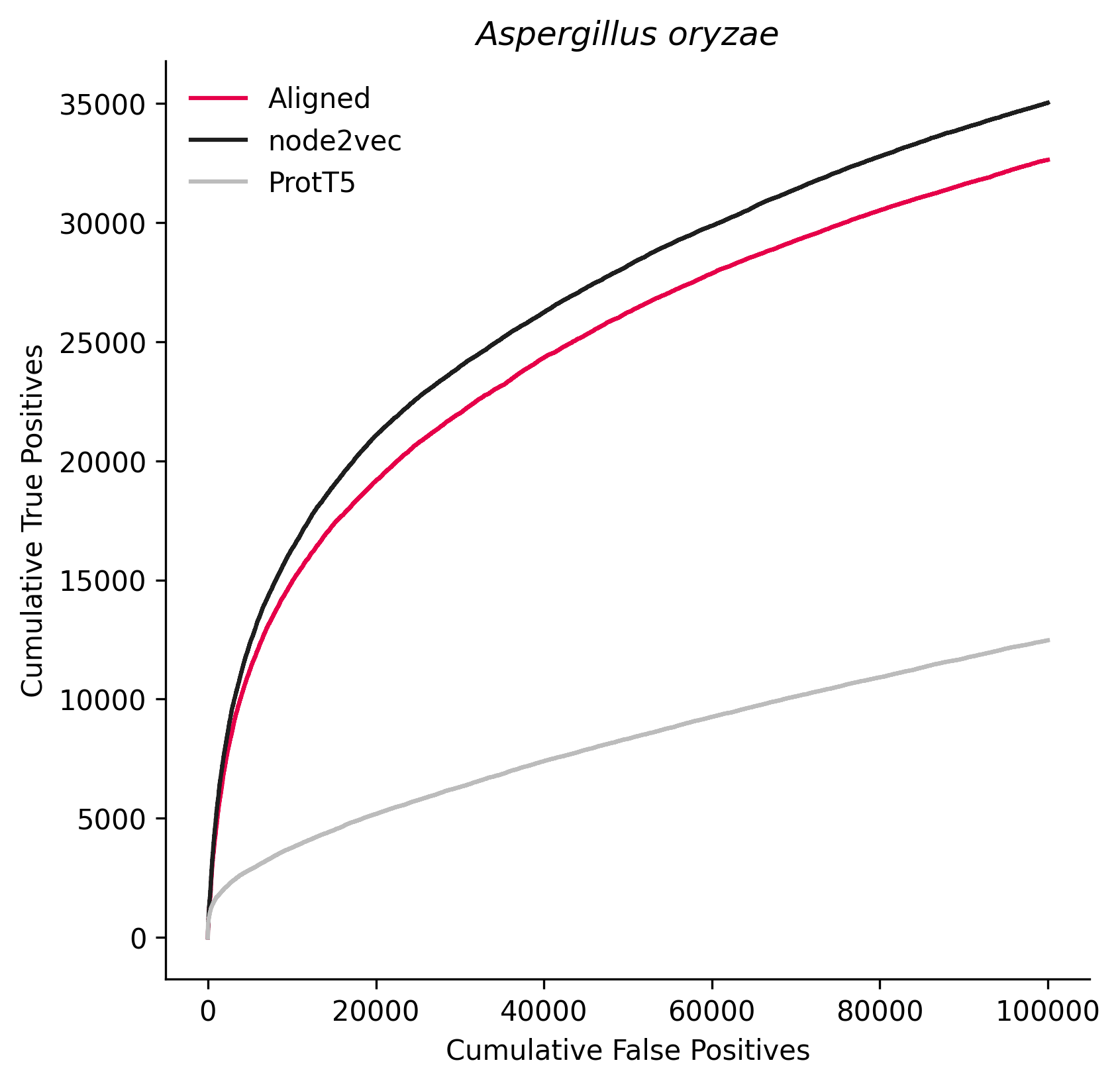

### 578455.png

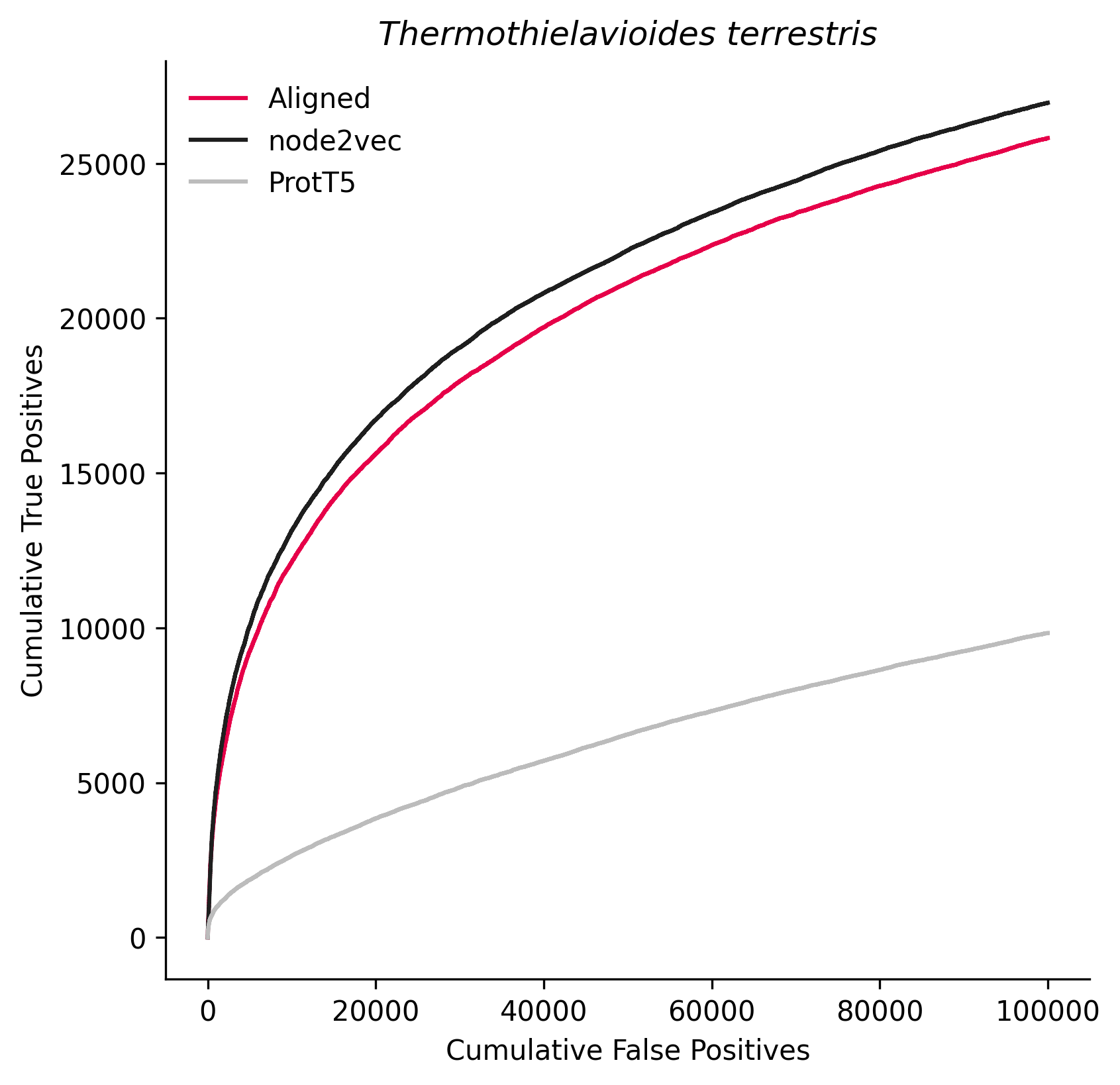

### 578458.png

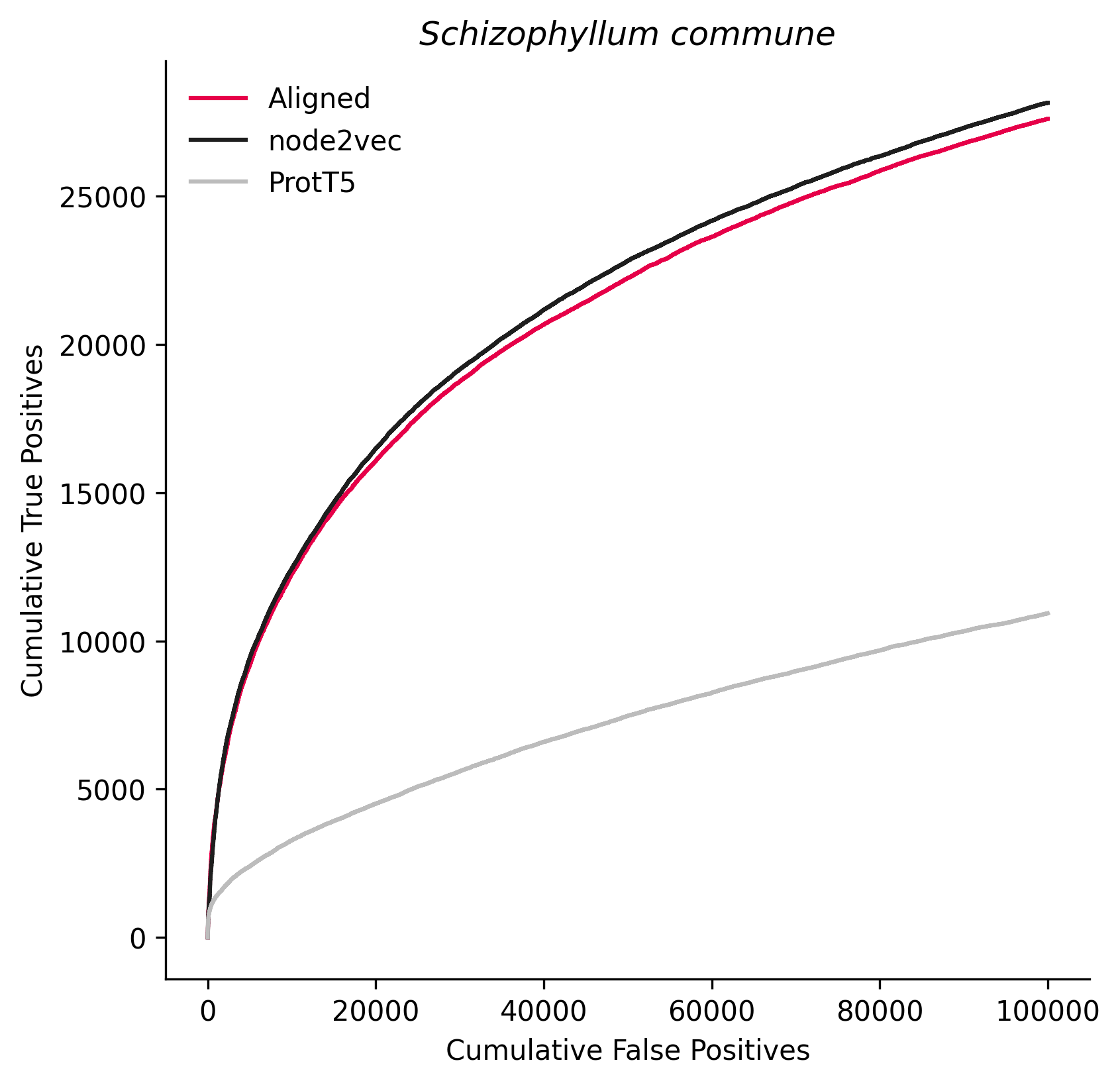

### 590646.png

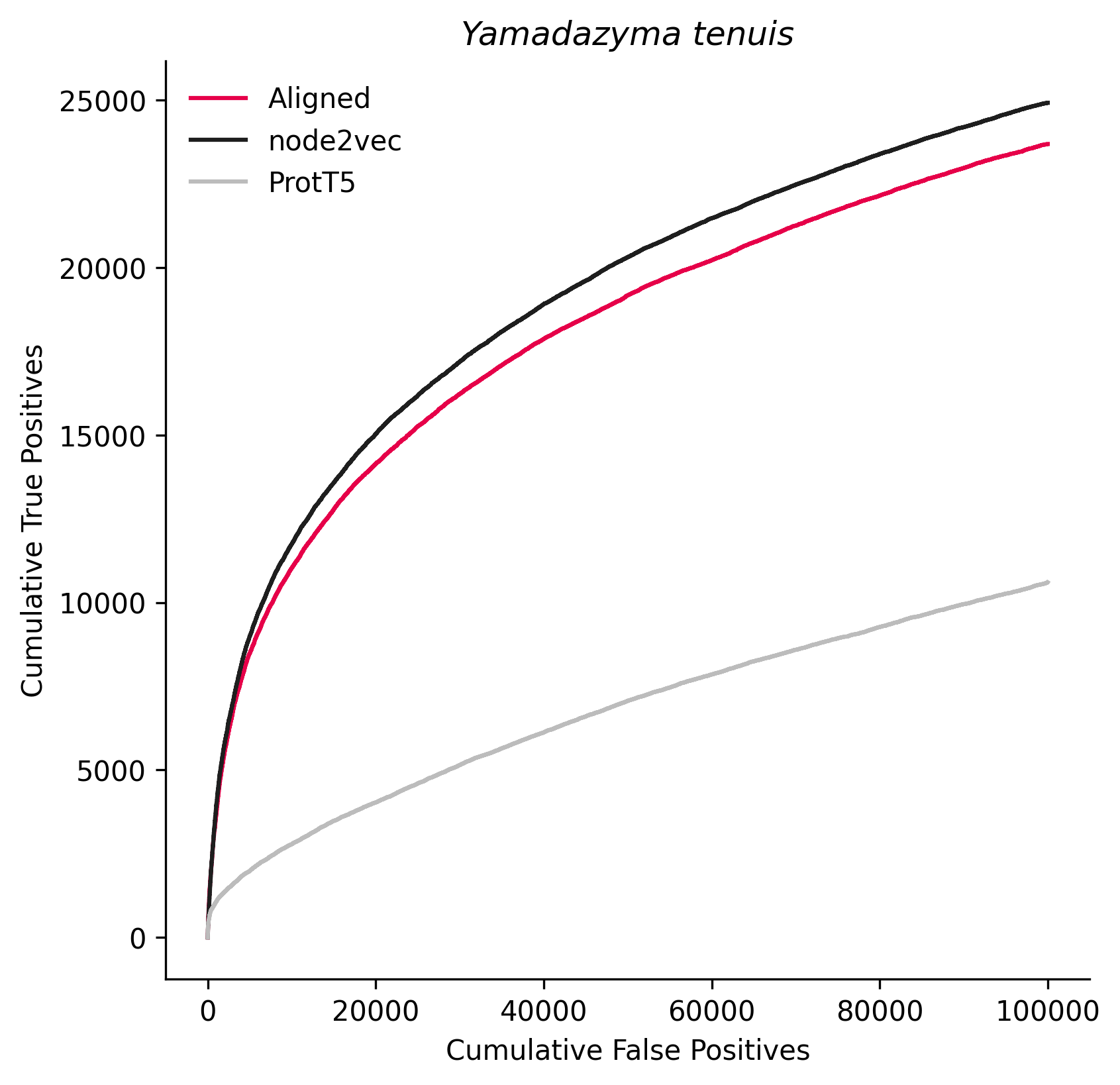

### 644223.png

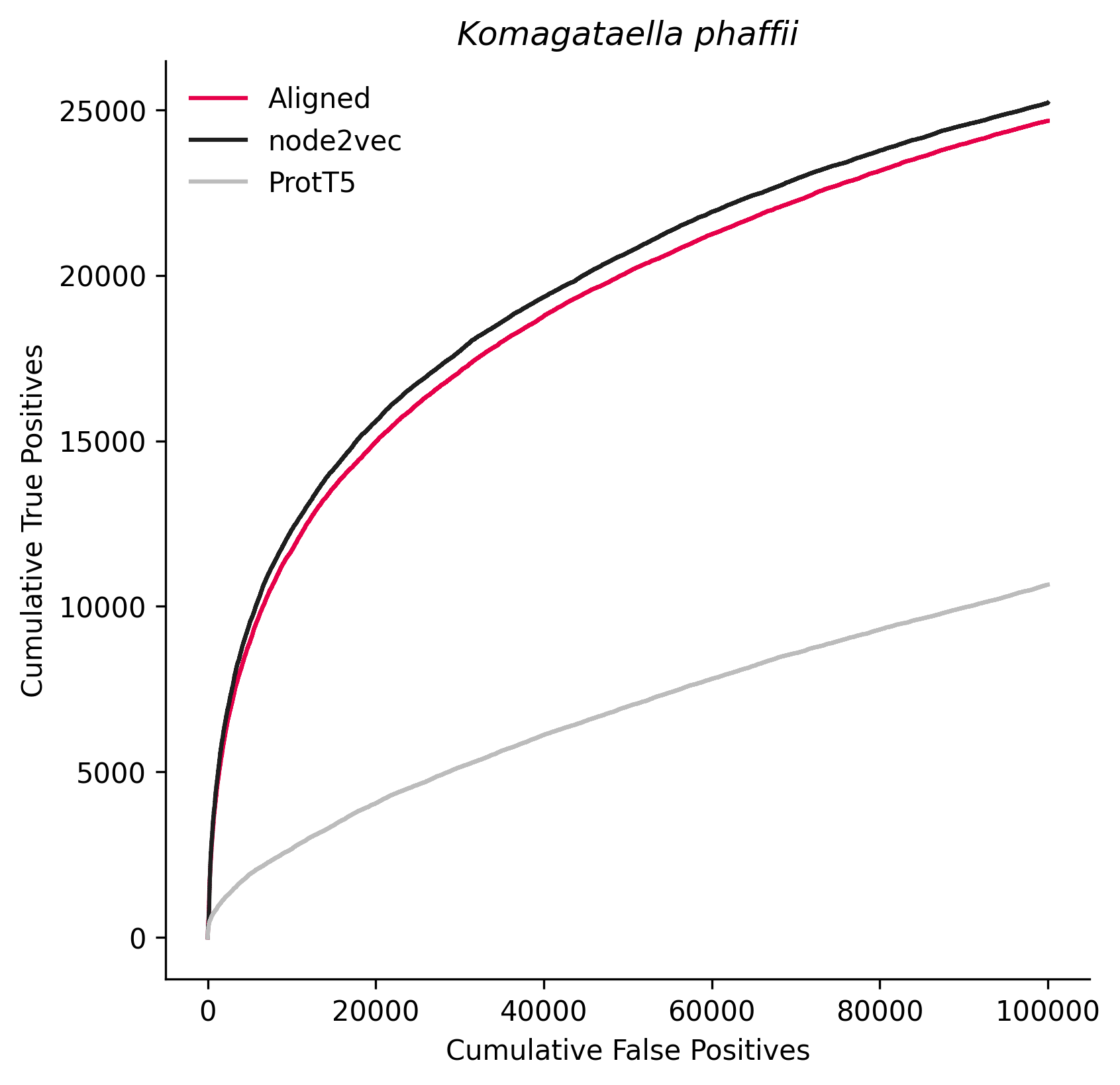

### 655827.png

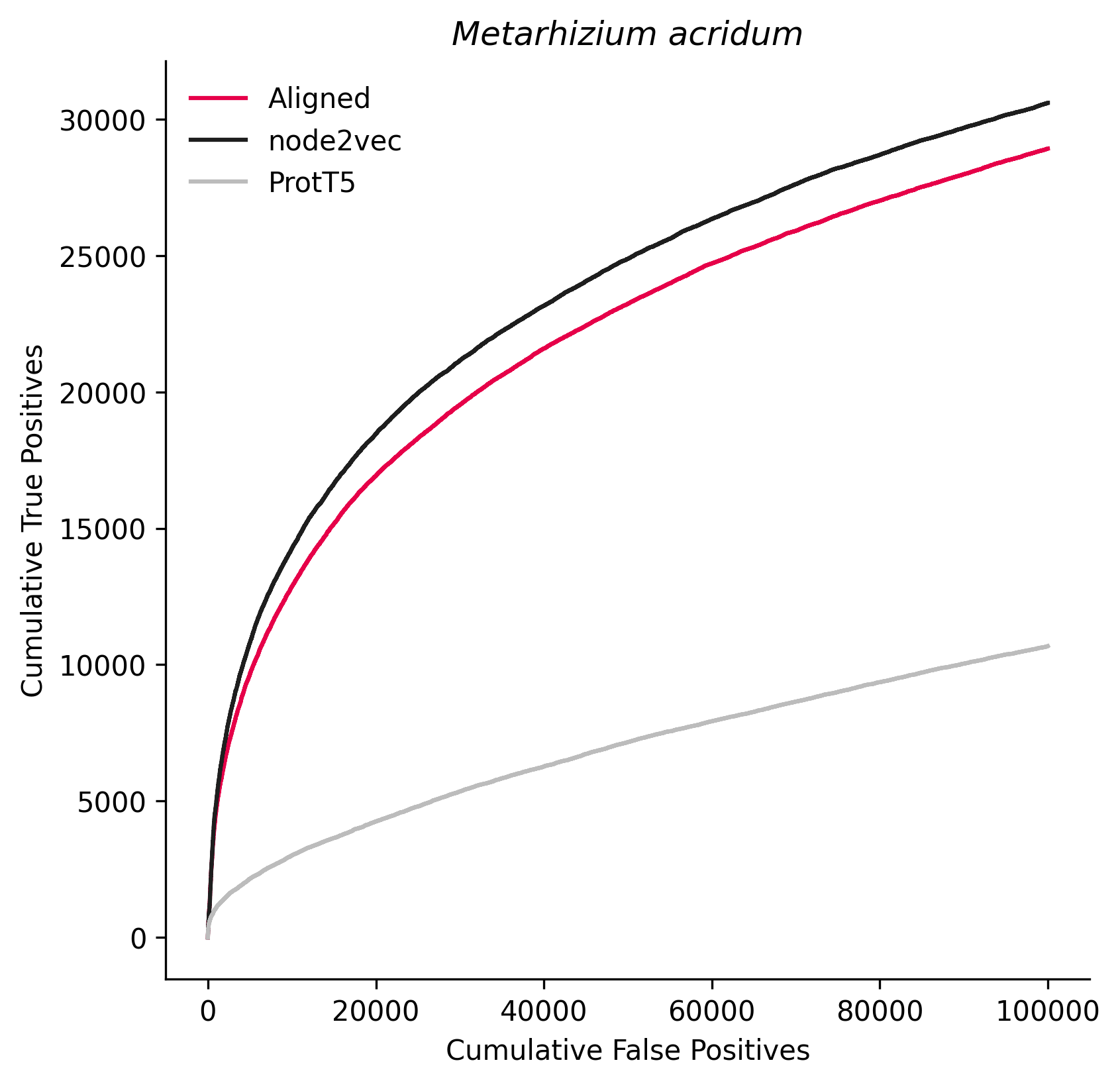

### 660122.png

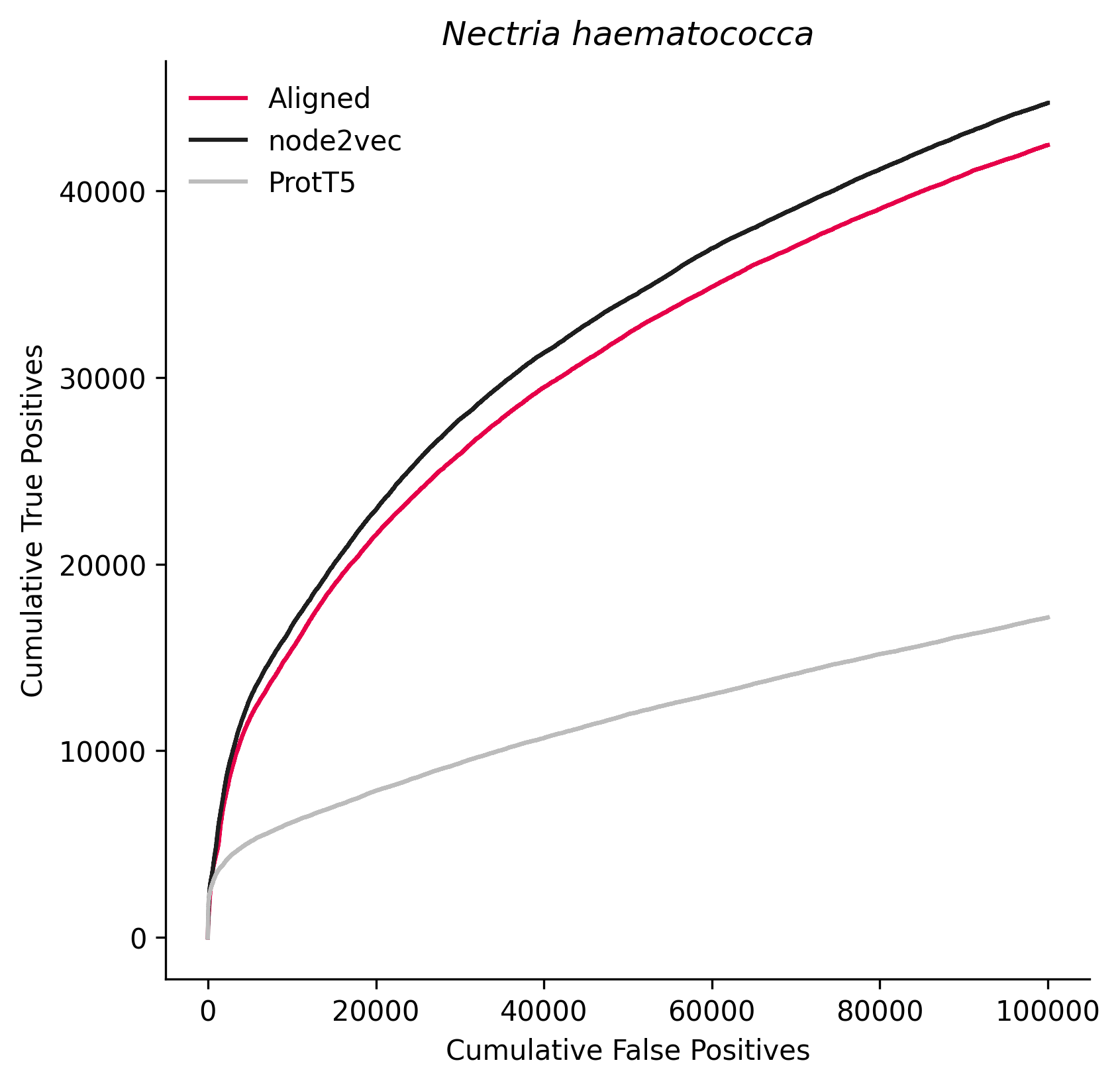

### 671144.png

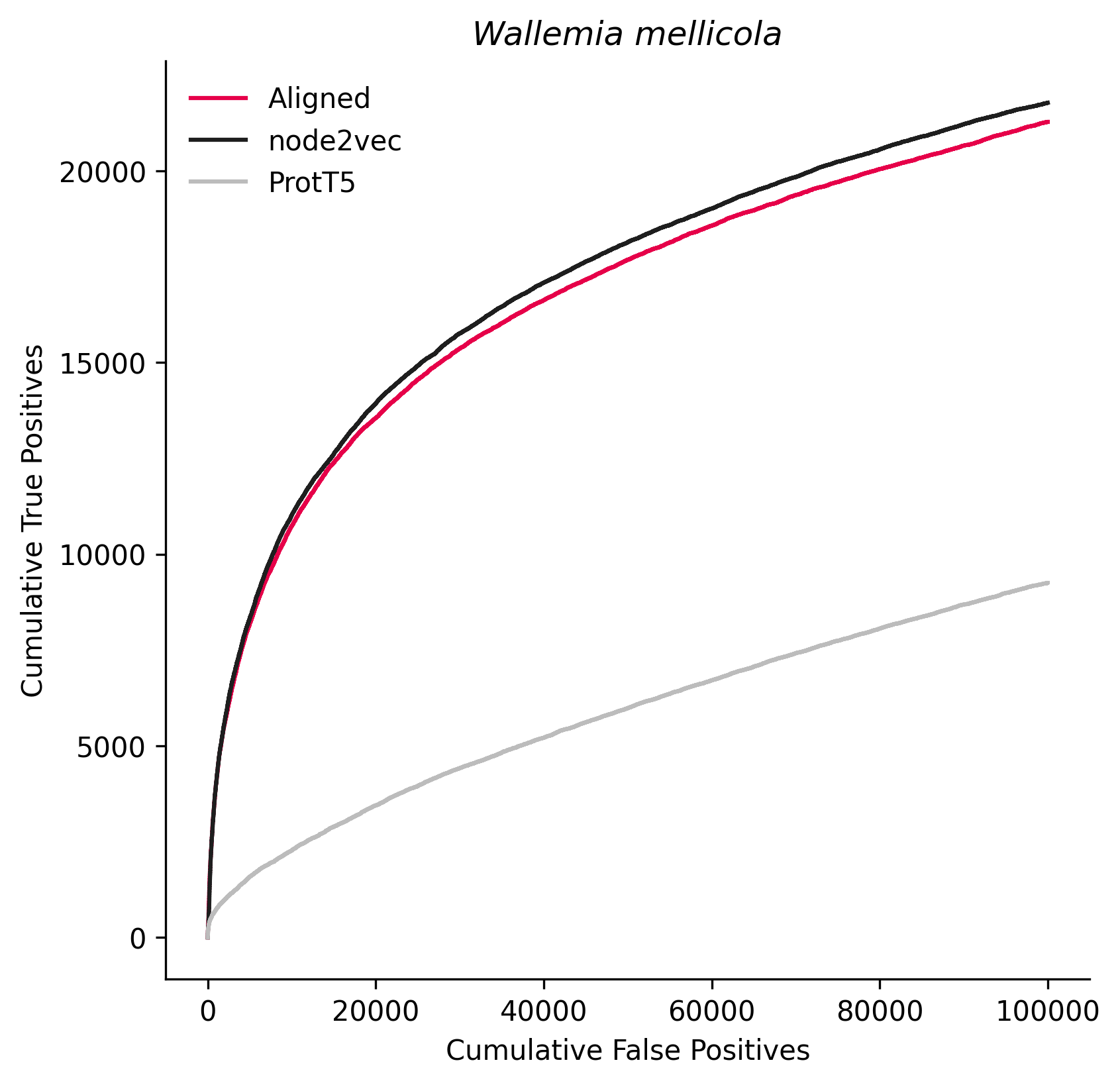

### 695850.png

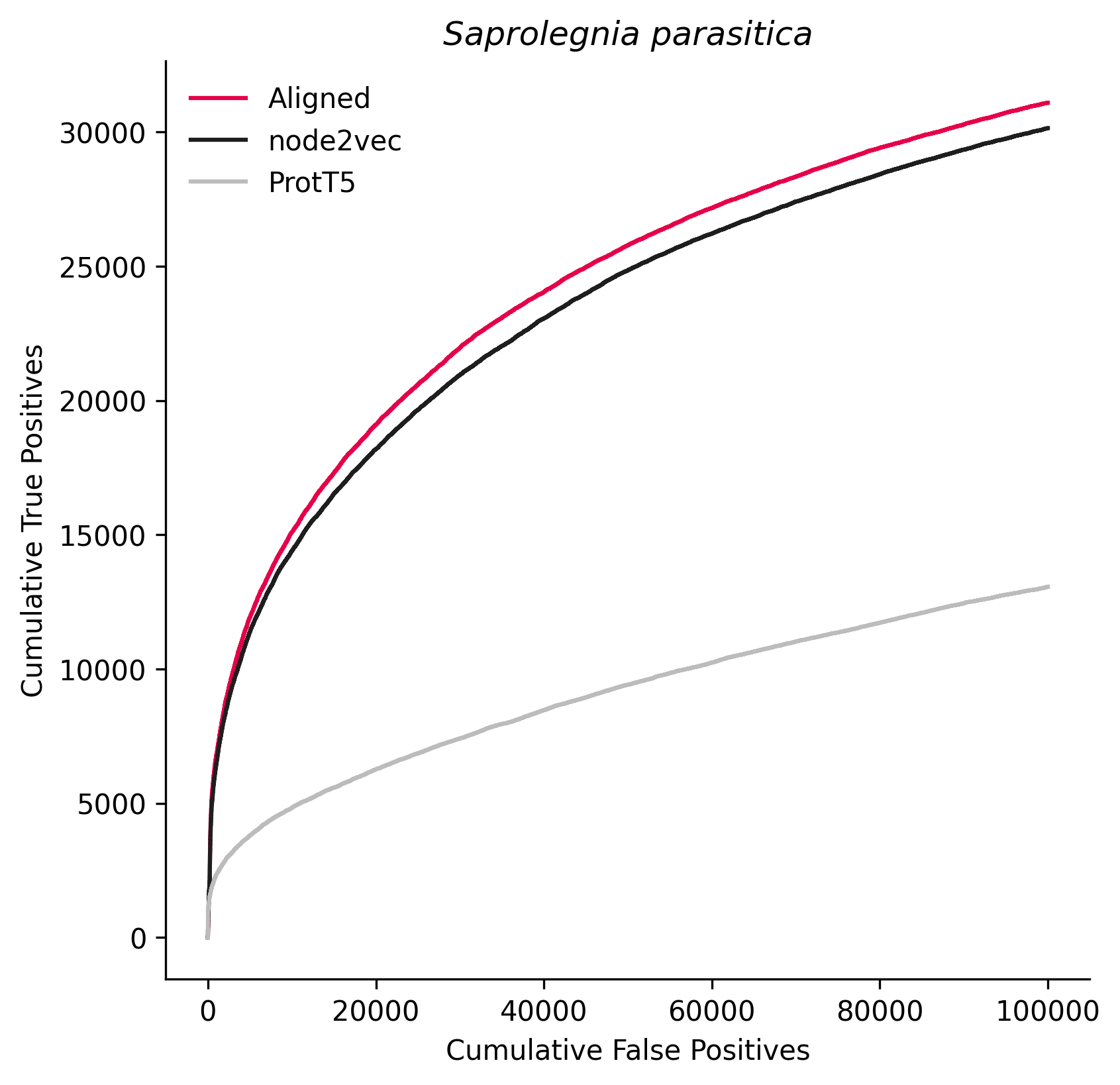

### 869250.png

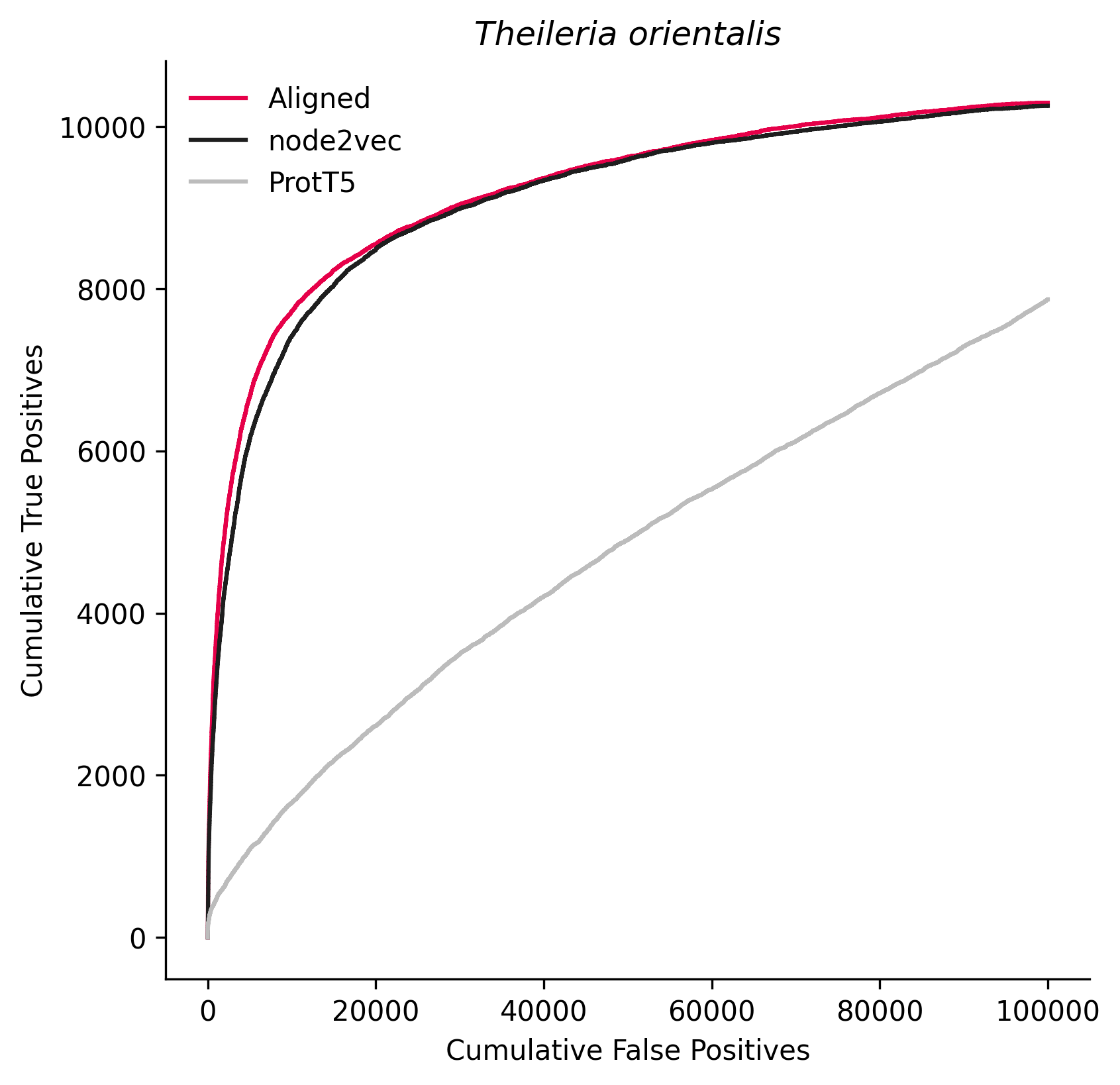

### 906689.png

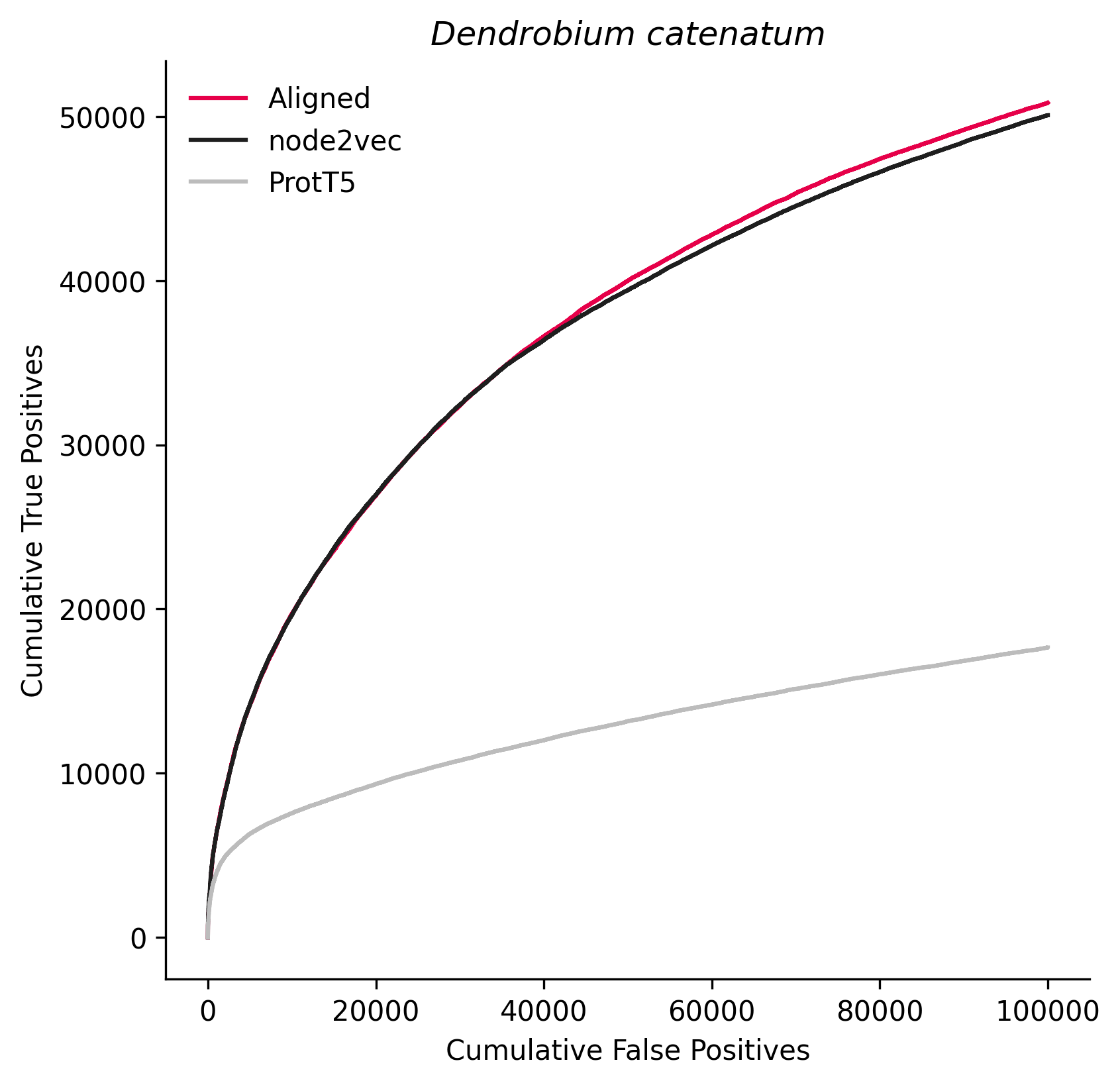

### 1026970.png

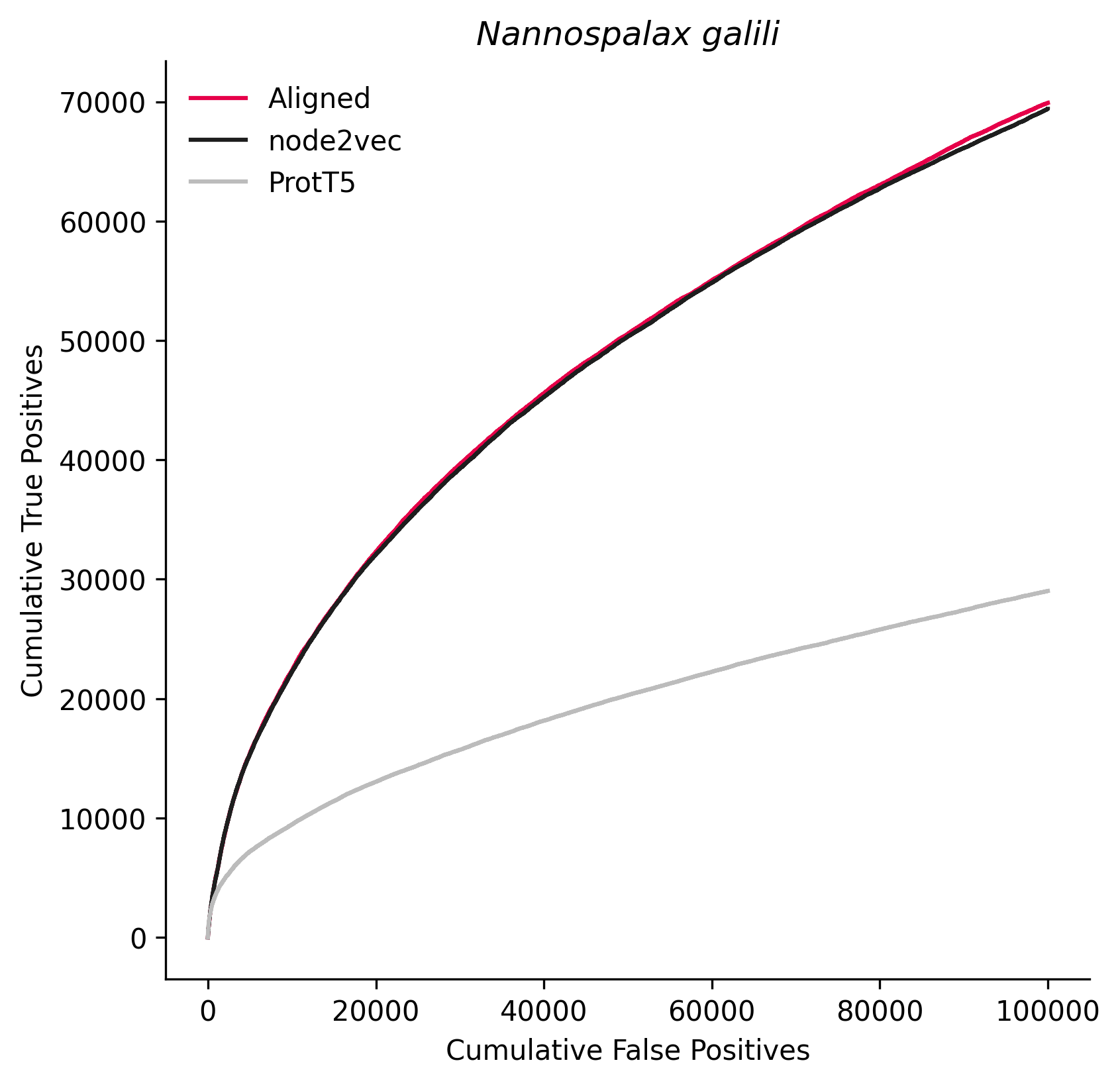

### 1054147.png

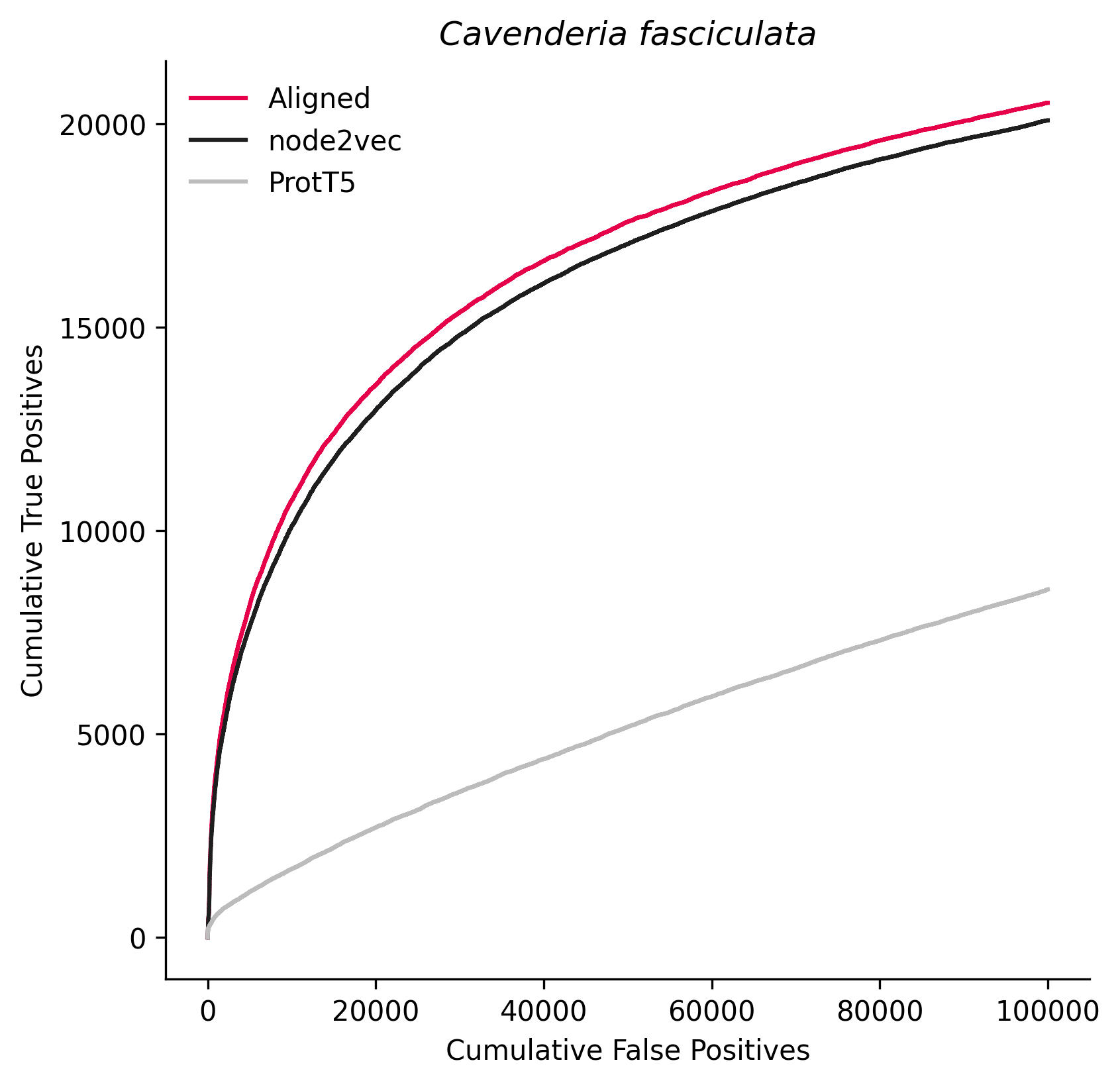

### 1071381.png

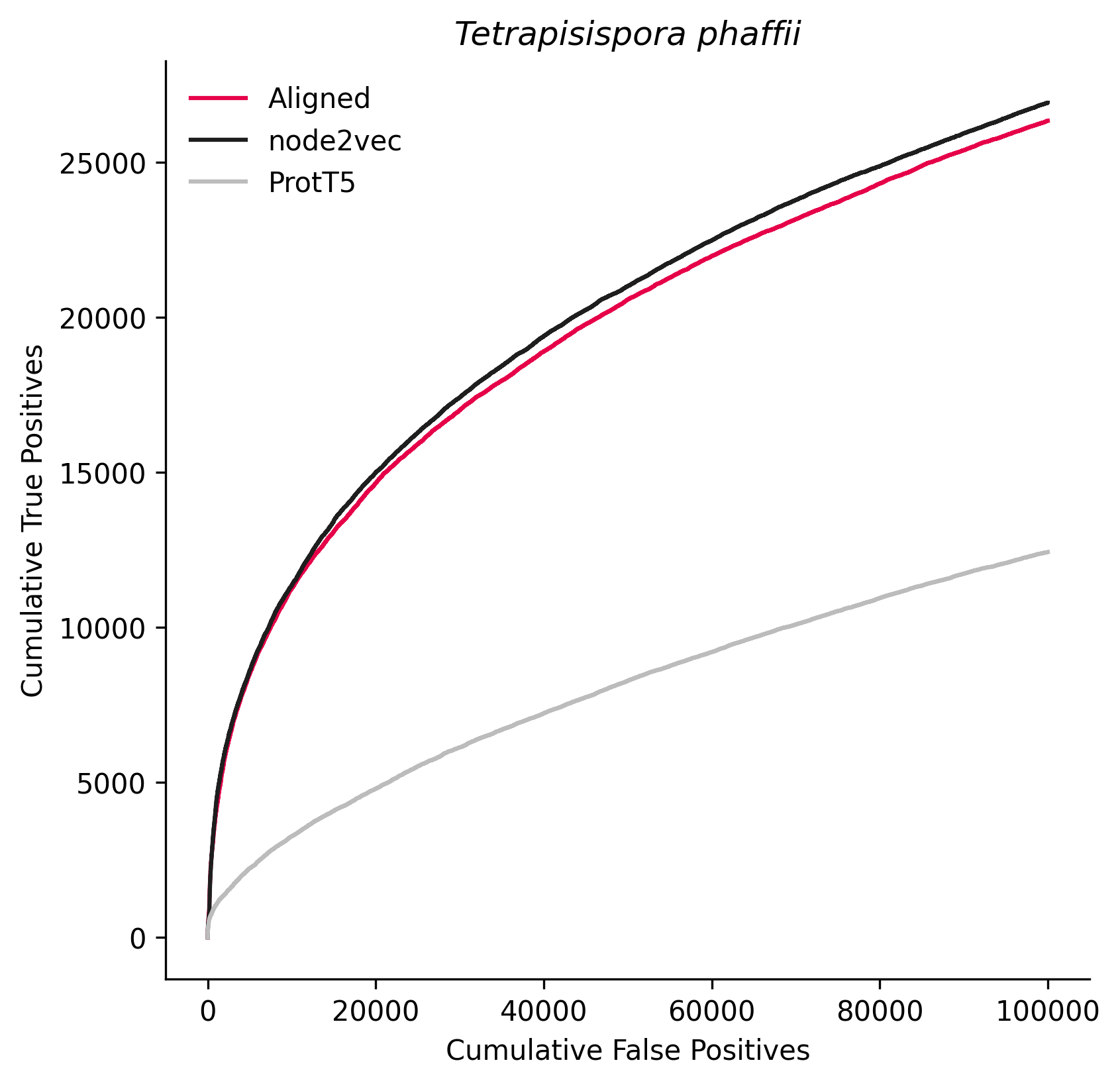

### 1094619.png

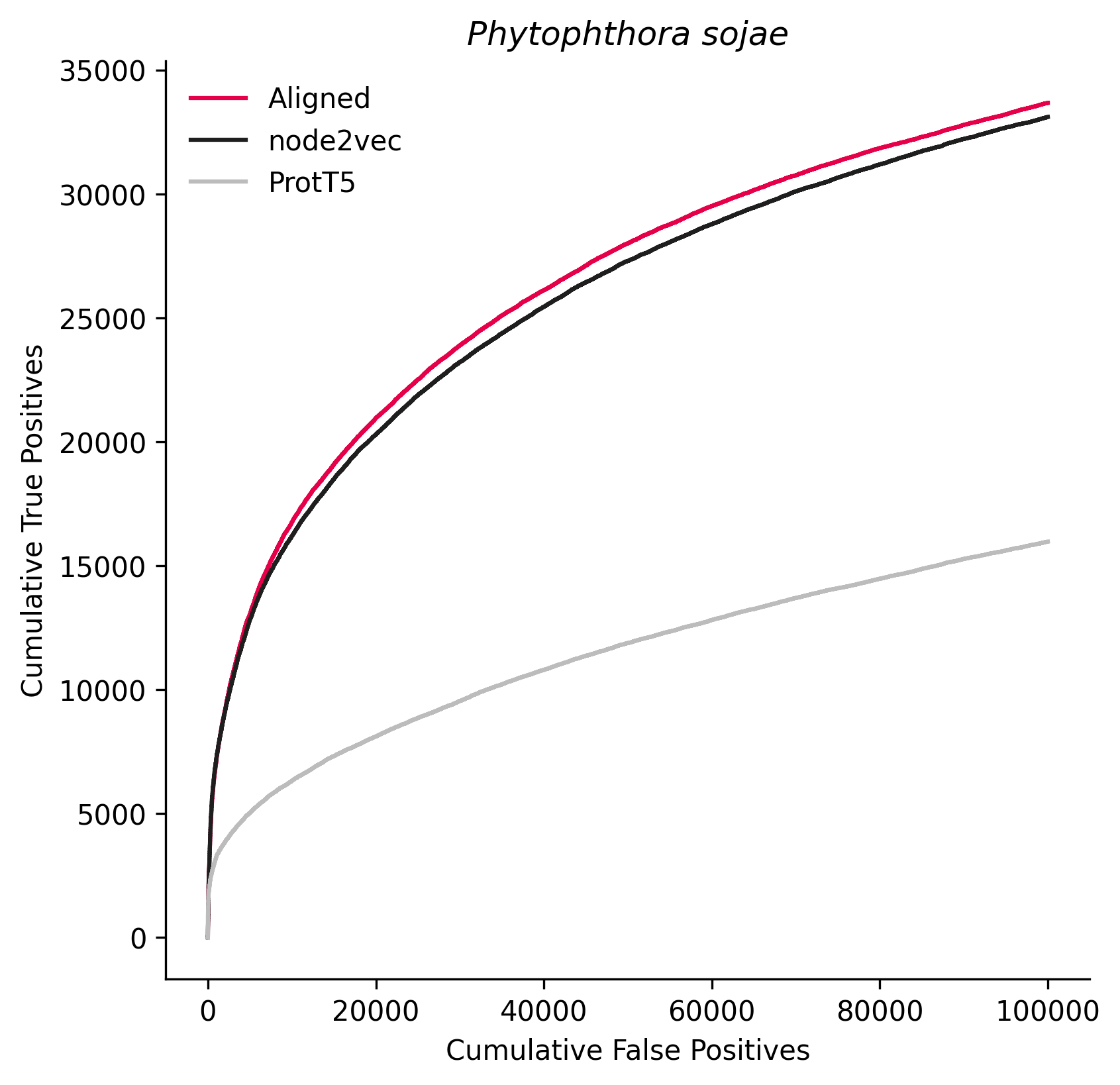

### 1116229.png

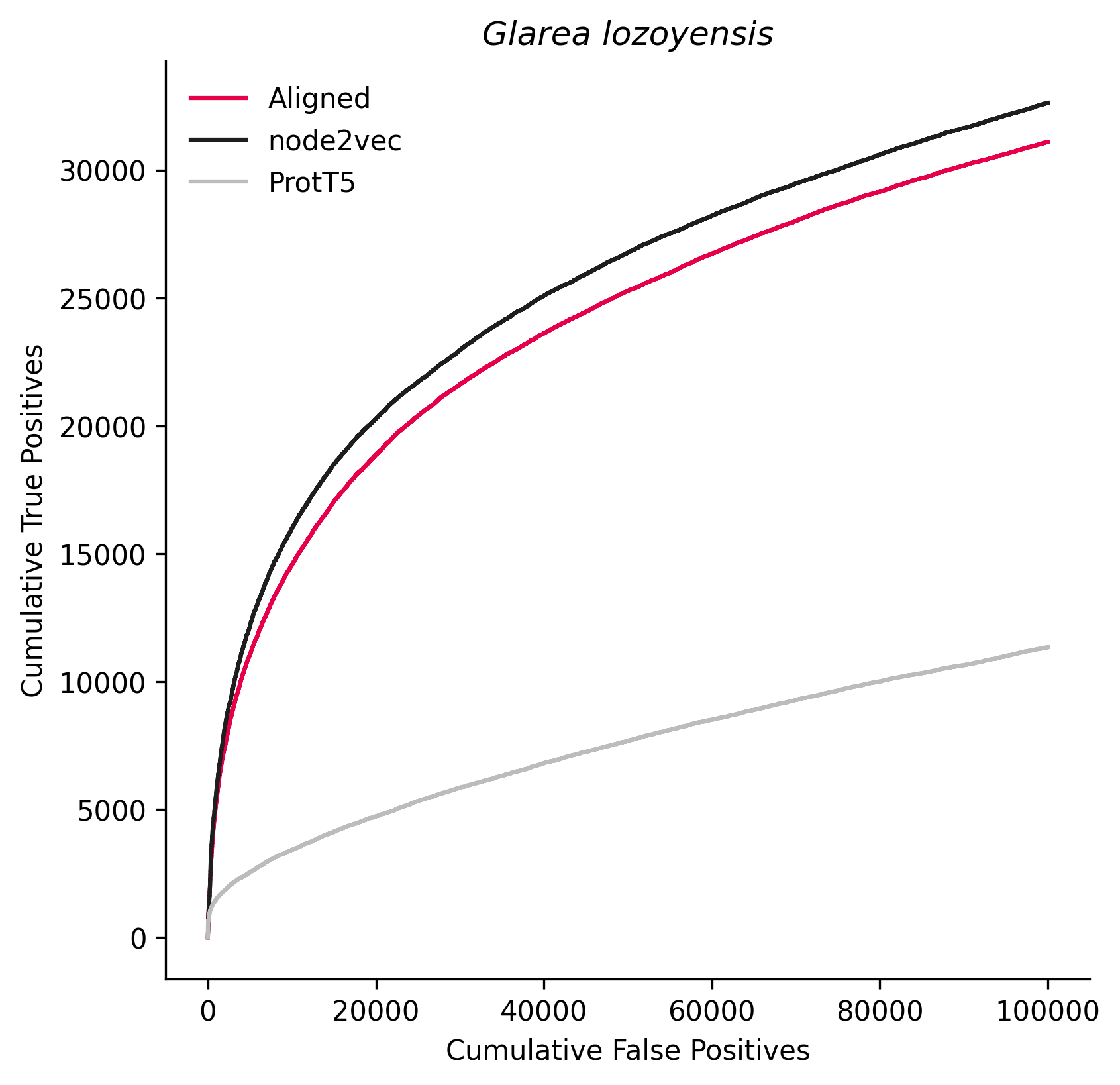

### 1257118.png

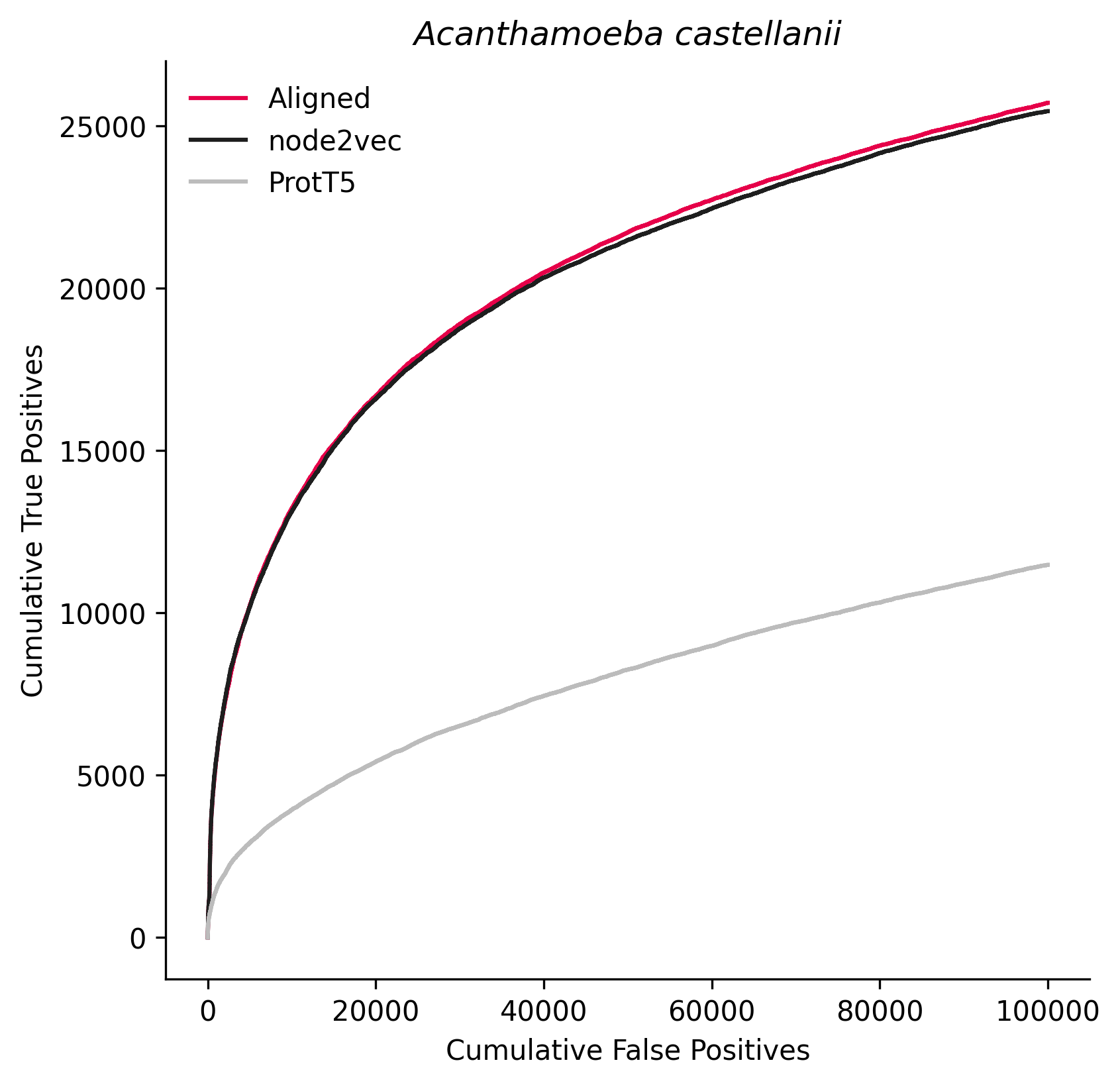

### 1287681.png

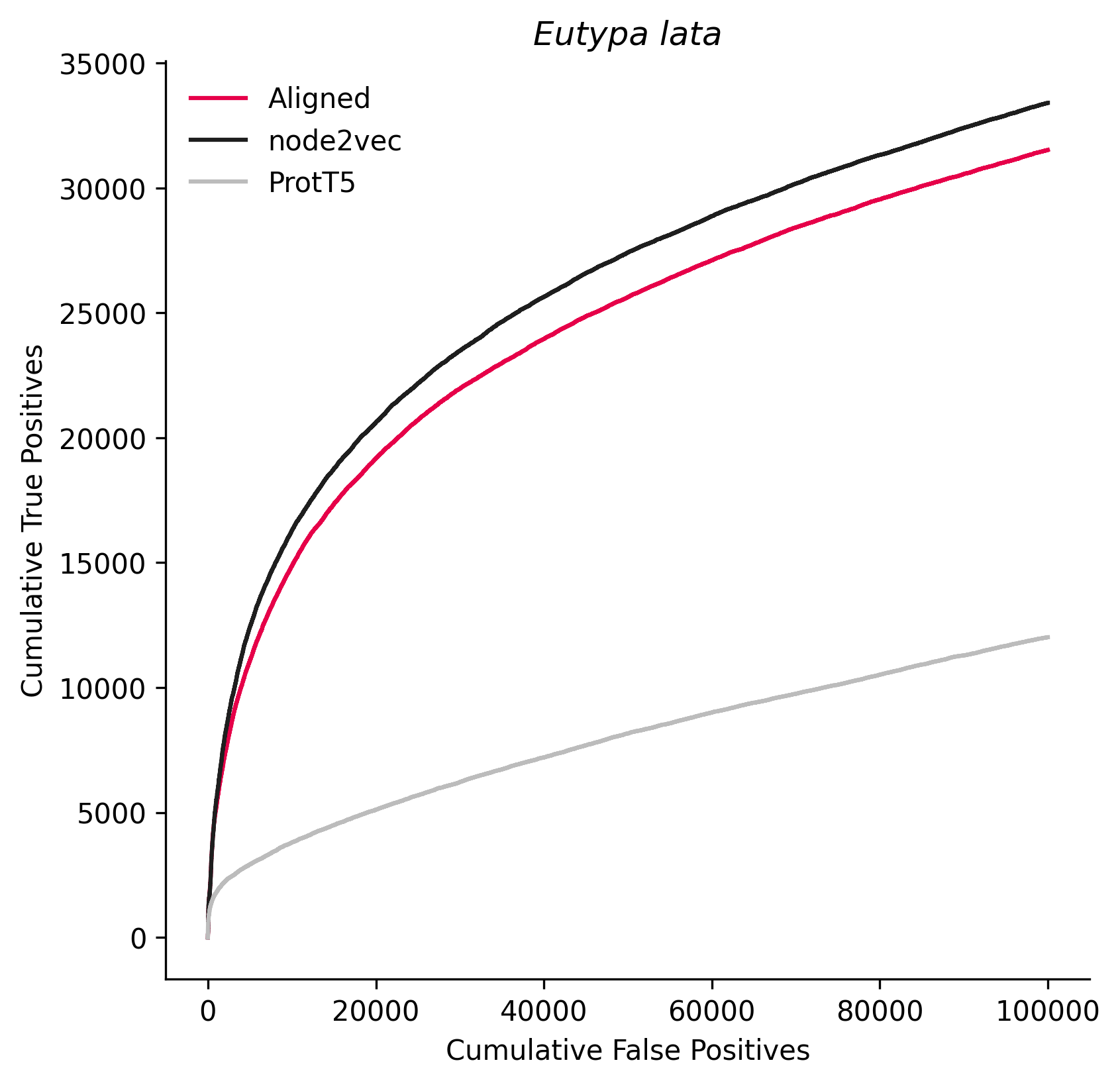

### 1299270.png

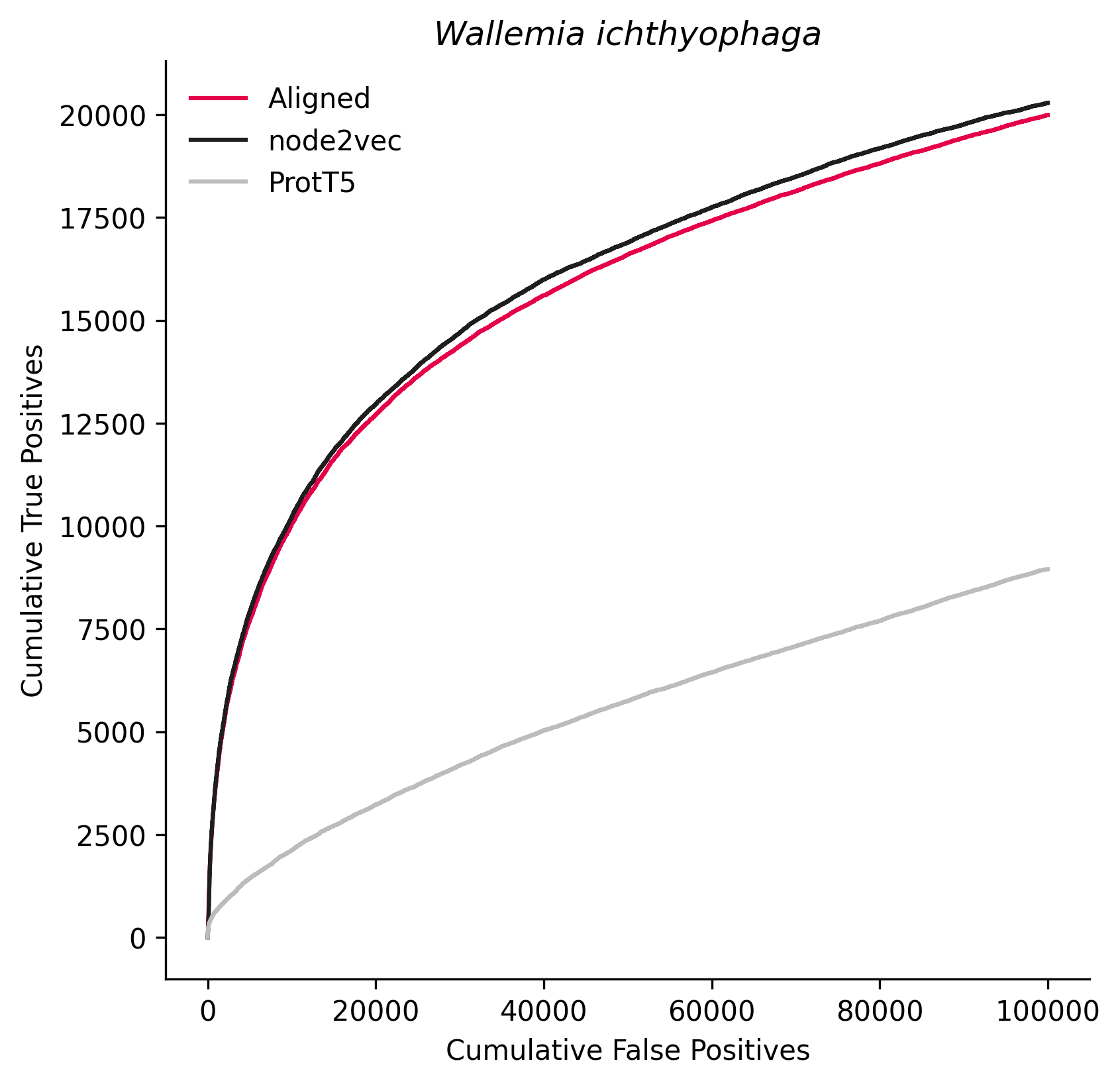

### 1445577.png

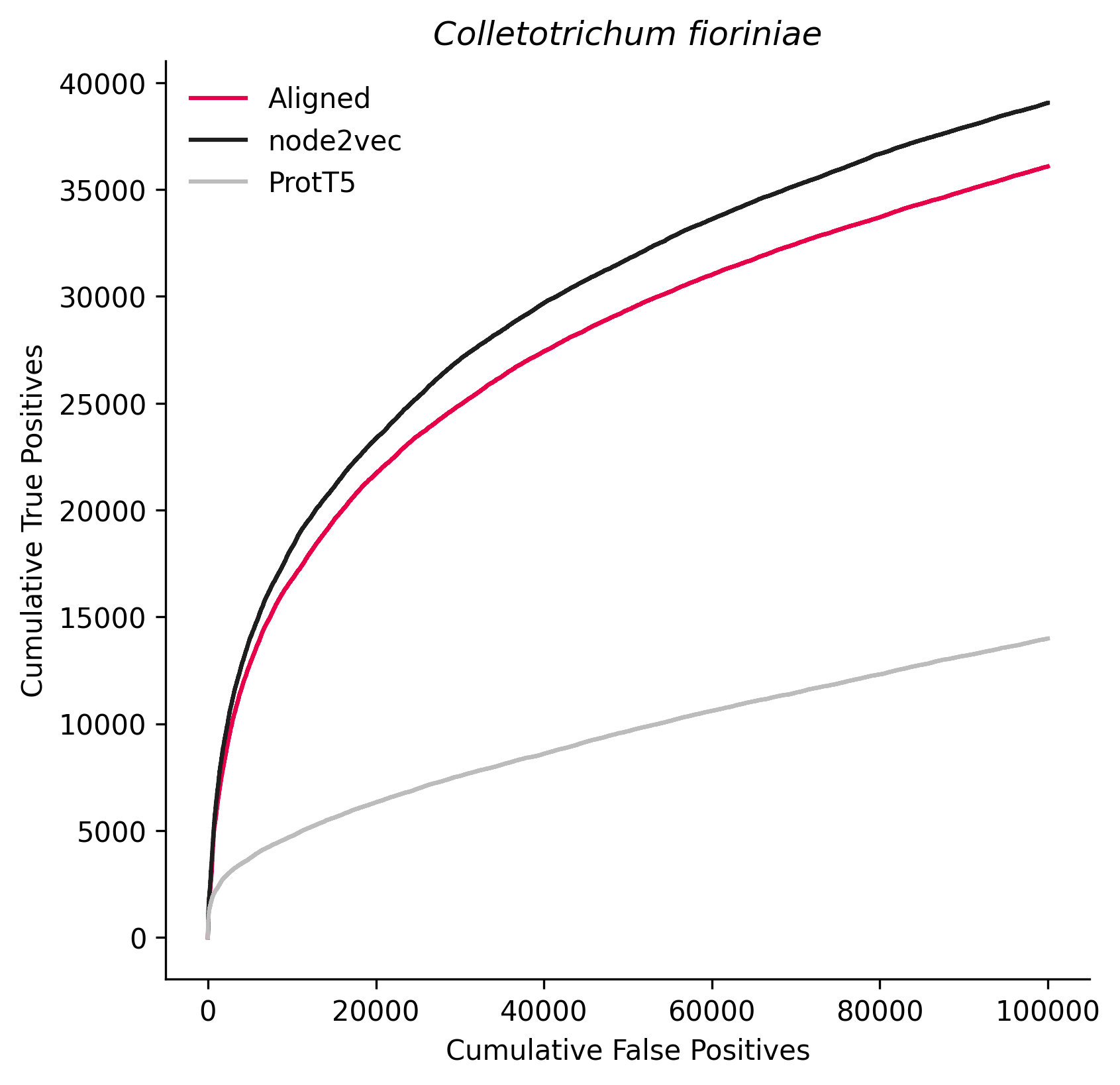

### 1537102.png

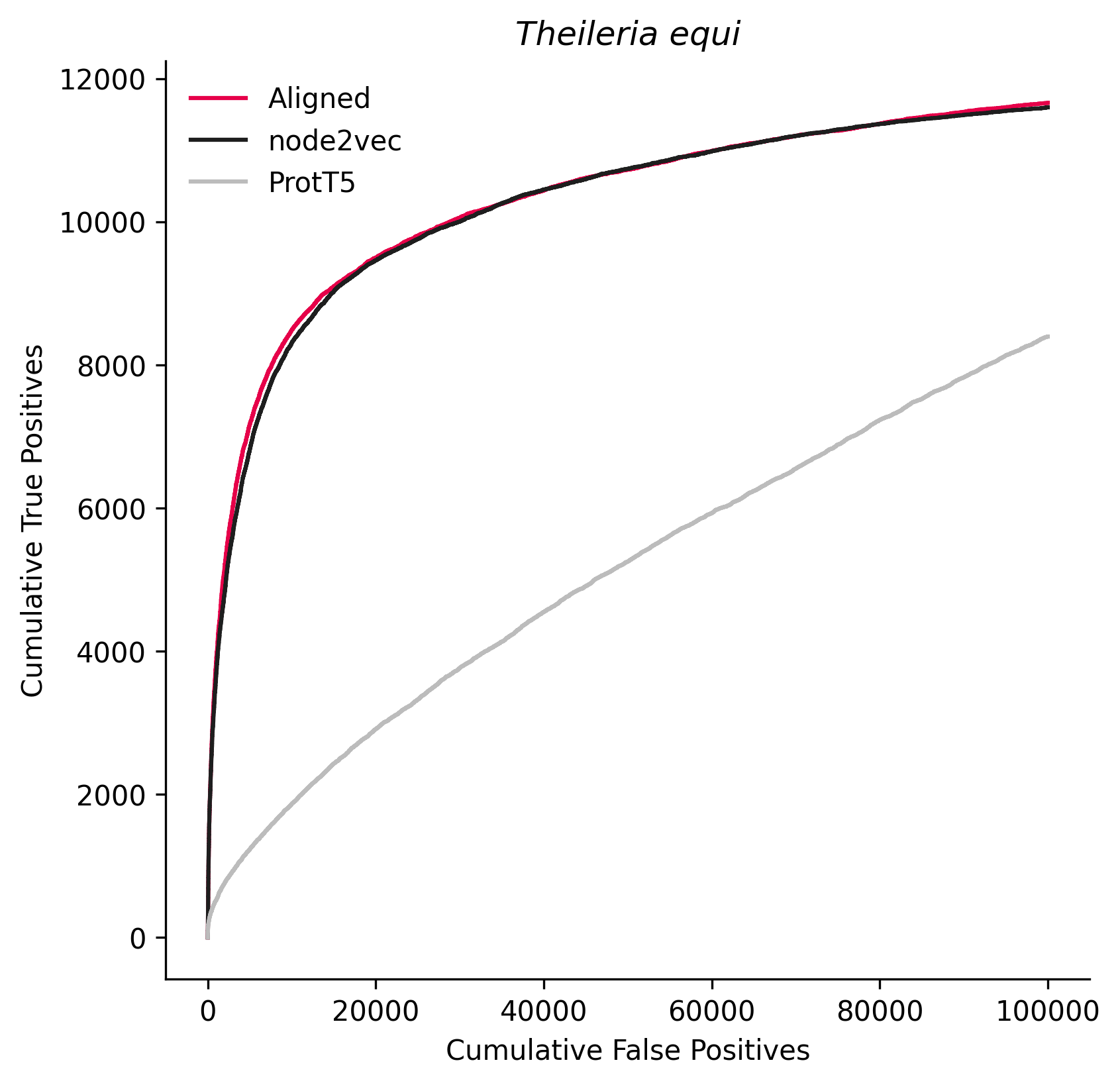

### 1841481.png

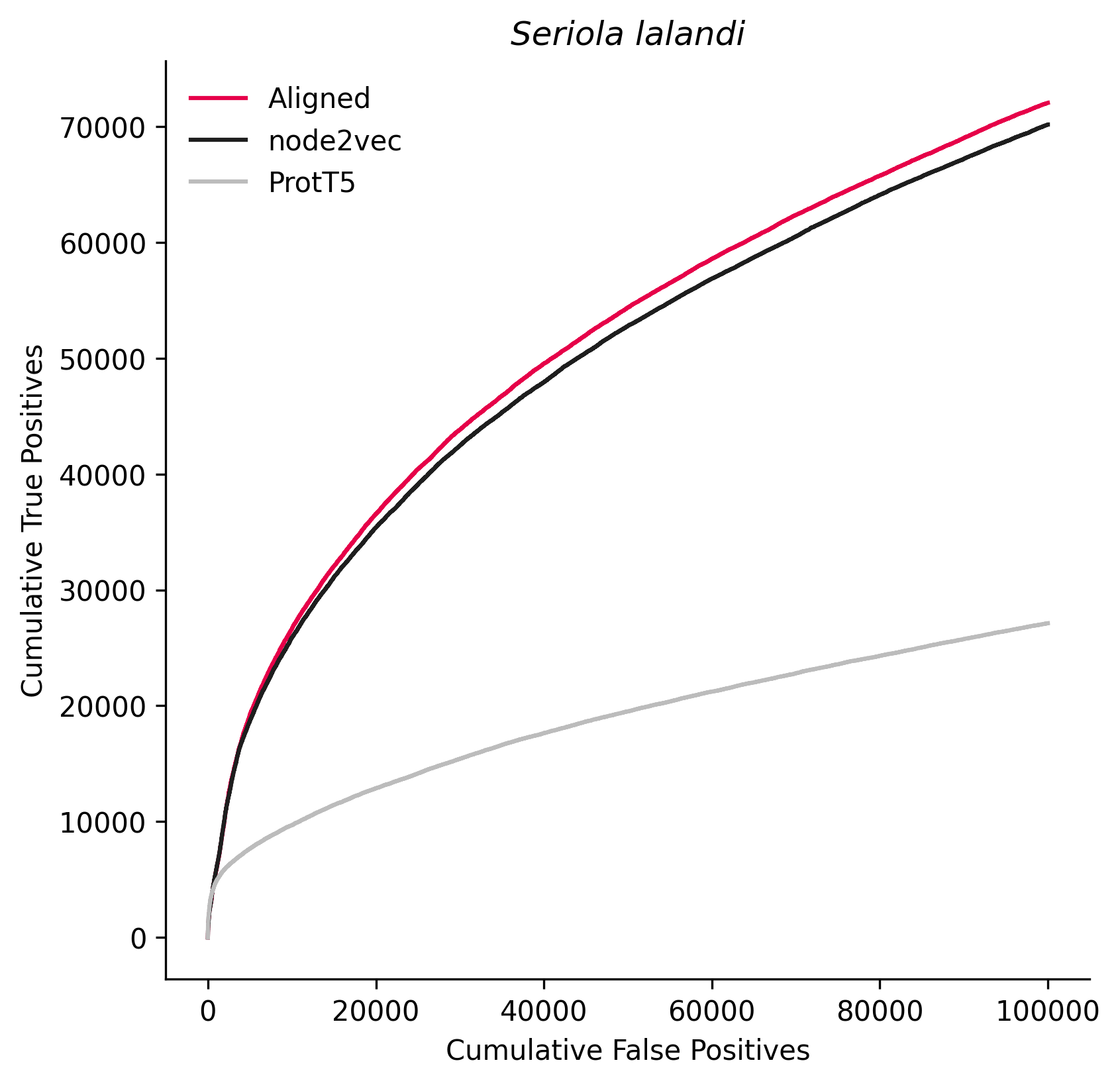
