## Supplementary figures and images for "SPACE: STRING proteins as complementary embeddings"

### 2903.png

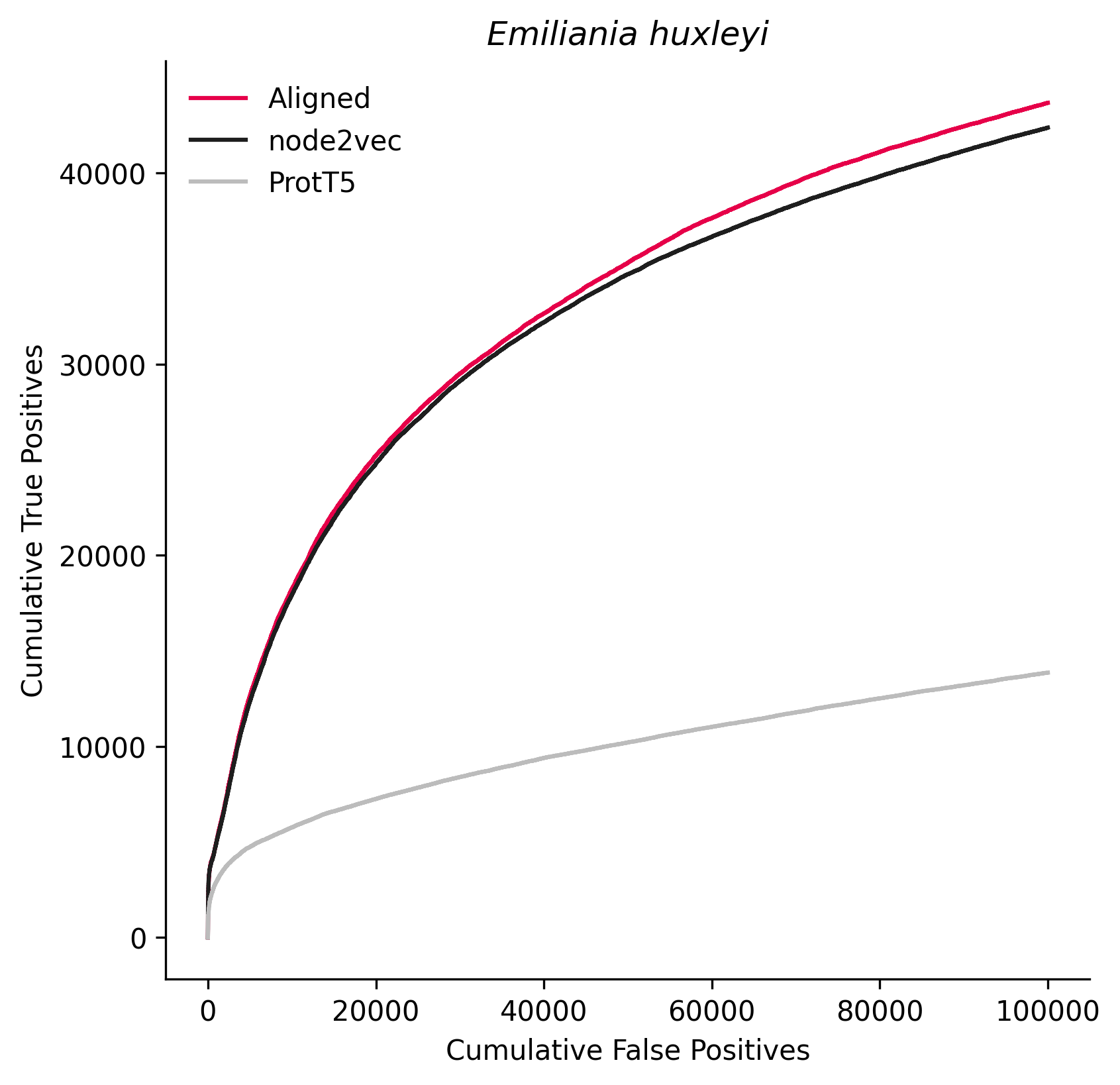

### 3708.png

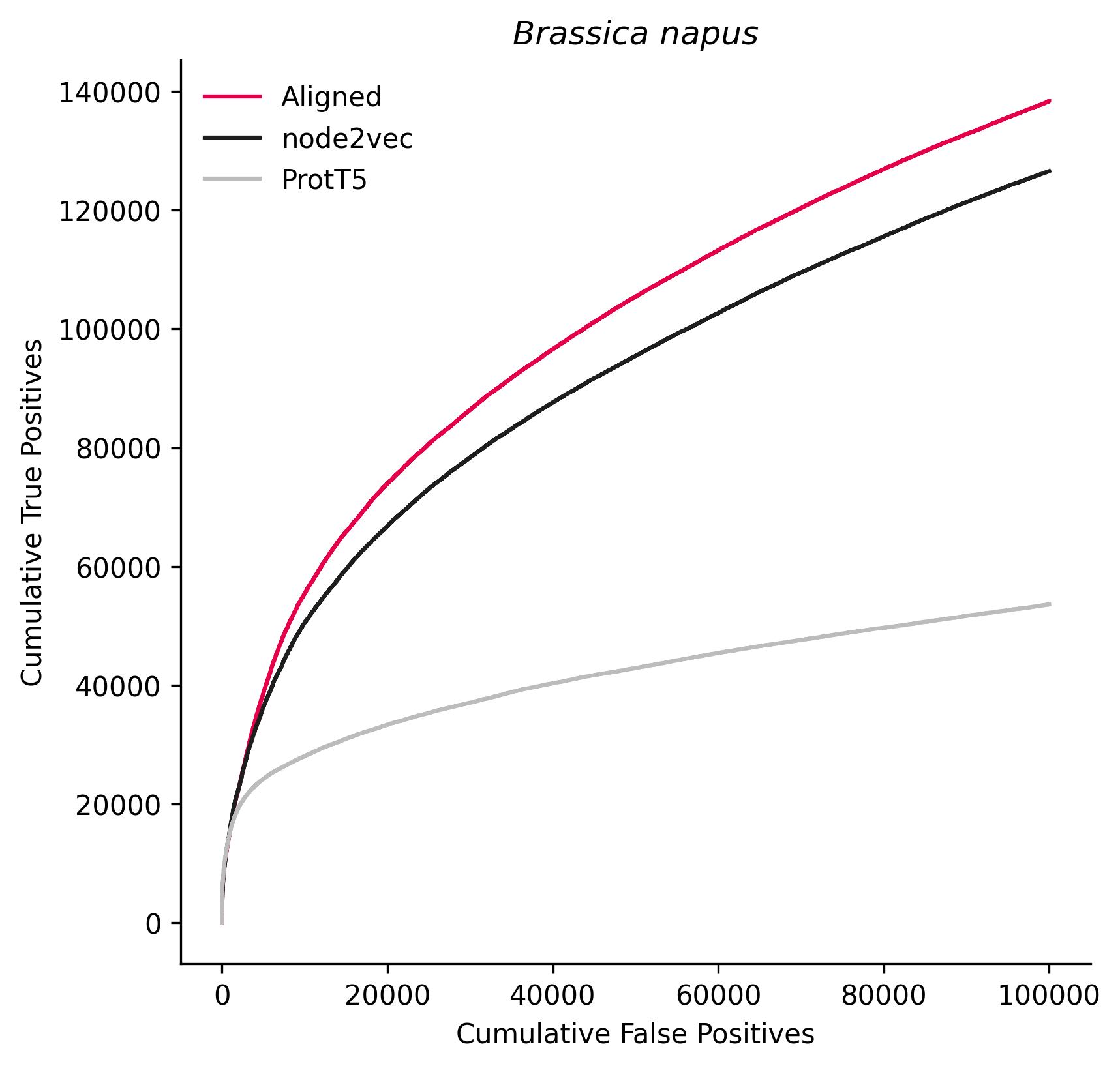

### 3827.png

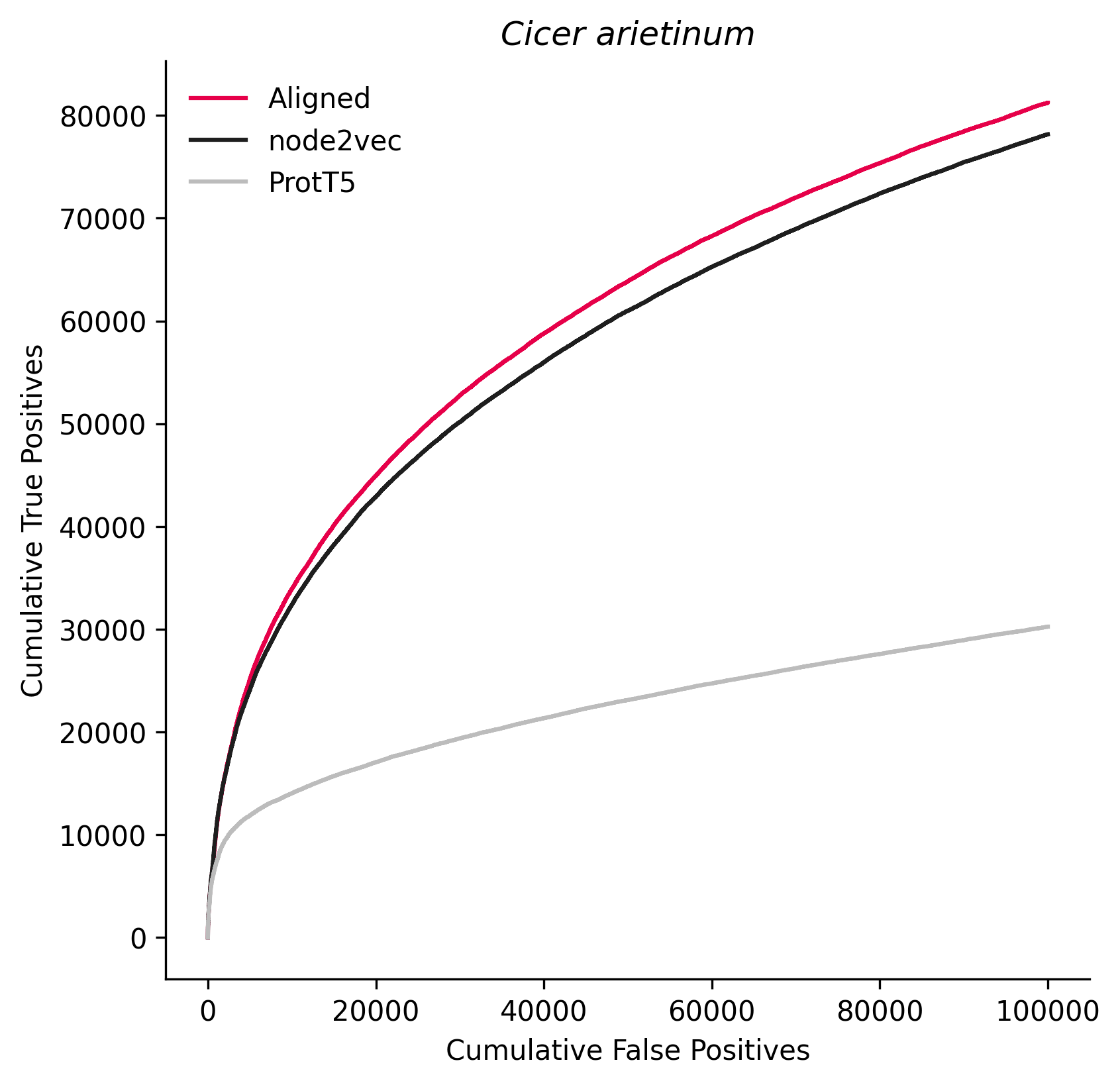

### 3983.png

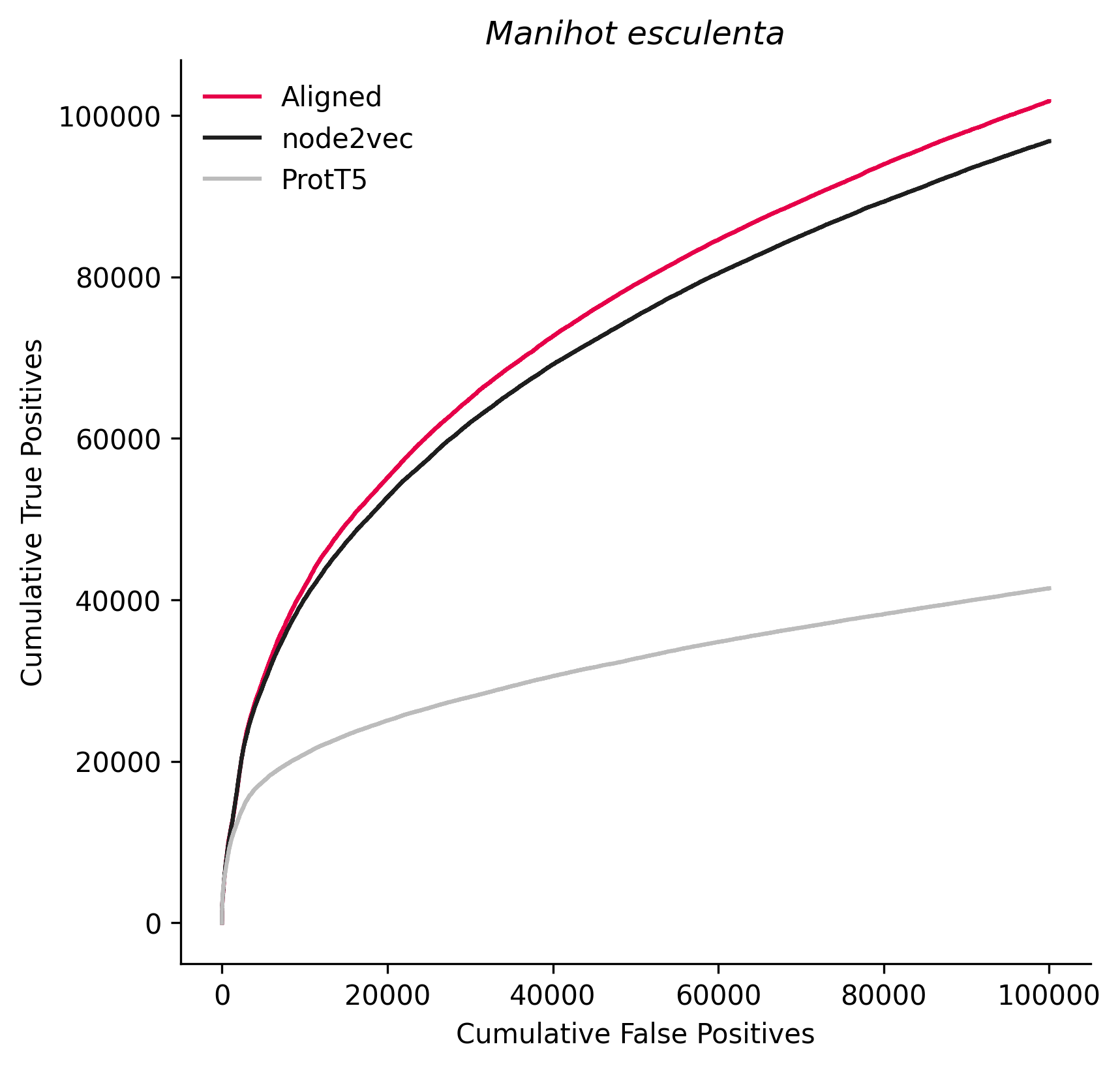

### 4072.png

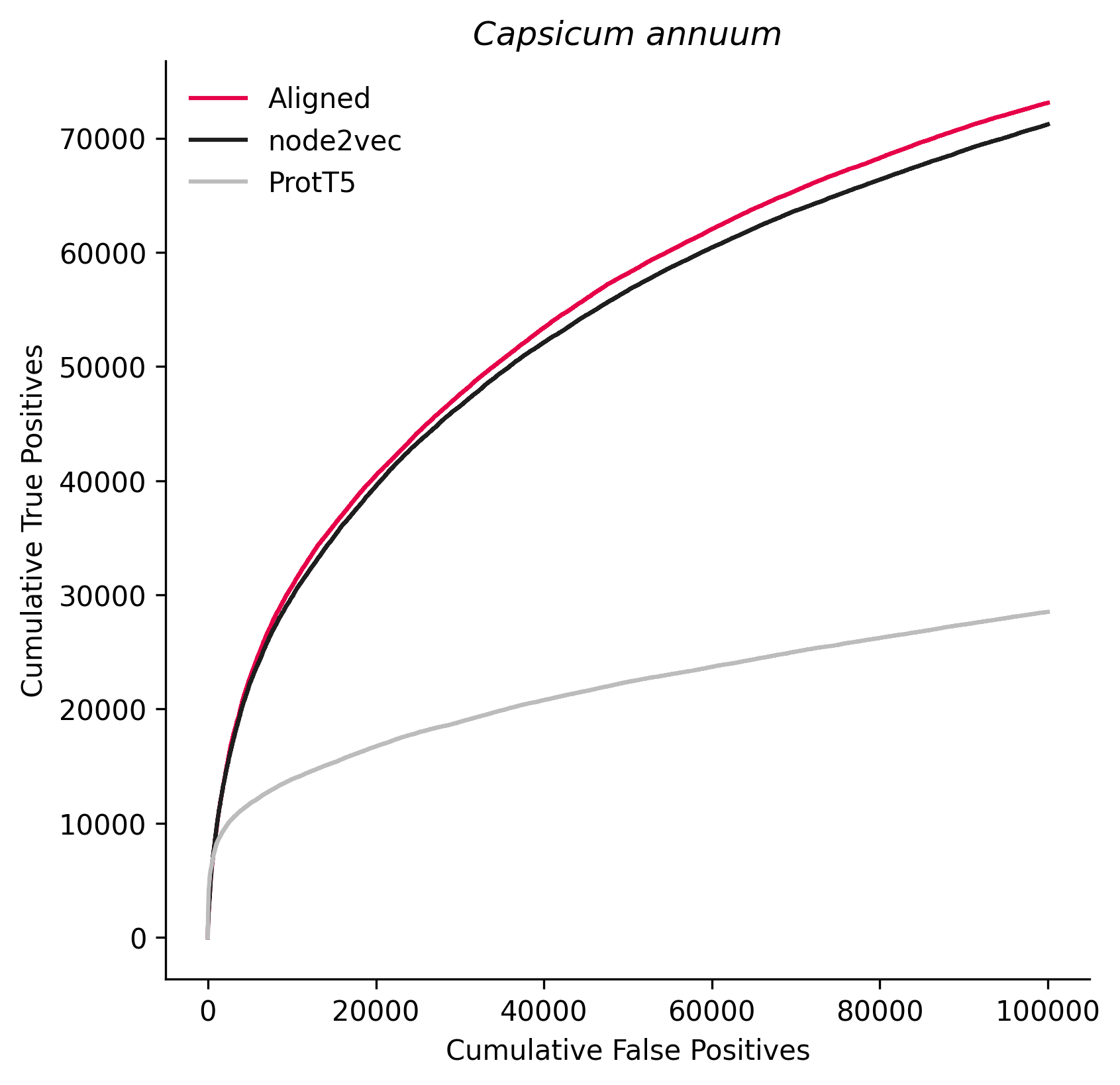

### 4081.png

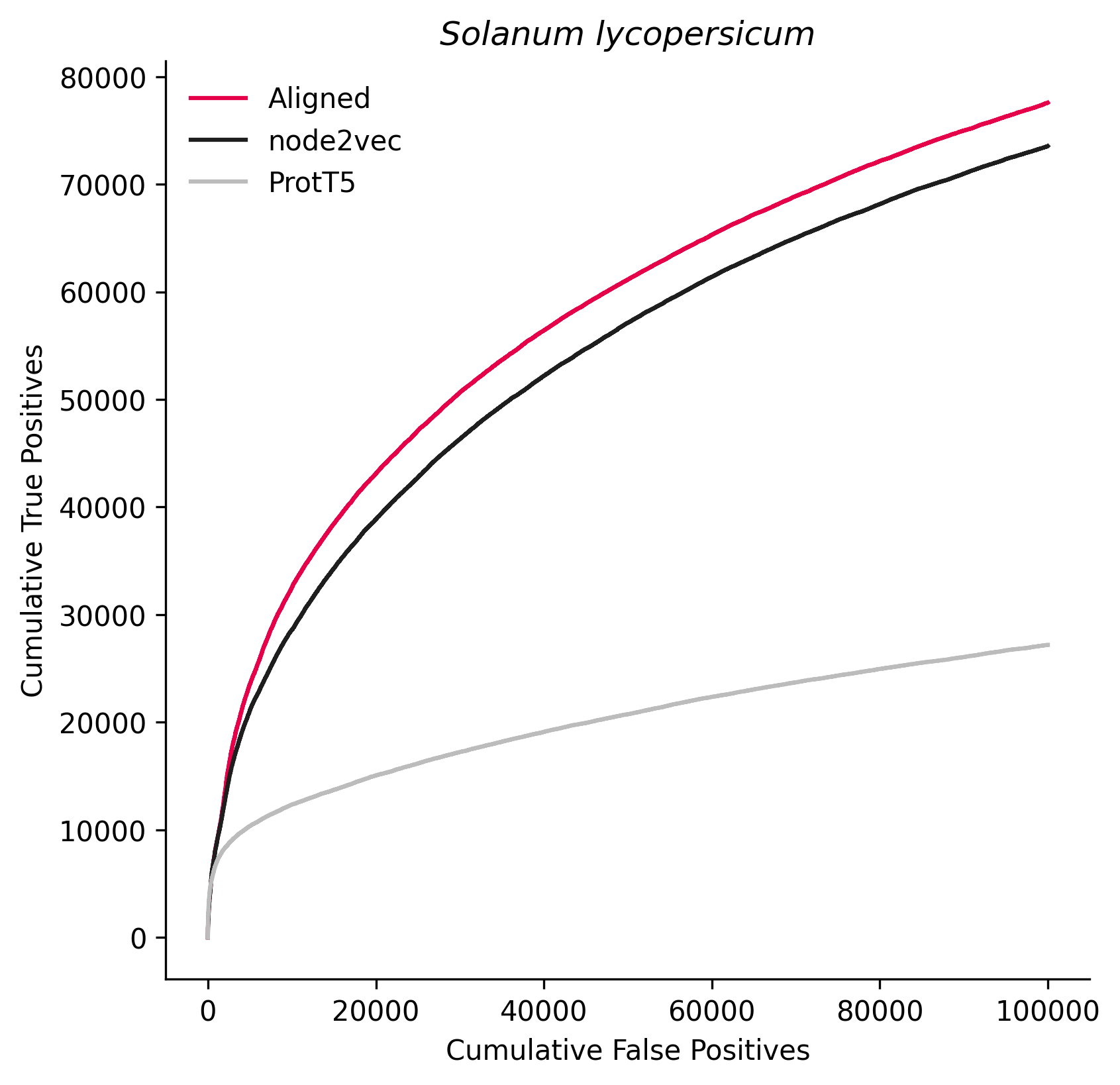

### 4232.png

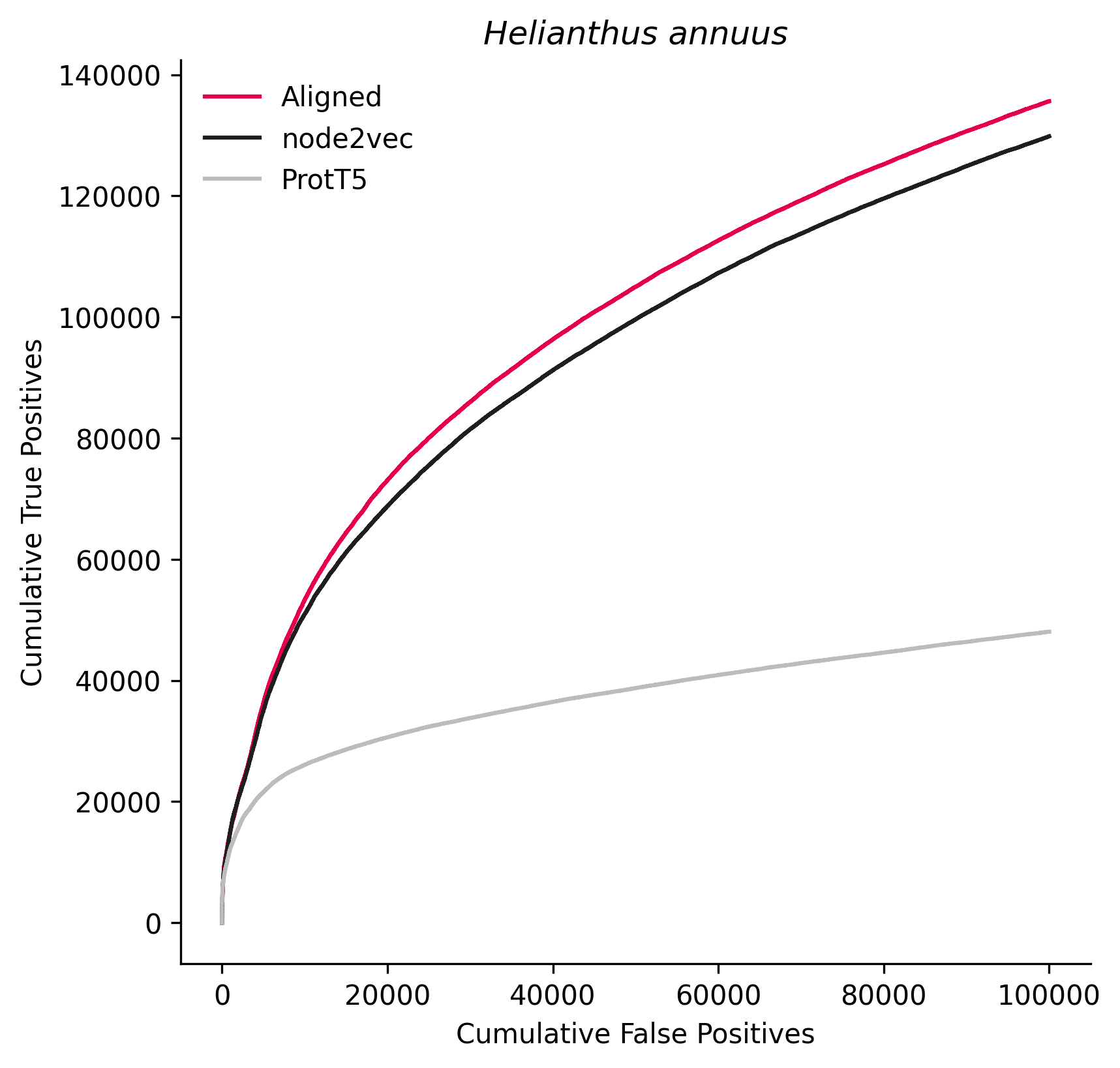

### 4577.png

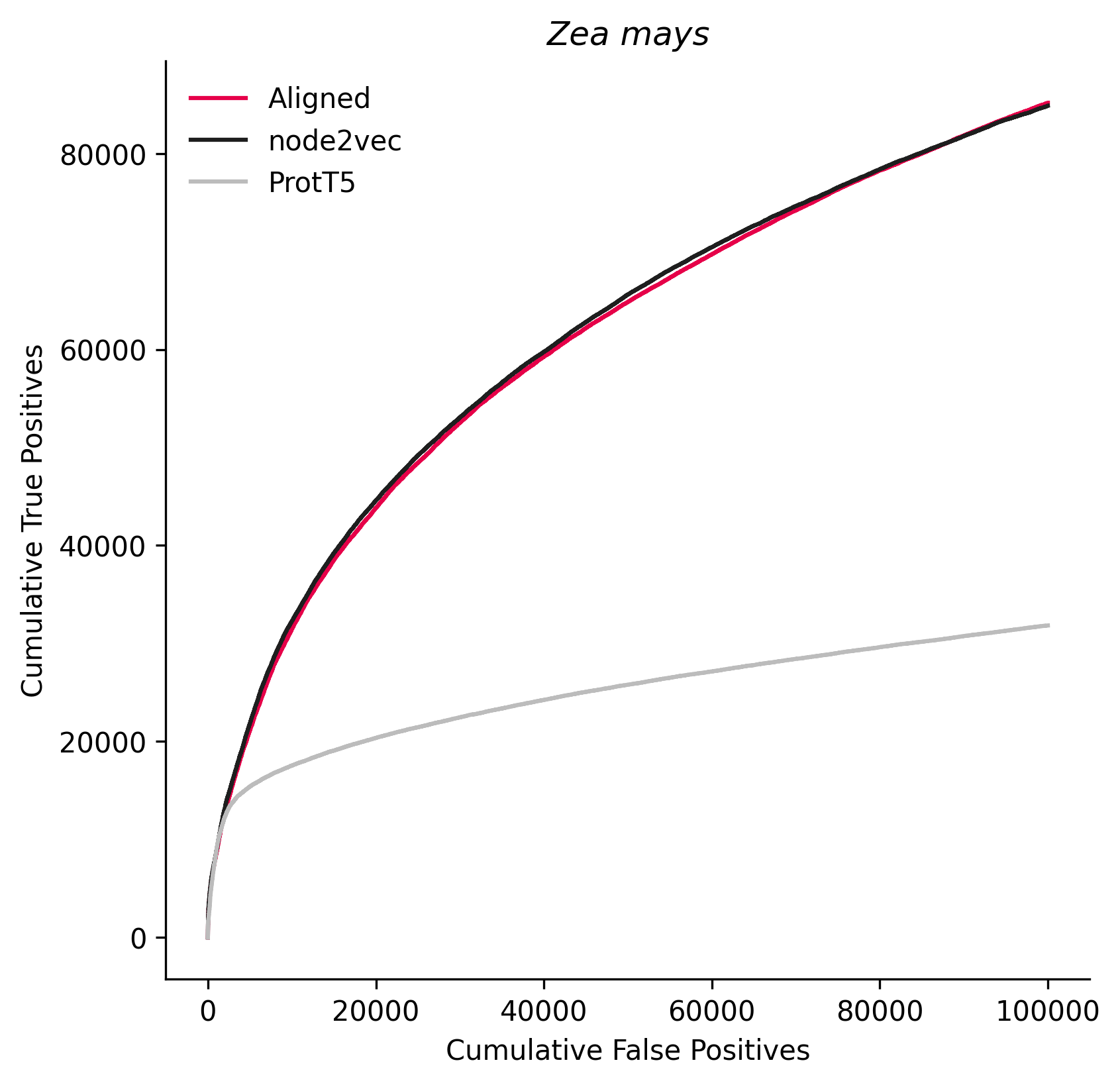

### 5599.png

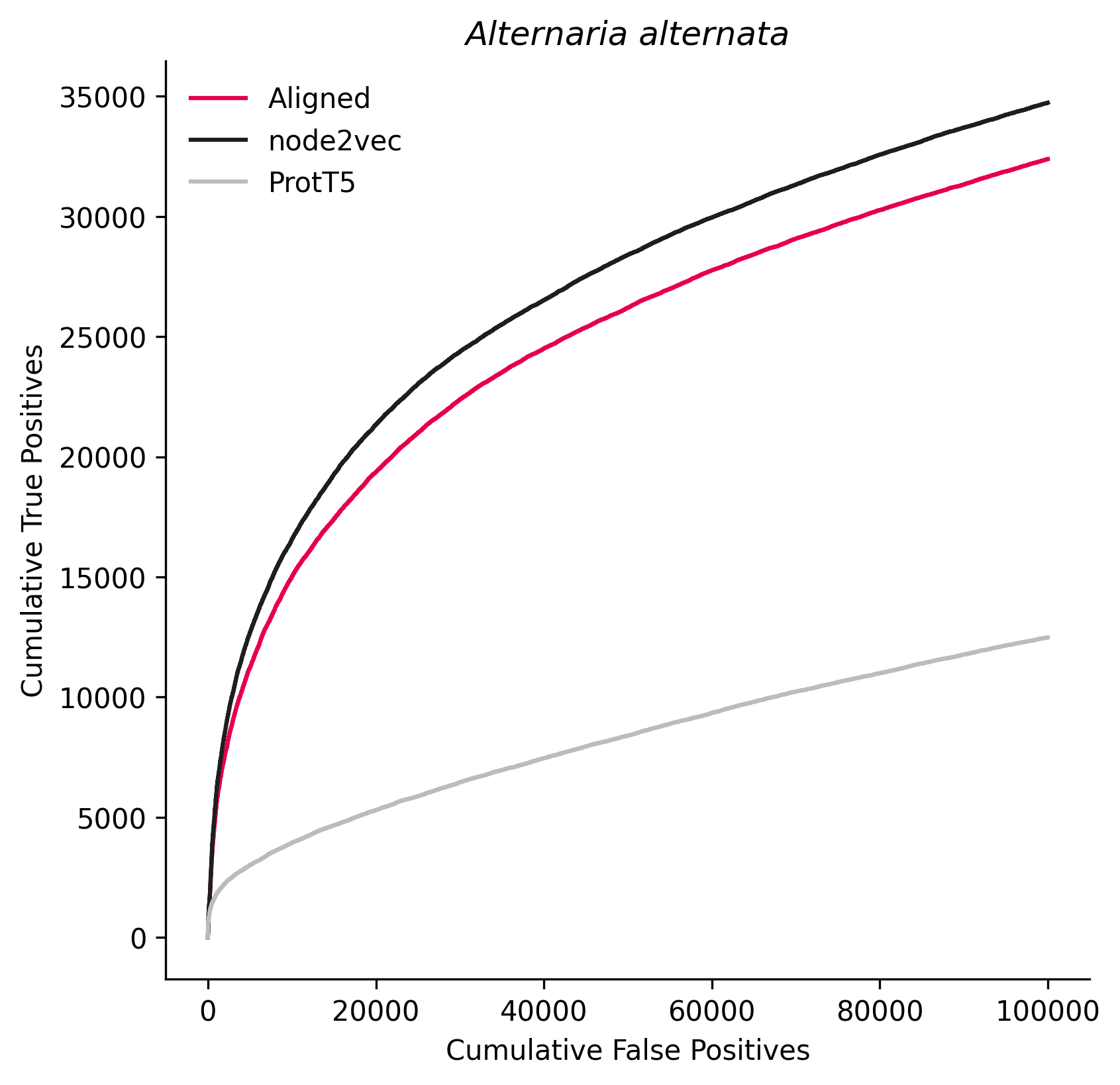

### 5759.png

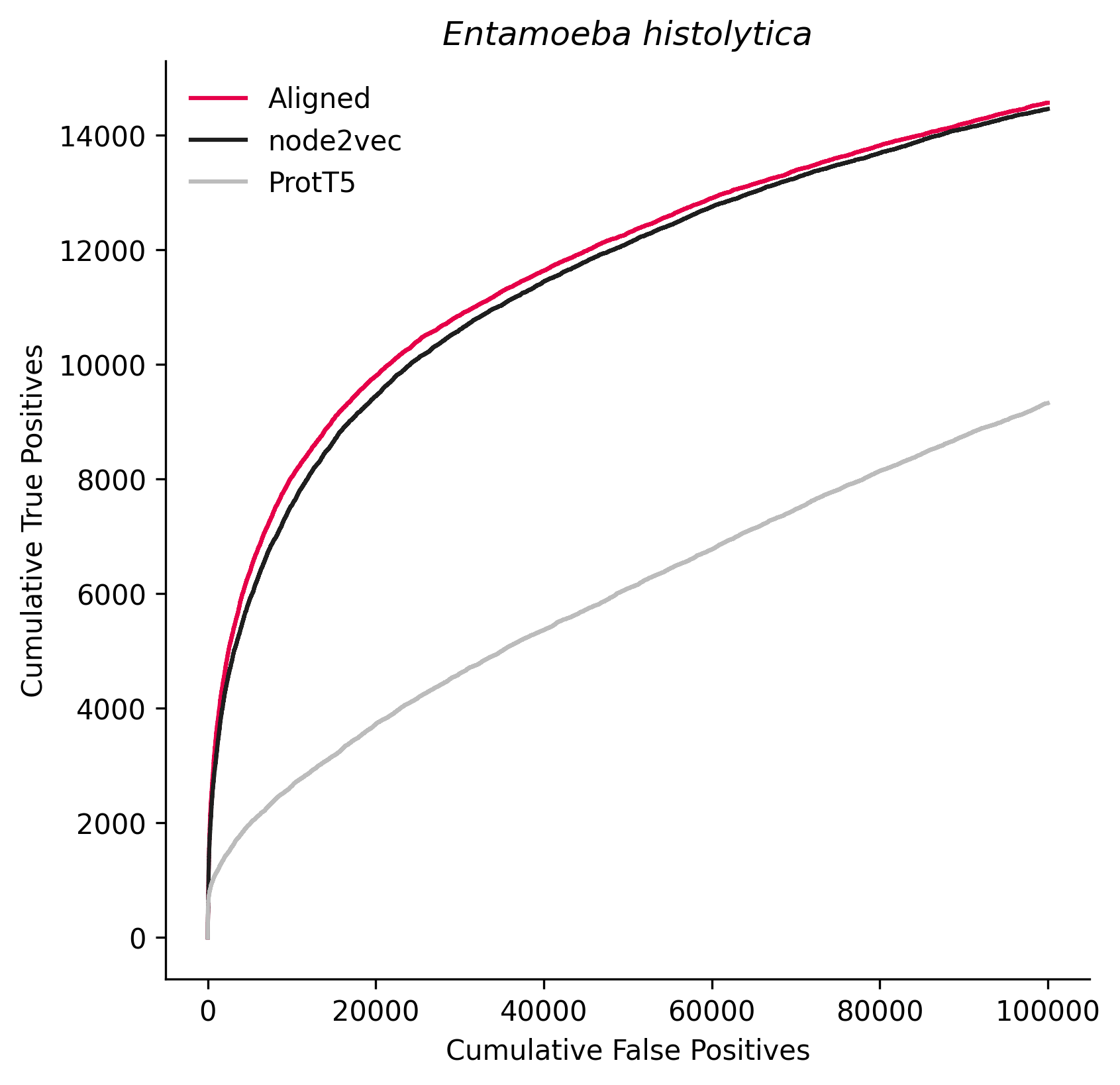

### 6185.png

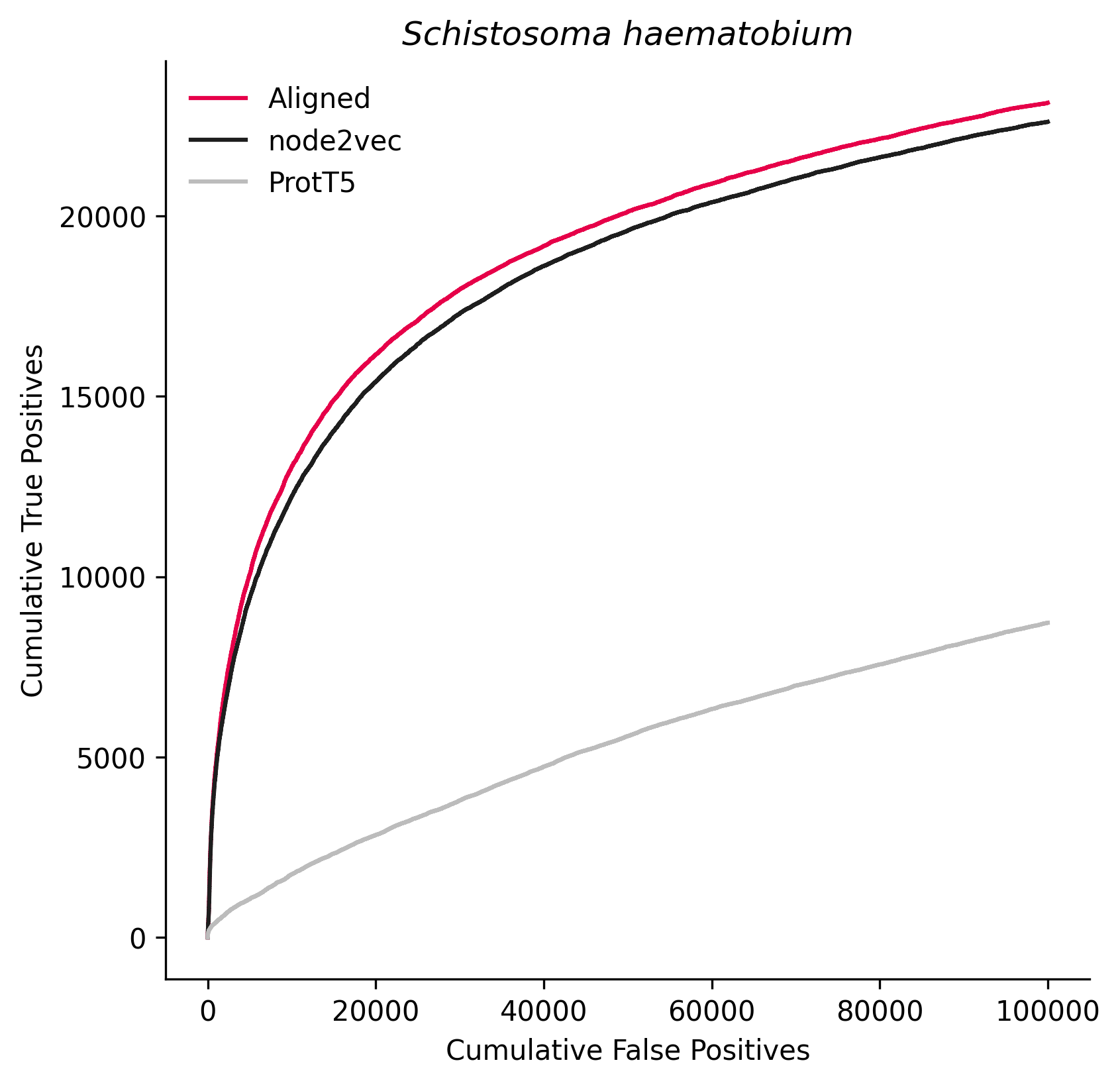

### 6198.png

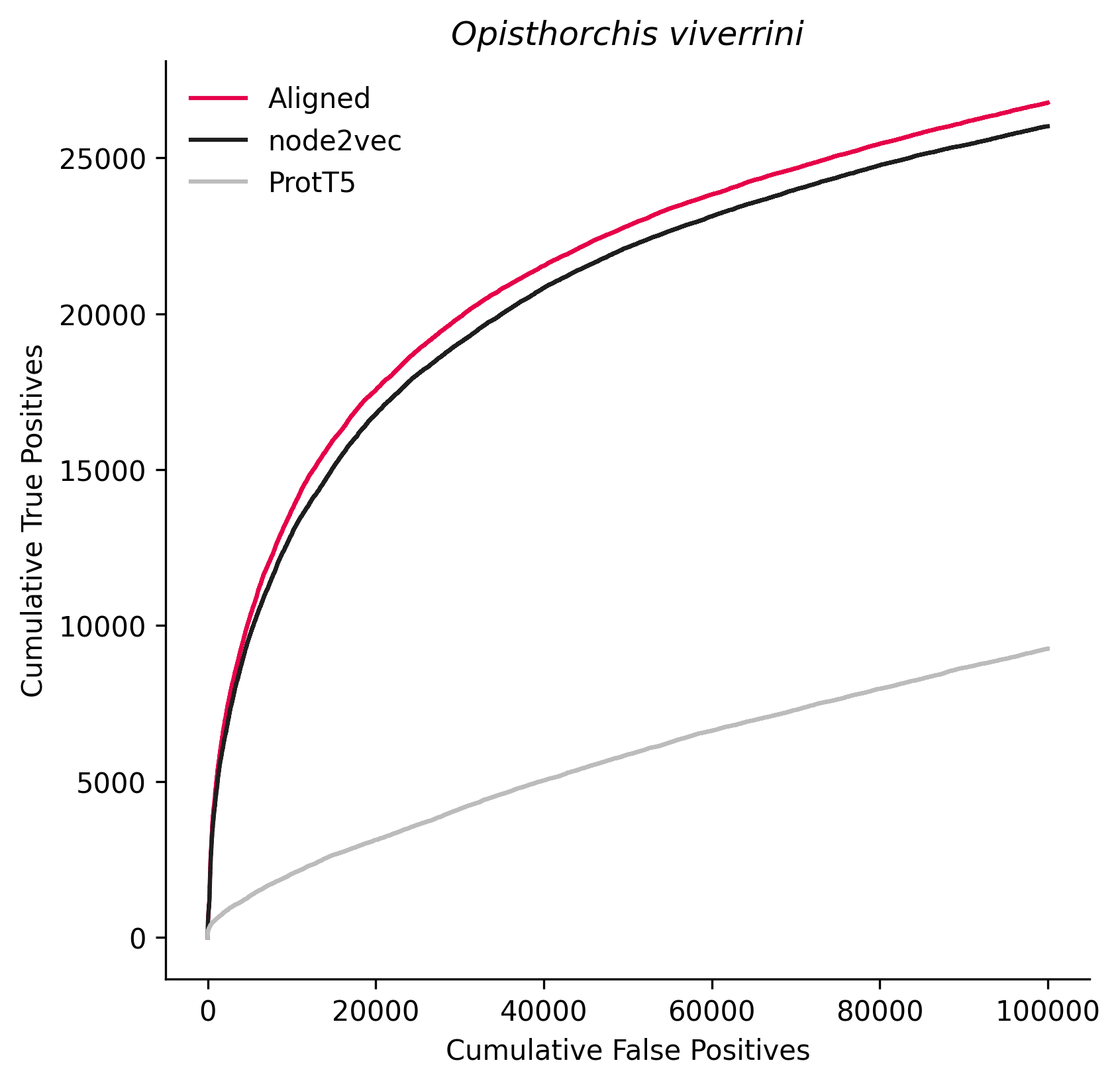

### 6279.png

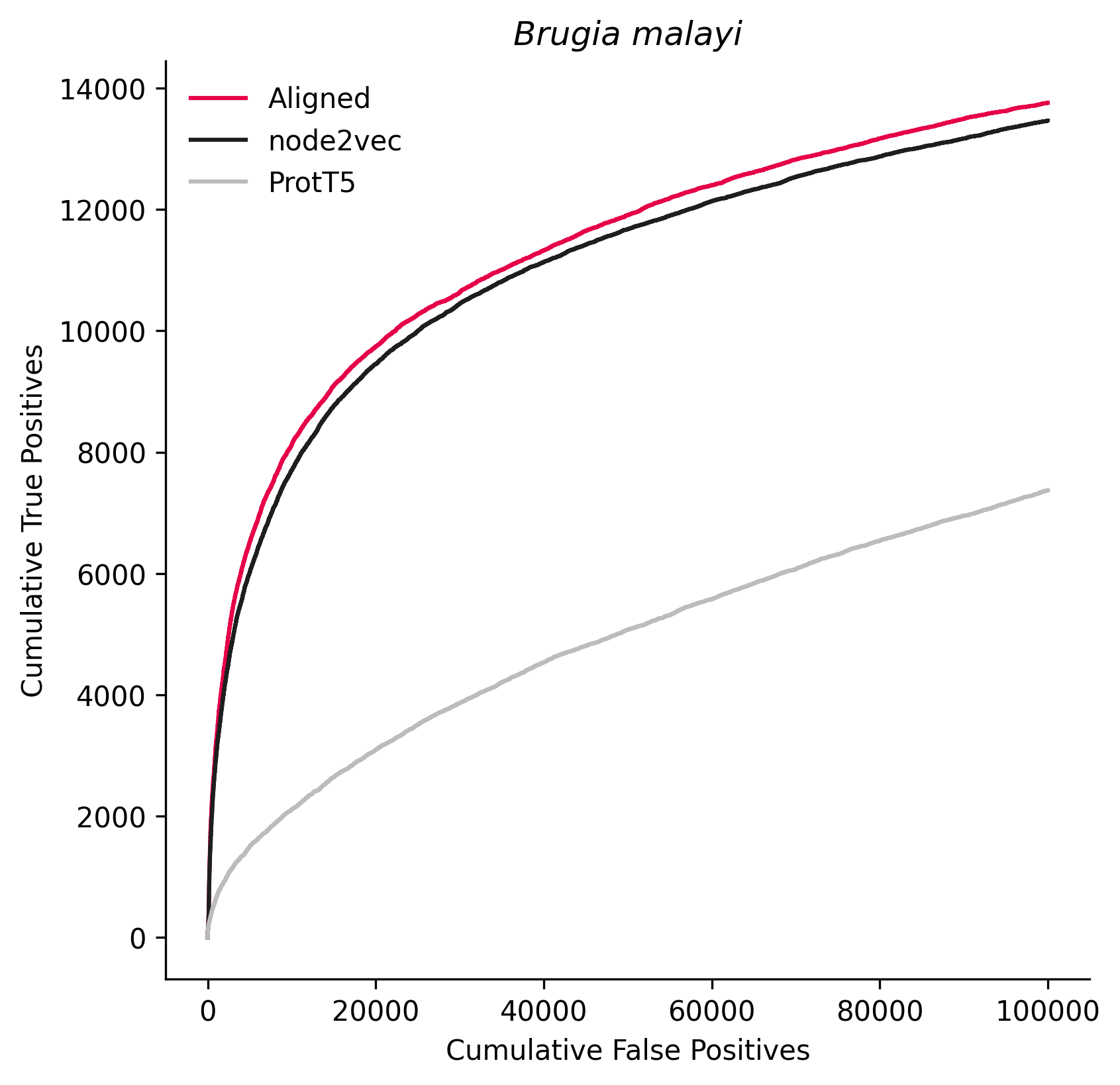

### 6412.png

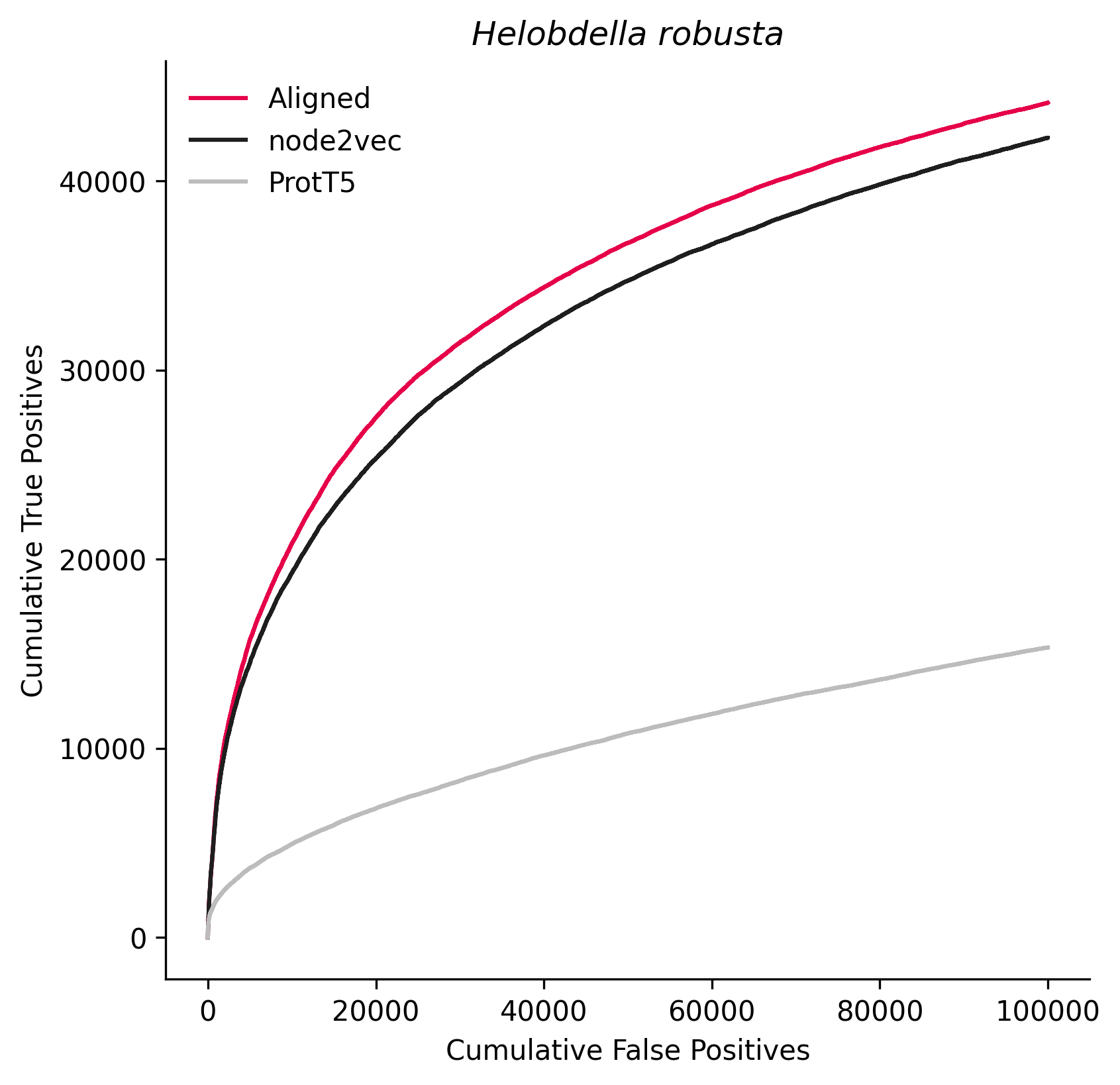

### 7070.png

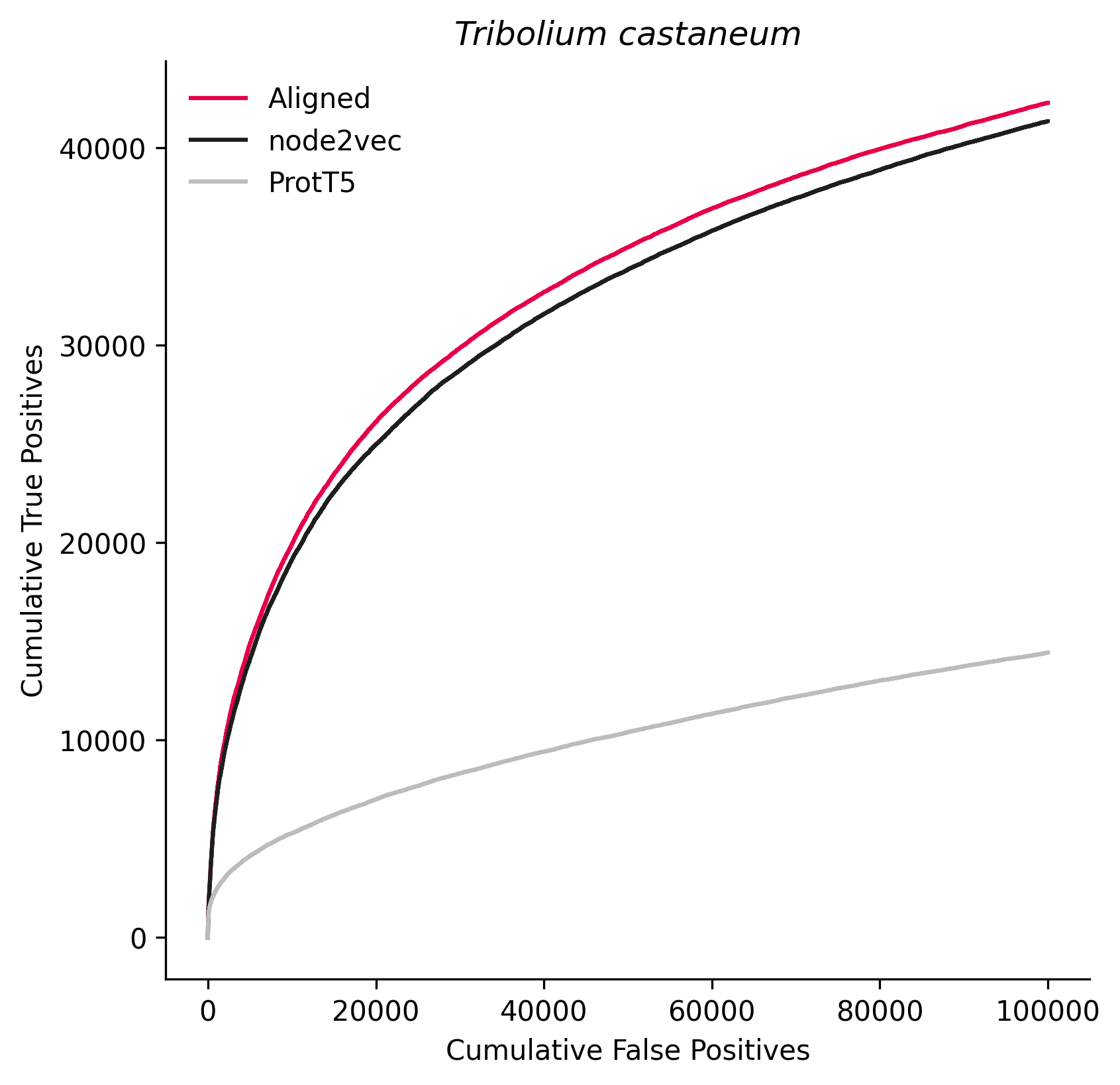

### 7227.png

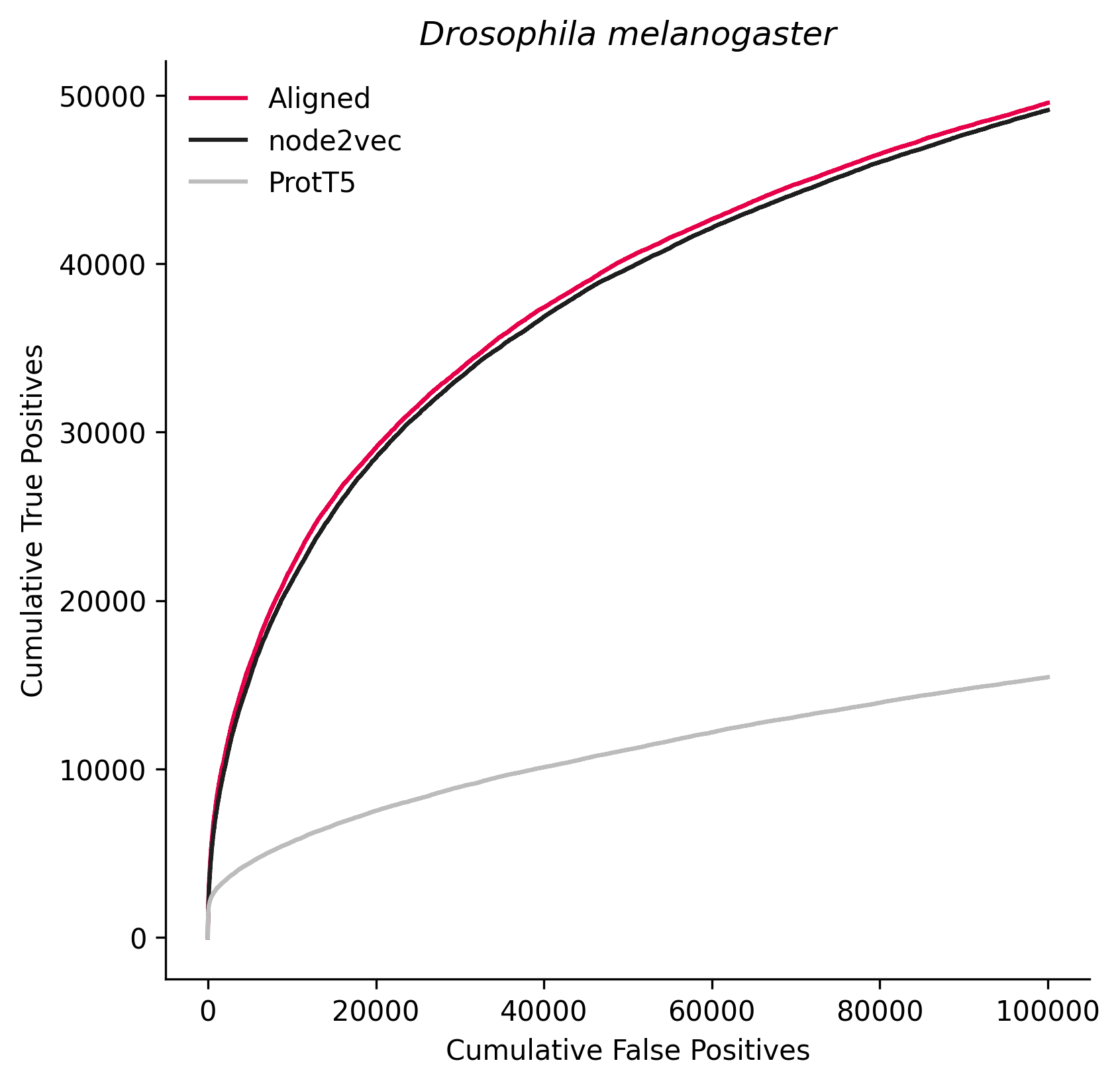

### 7240.png

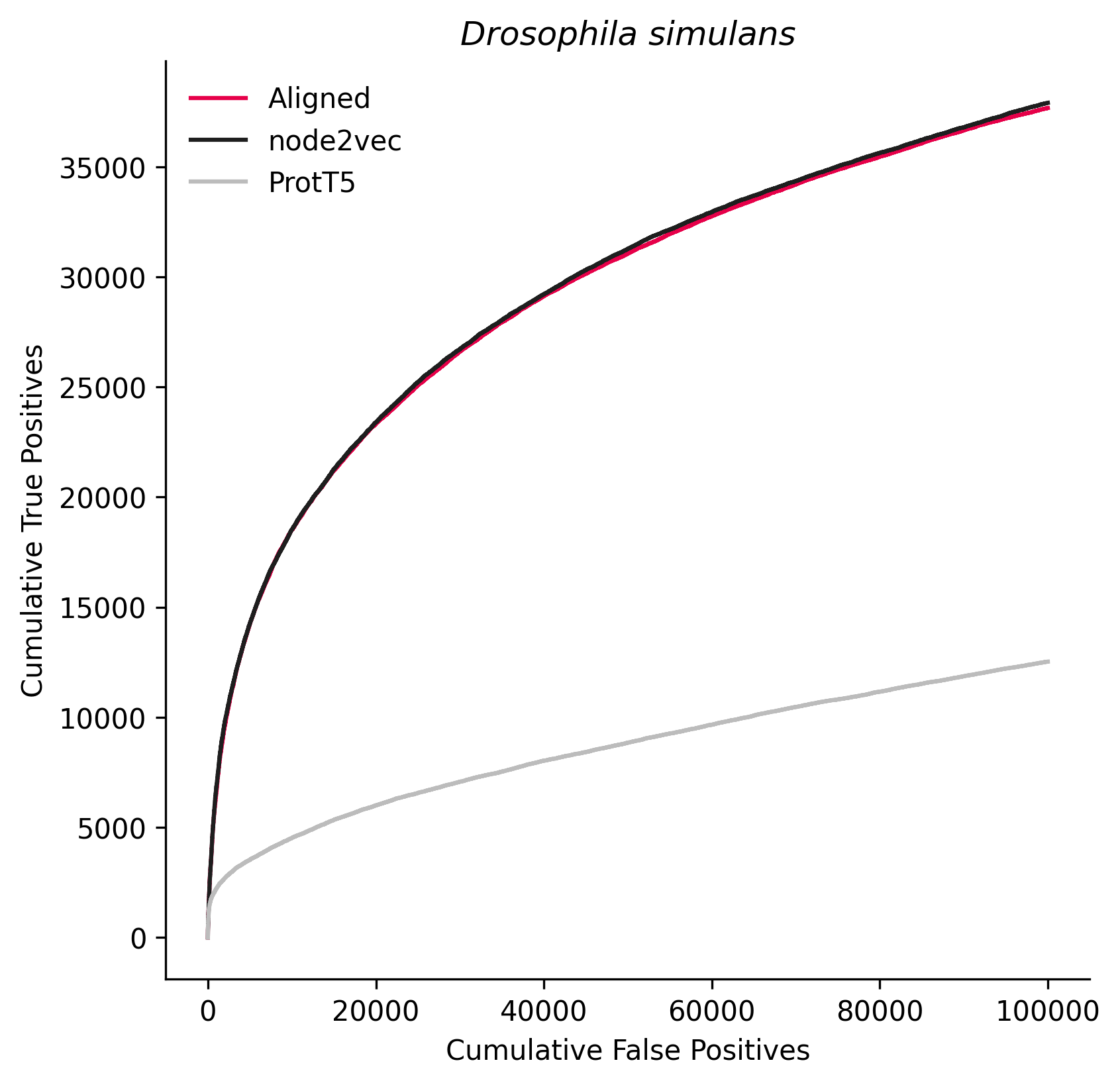

### 7244.png

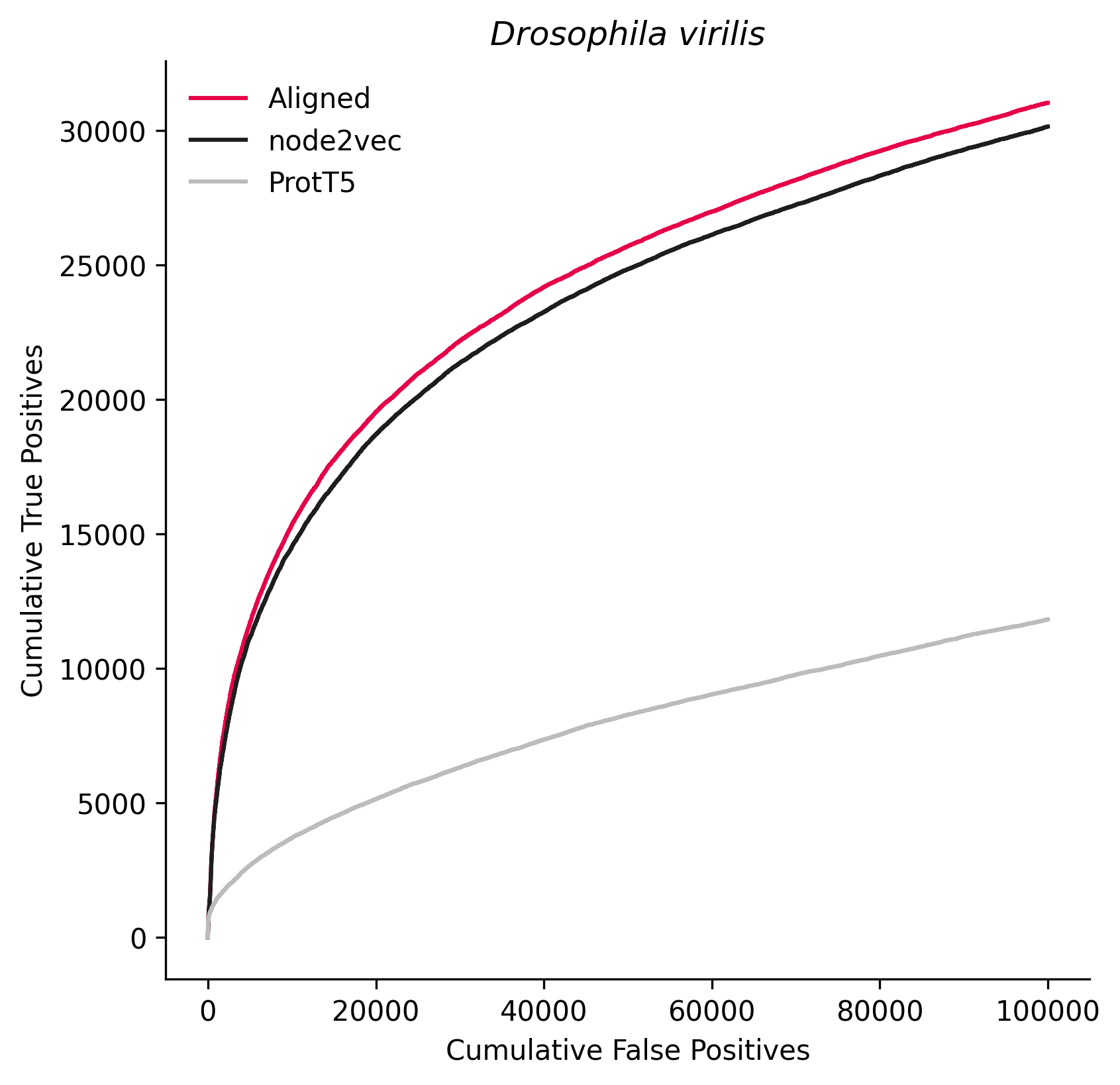

### 7260.png

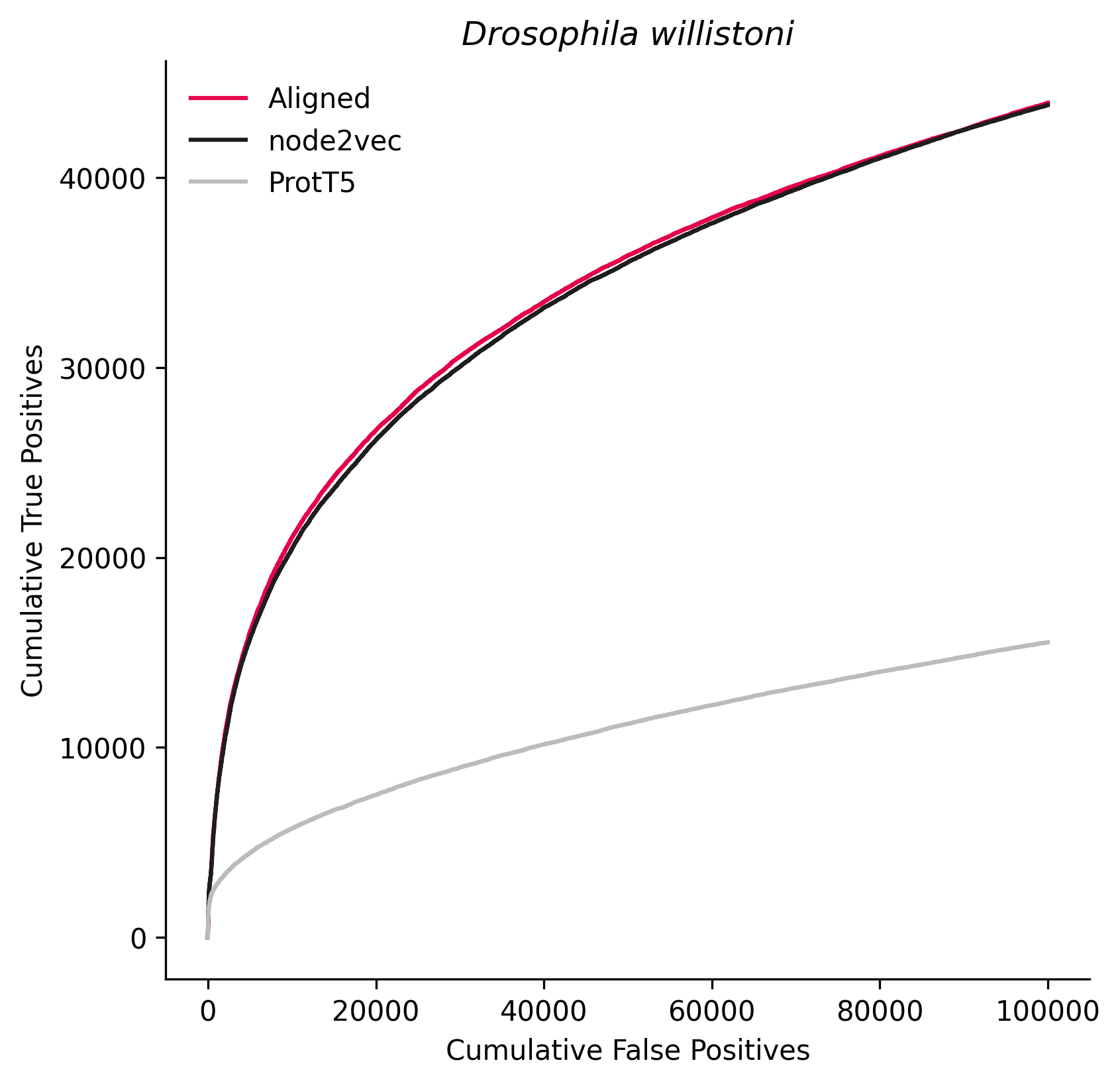

### 7460.png

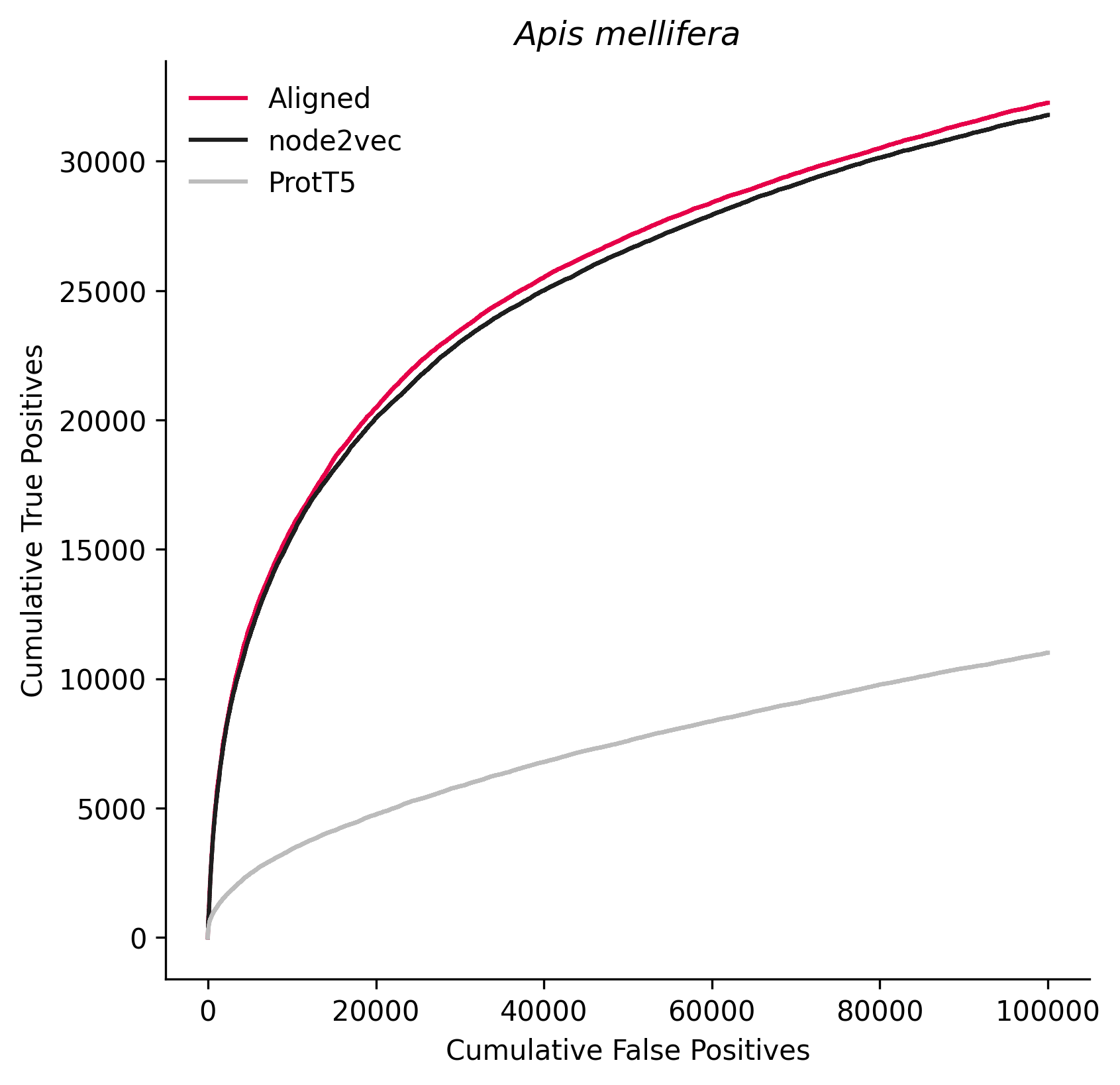

### 7955.png

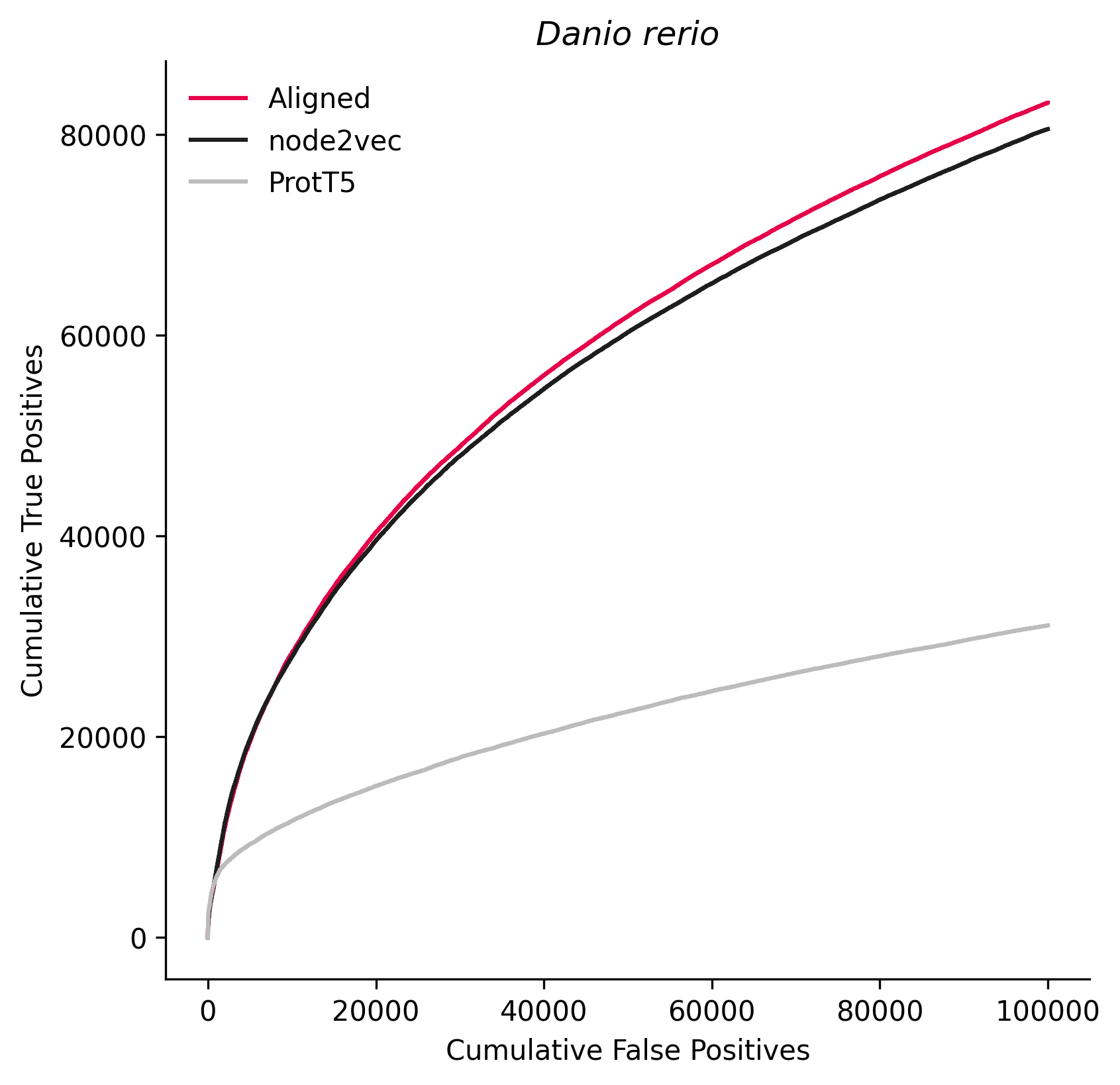

### 7998.png

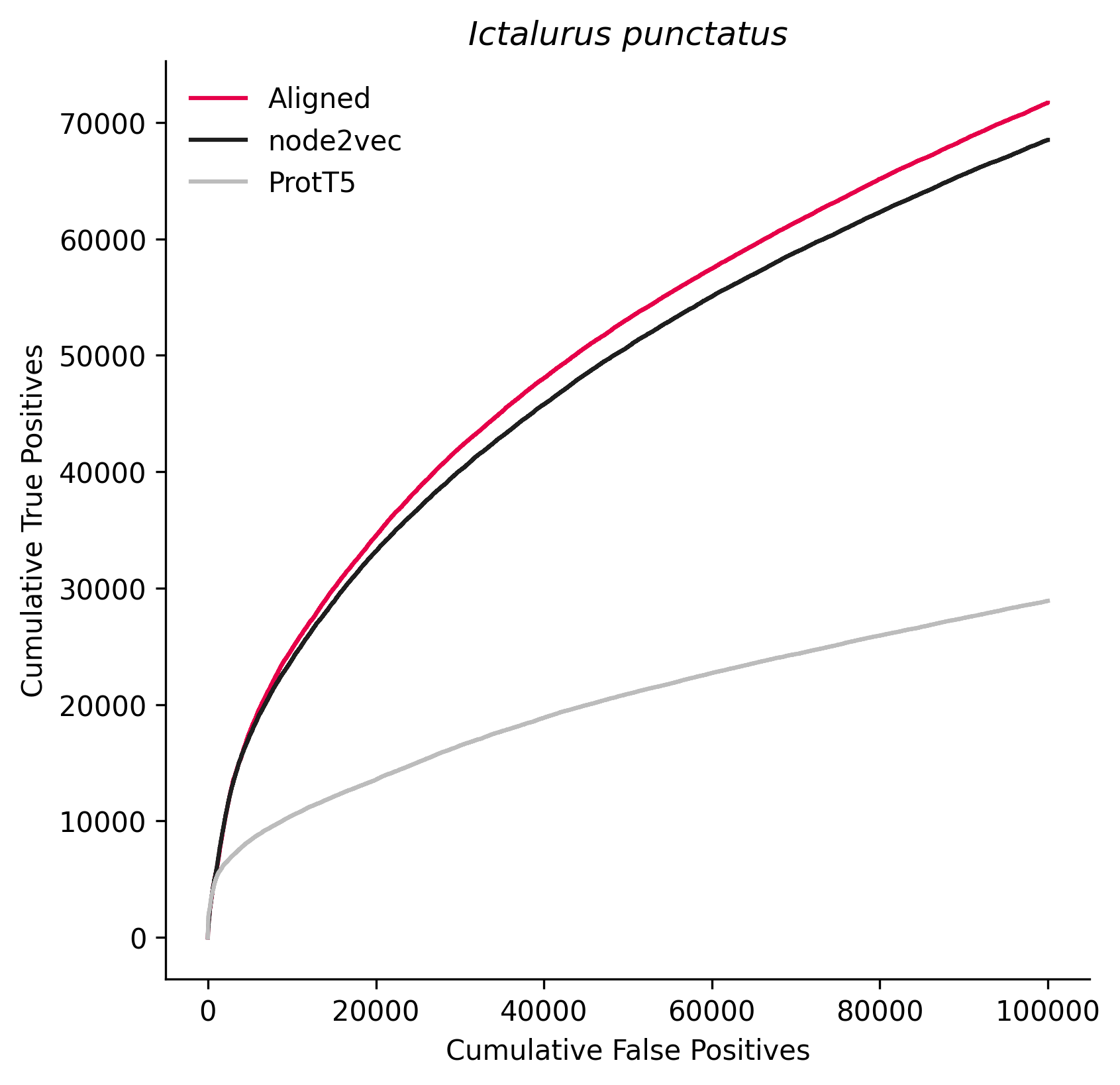

### 8010.png

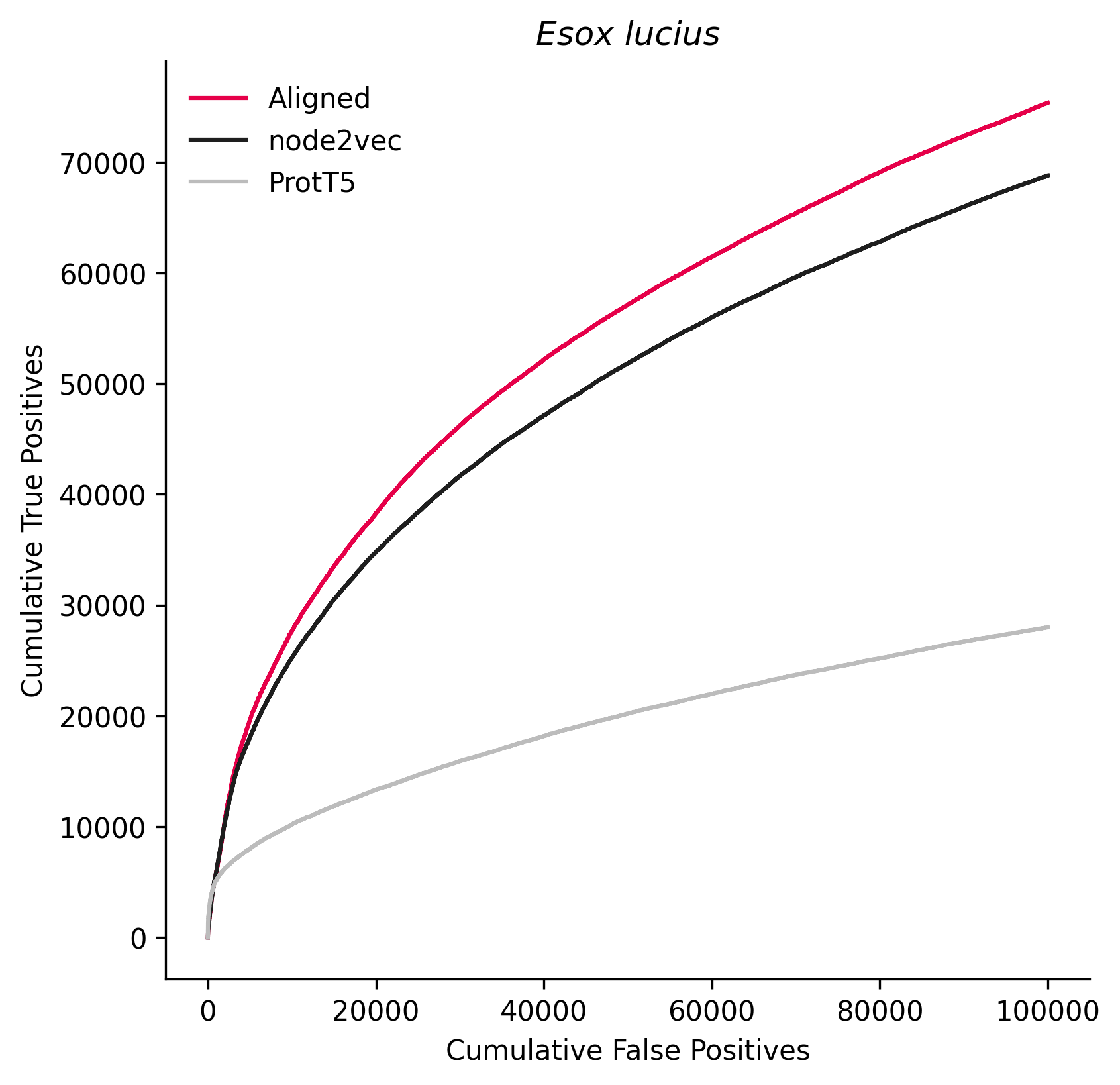

### 8355.png

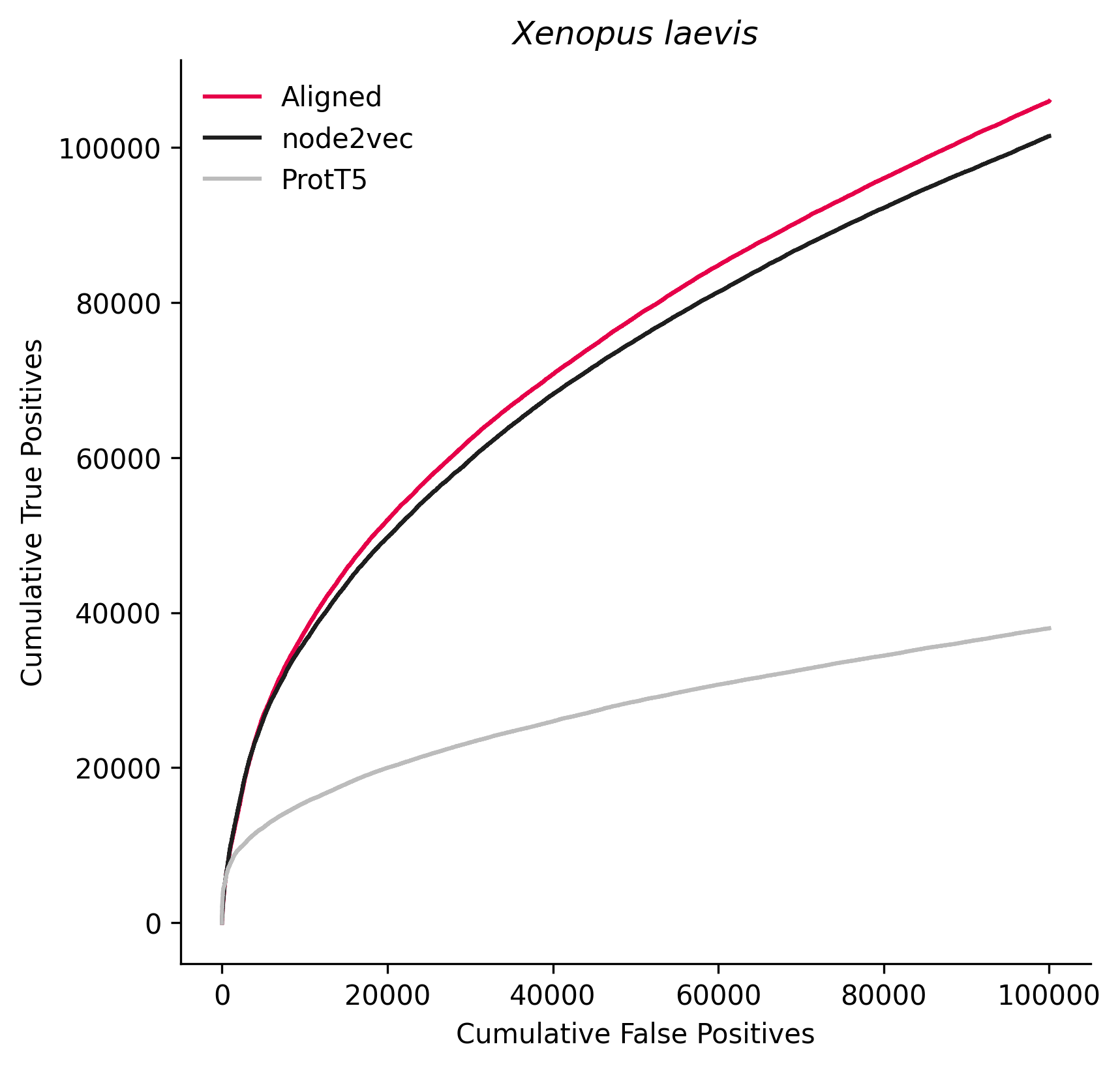

### 8469.png

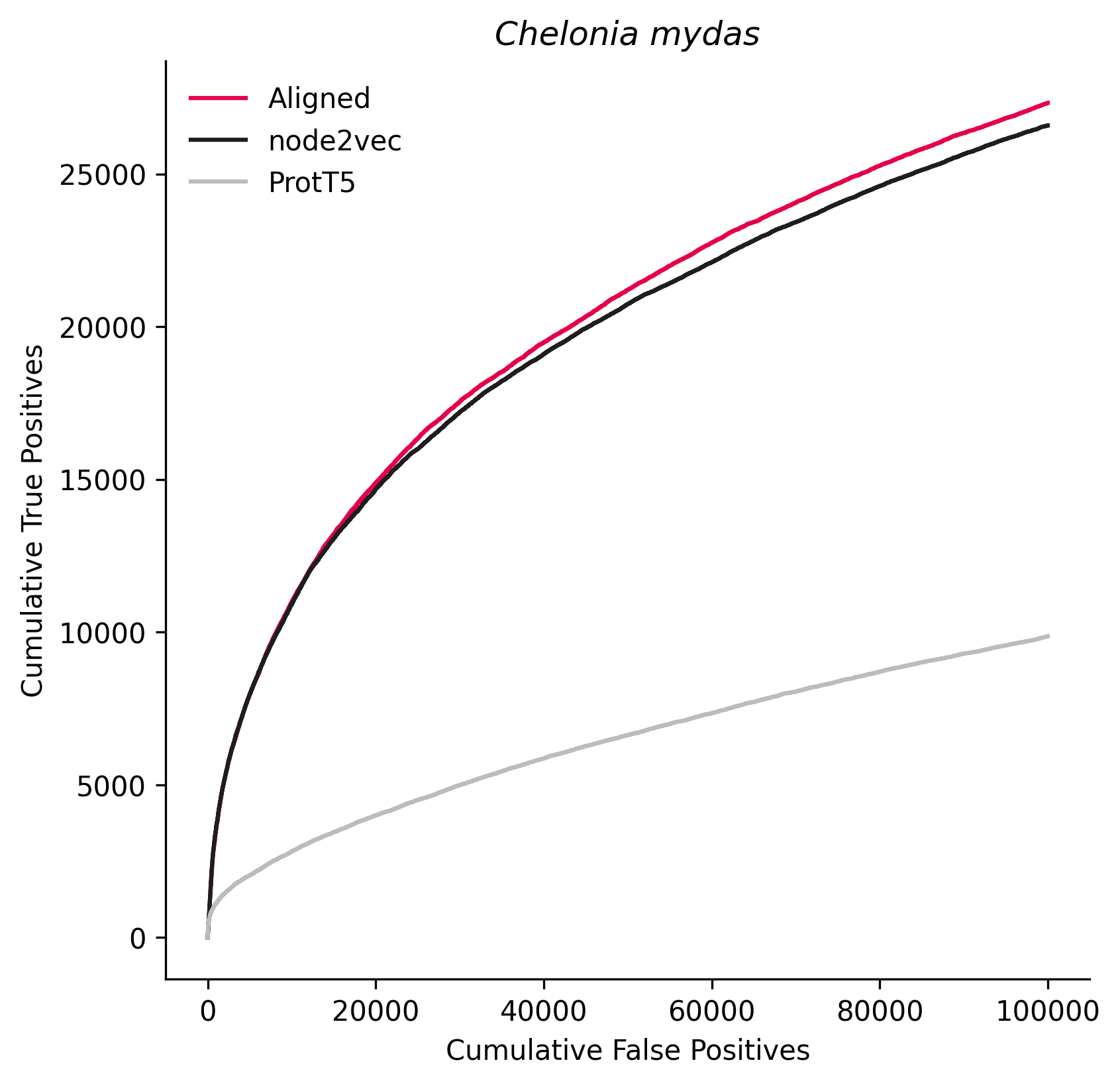

### 8840.png

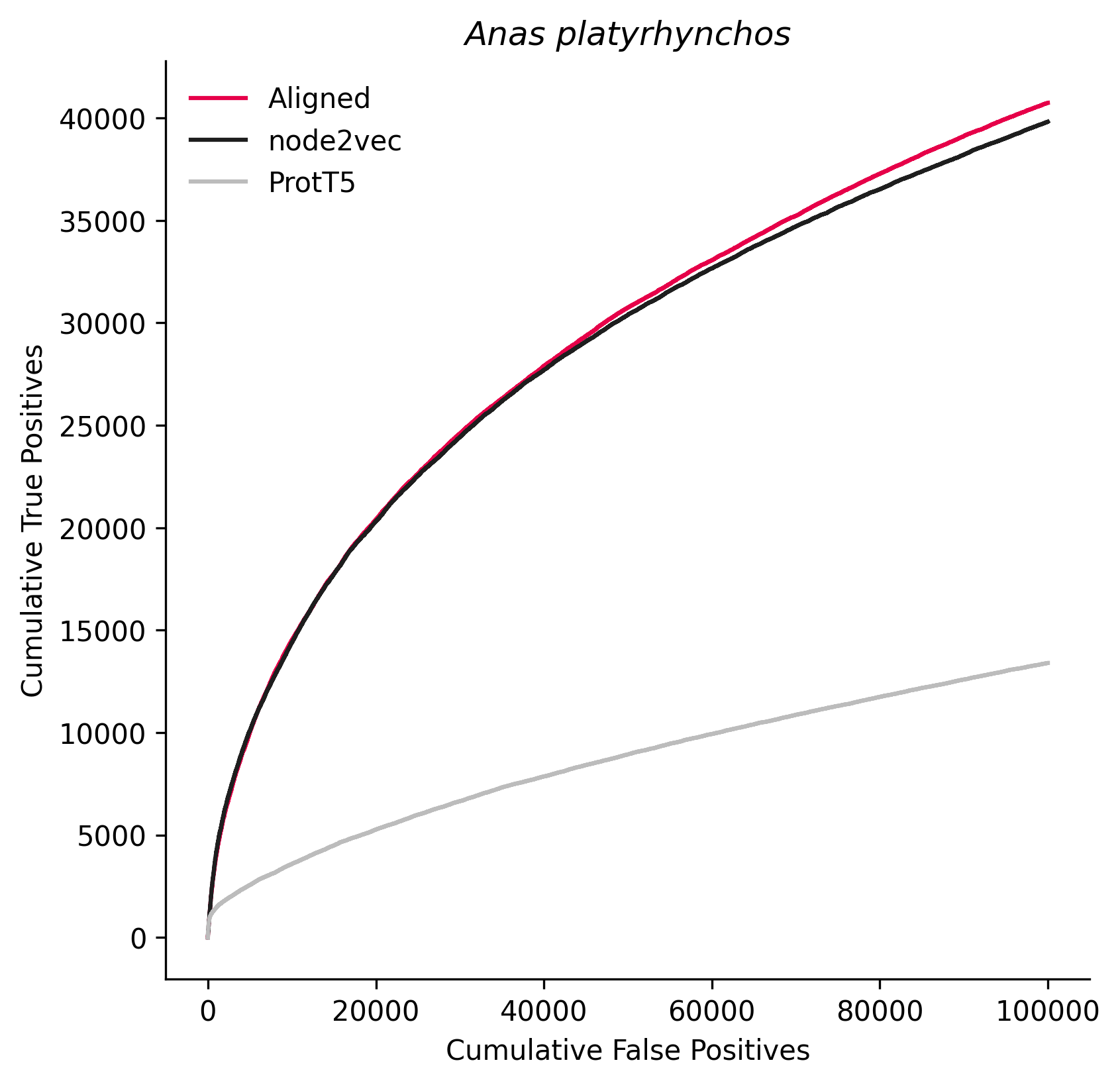

### 8845.png

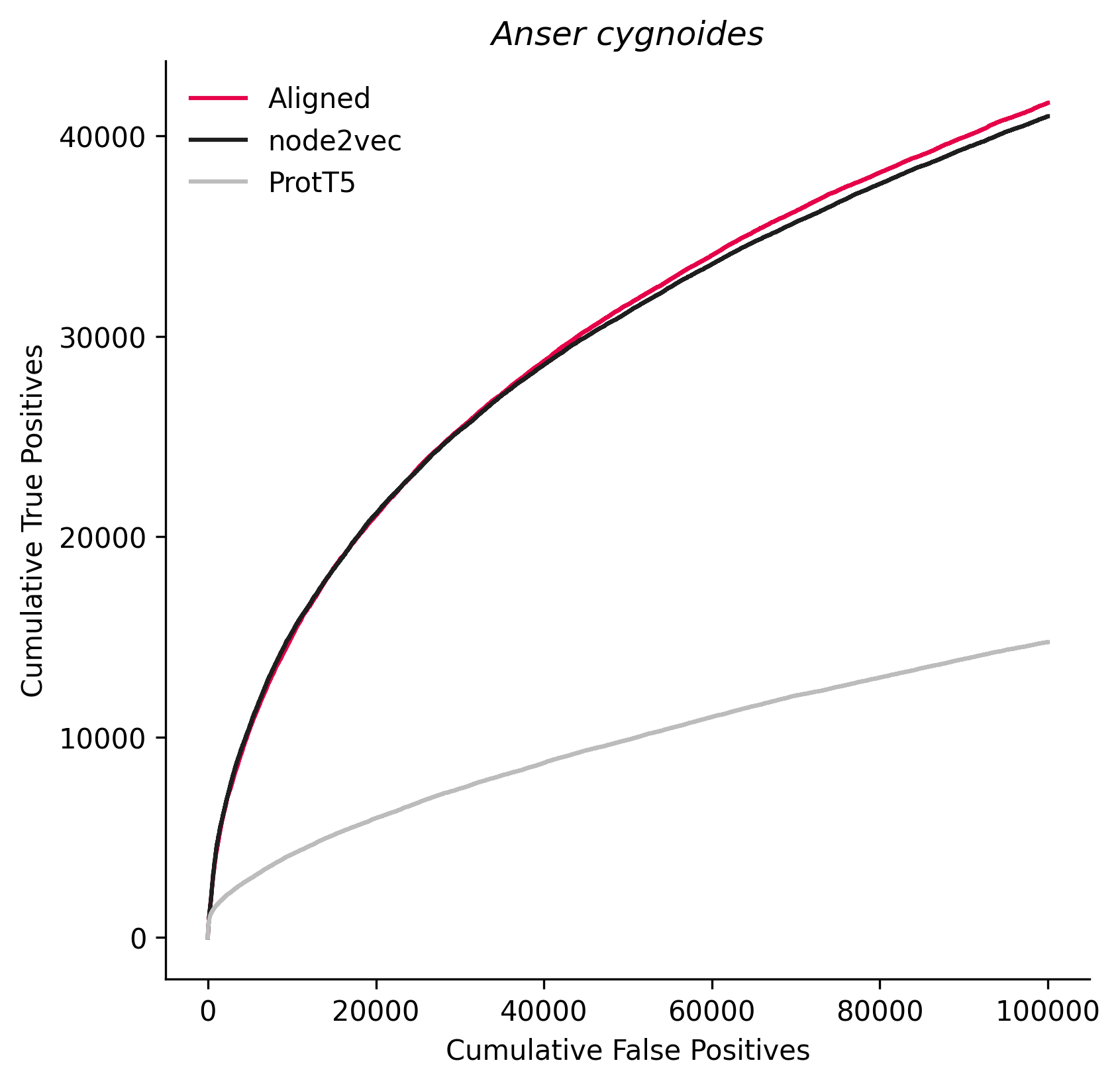

### 9593.png

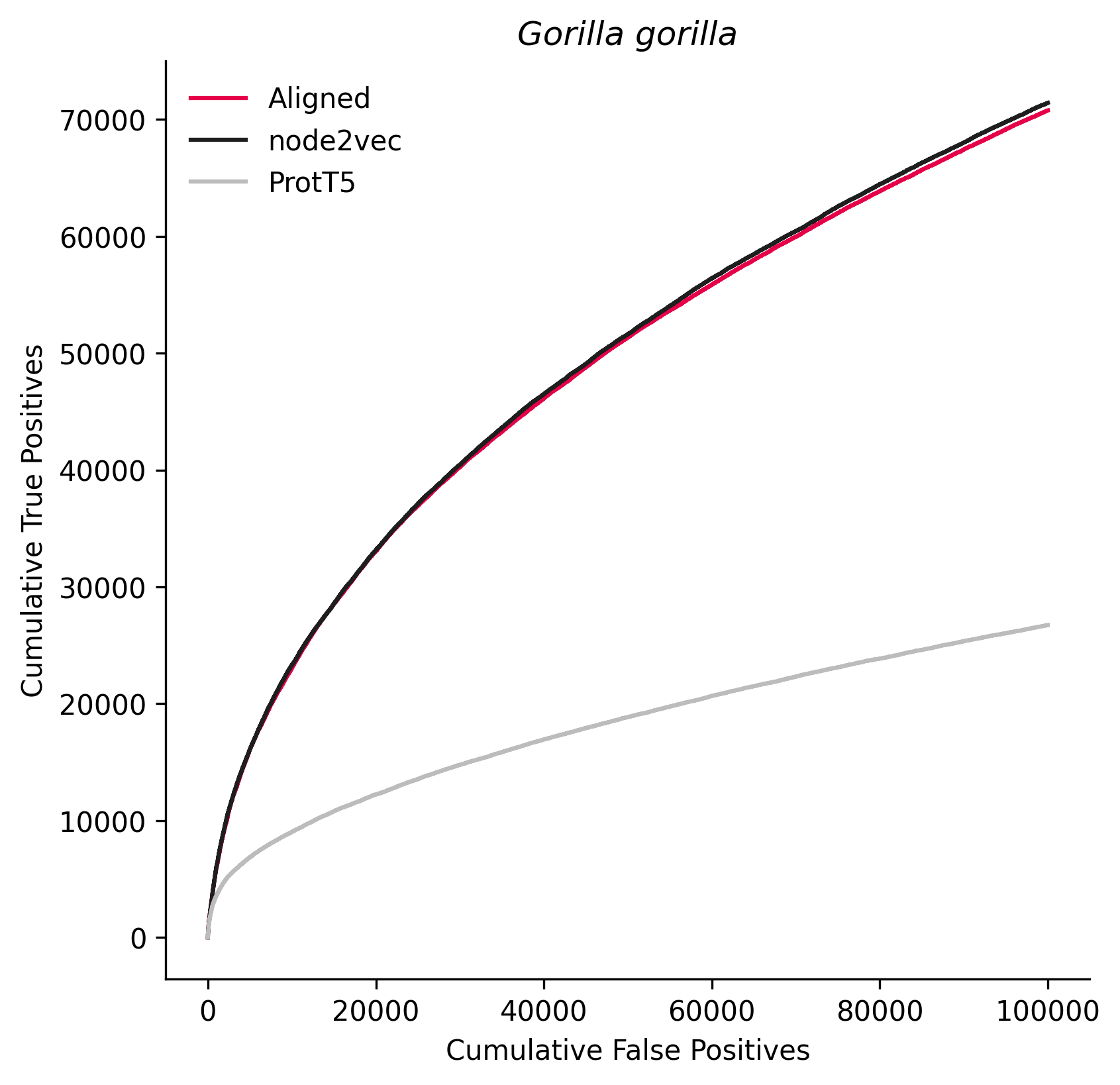

### 9615.png

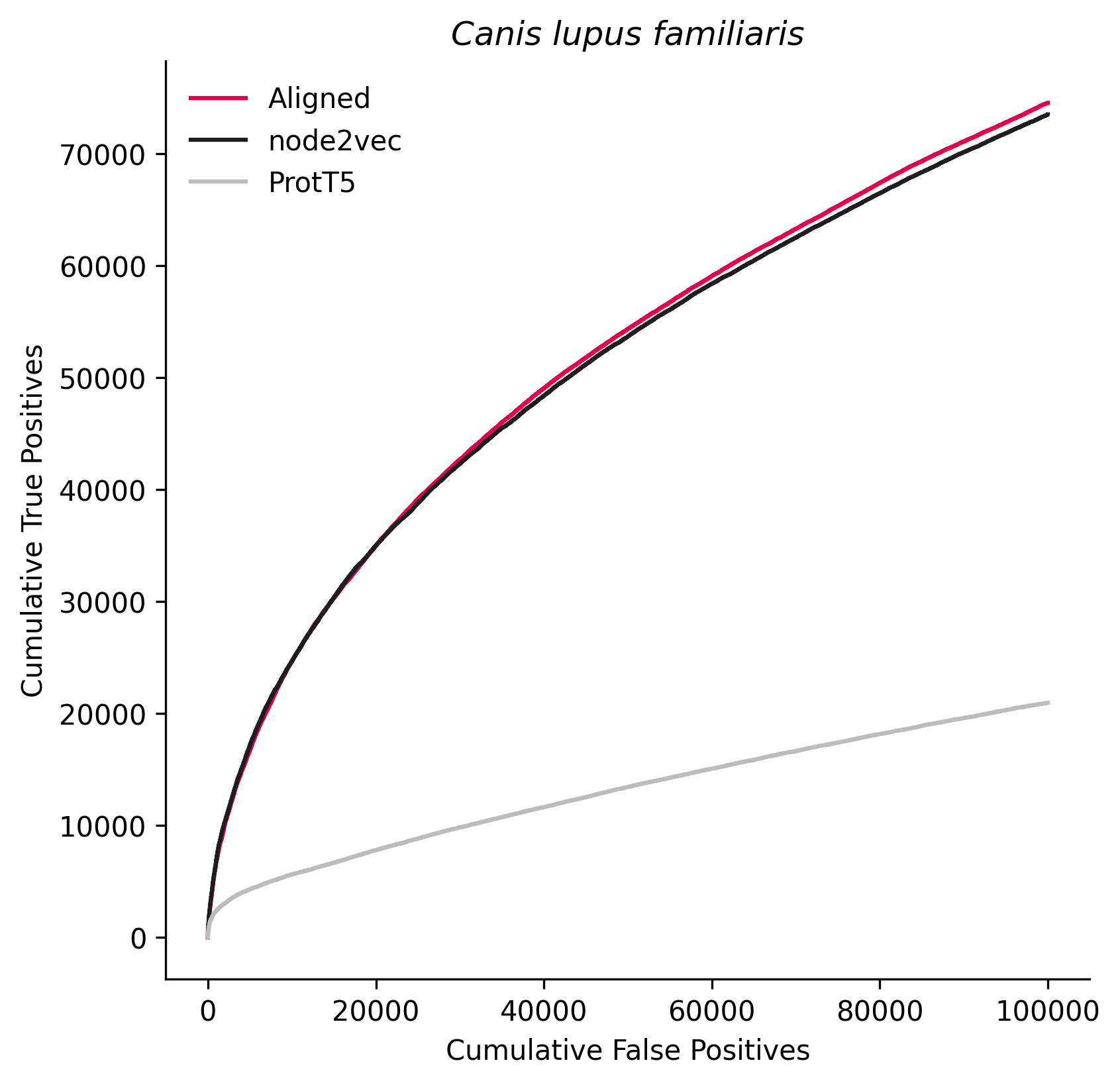

### 9685.png

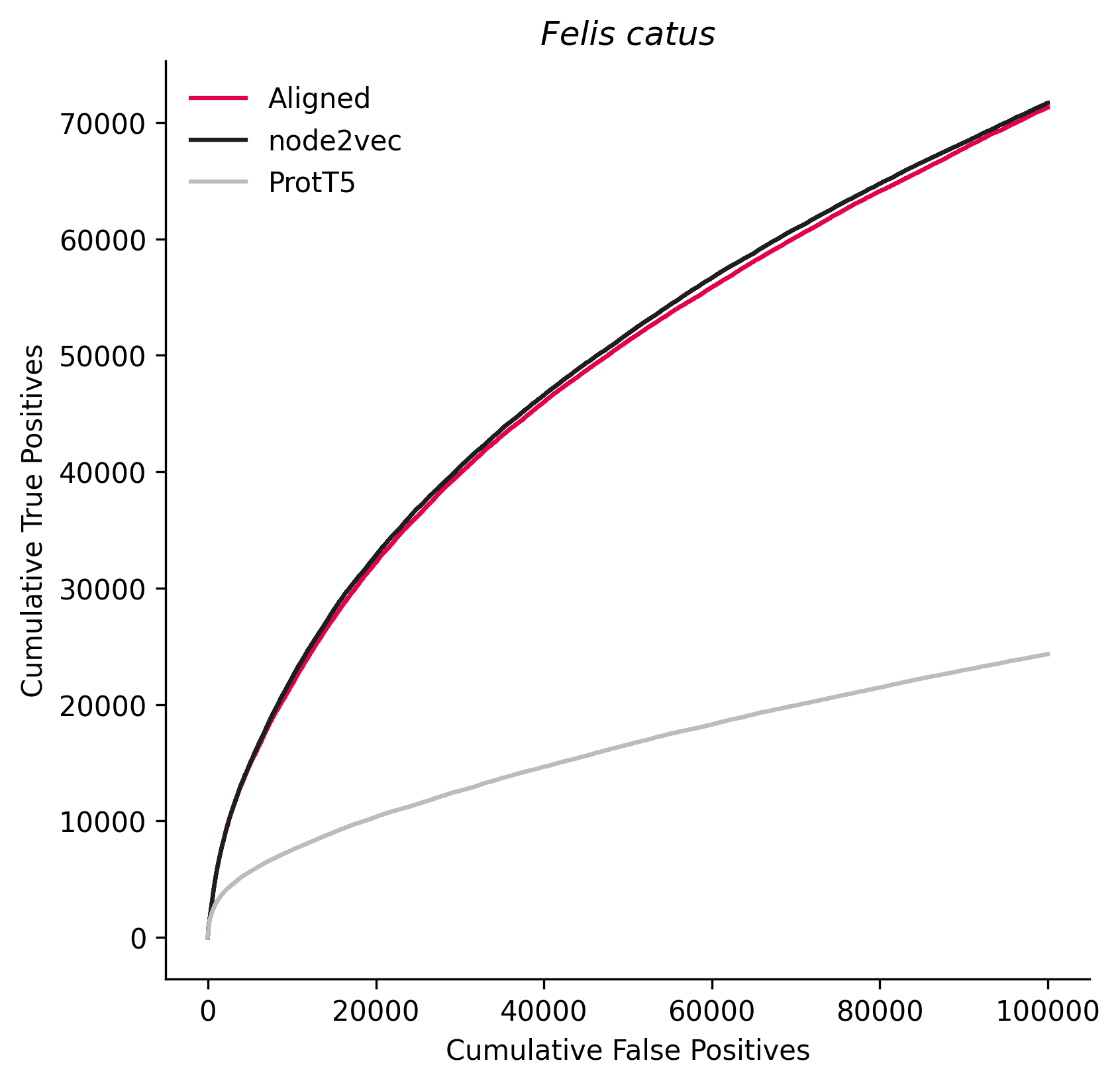

### 9691.png

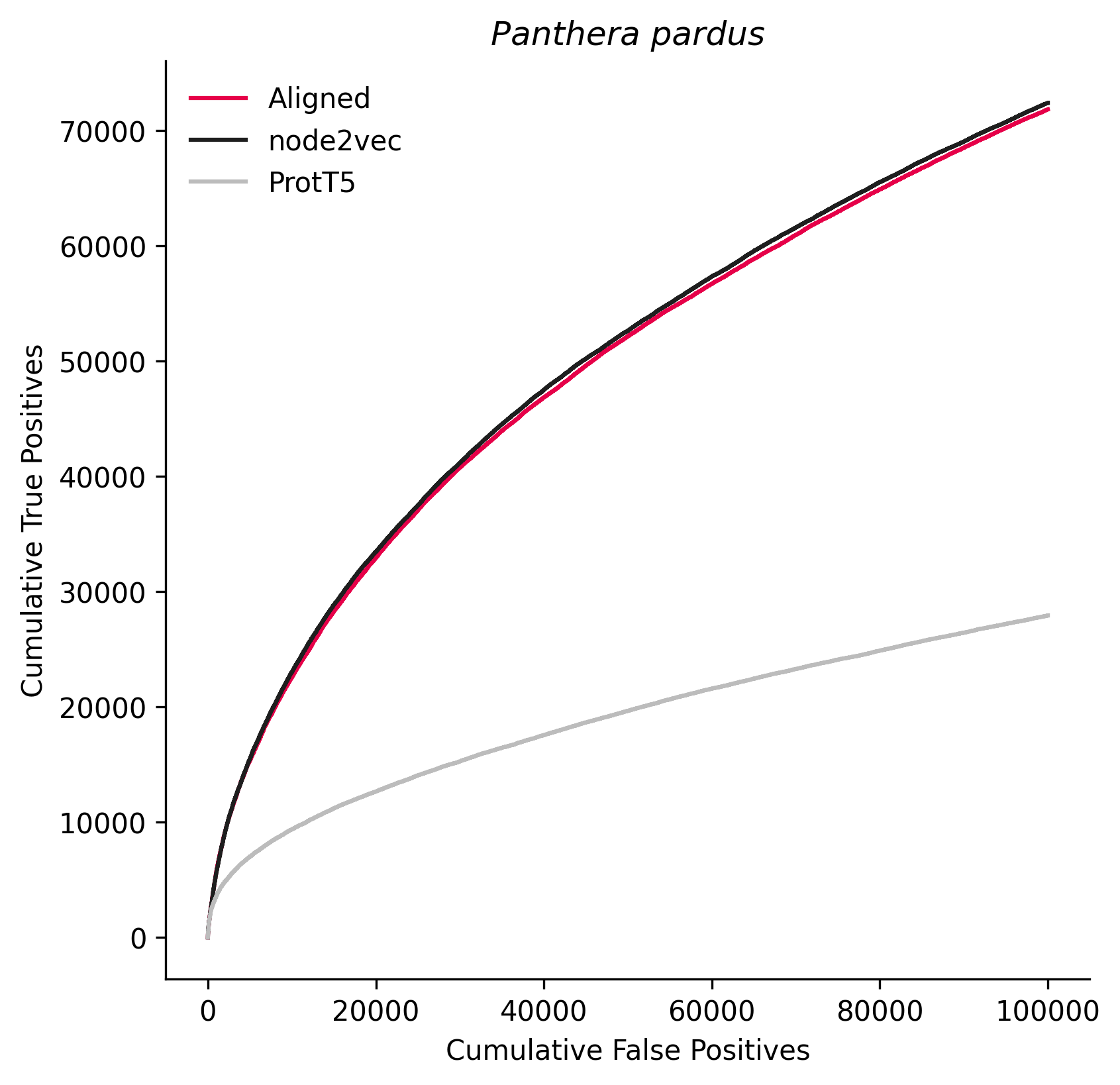

### 9708.png

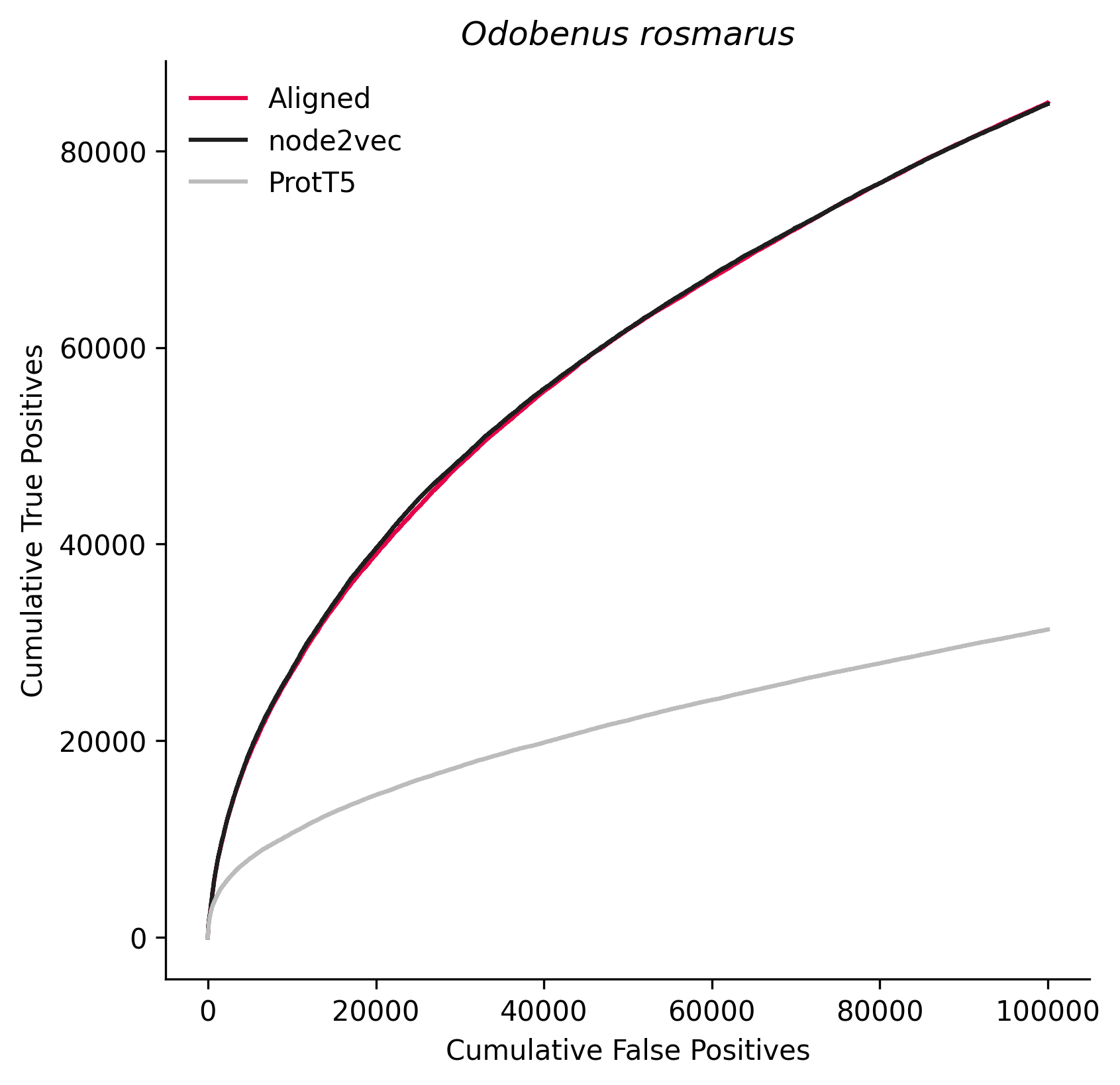

### 9785.png

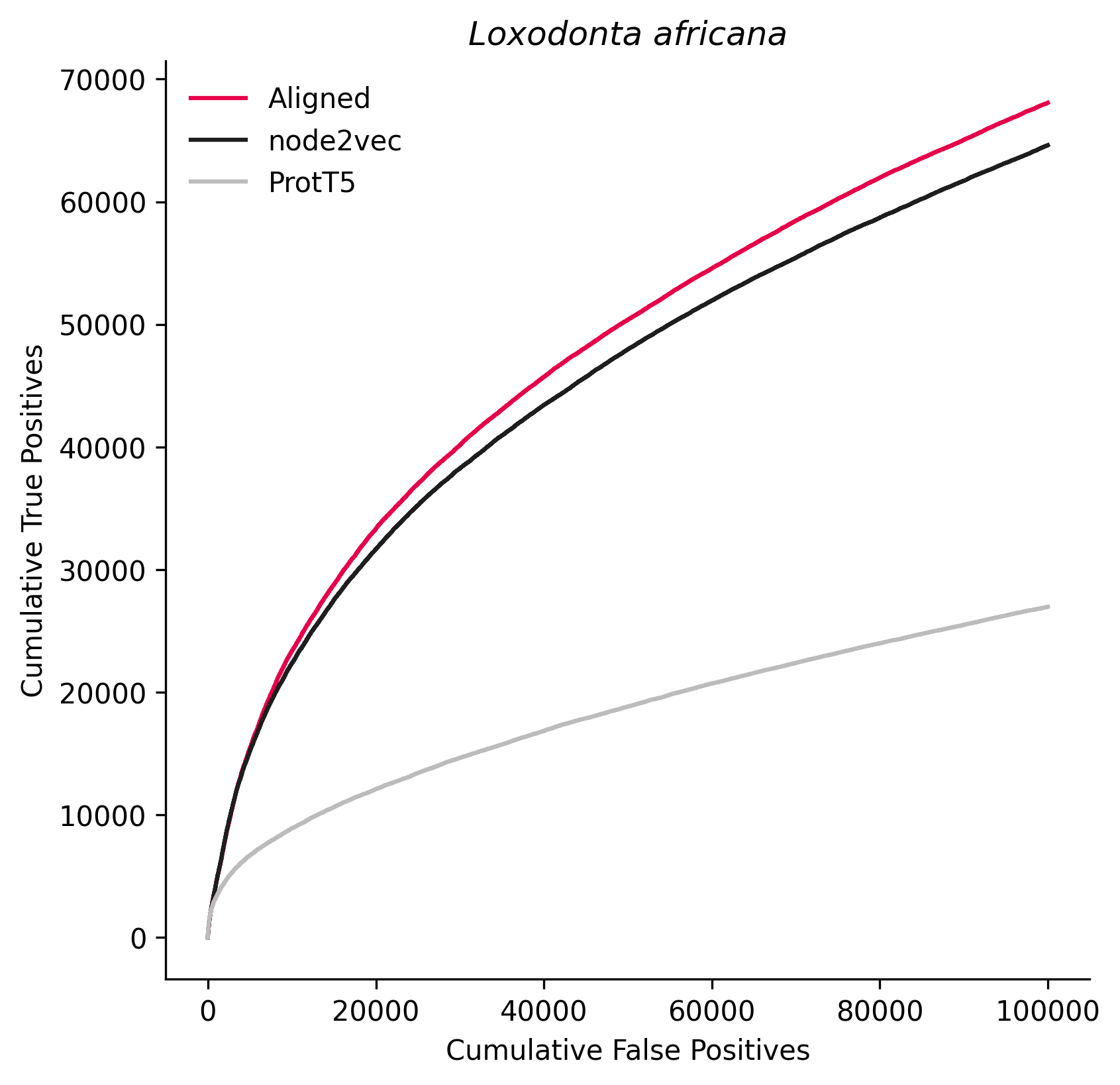

### 9925.png

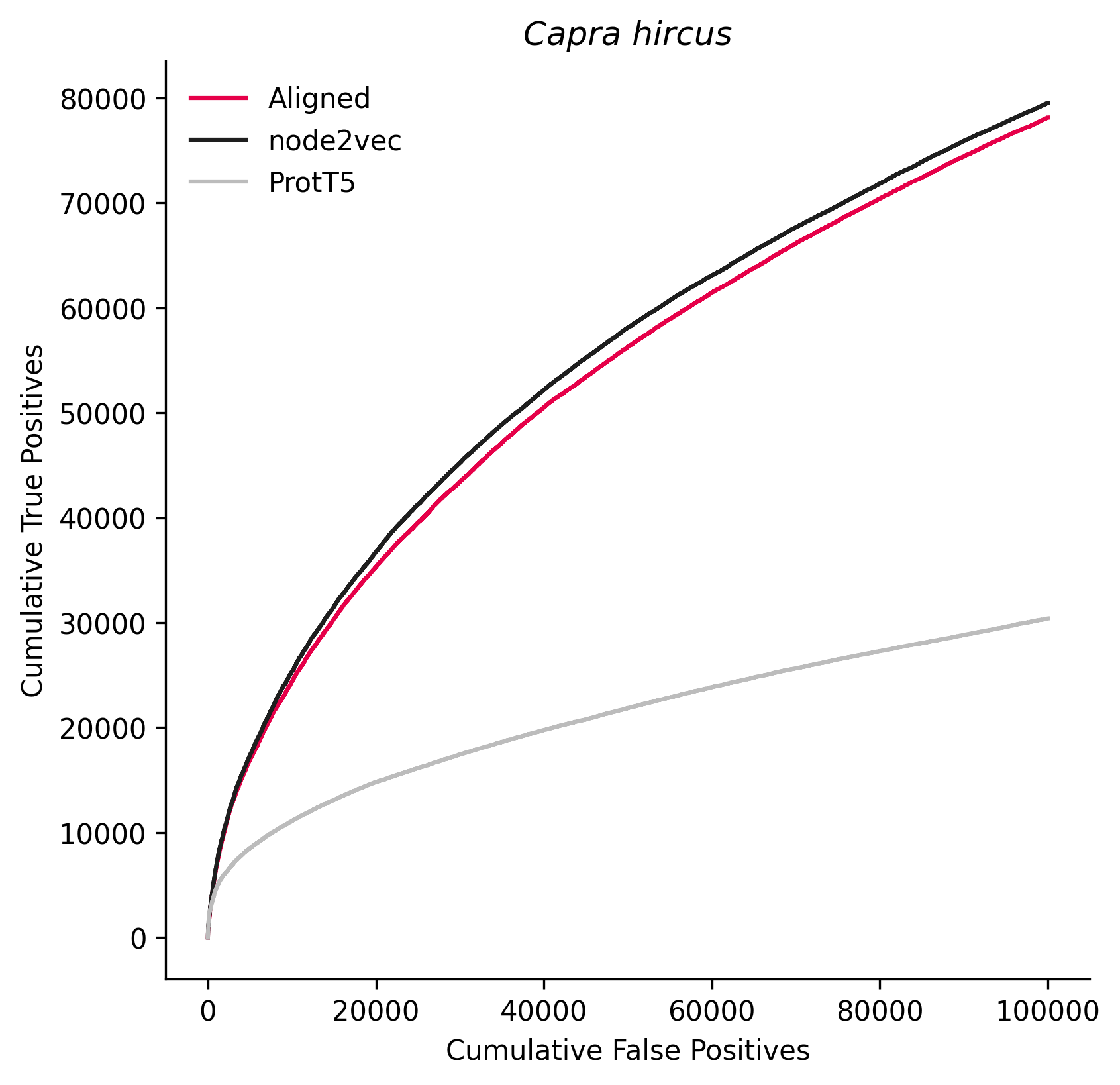

### 12957.png

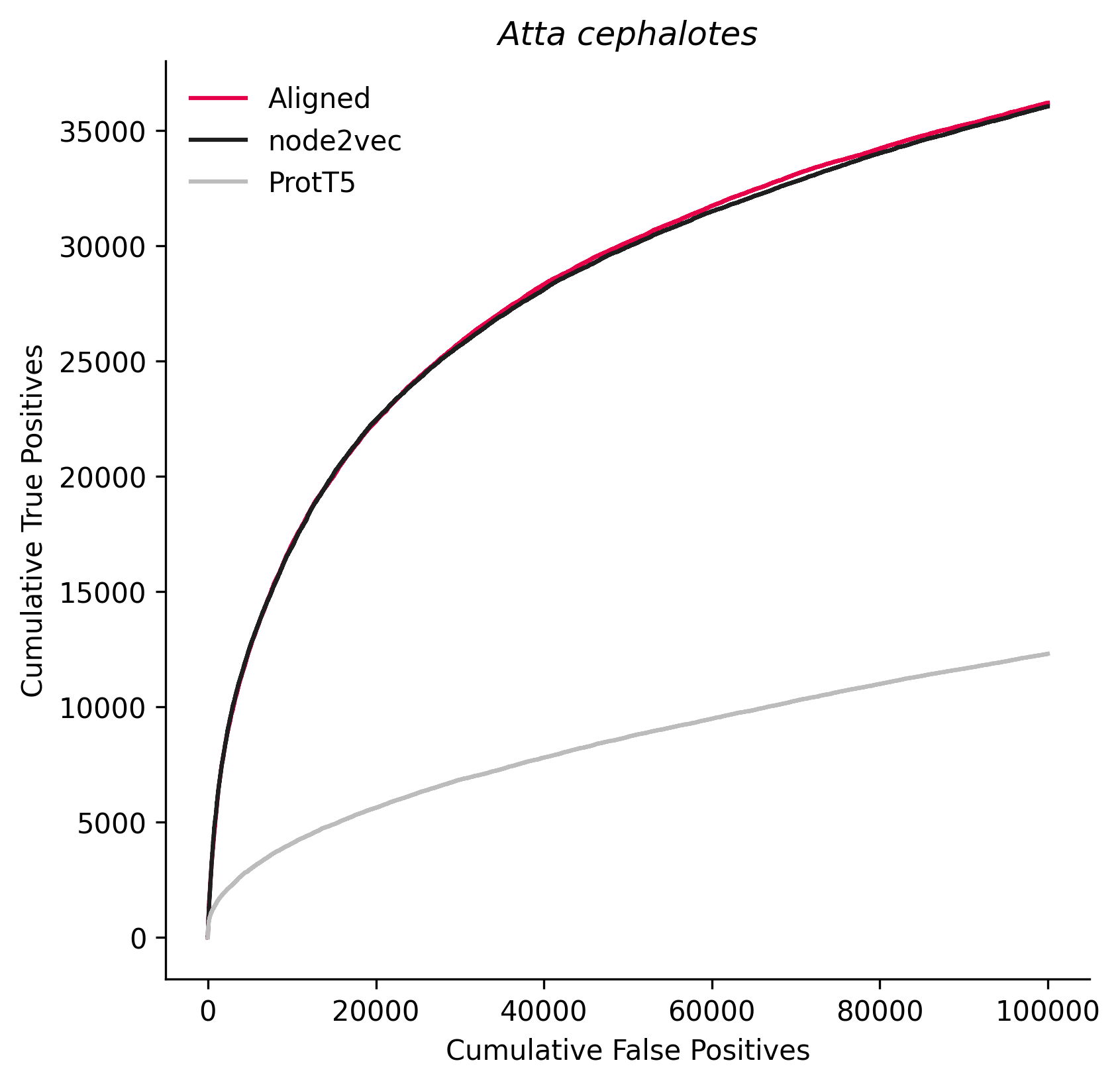

### 13735.png

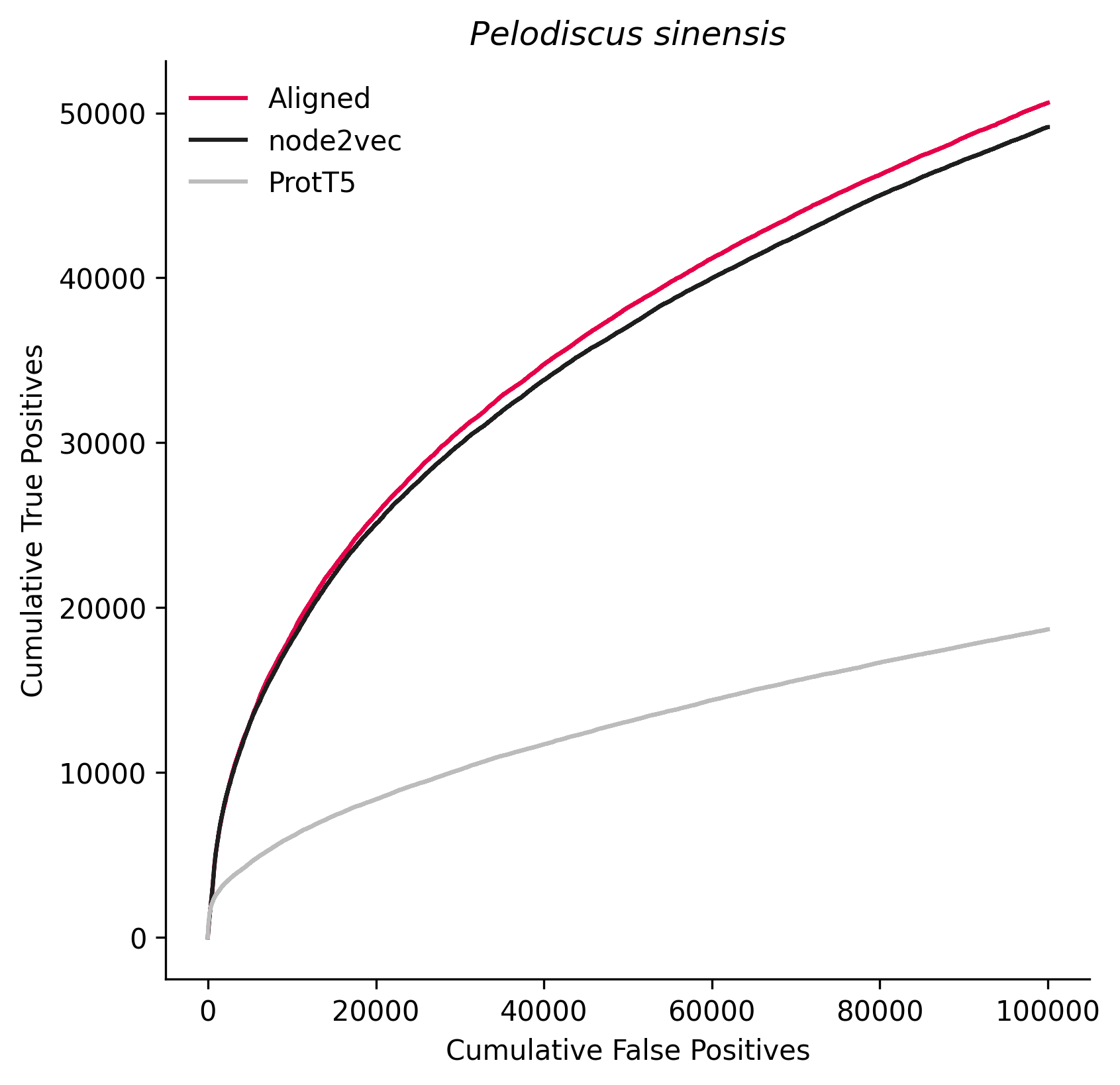

### 15368.png

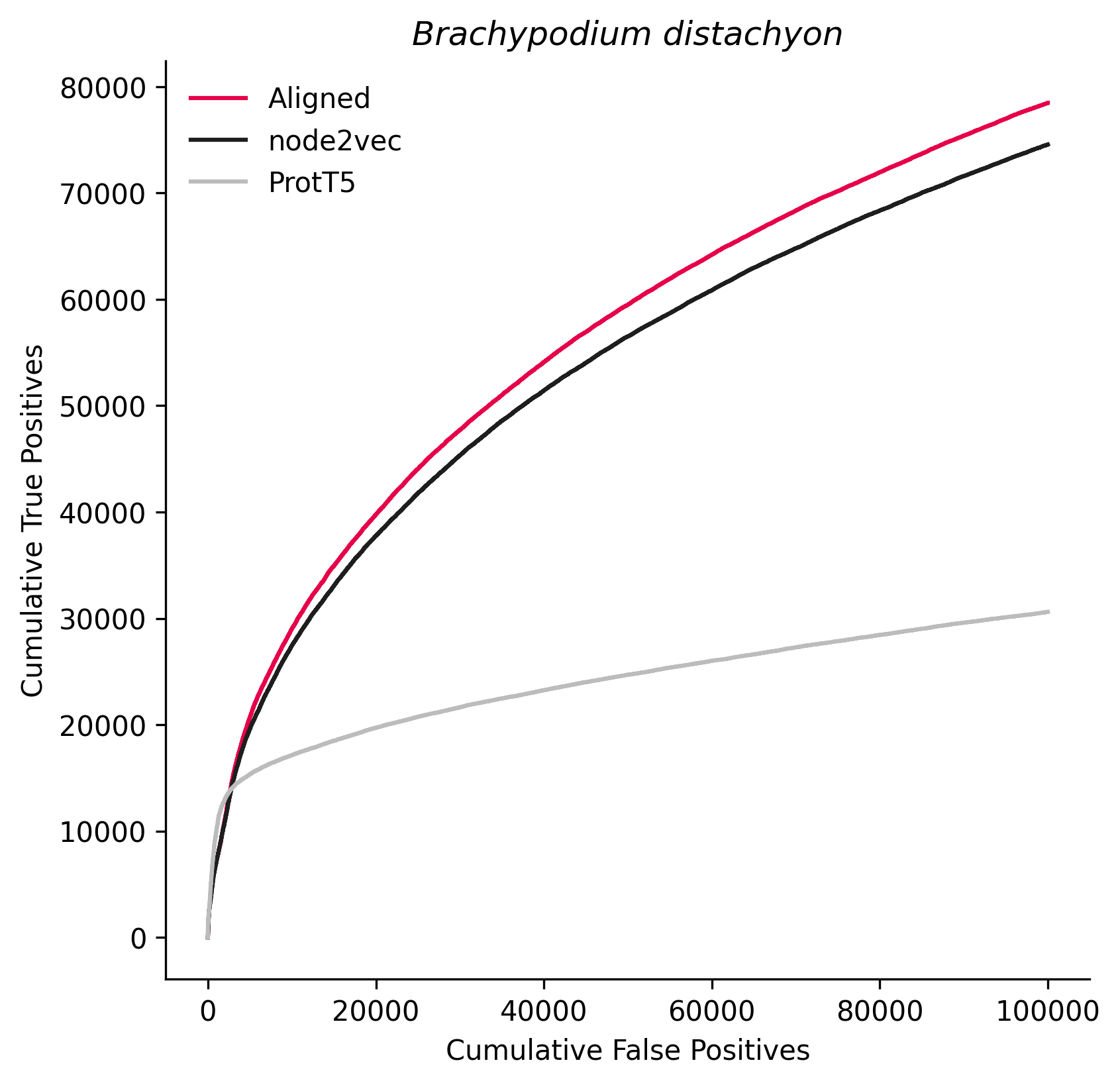

### 29073.png

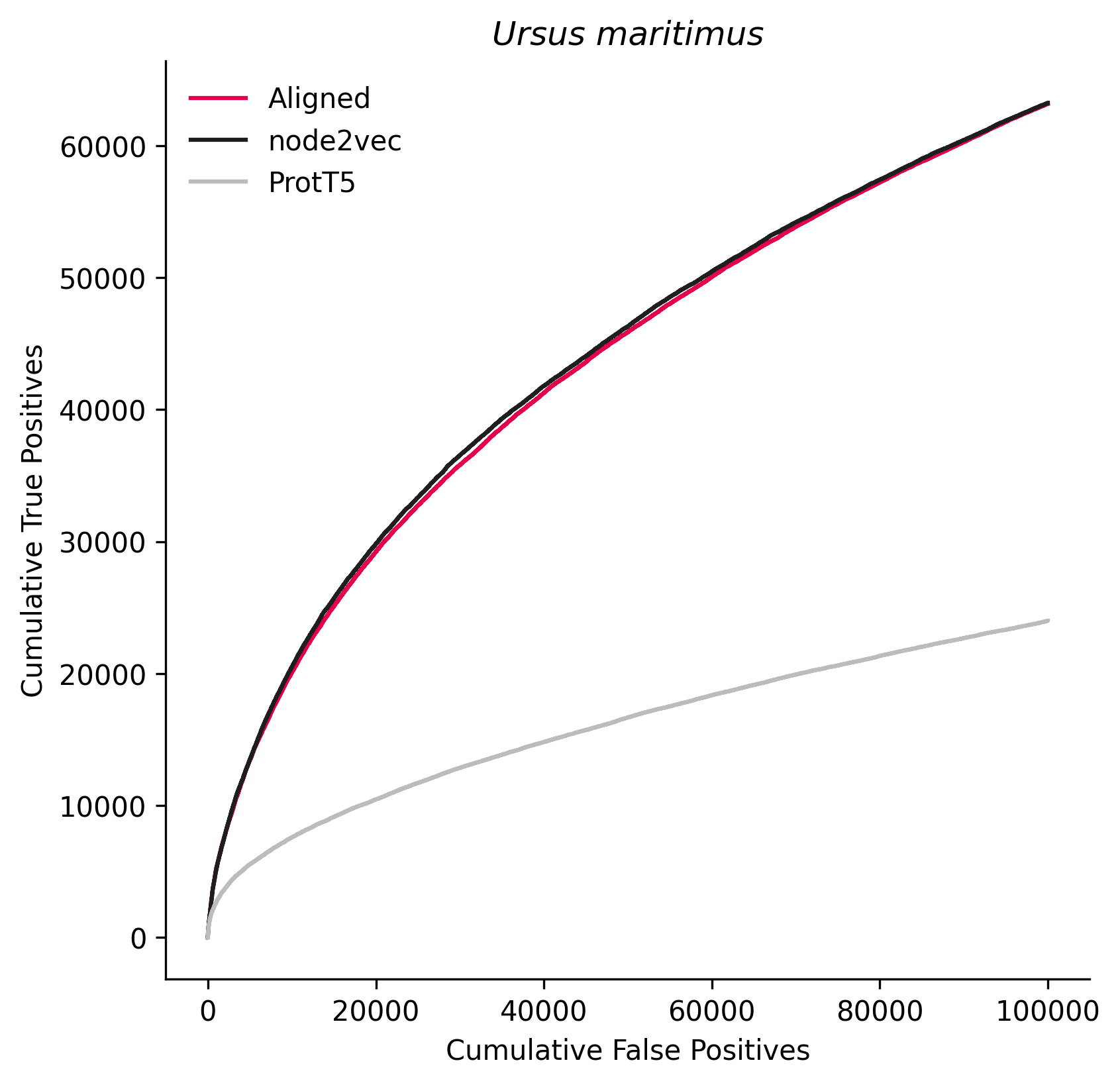

### 29730.png

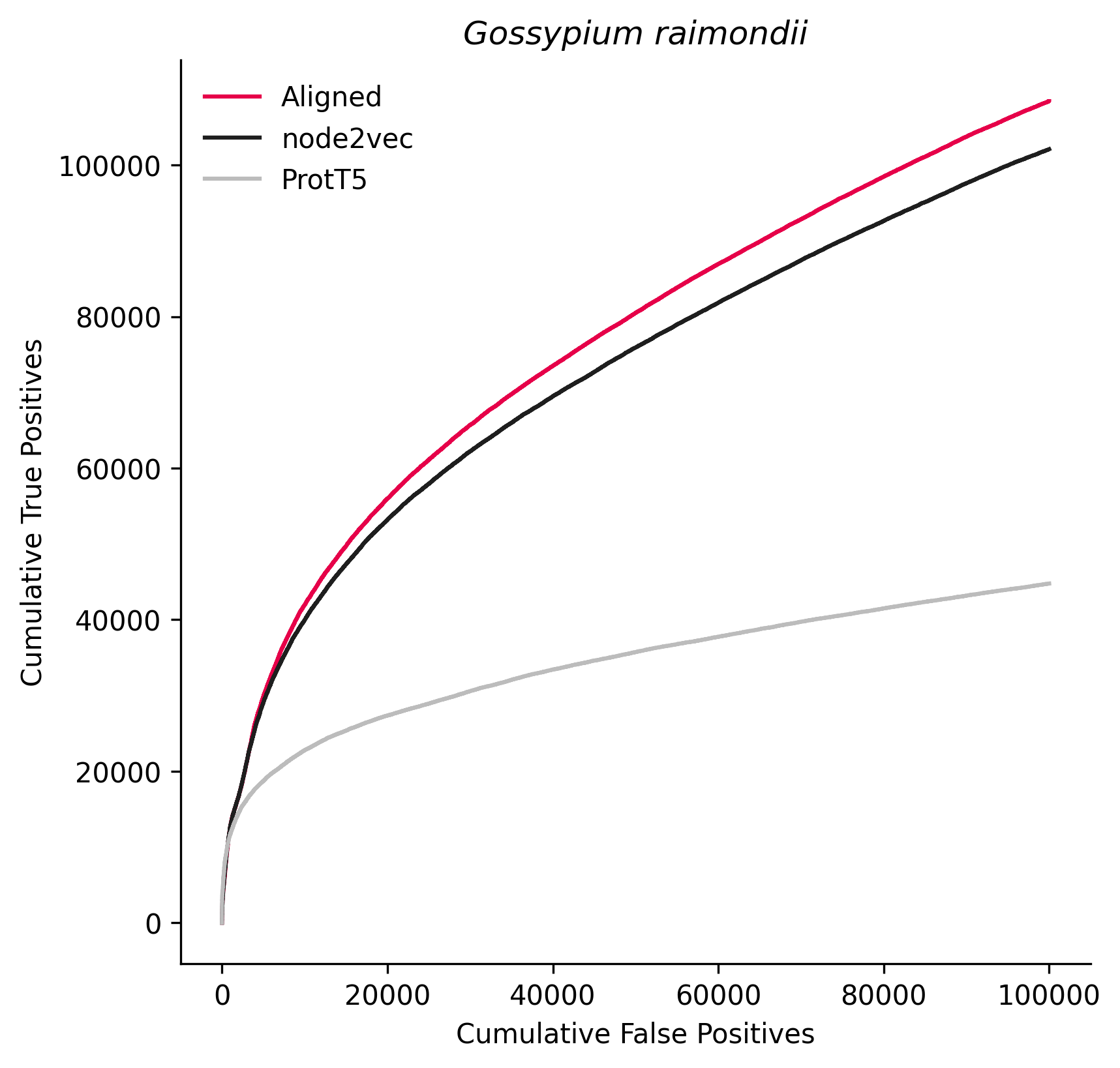

### 29760.png

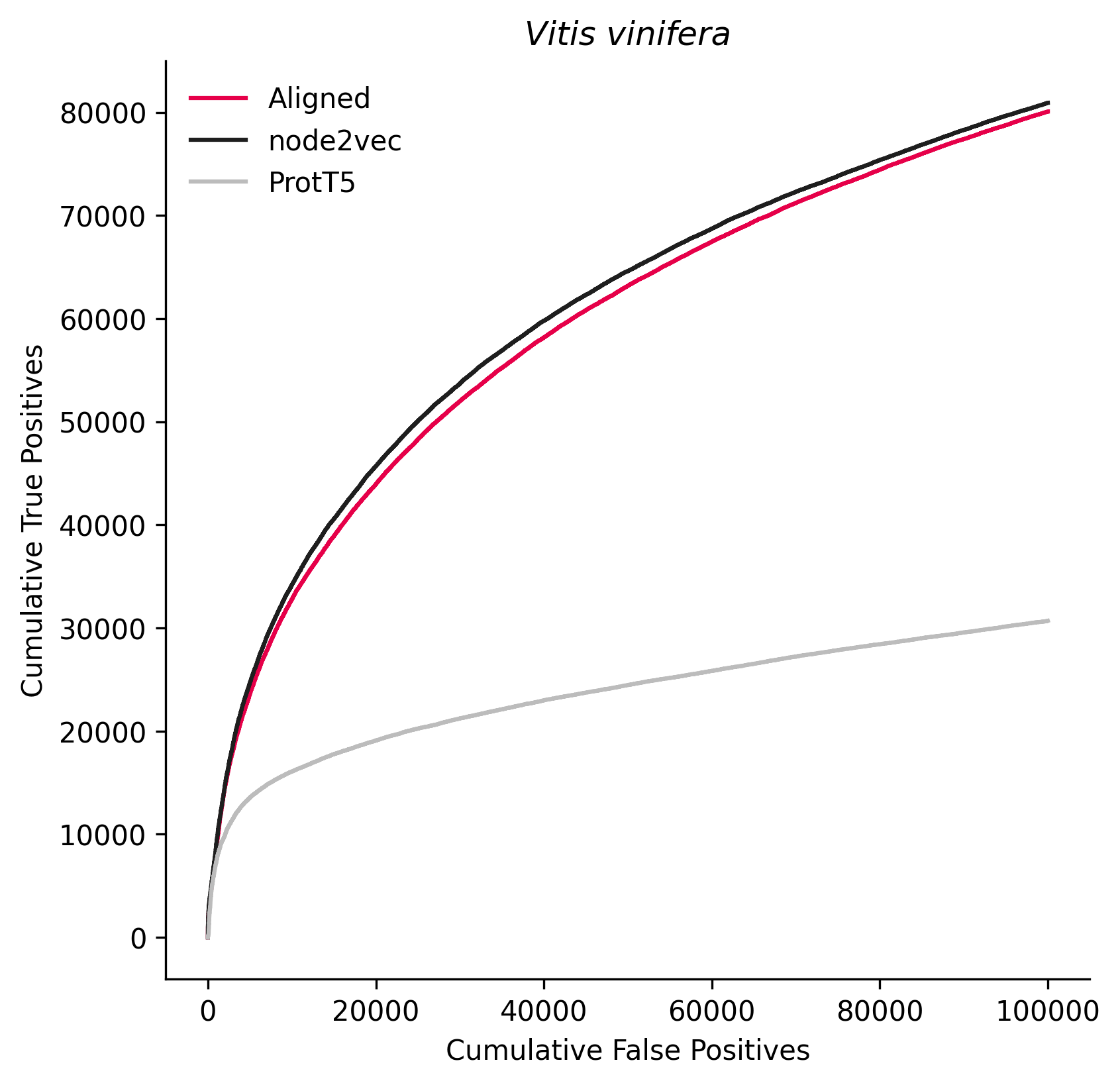

### 32473.png

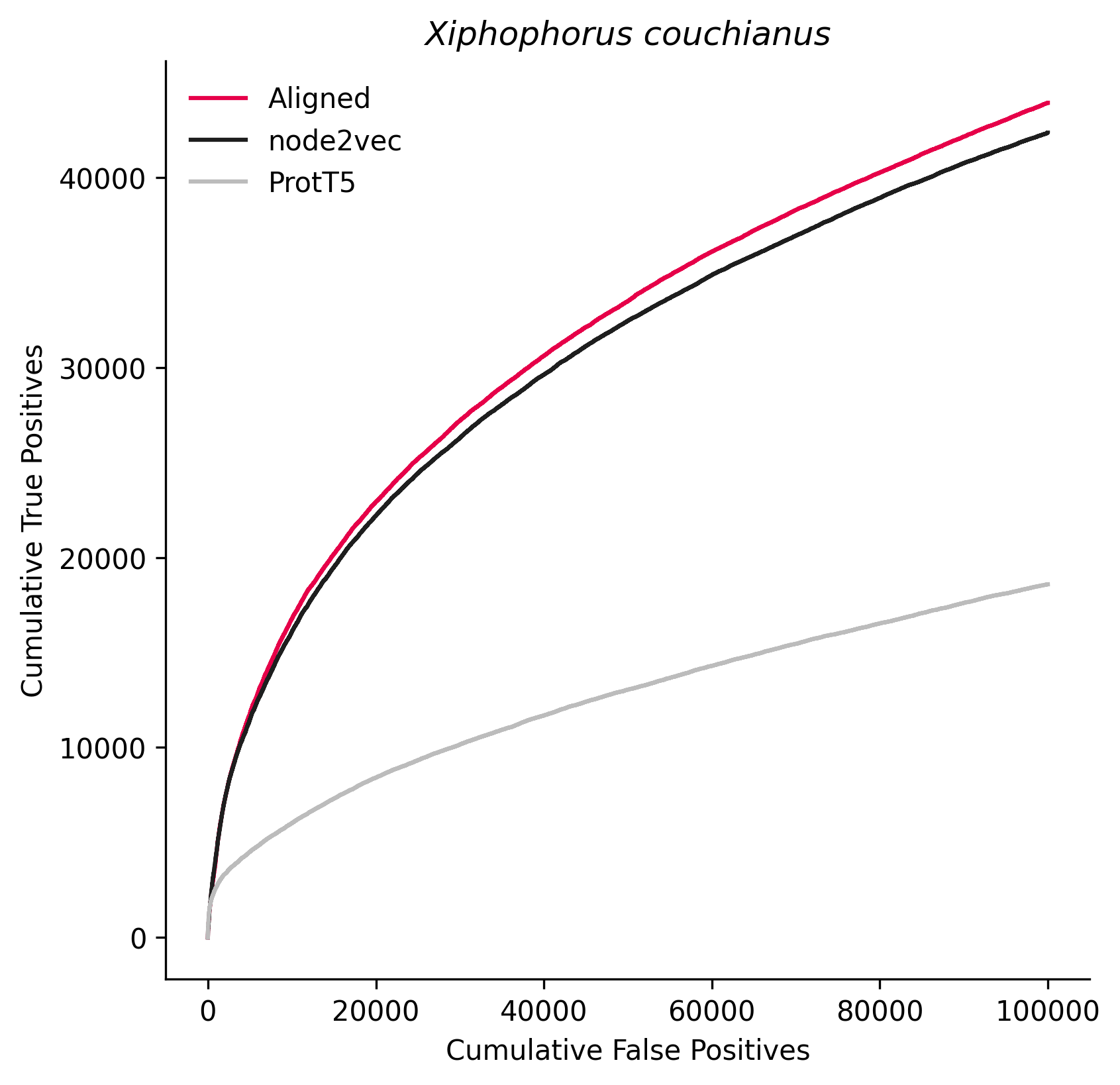

### 35128.png

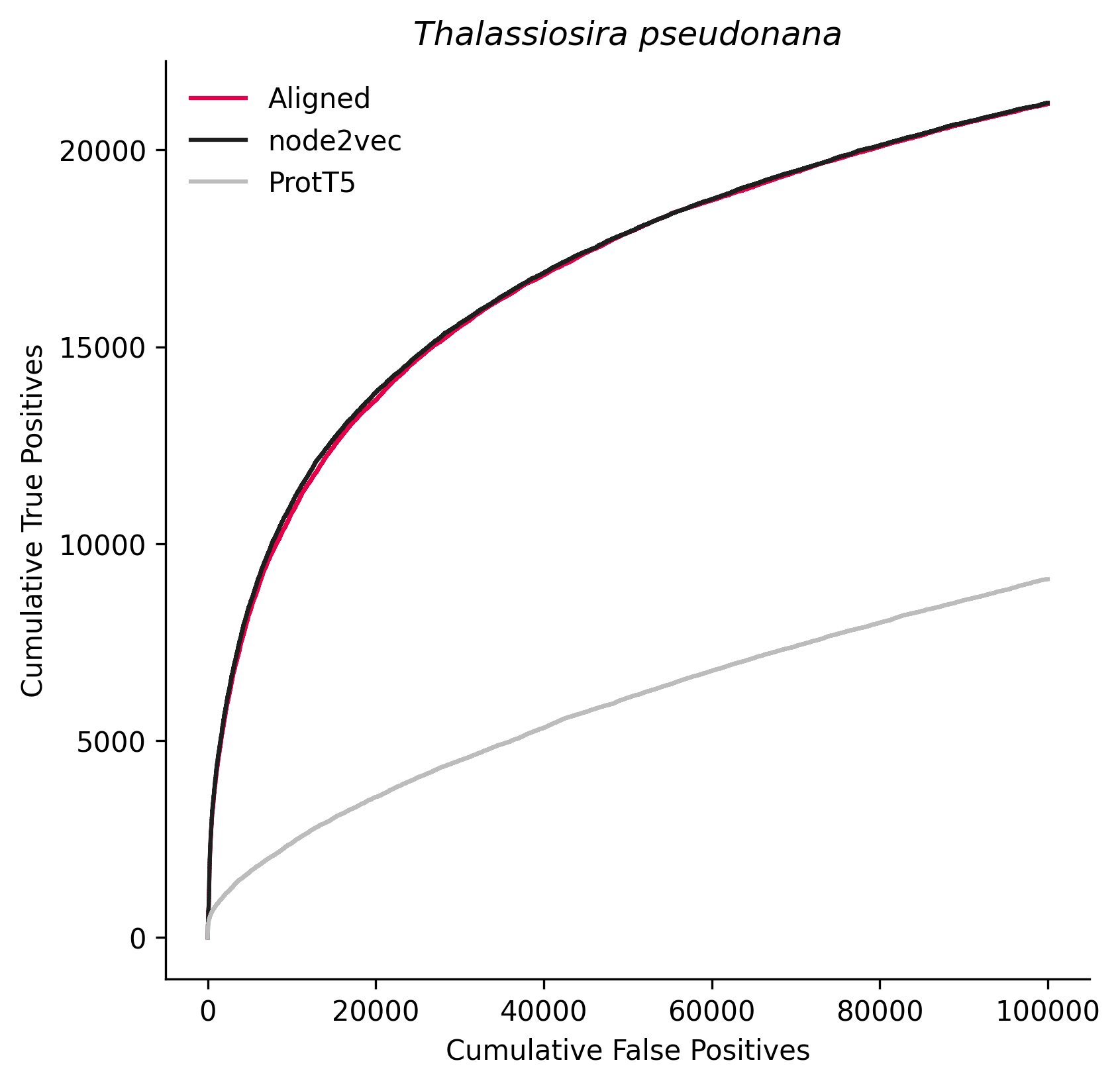

### 38654.png

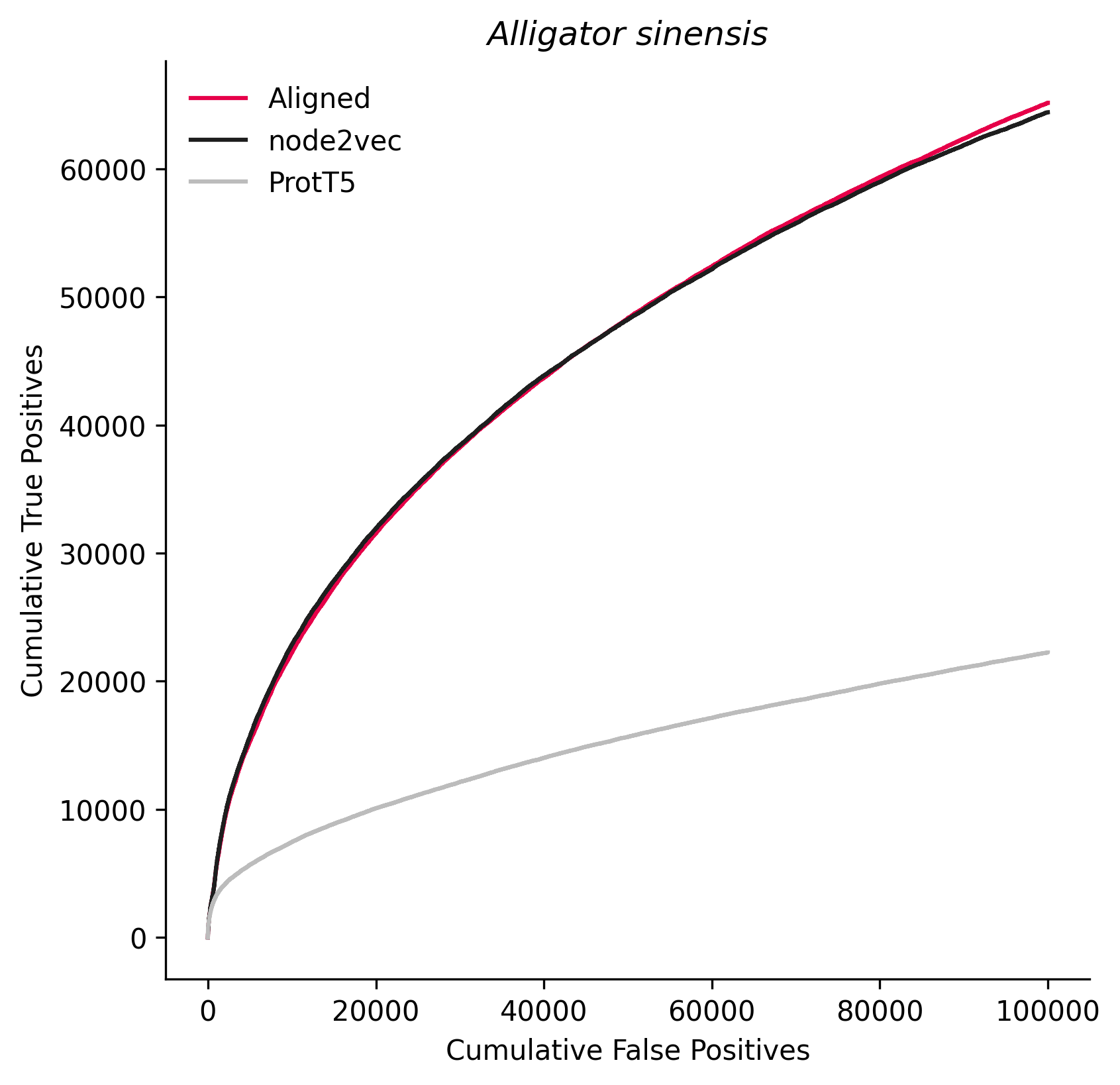

### 39432.png

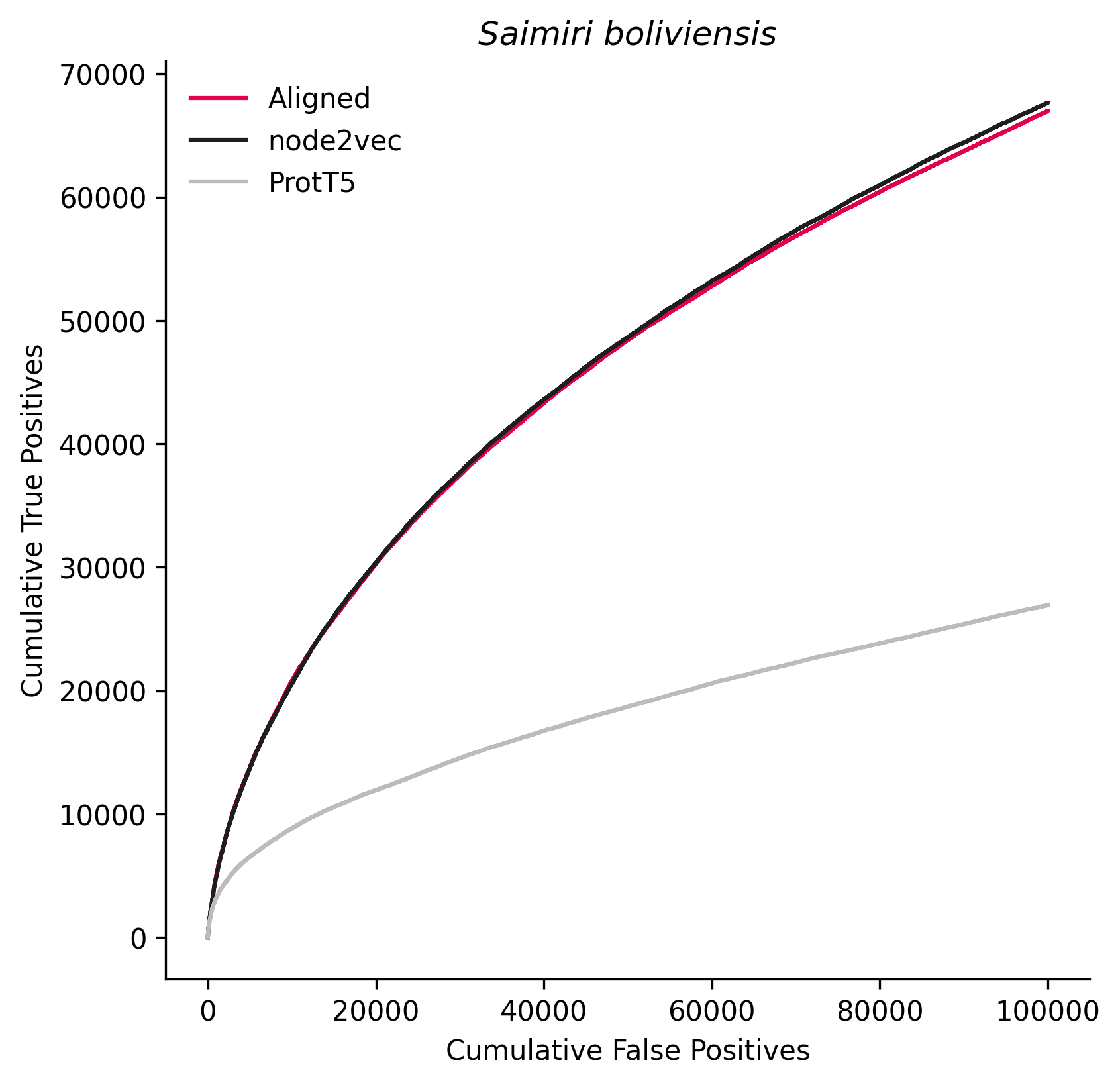

### 42345.png

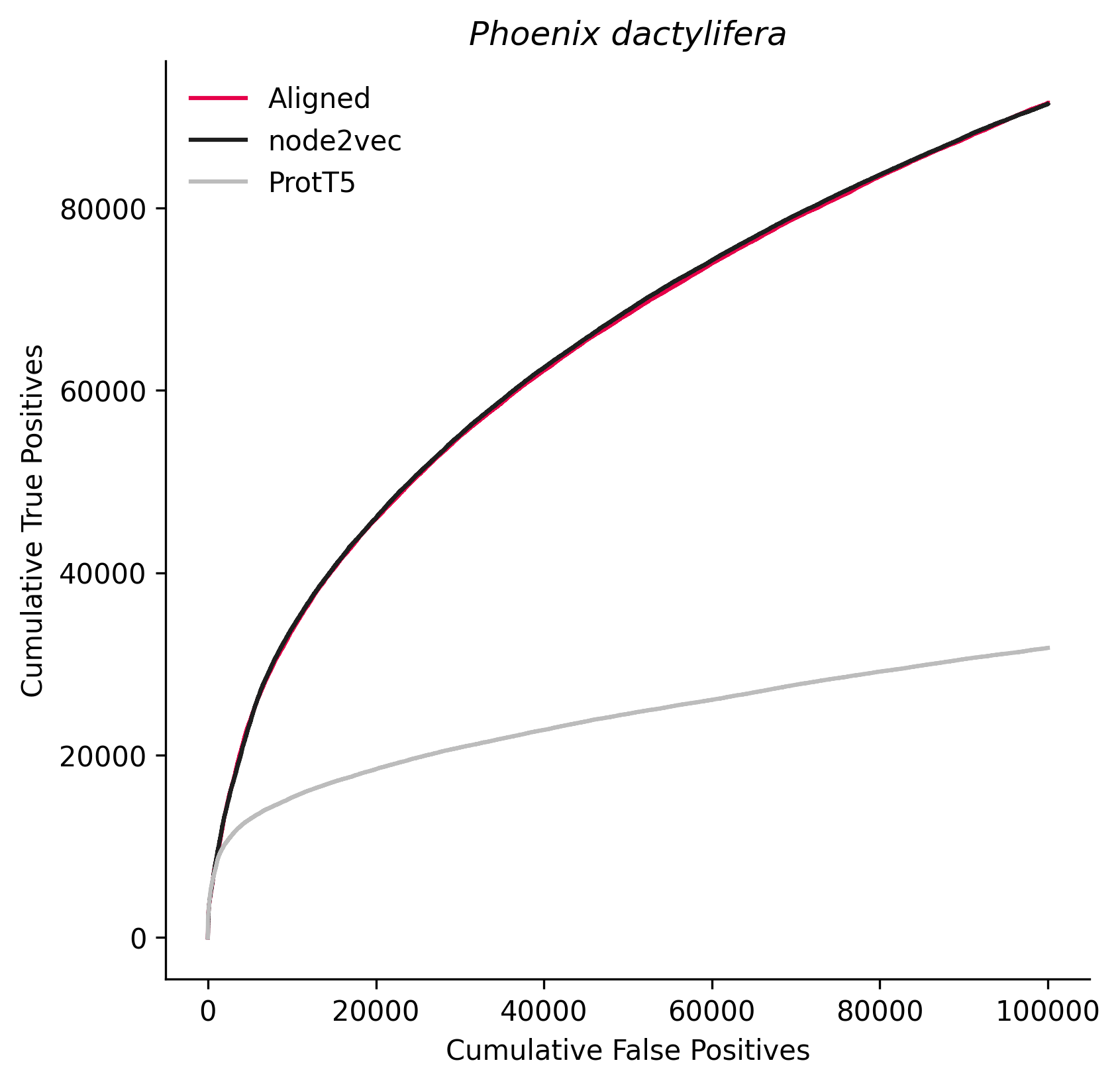

### 45351.png

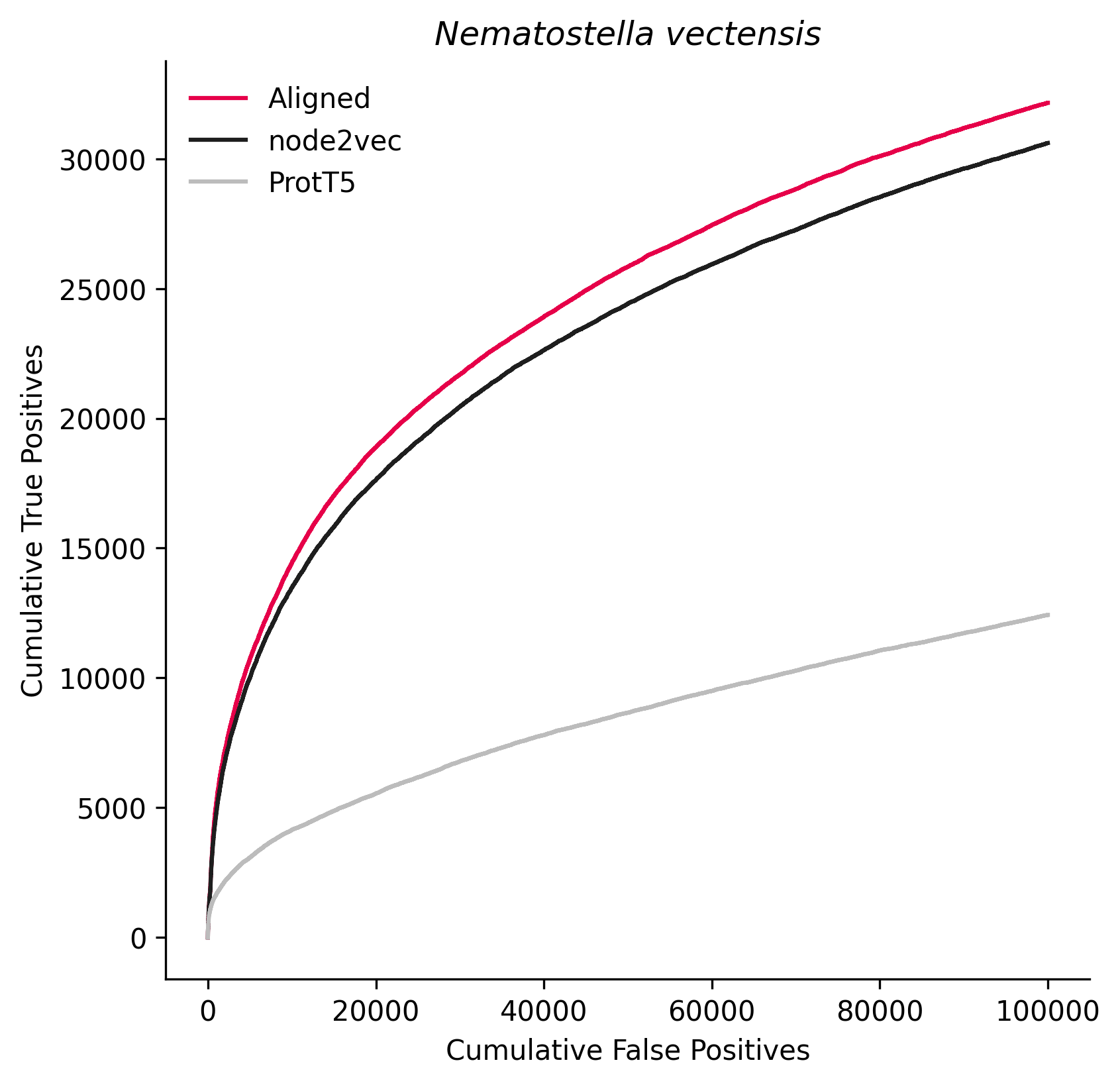

### 46731.png

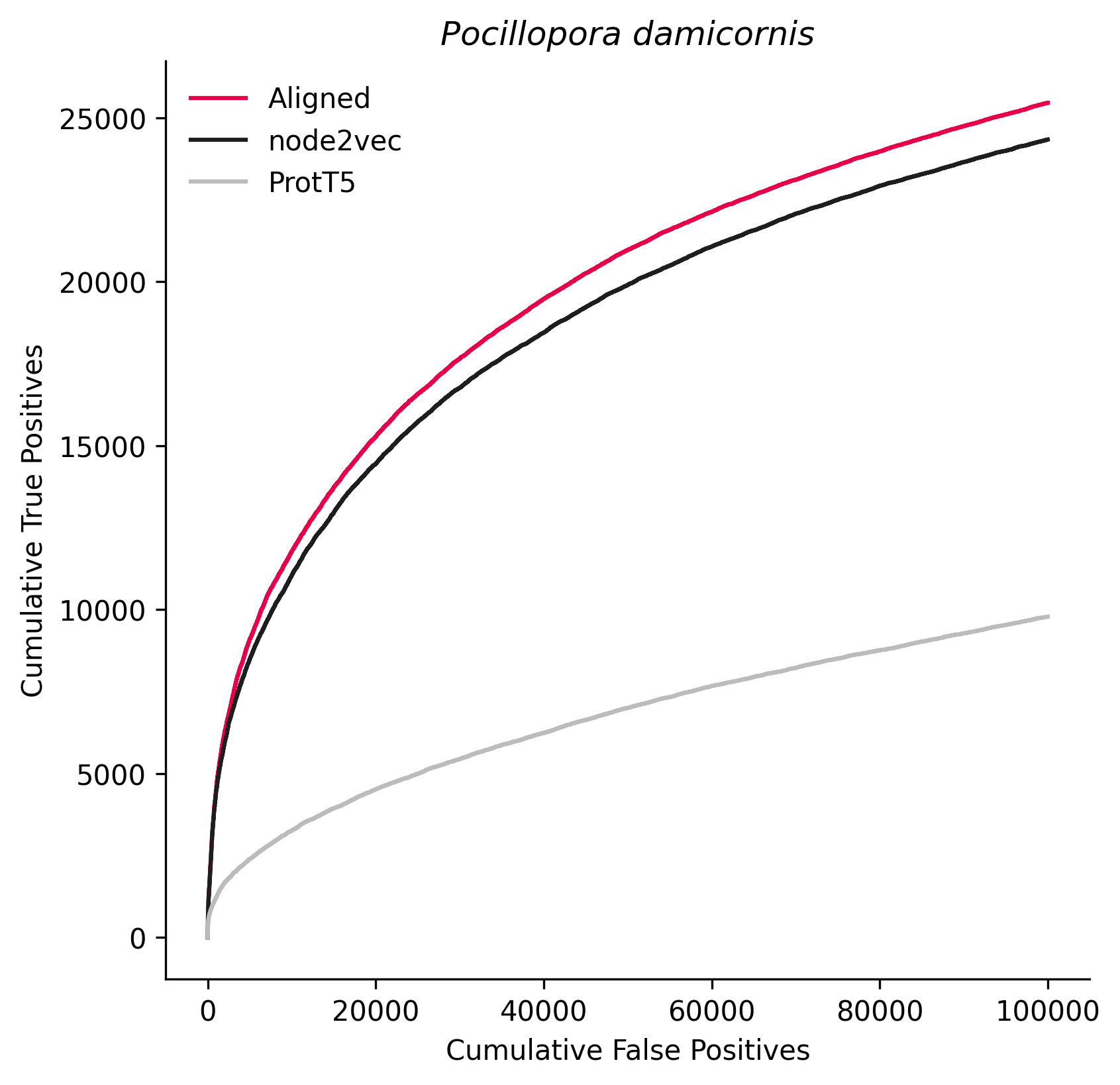

### 51351.png

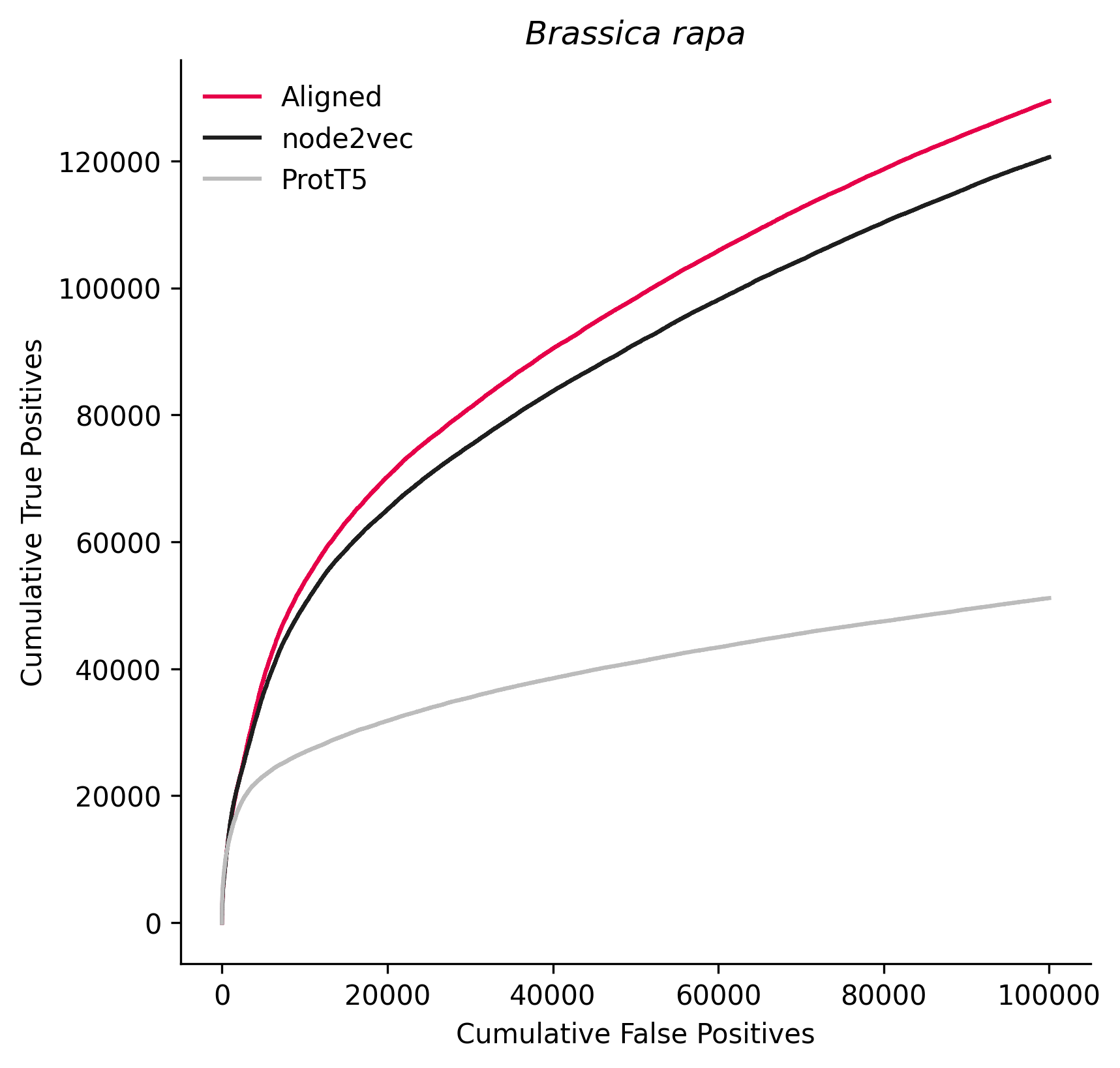

### 59729.png

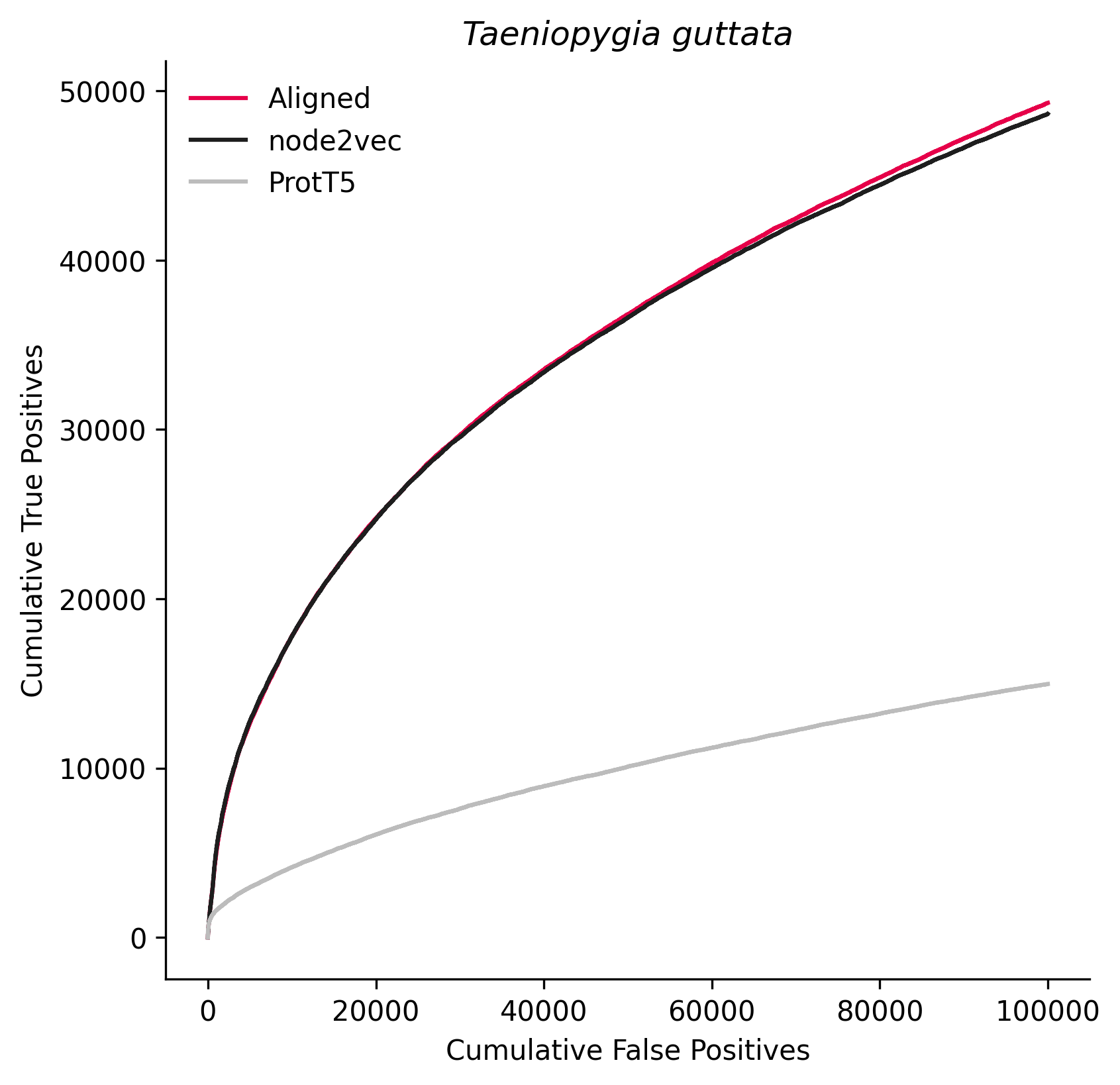

### 64176.png

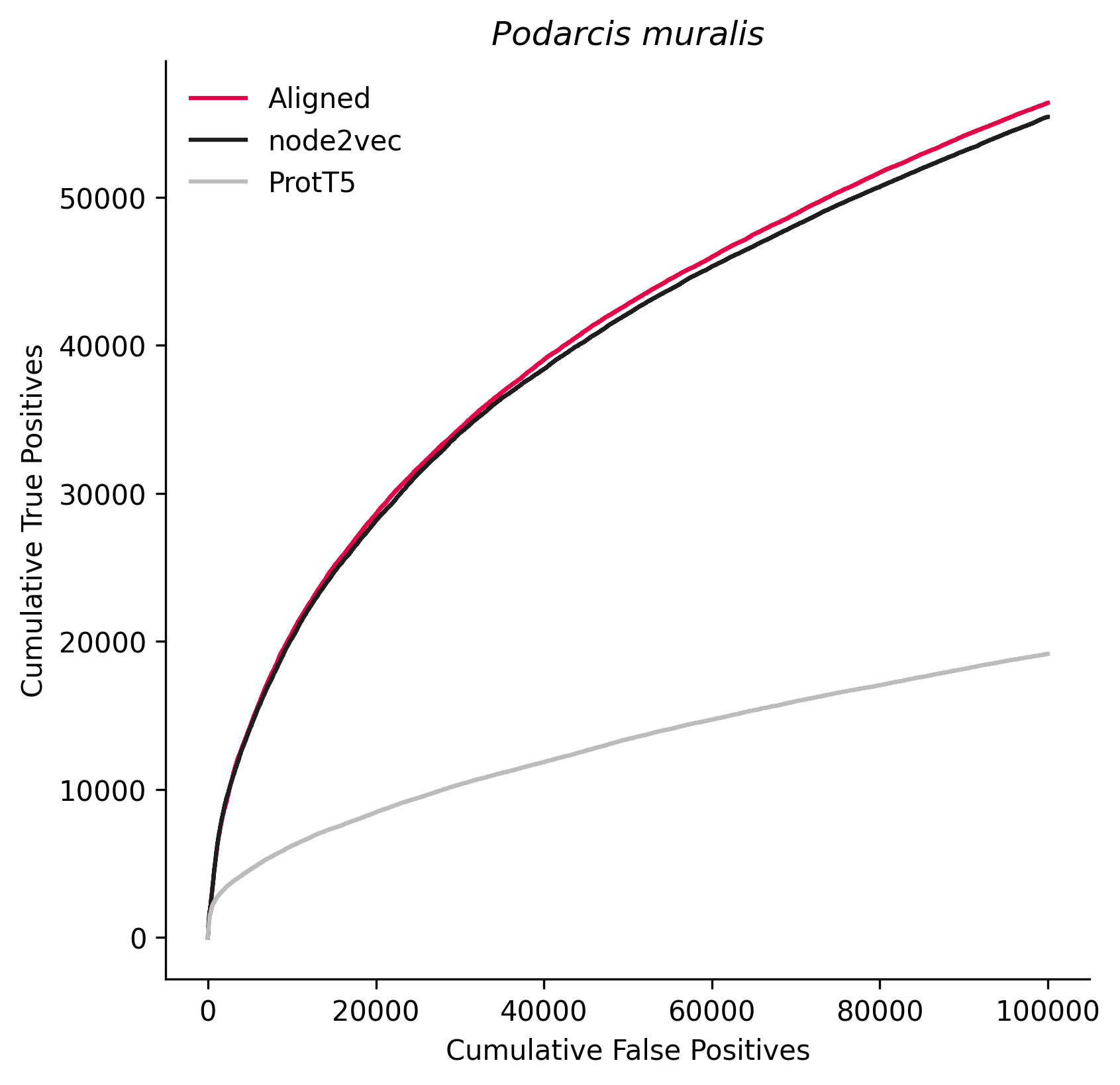

### 70448.png

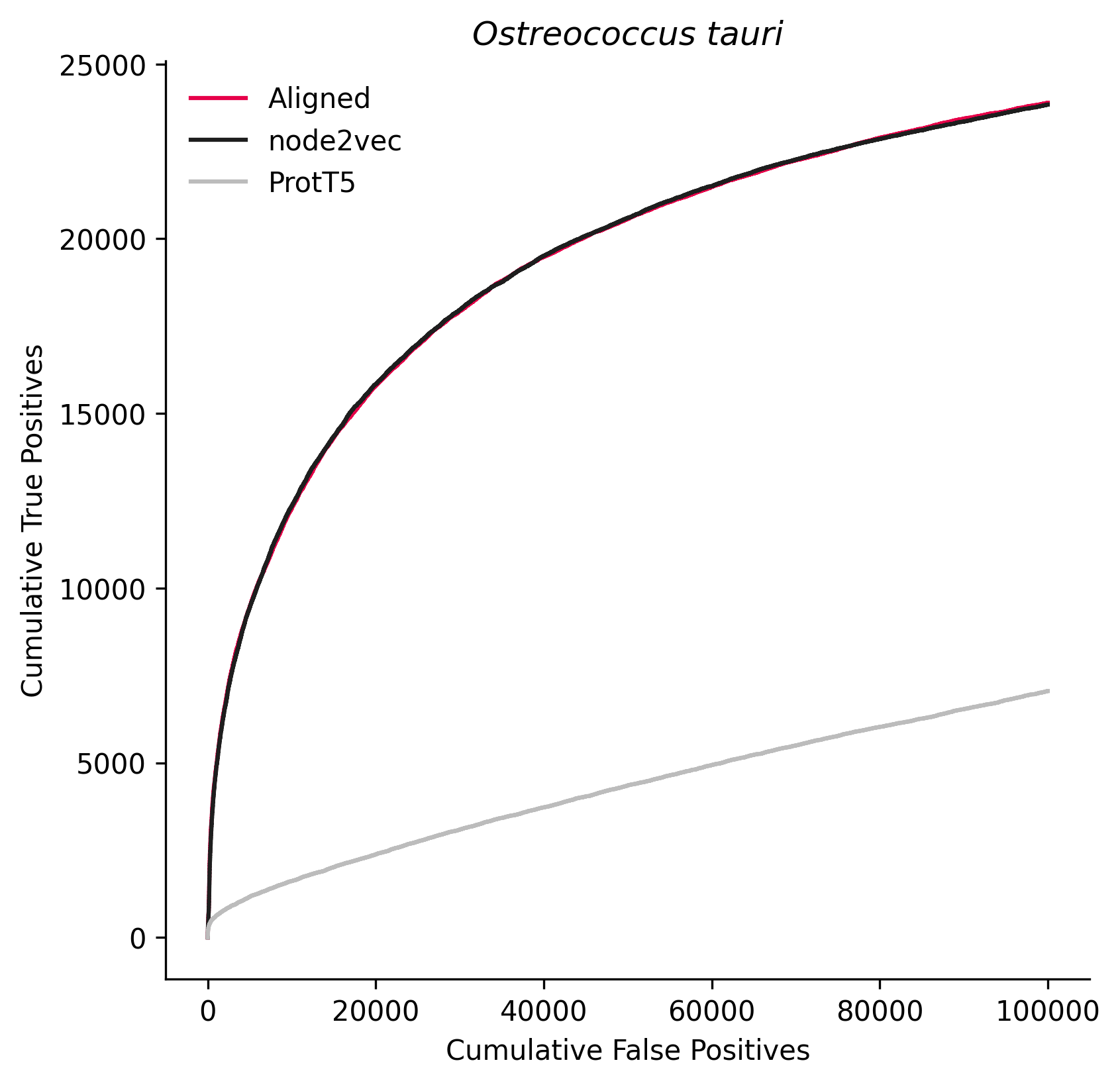

### 74533.png

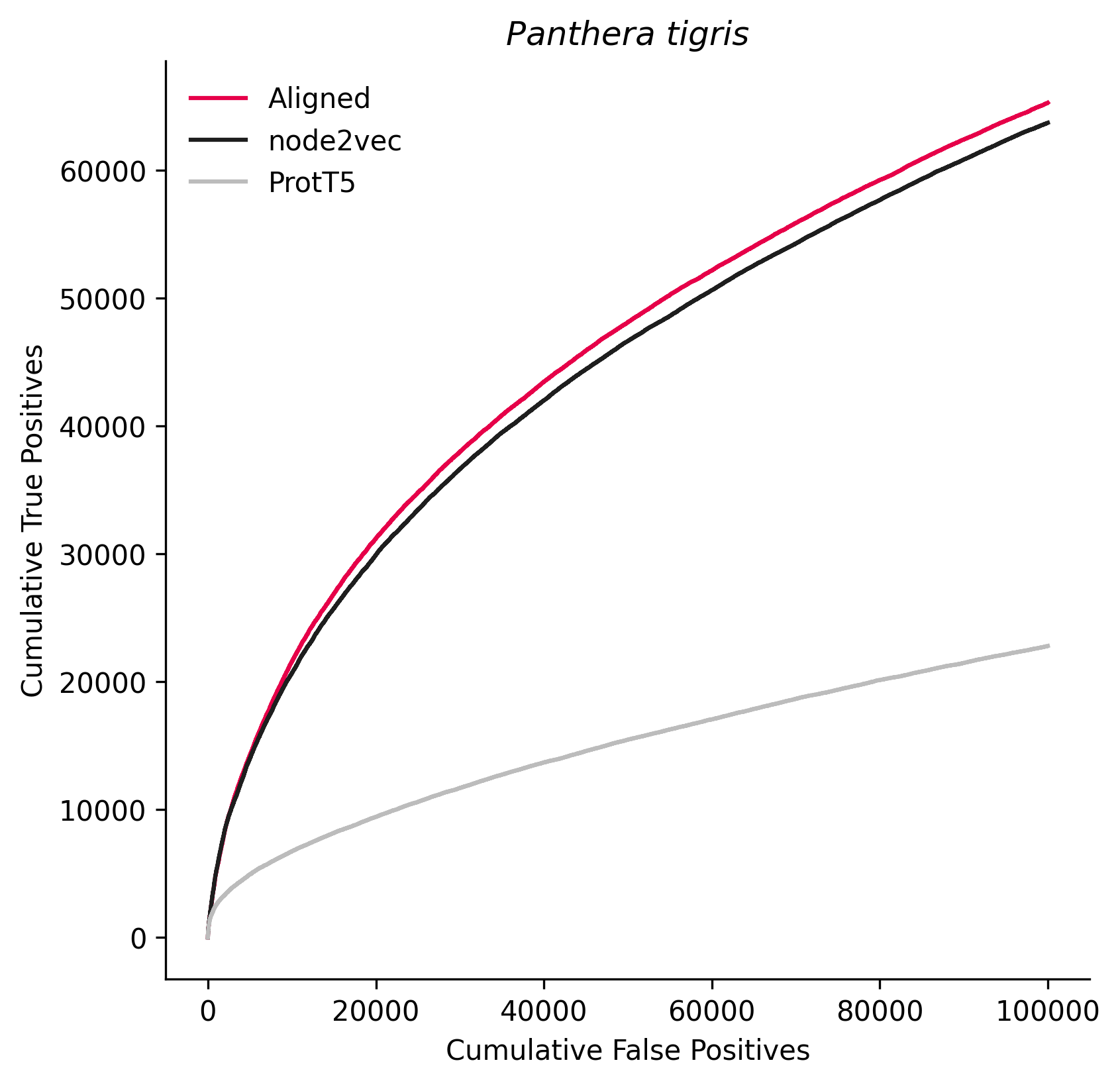

### 79200.png

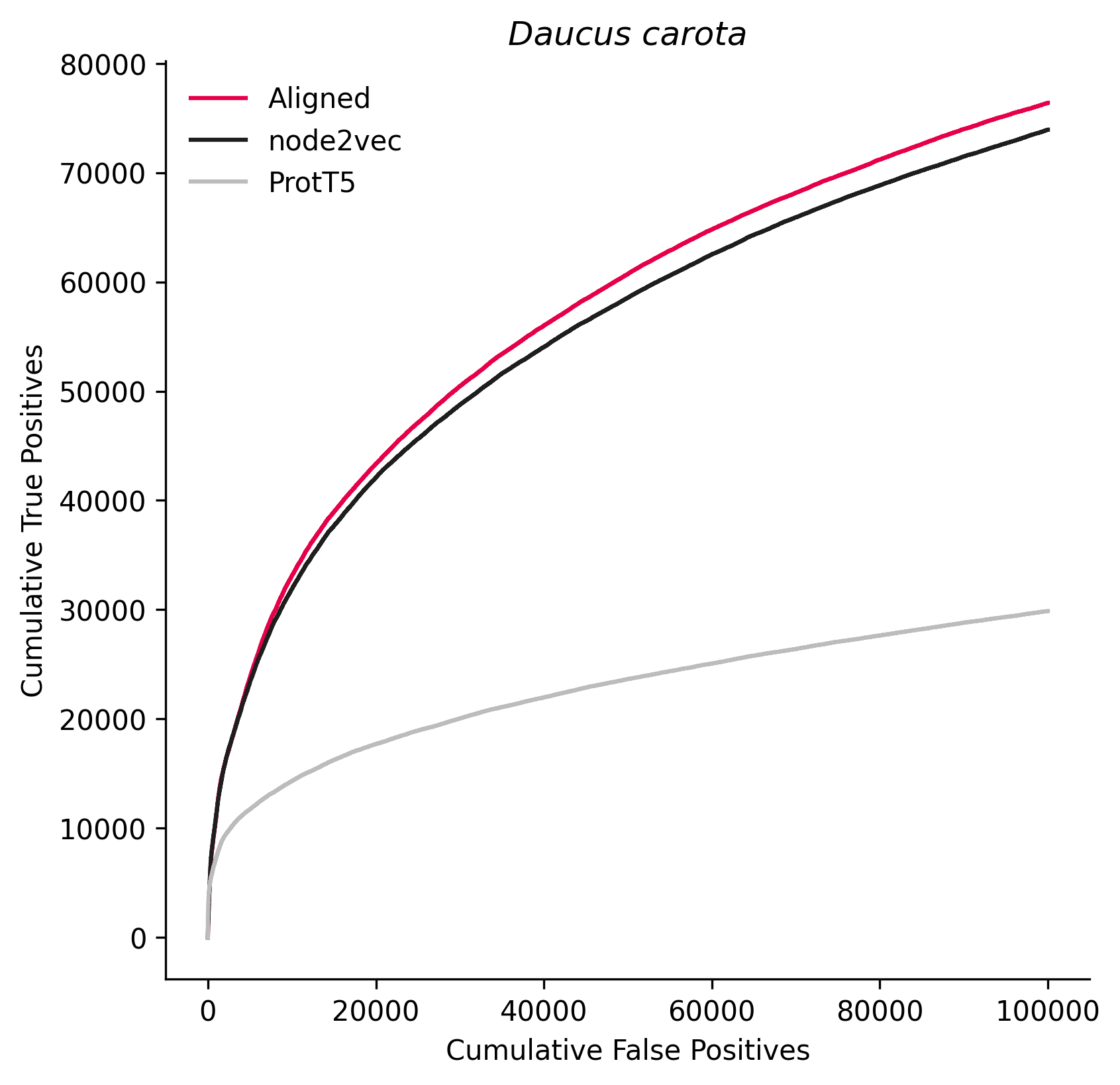

### 80972.png

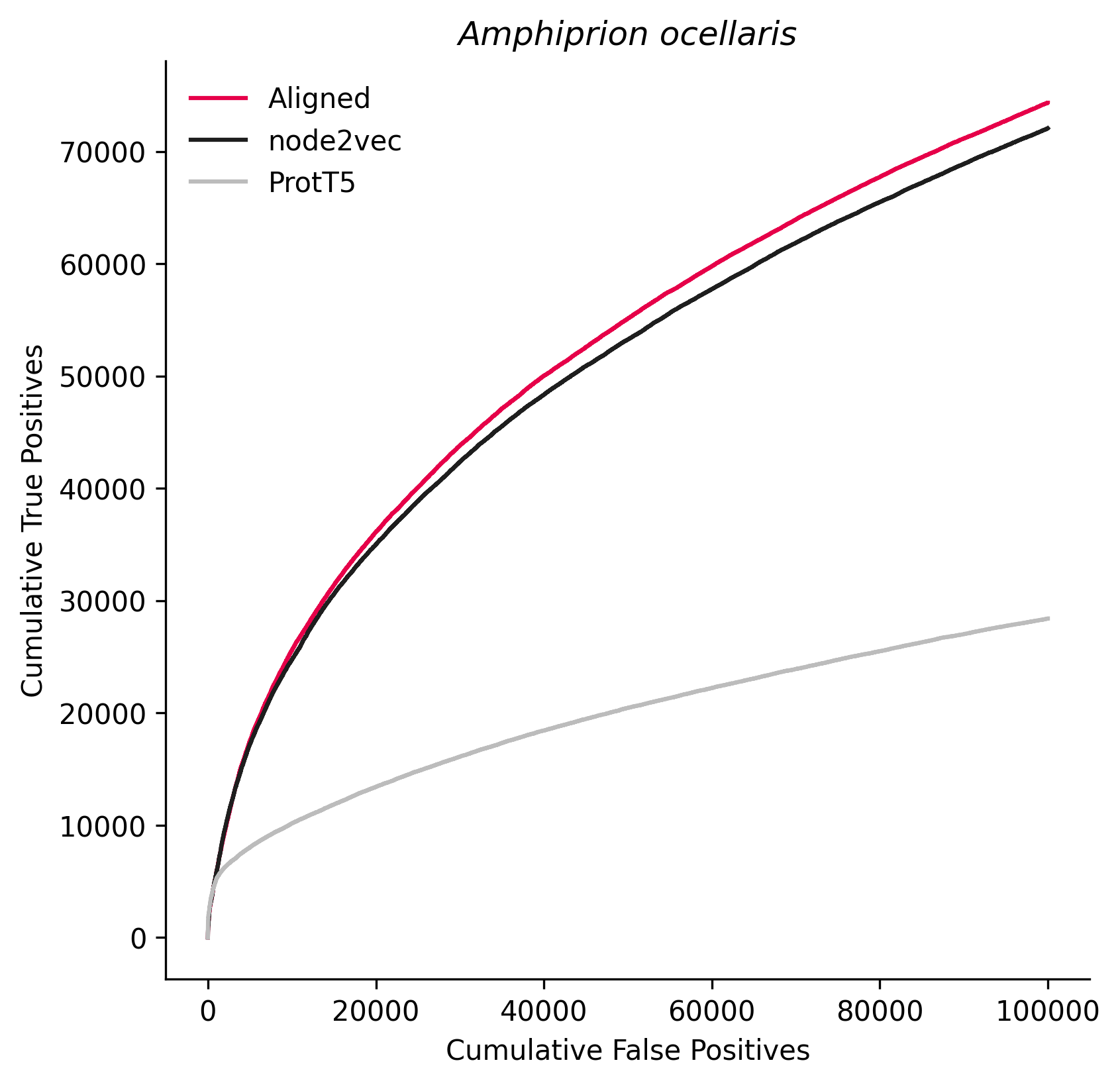

### 93934.png

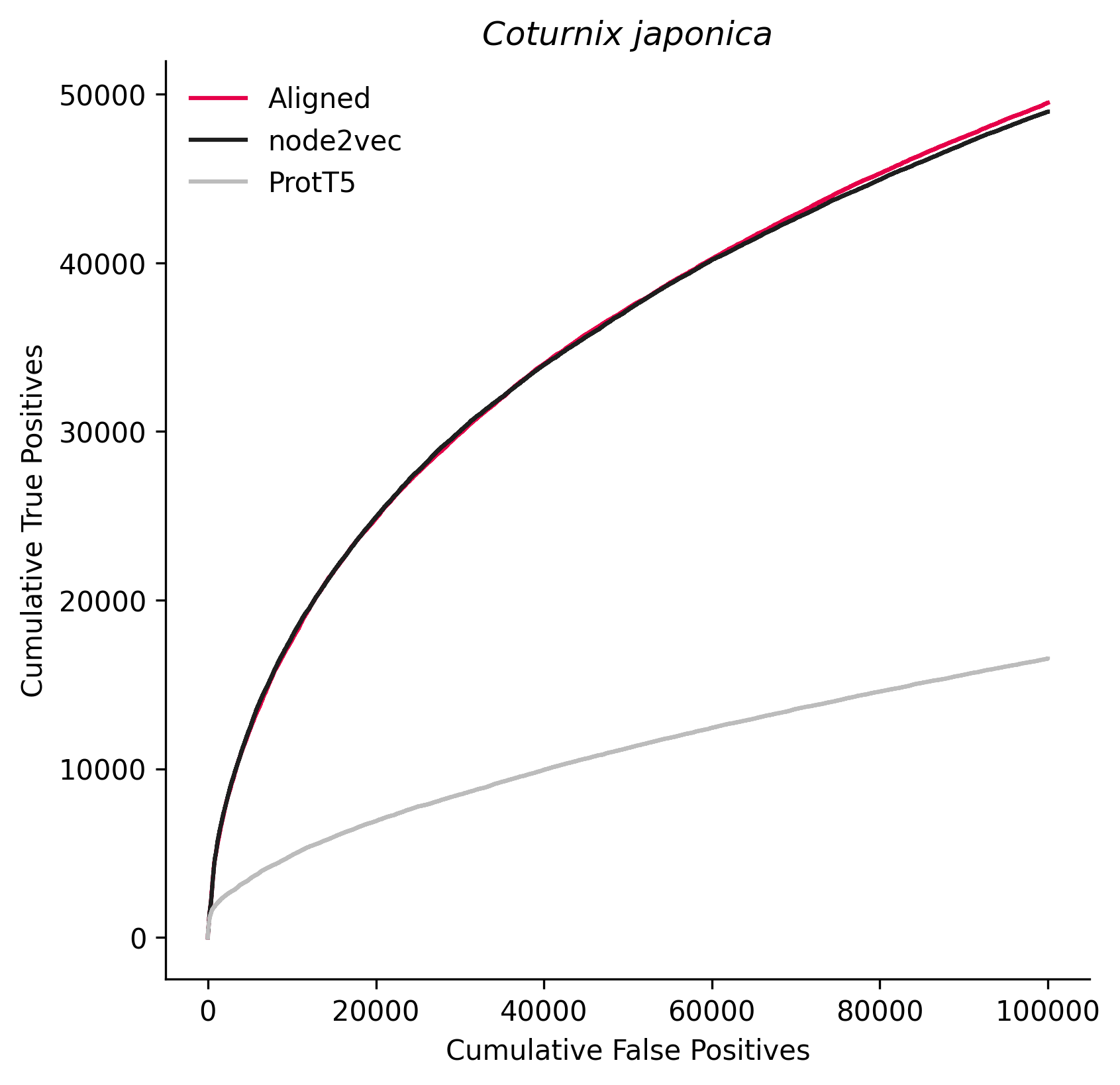

### 109280.png

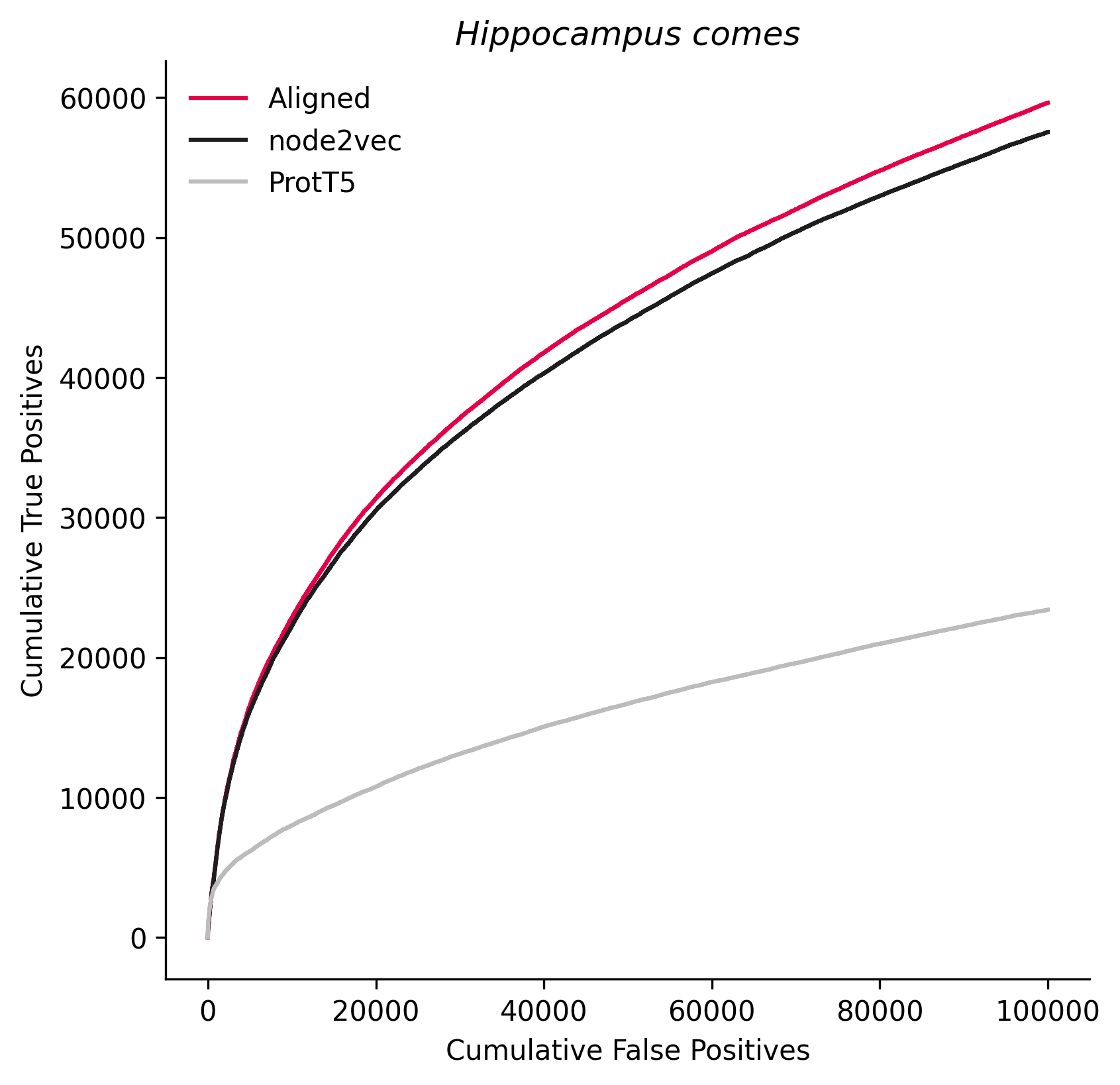

### 113540.png

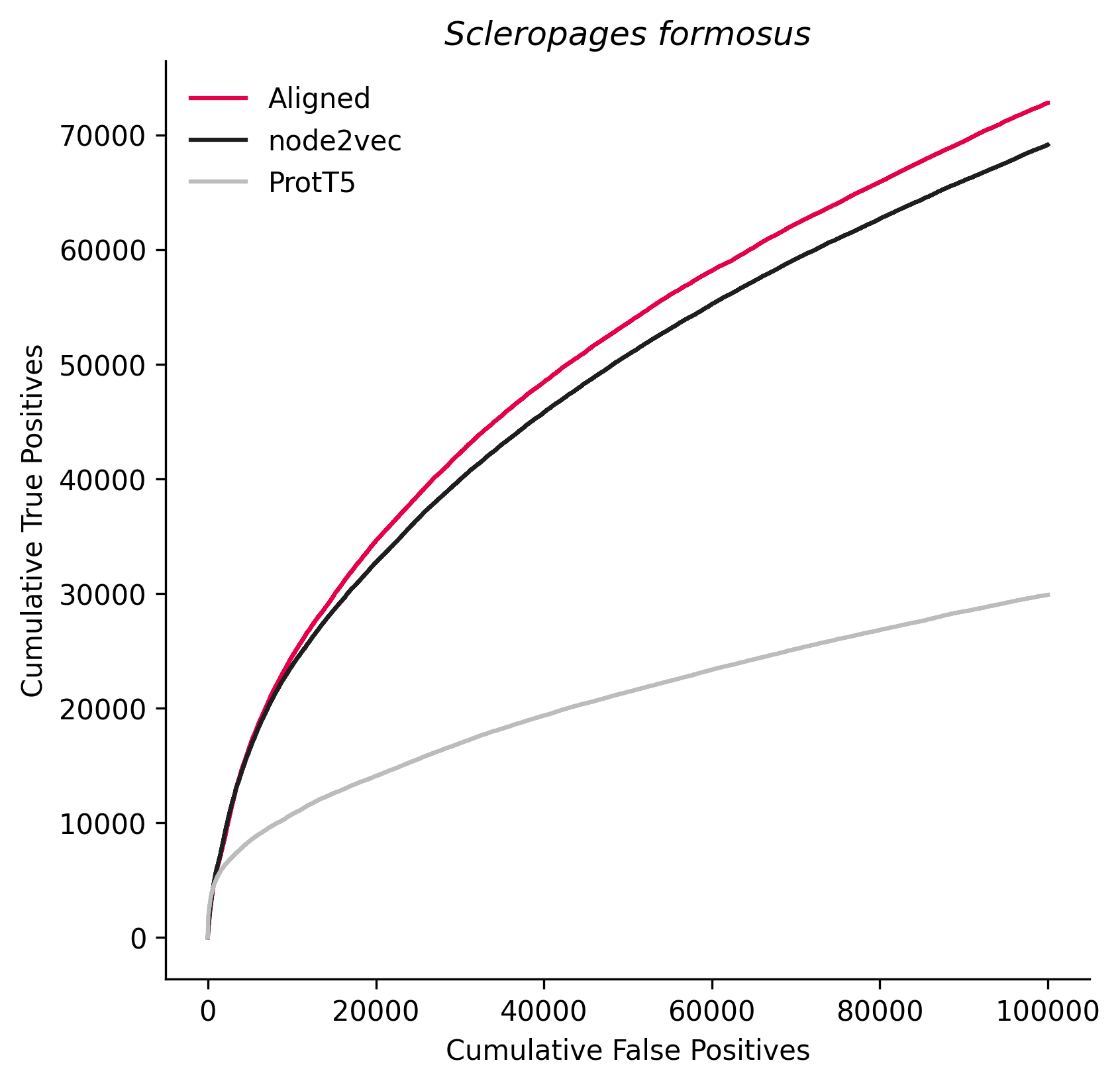

### 118797.png

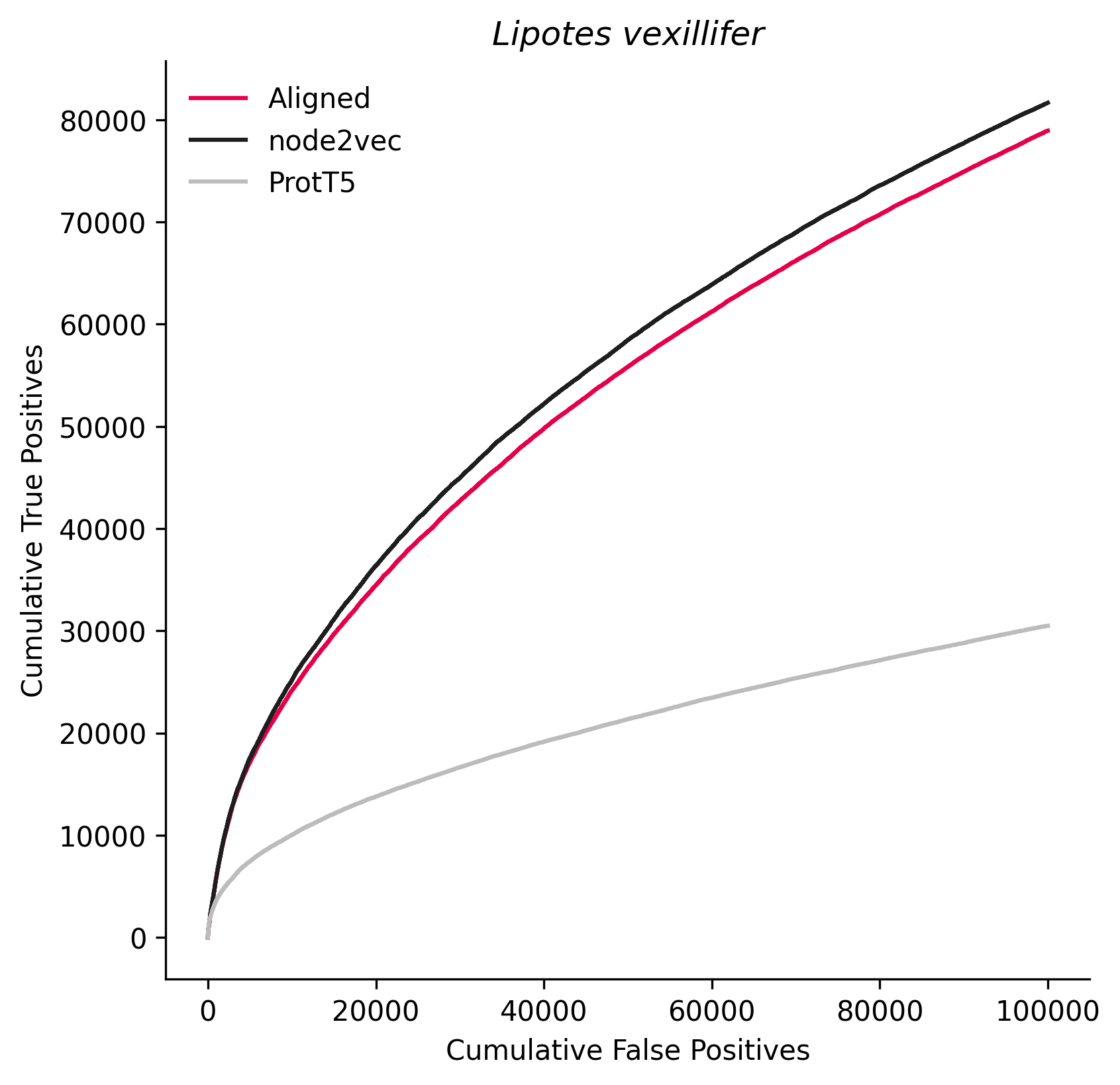

### 145388.png

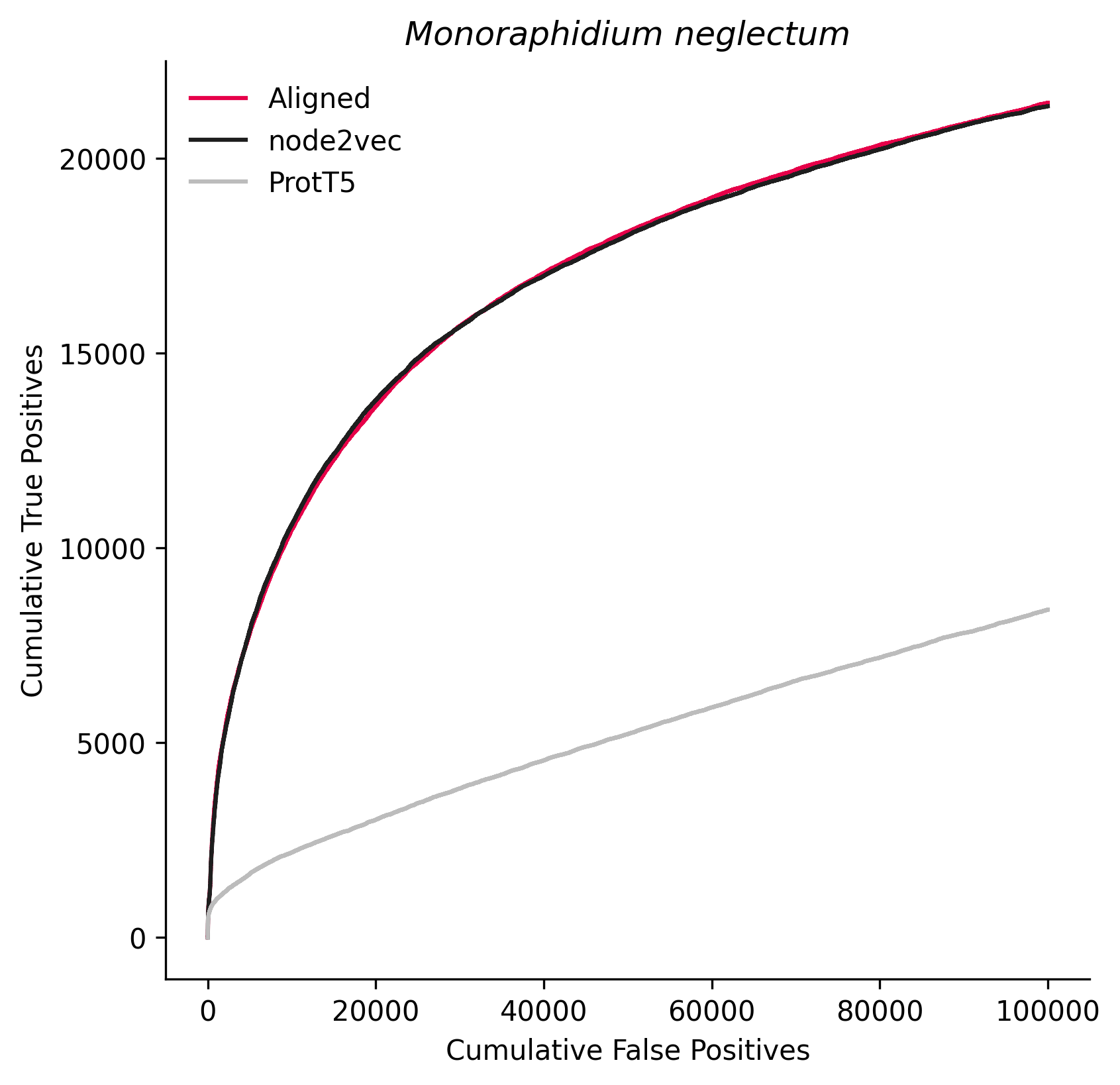

### 149040.png

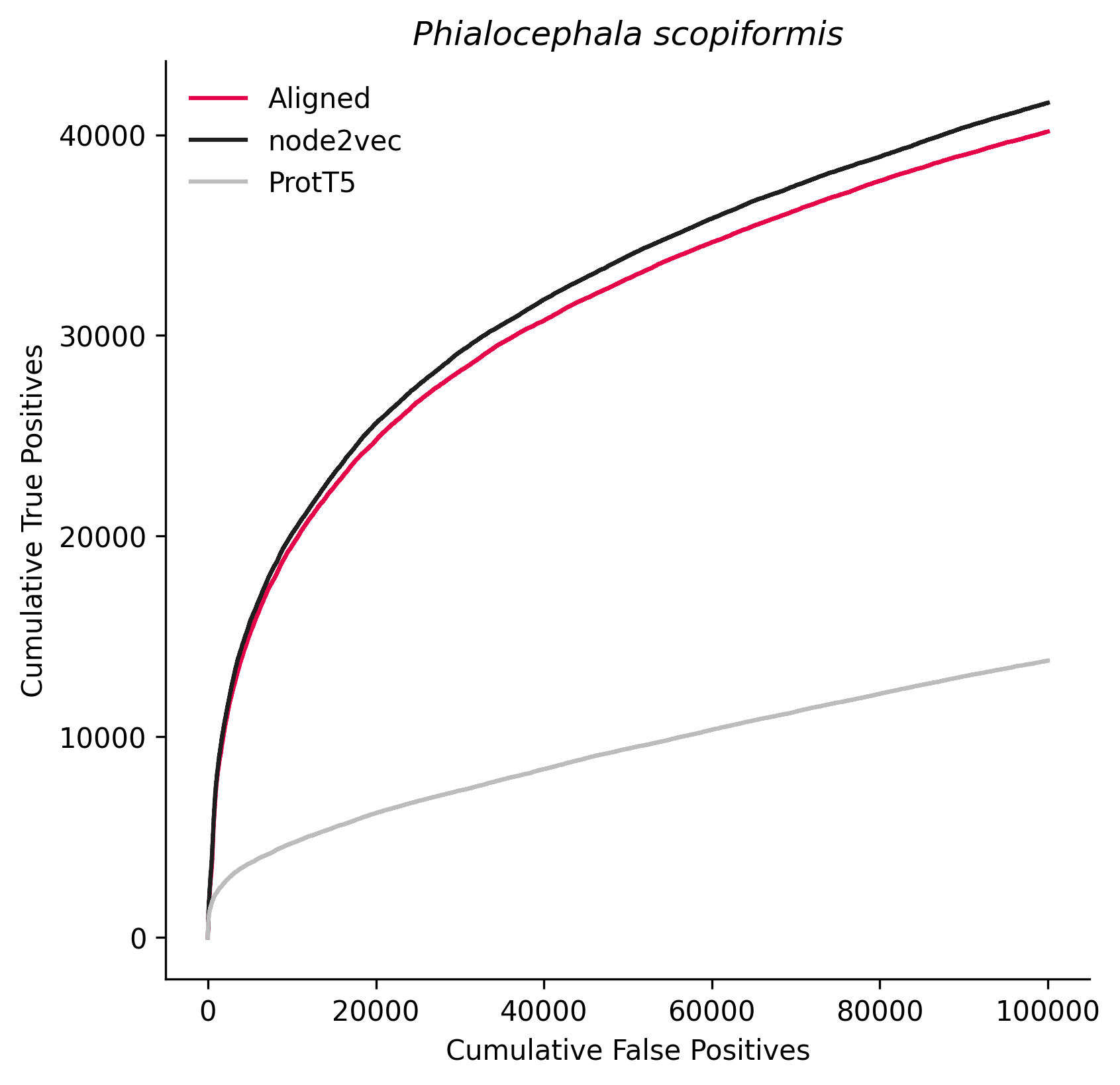

### 156563.png

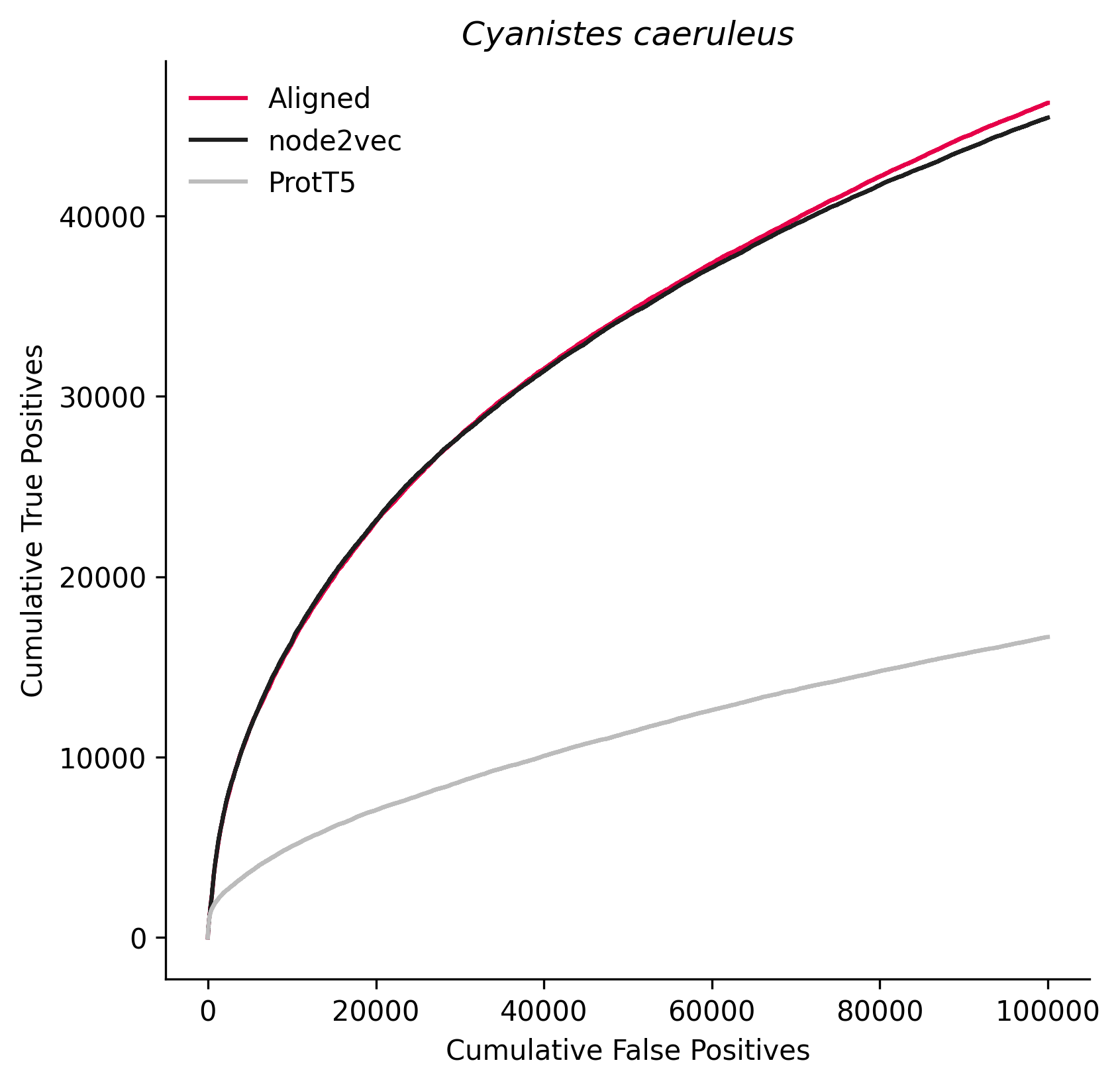

### 158441.png

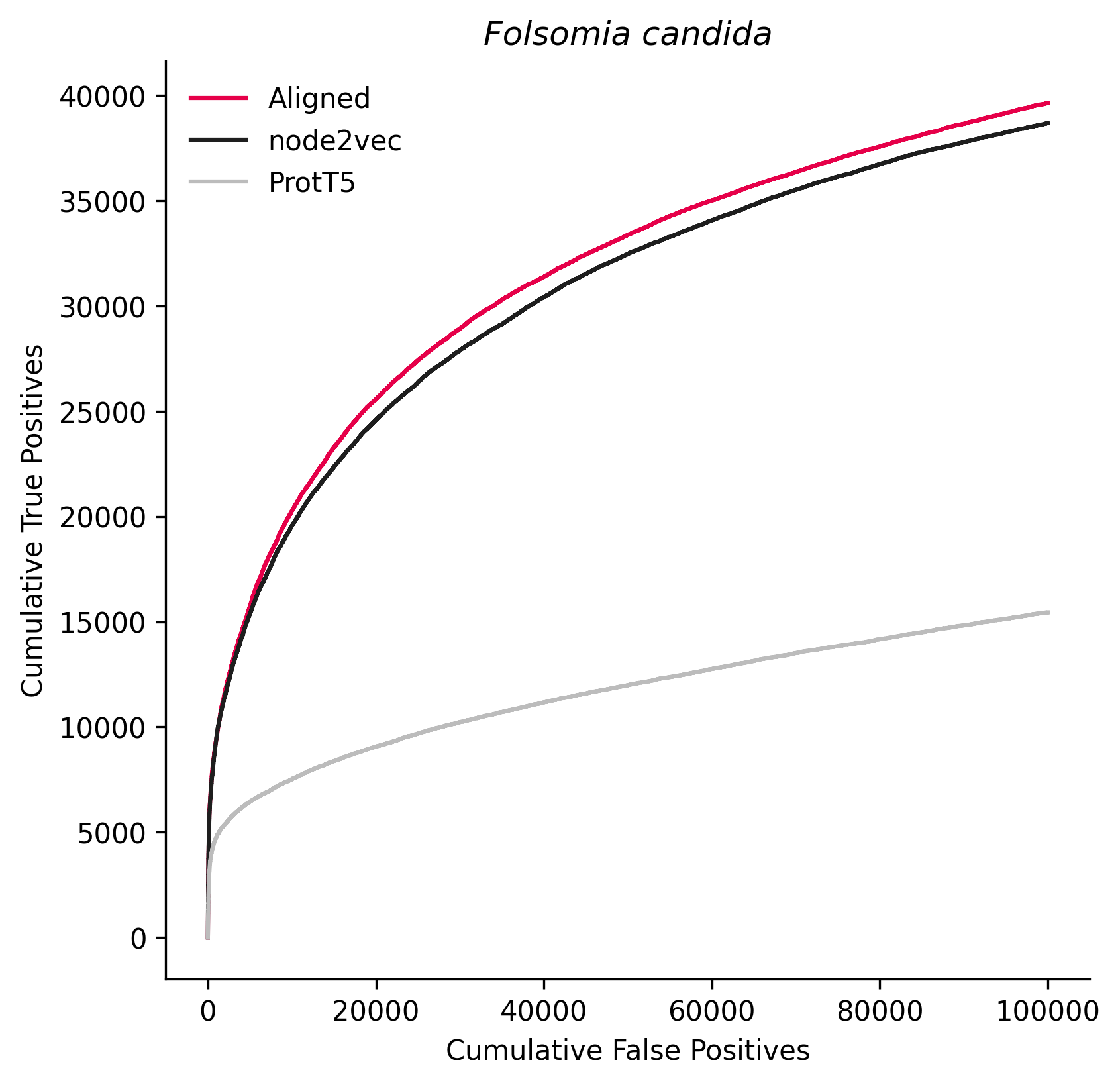

### 188379.png

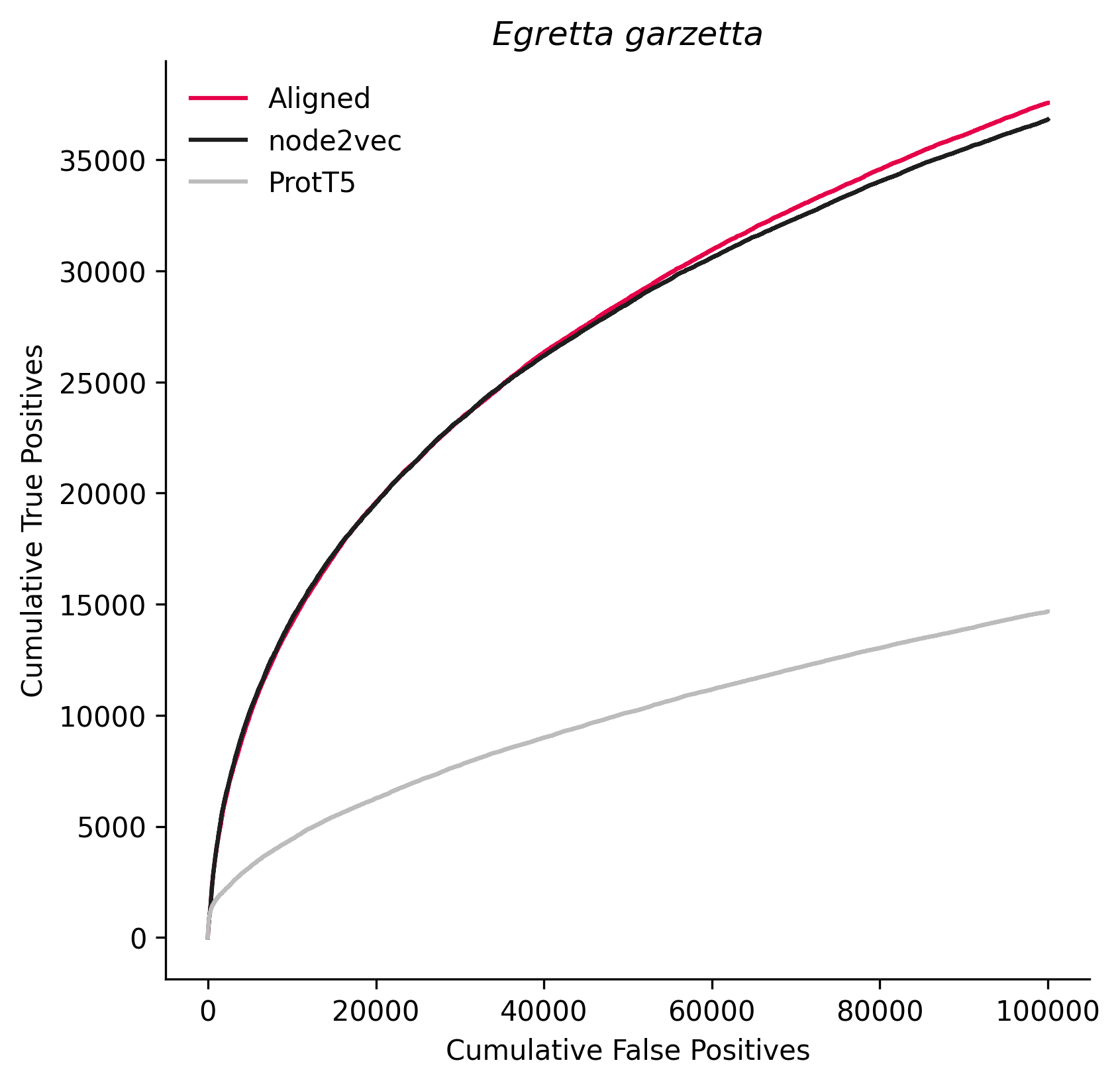

### 214684.png

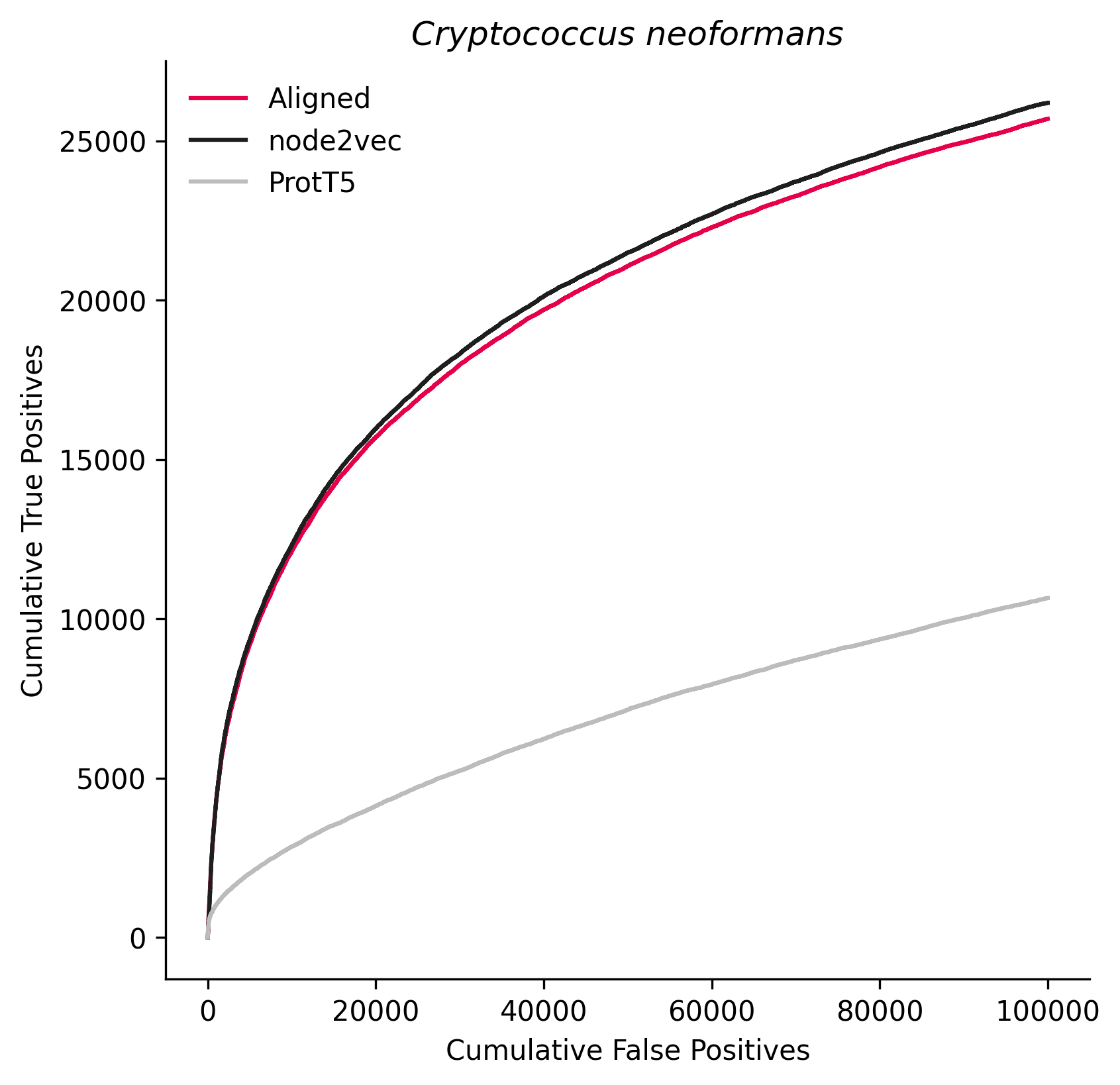

### 225164.png

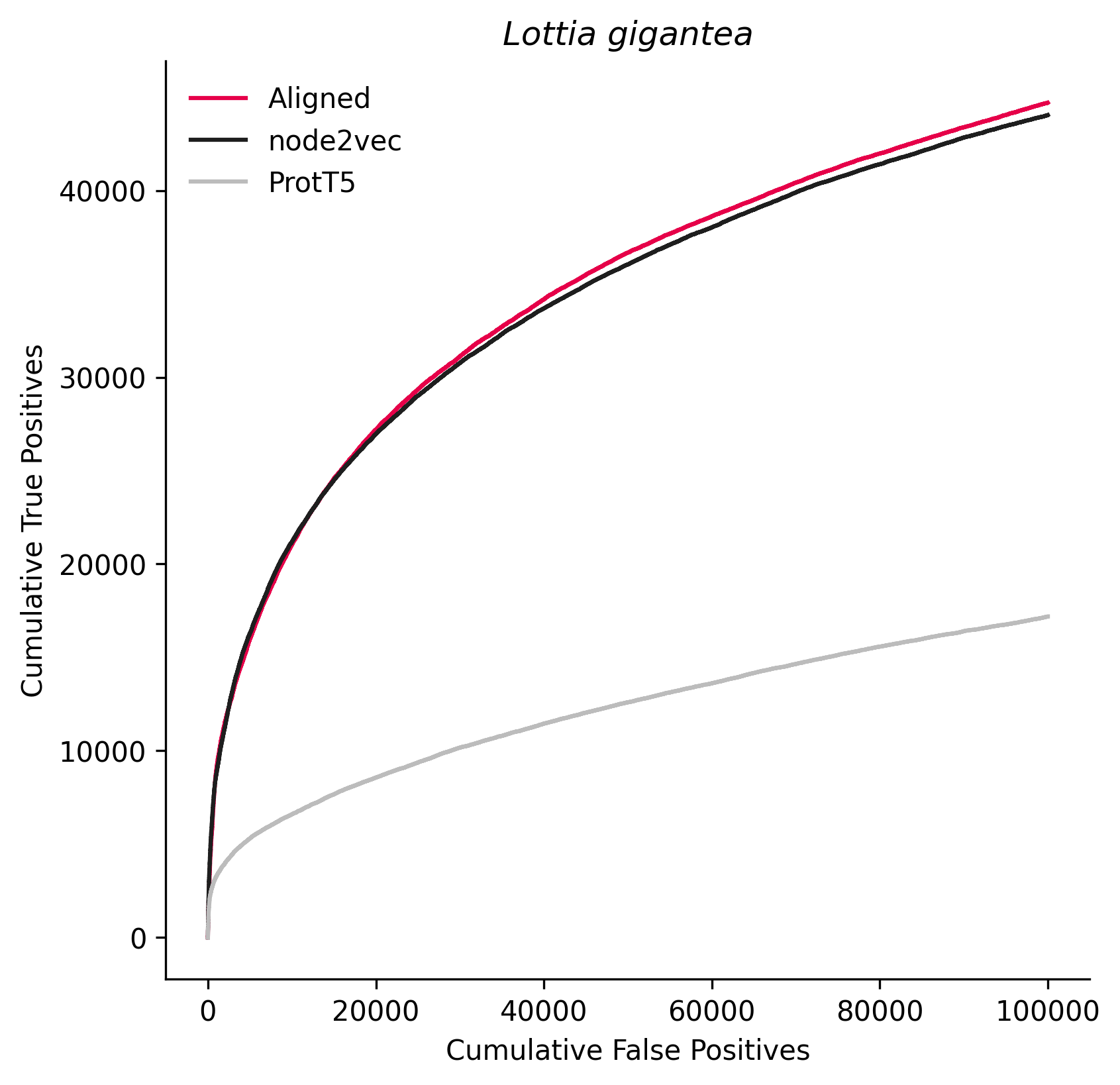

### 229533.png

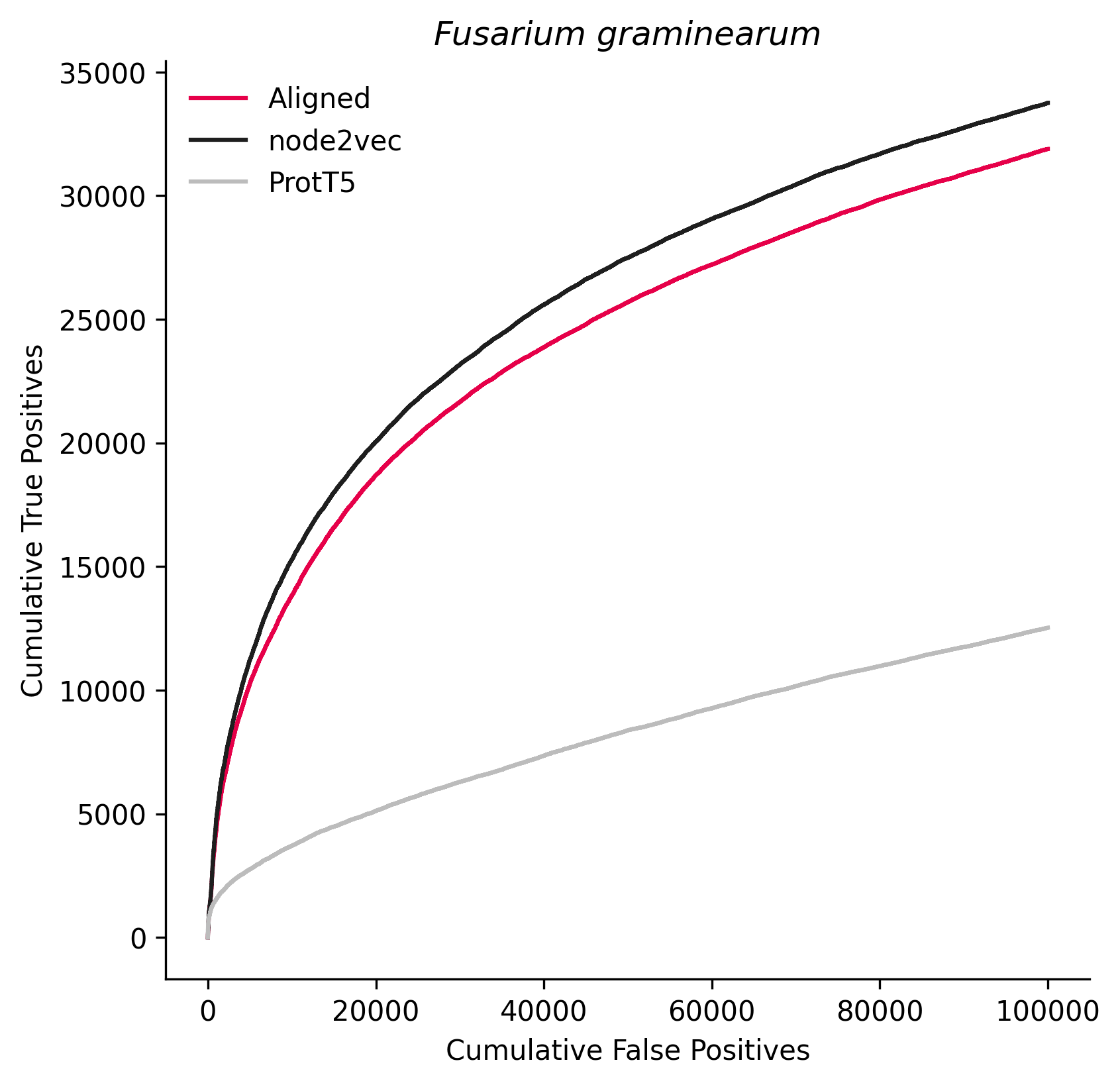

### 242507.png

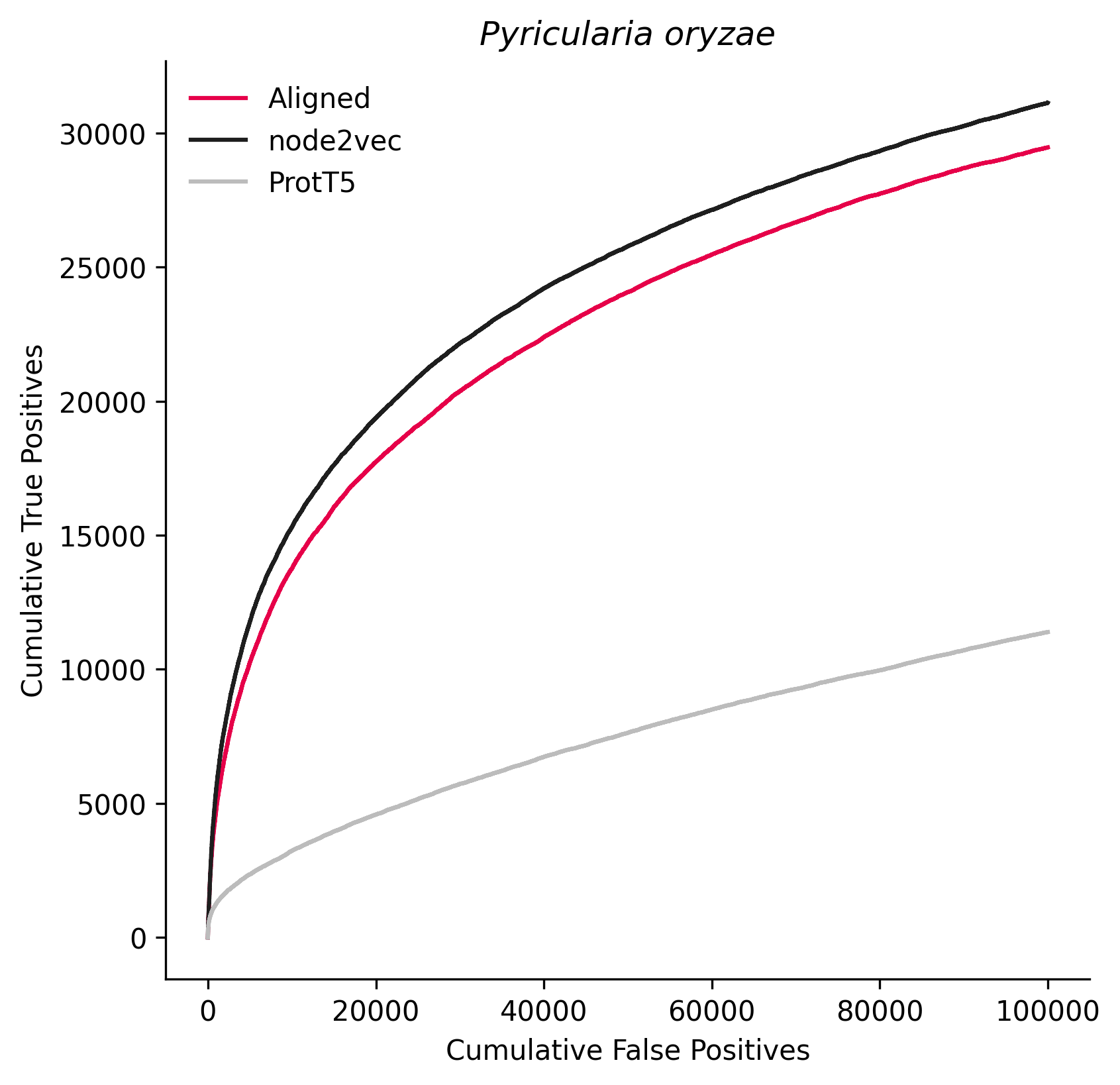

### 284590.png

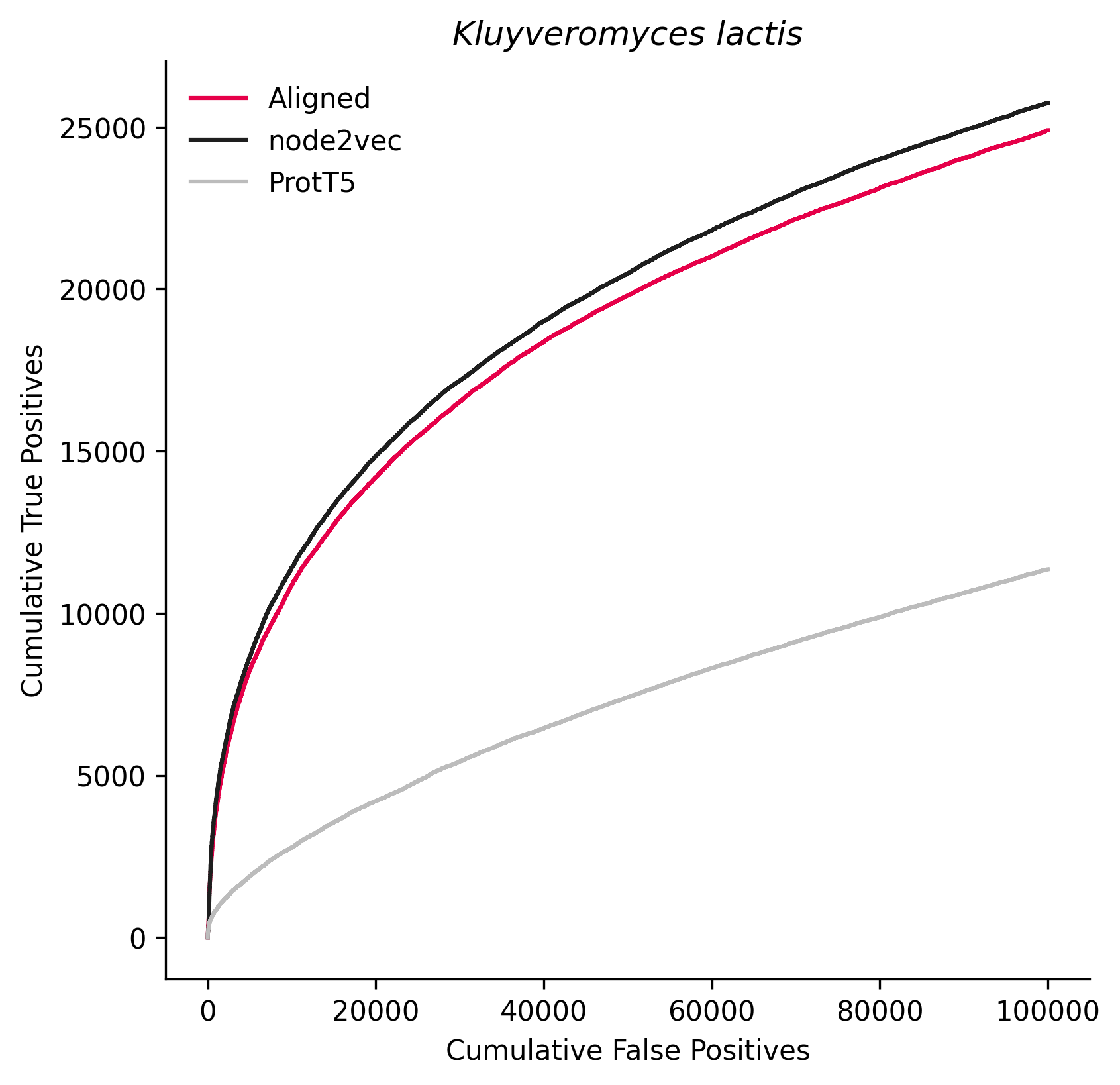

### 284592.png

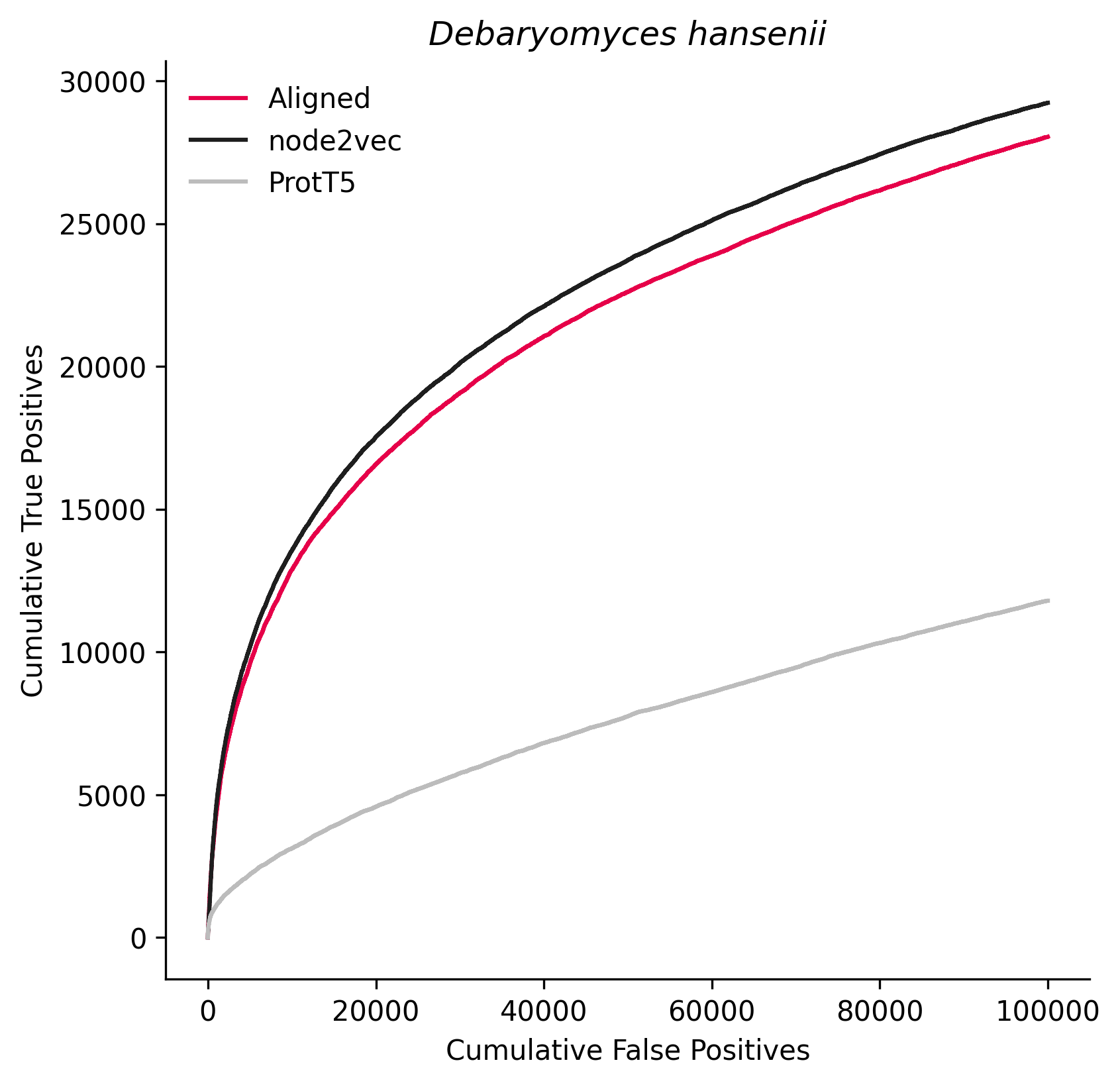

### 296587.png

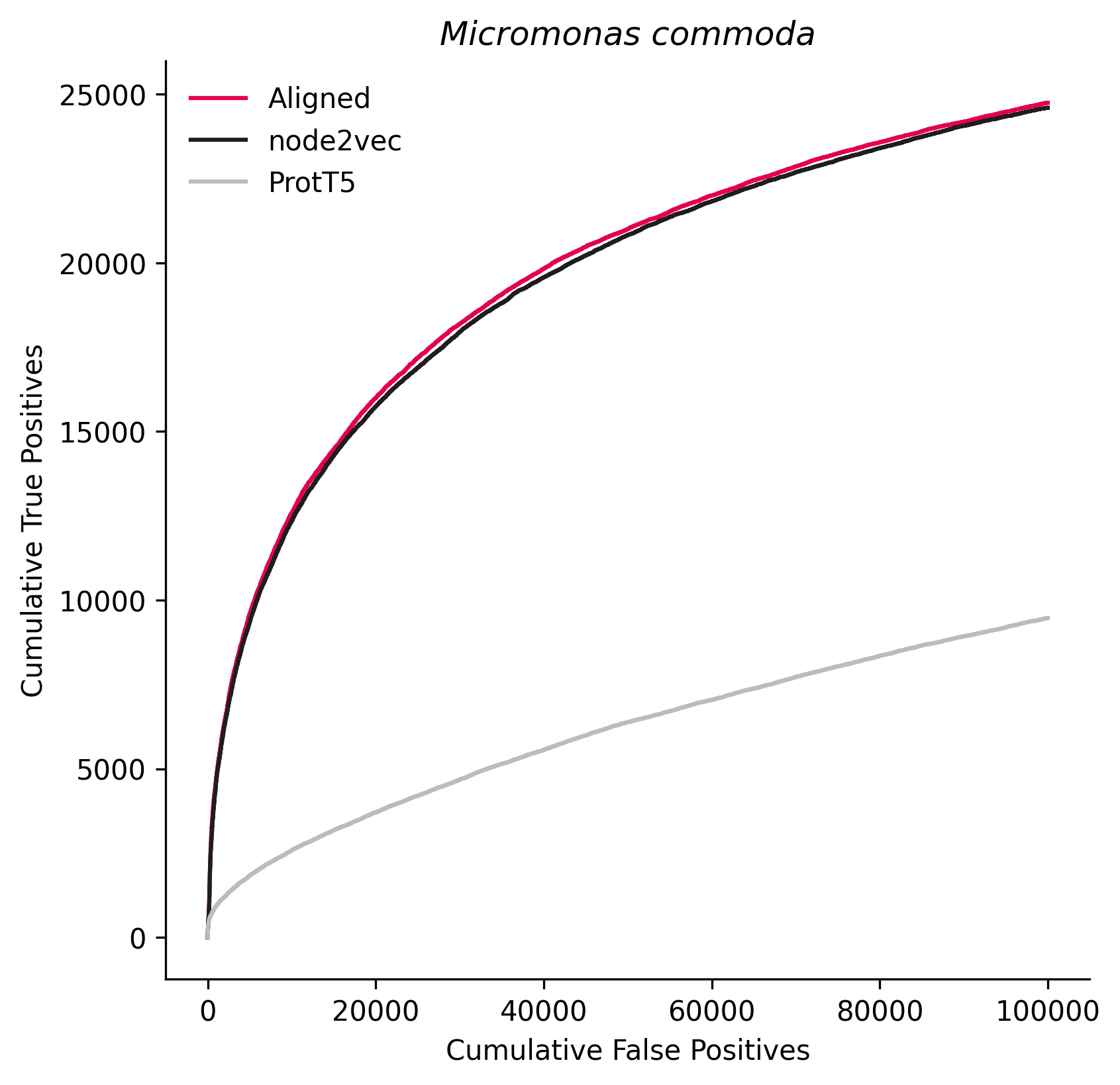

### 310752.png

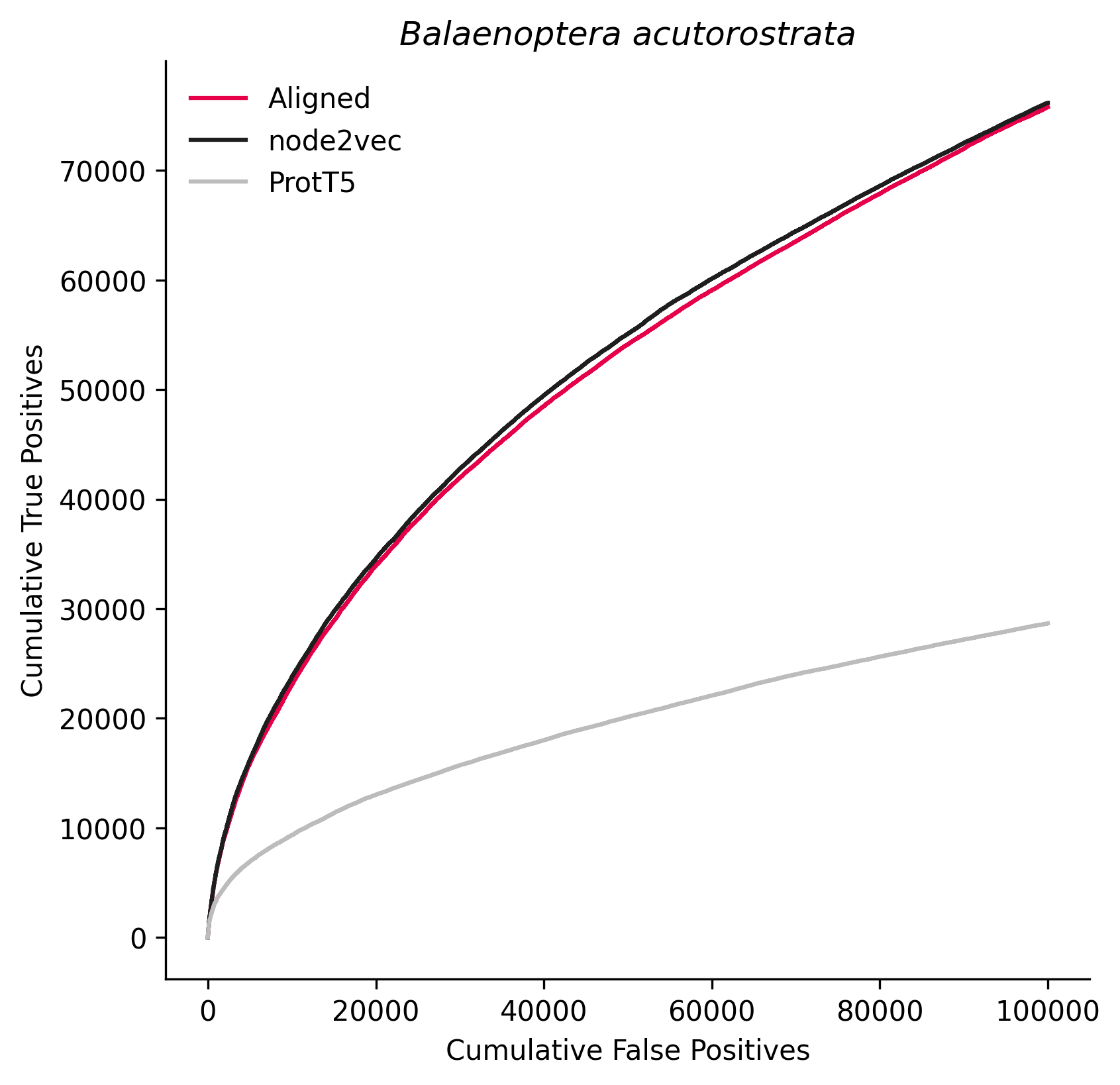

### 322104.png

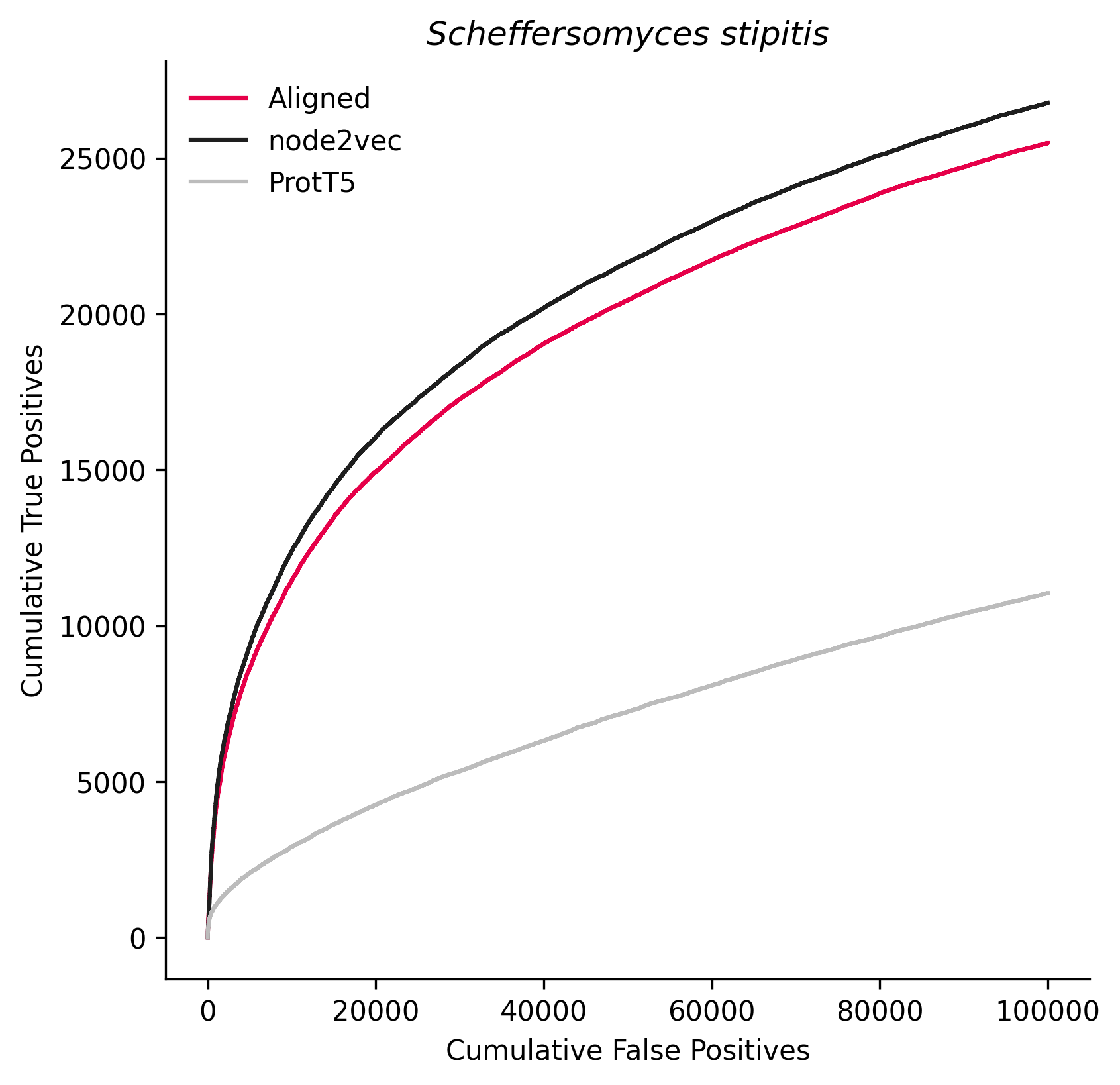

### 344612.png

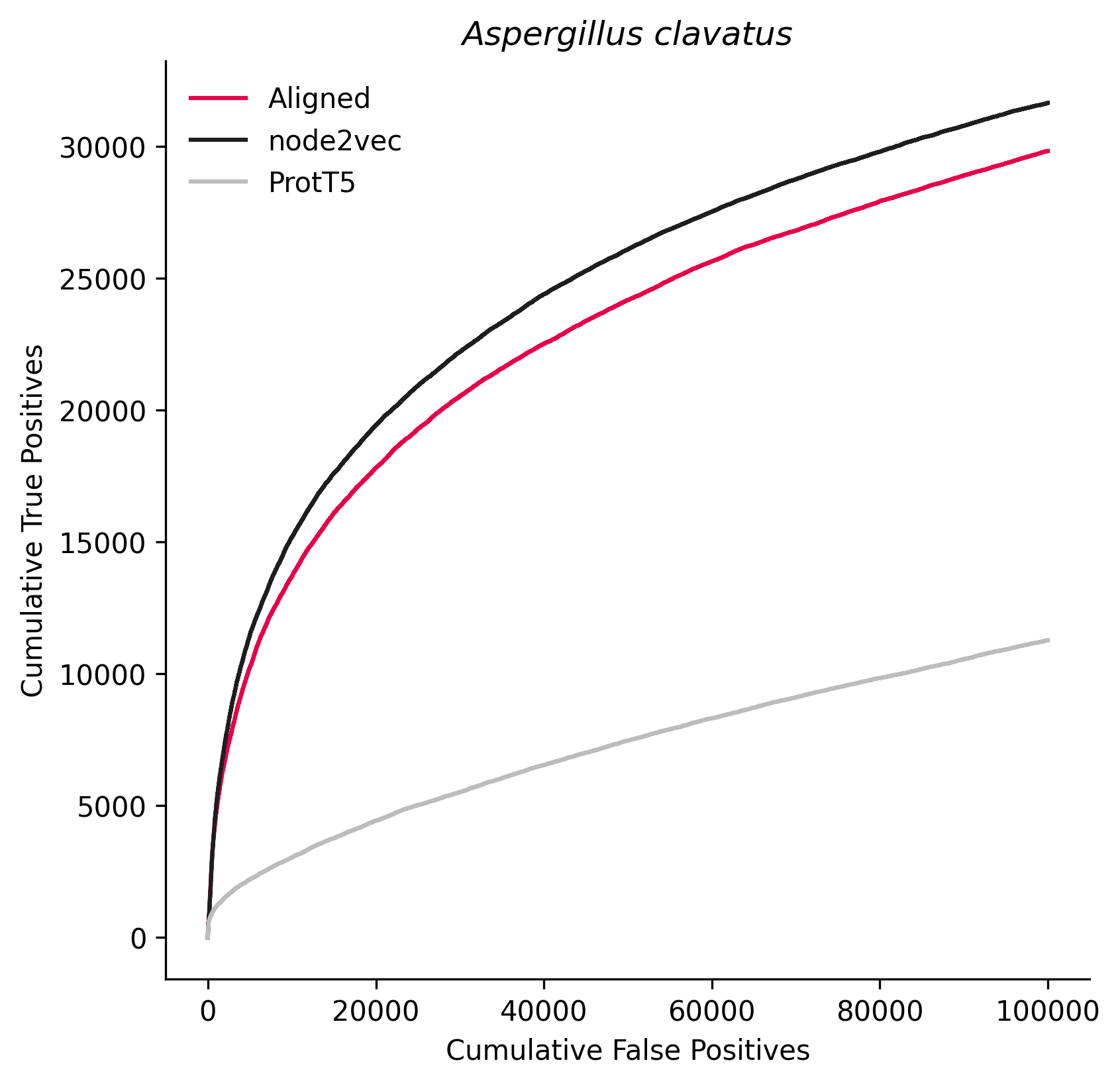

### 379508.png

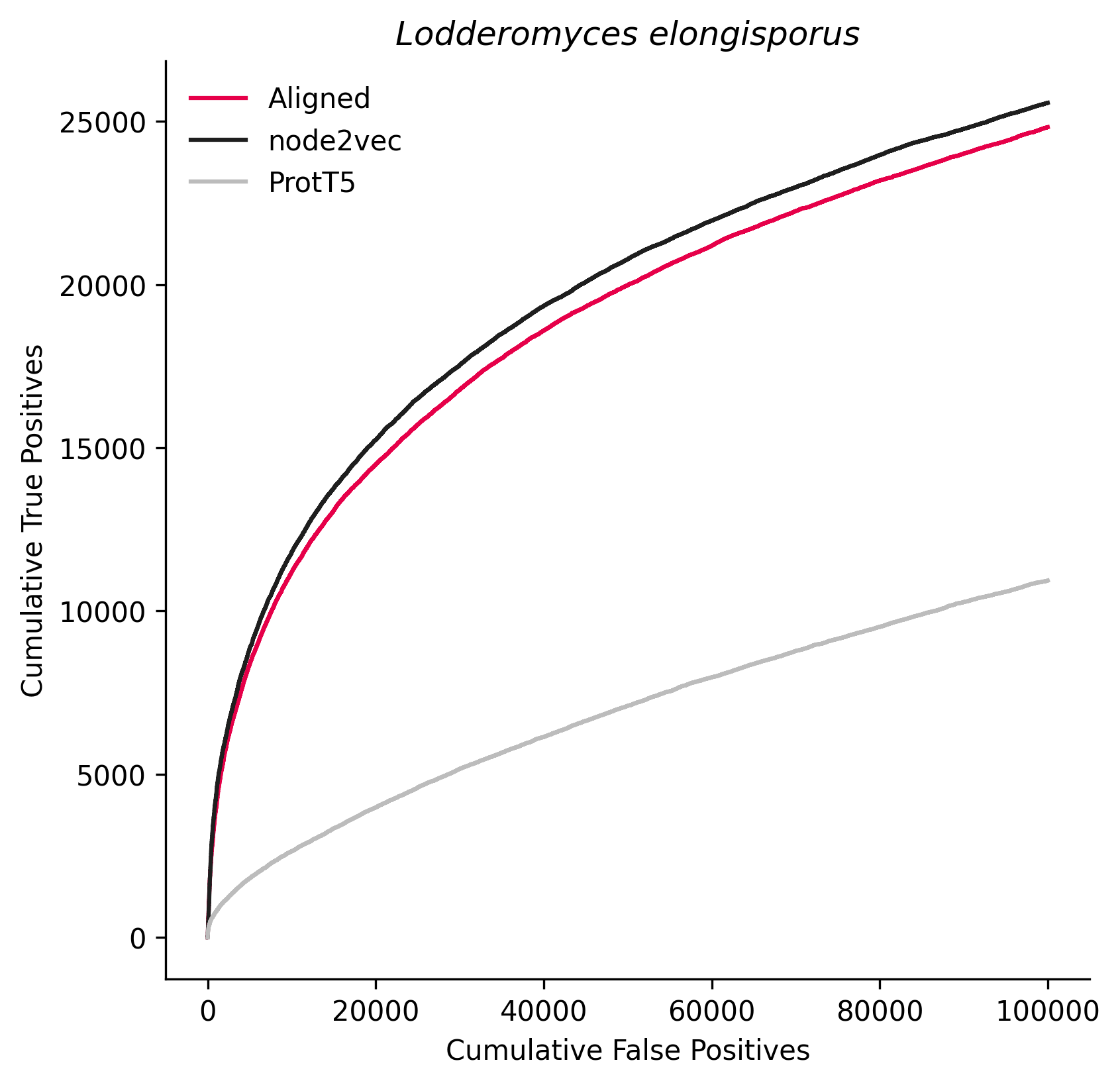

### 391180.png

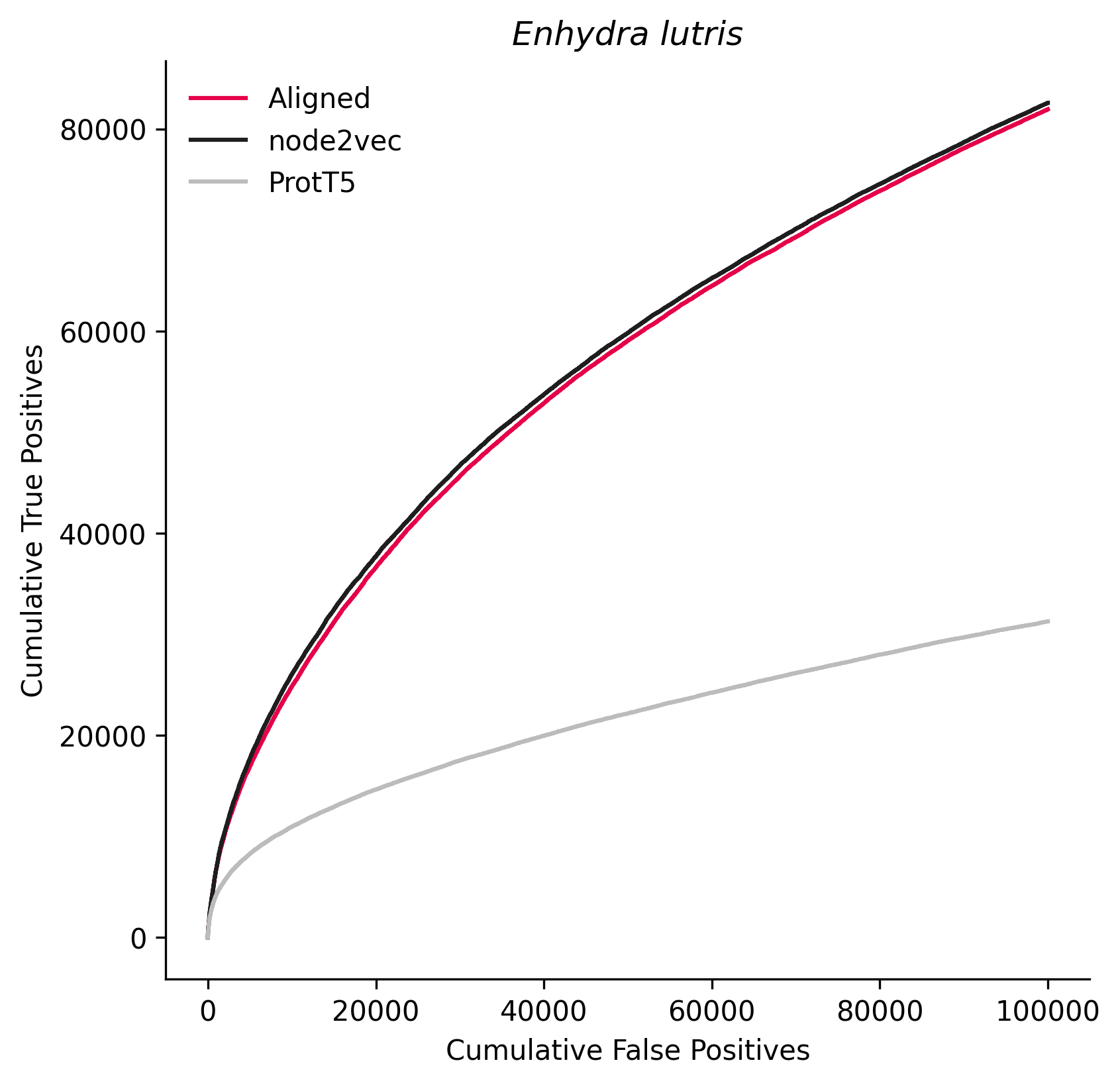

### 425265.png

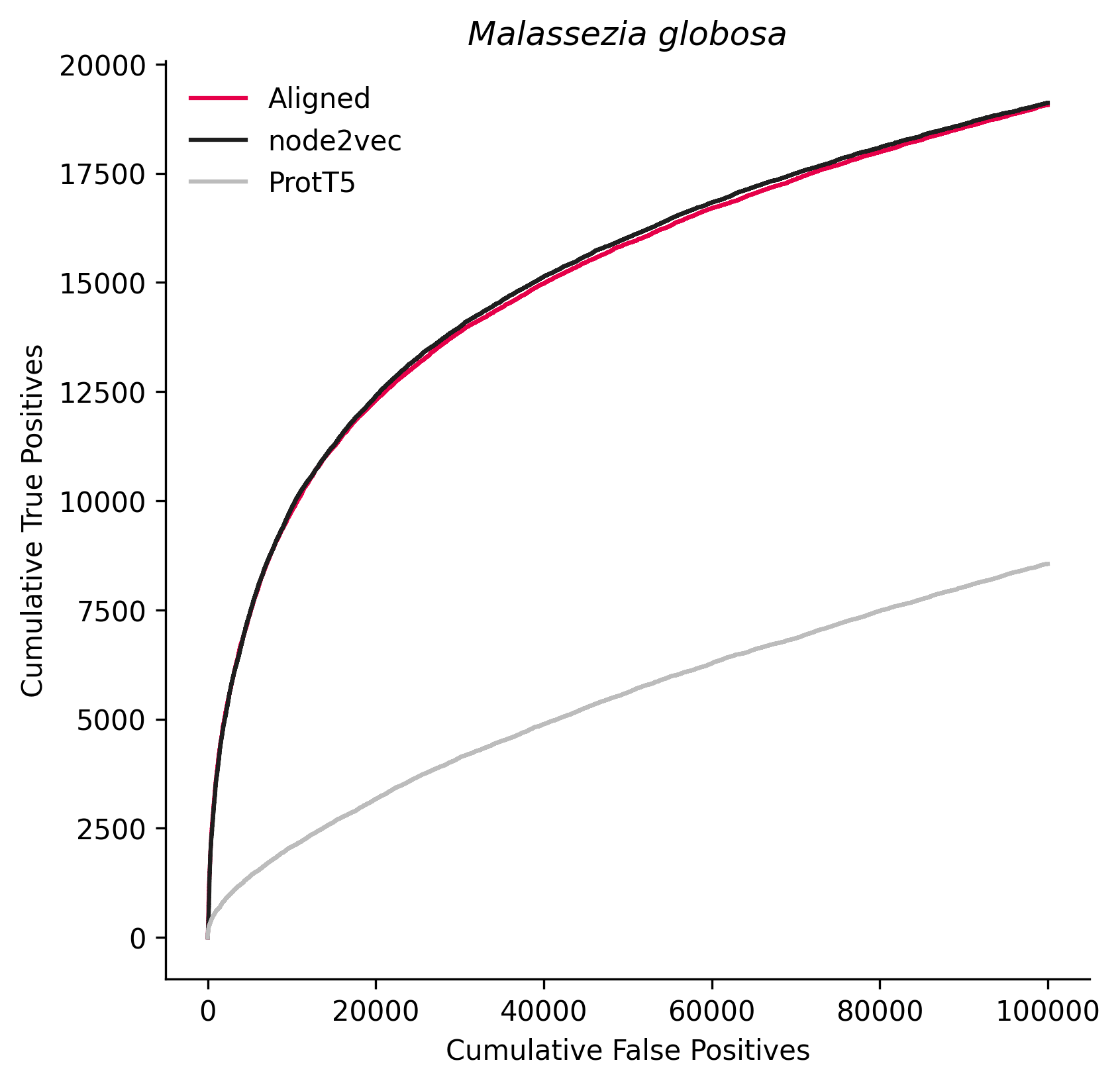

### 426428.png

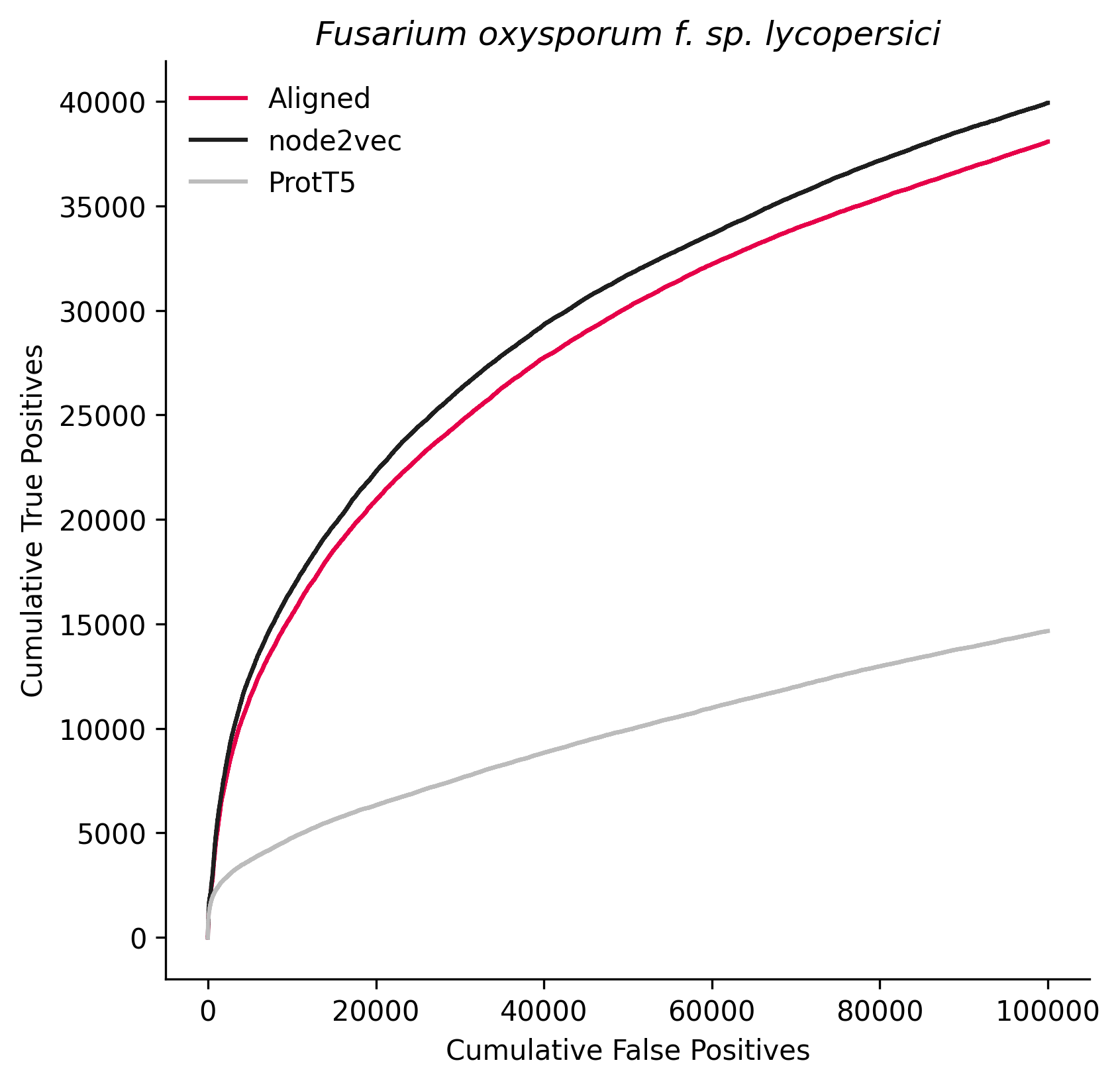

### 431241.png

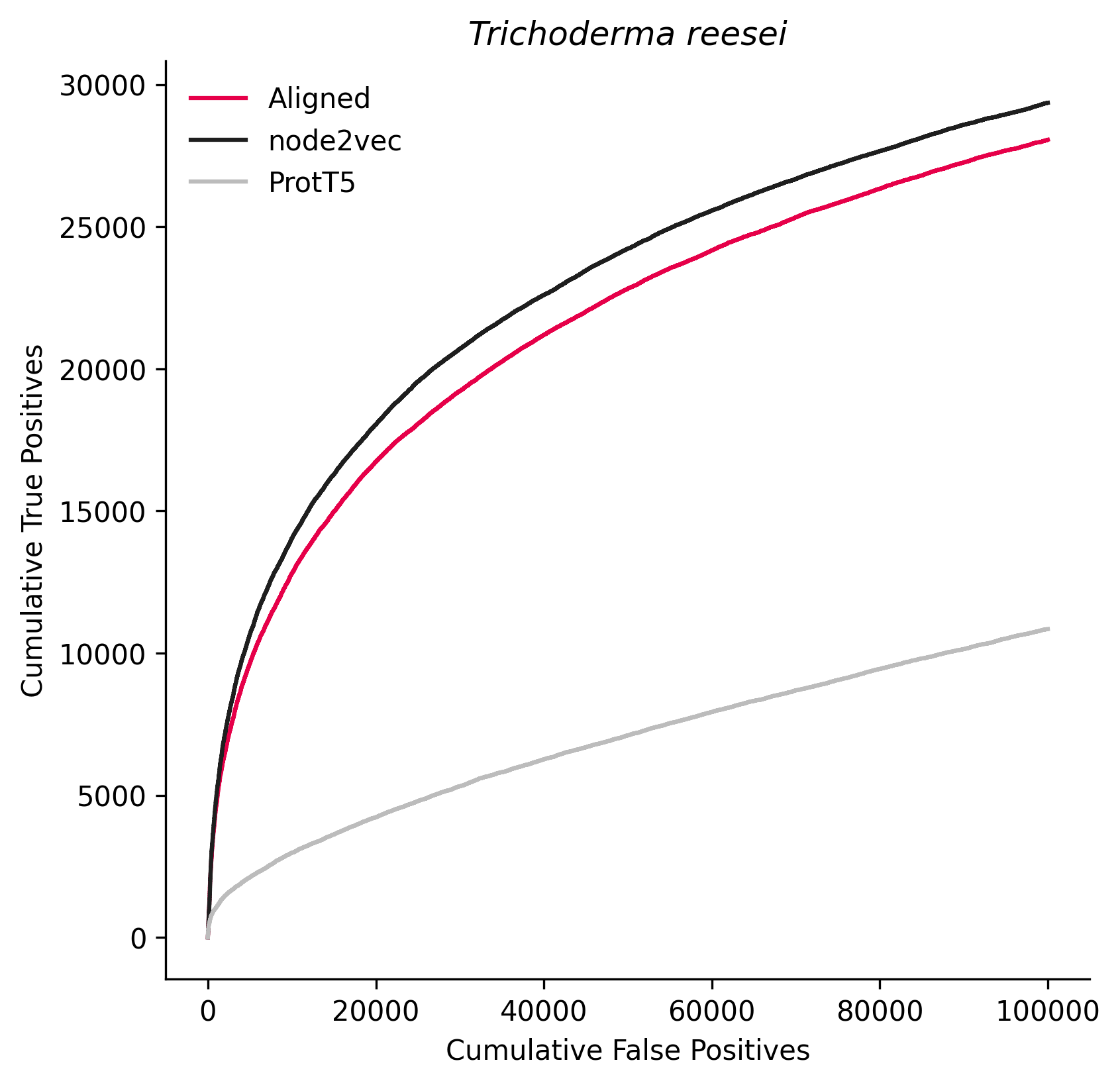

### 441960.png

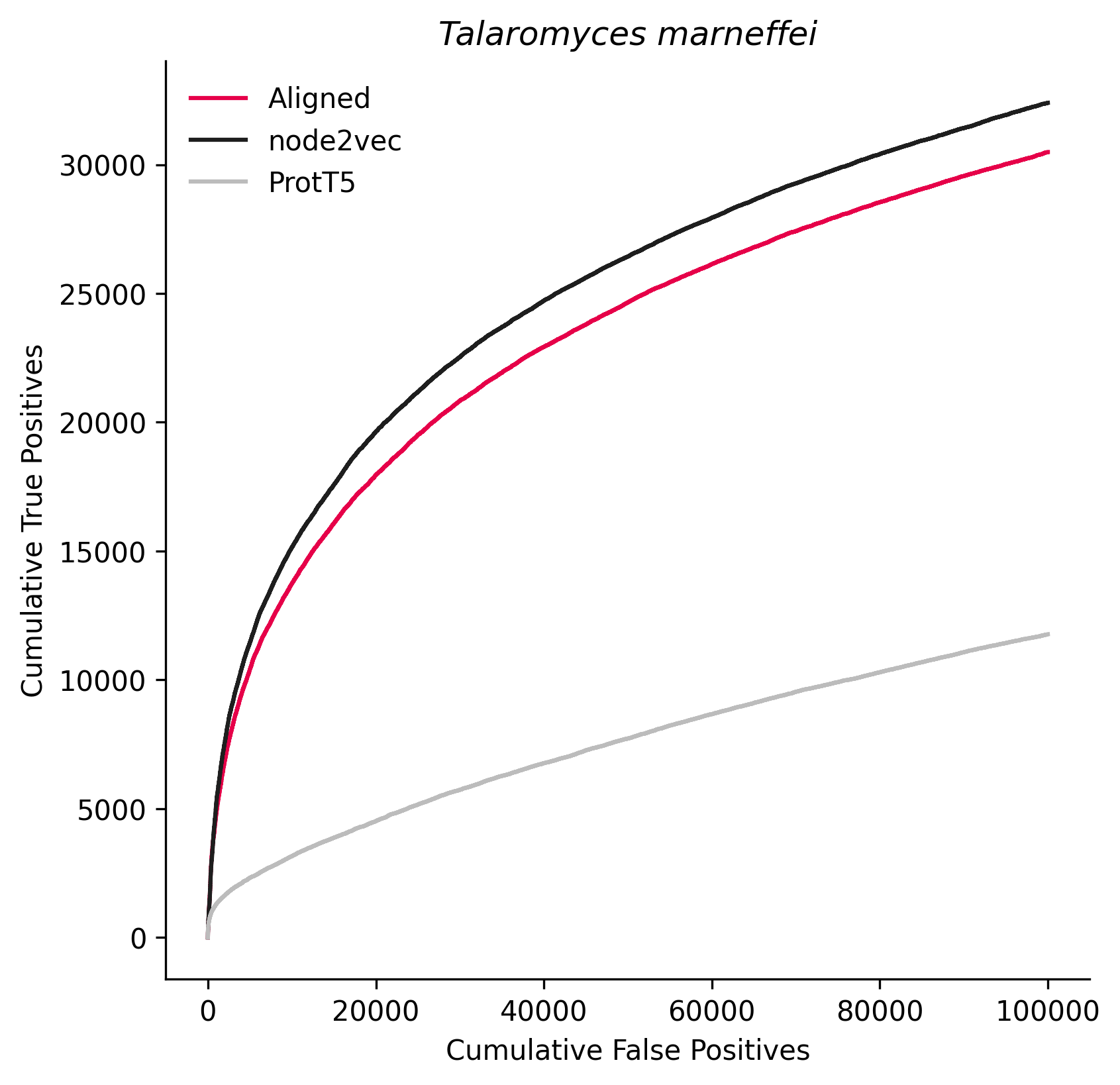

### 486041.png

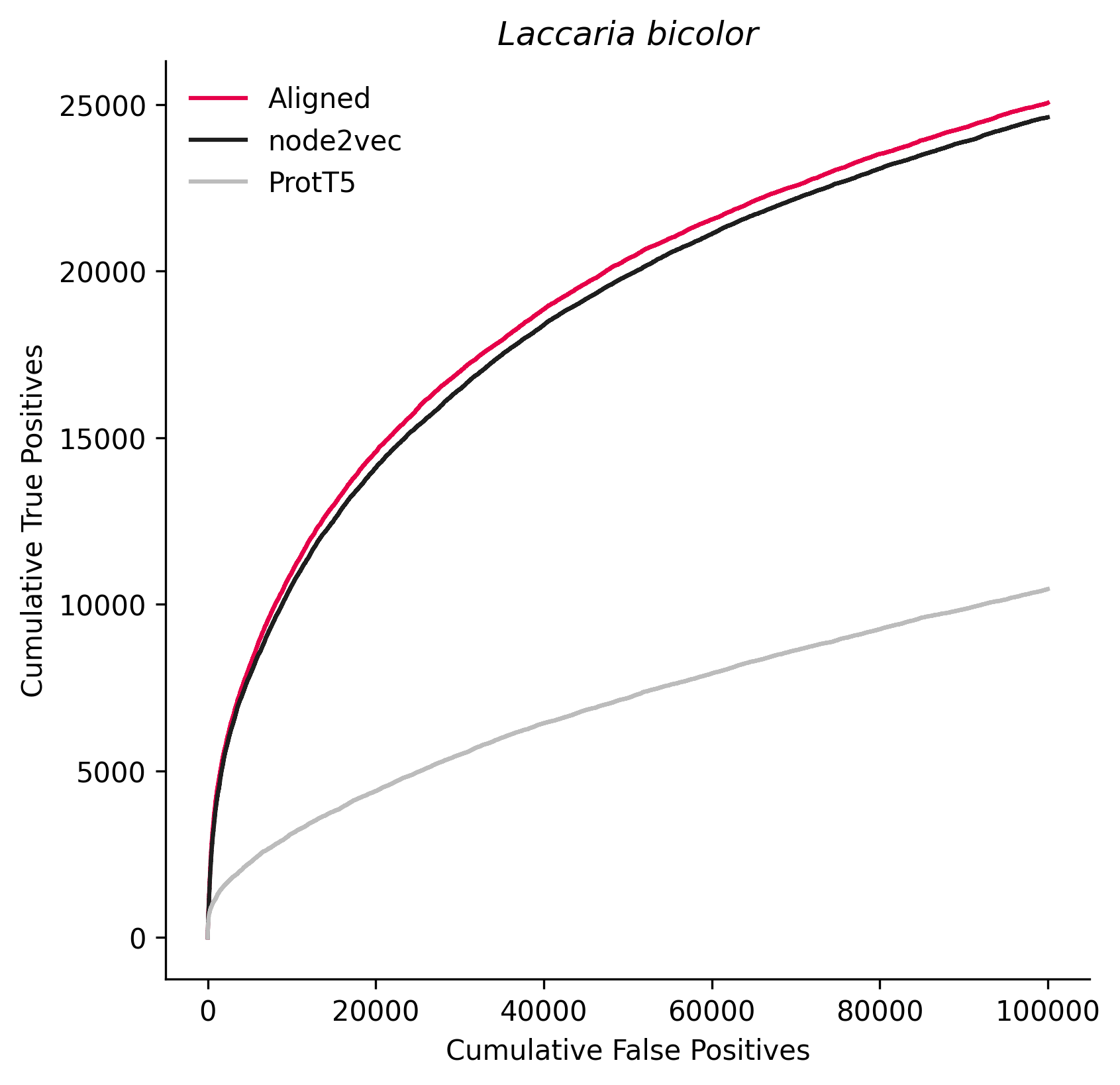

### 500485.png

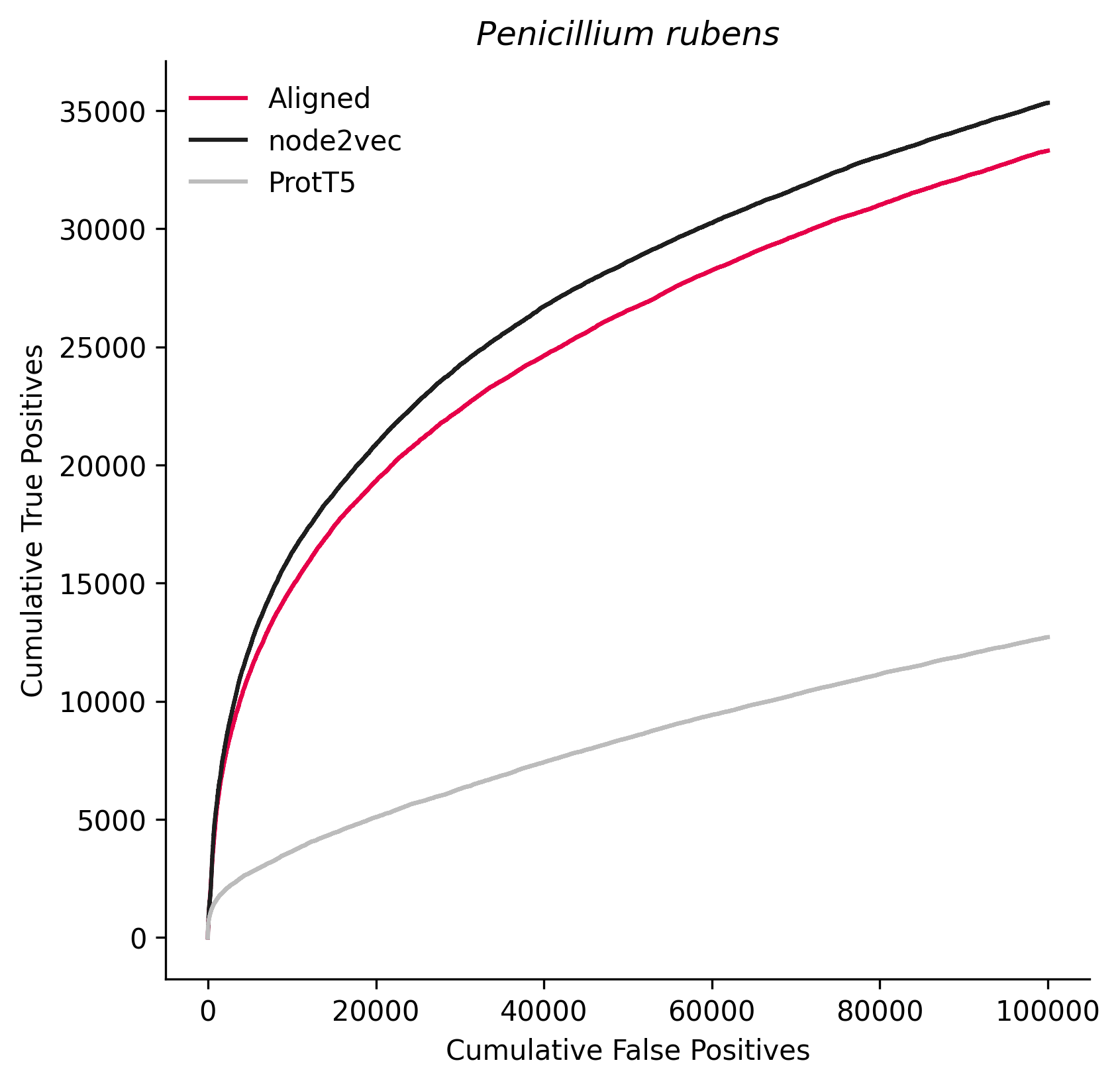

### 502779.png

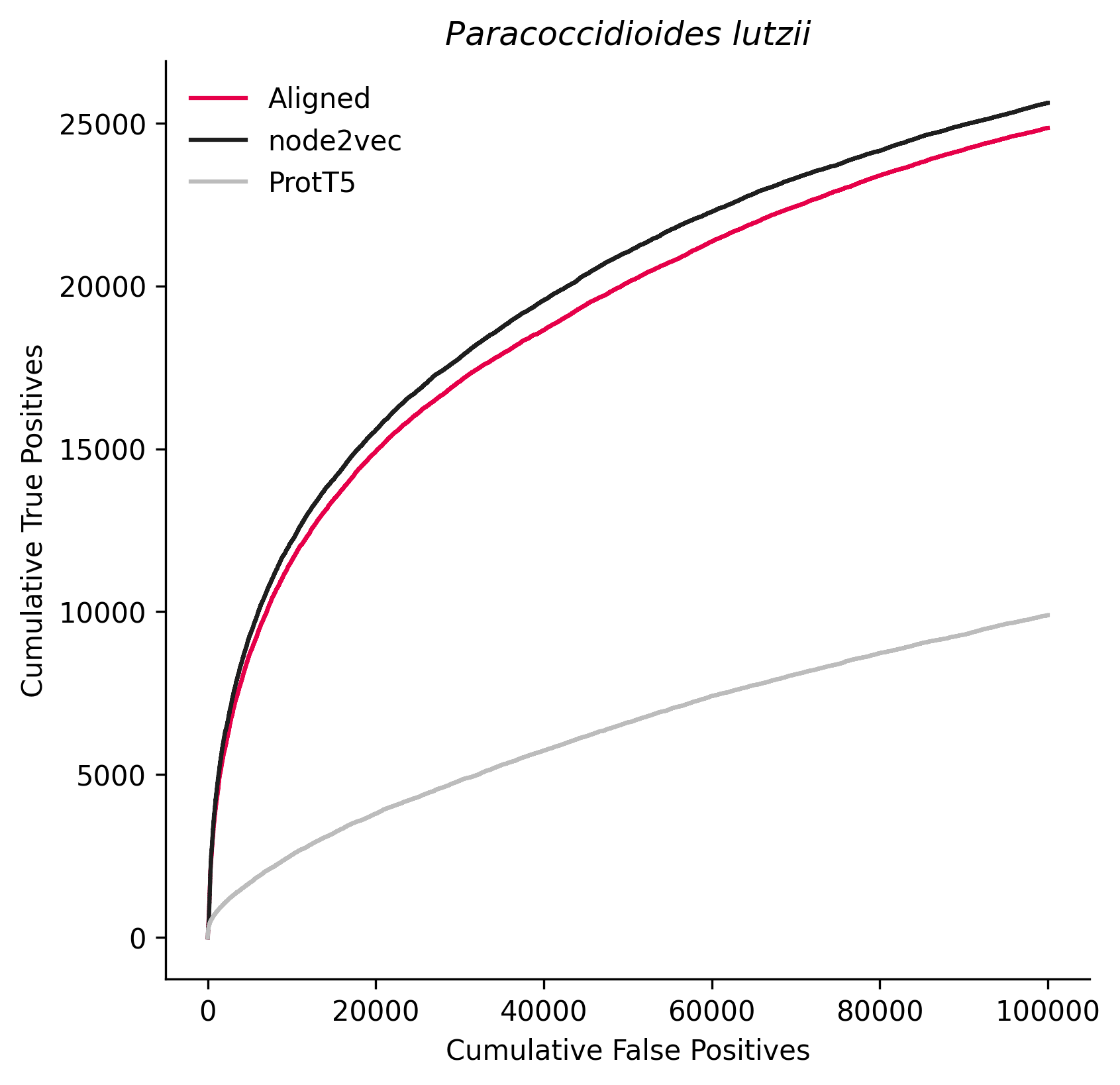

### 526221.png

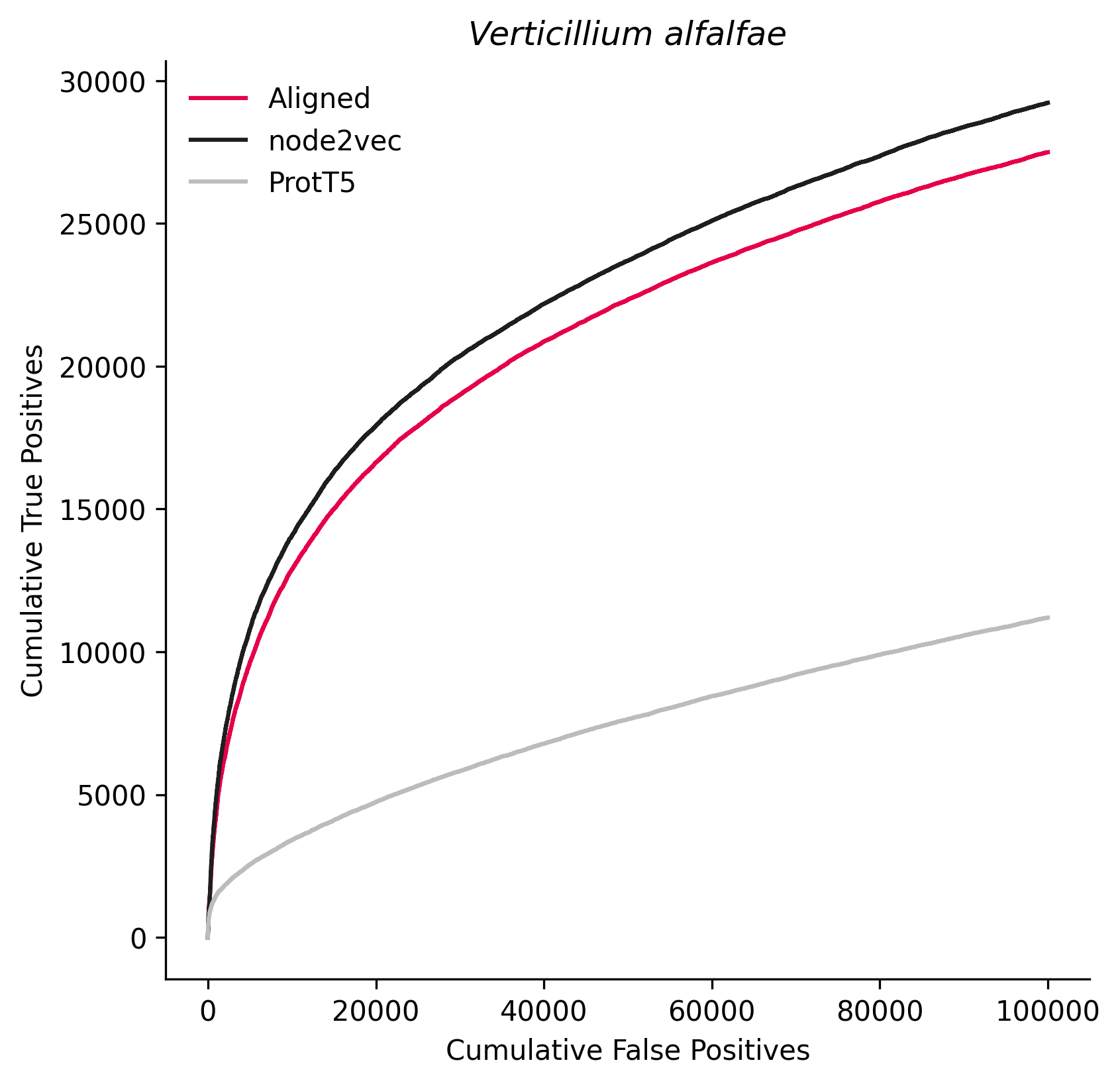

### 554065.png

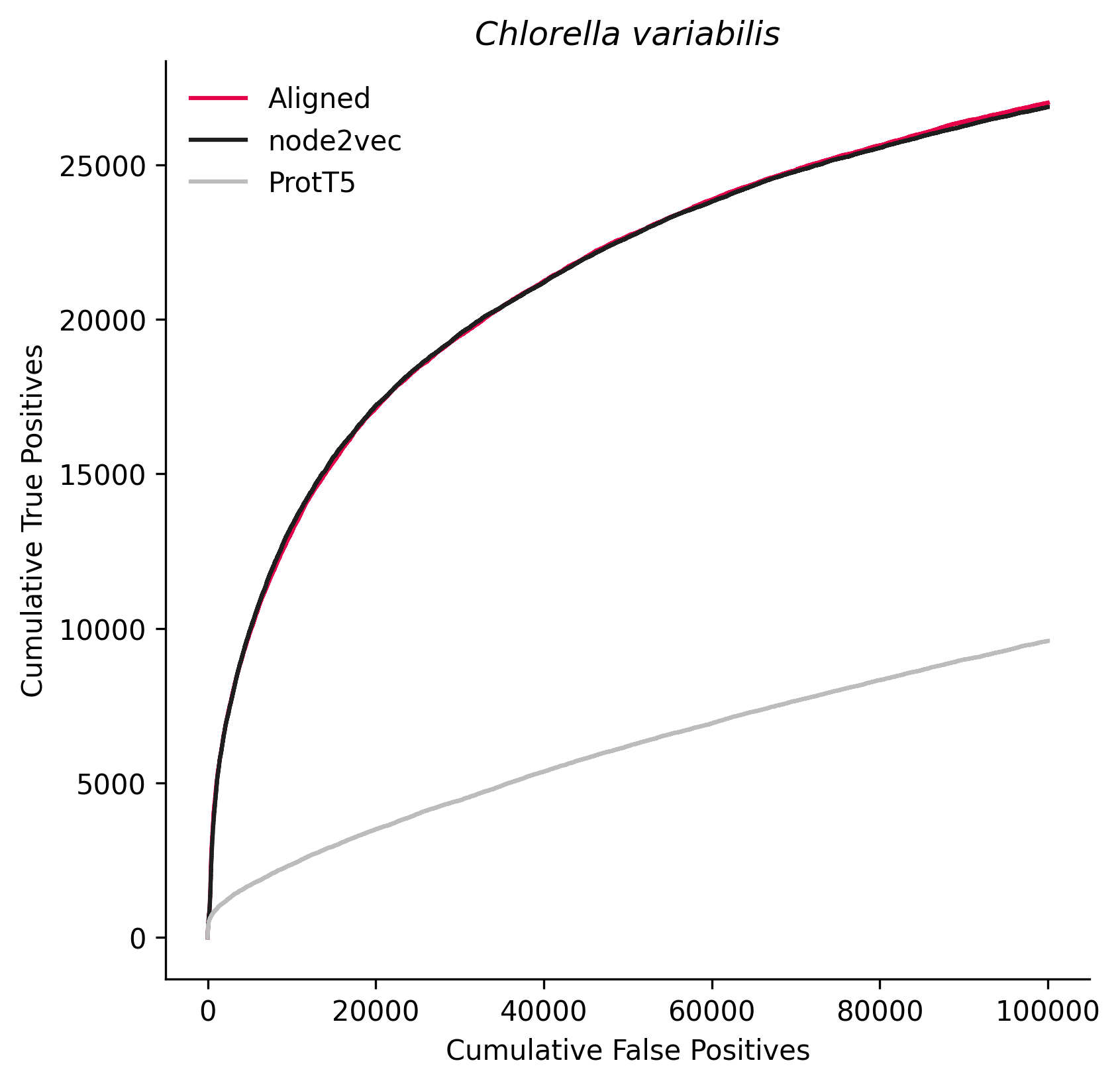

### 556484.png

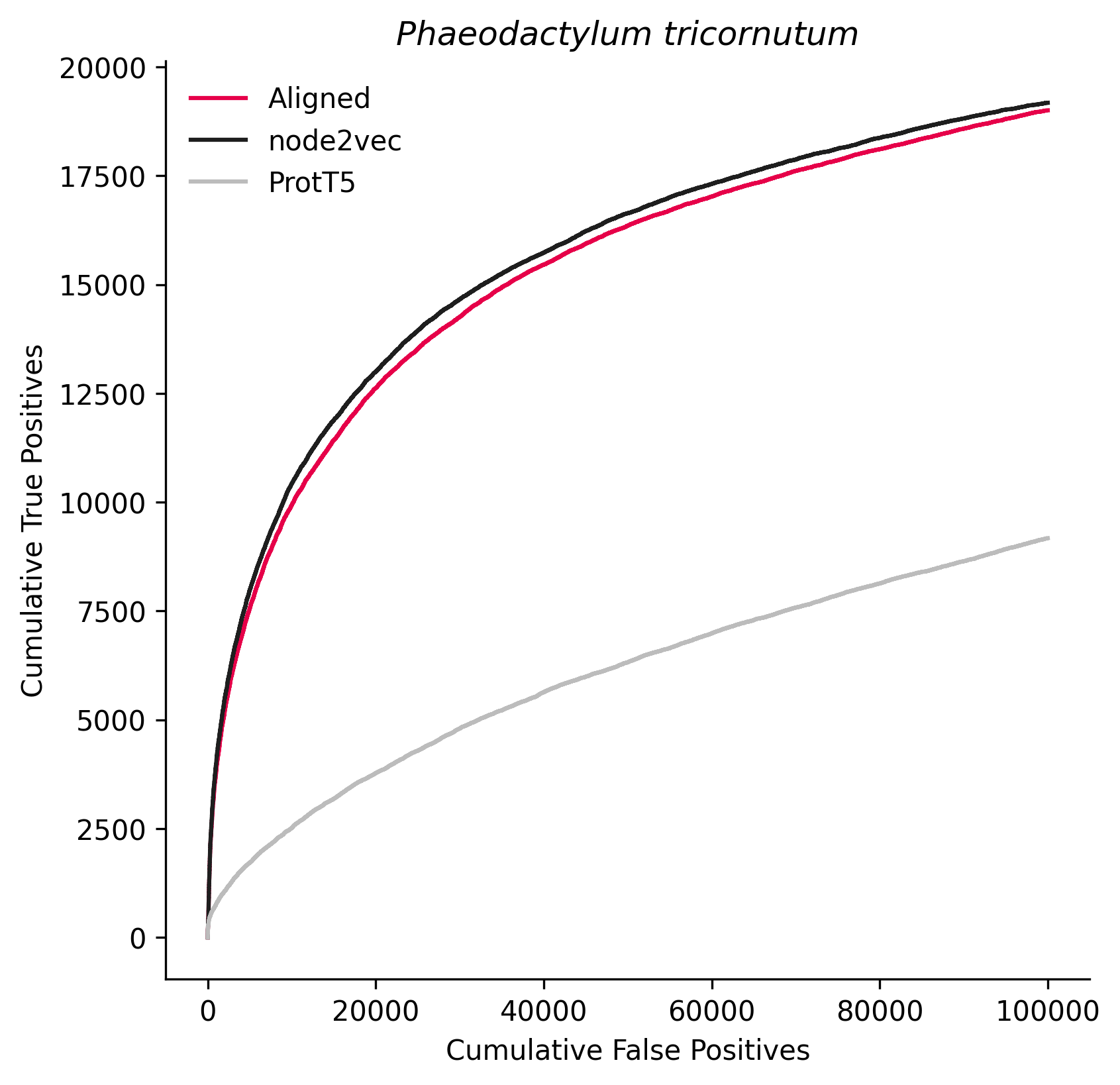

### 559298.png

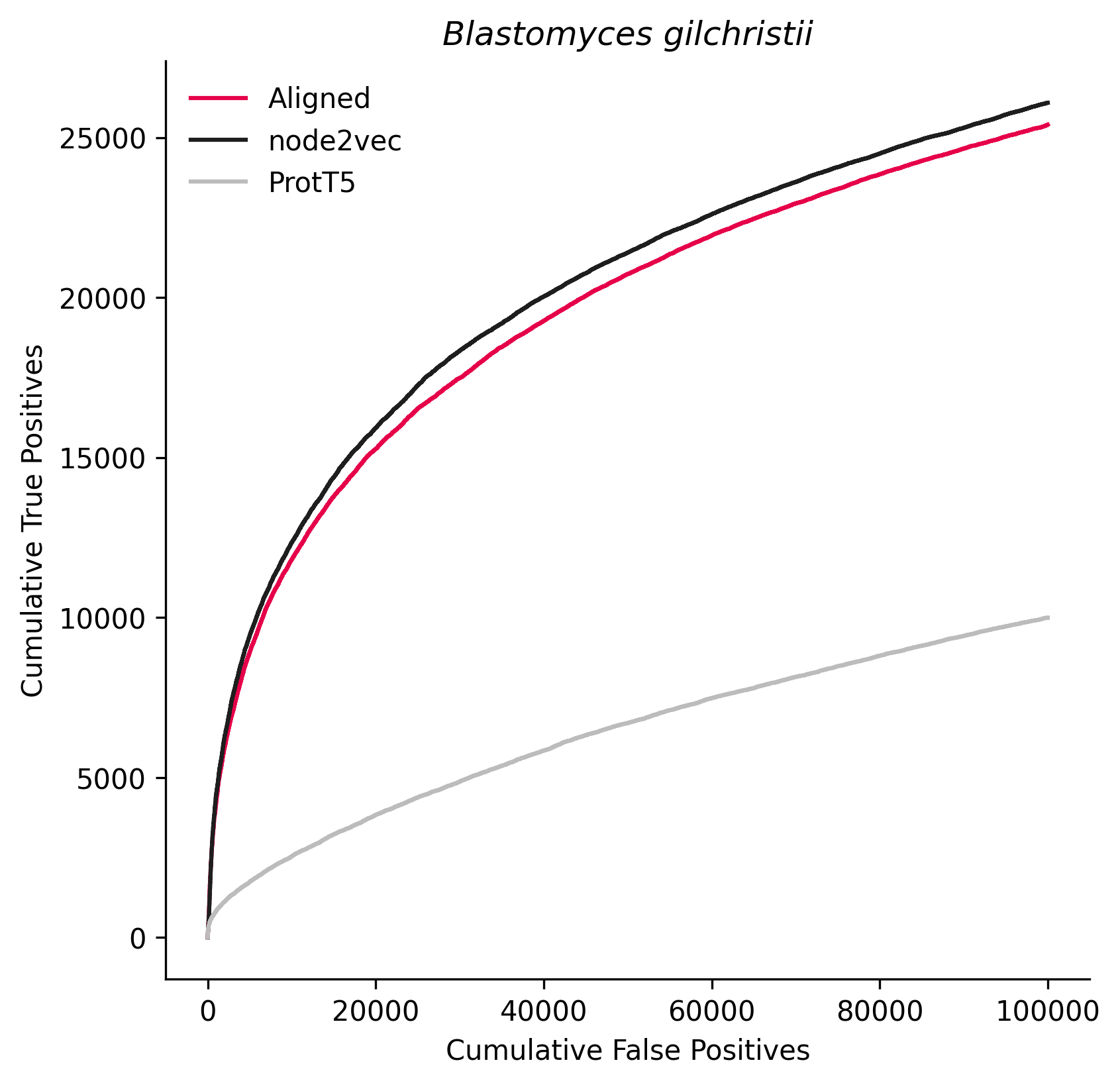

### 559307.png

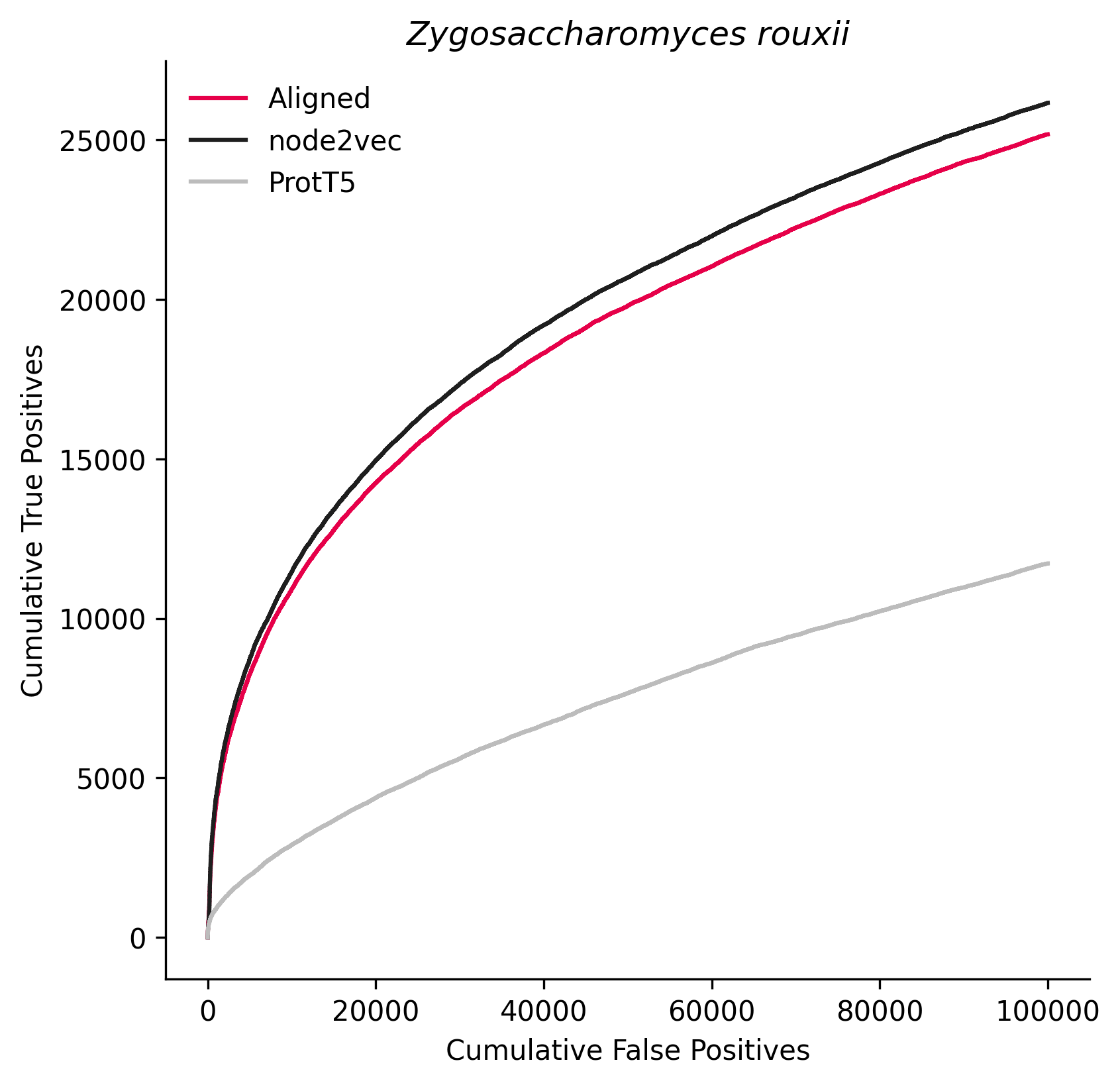

### 561896.png

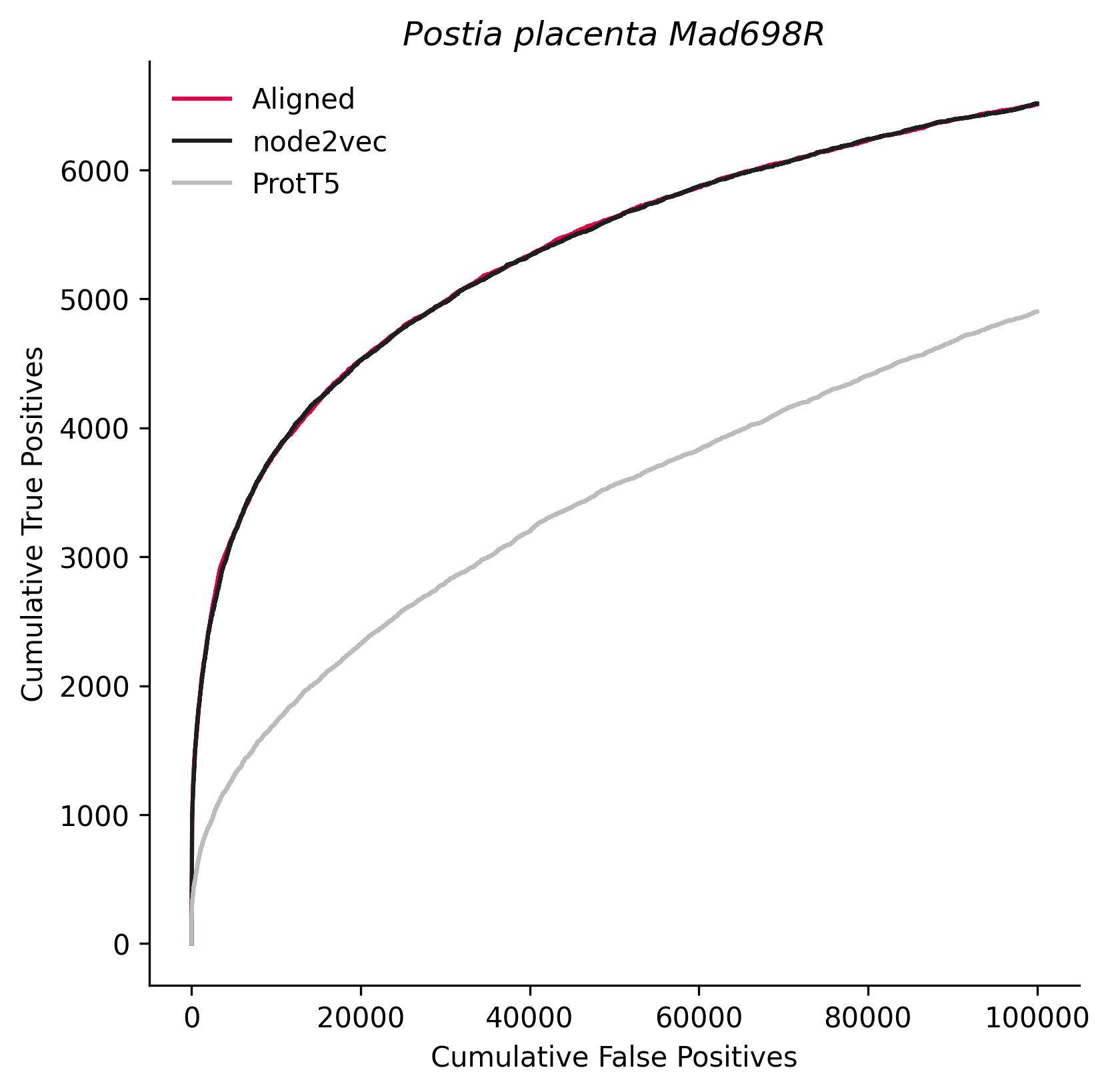

### 573729.png

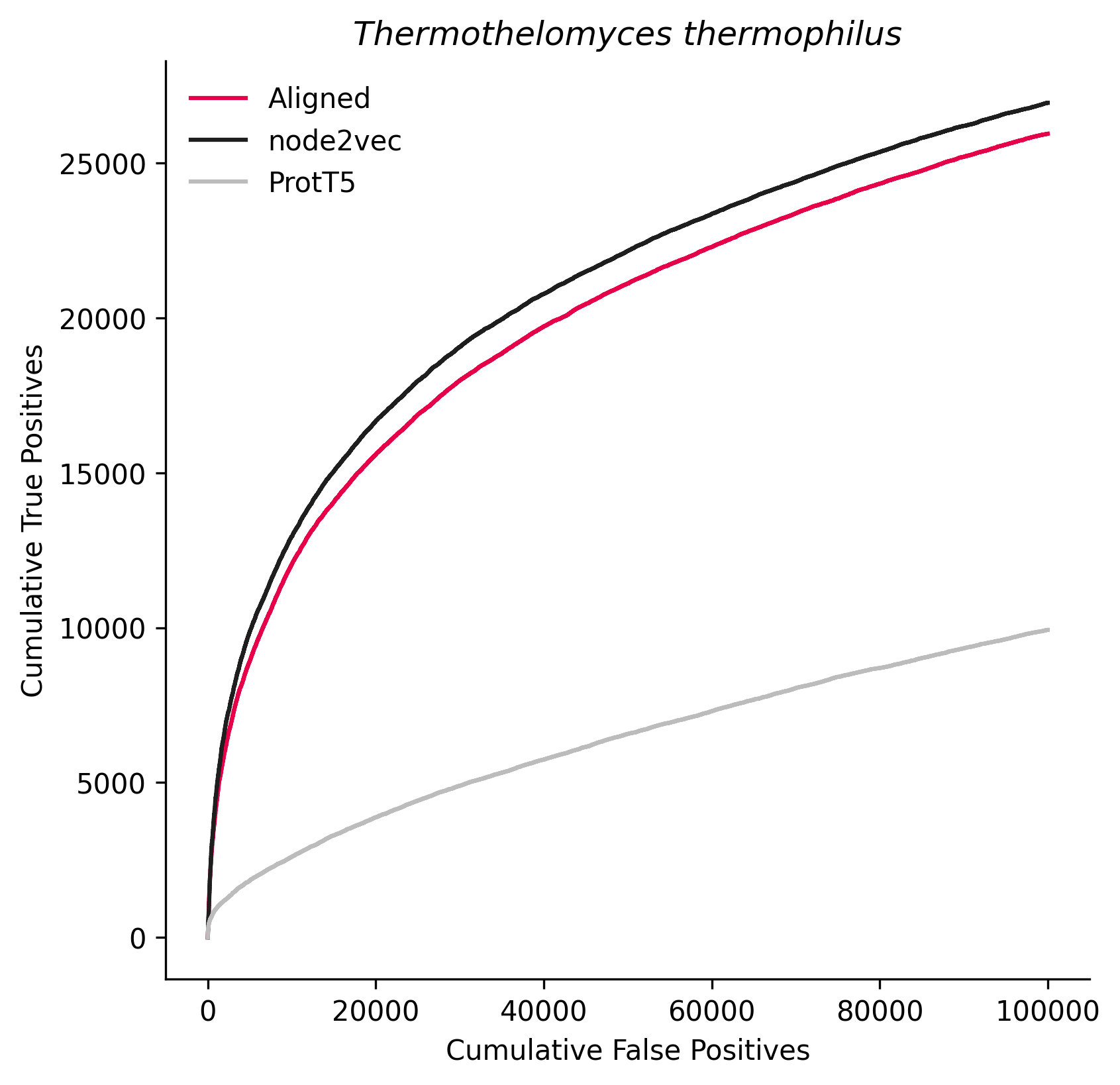

### 578460.png

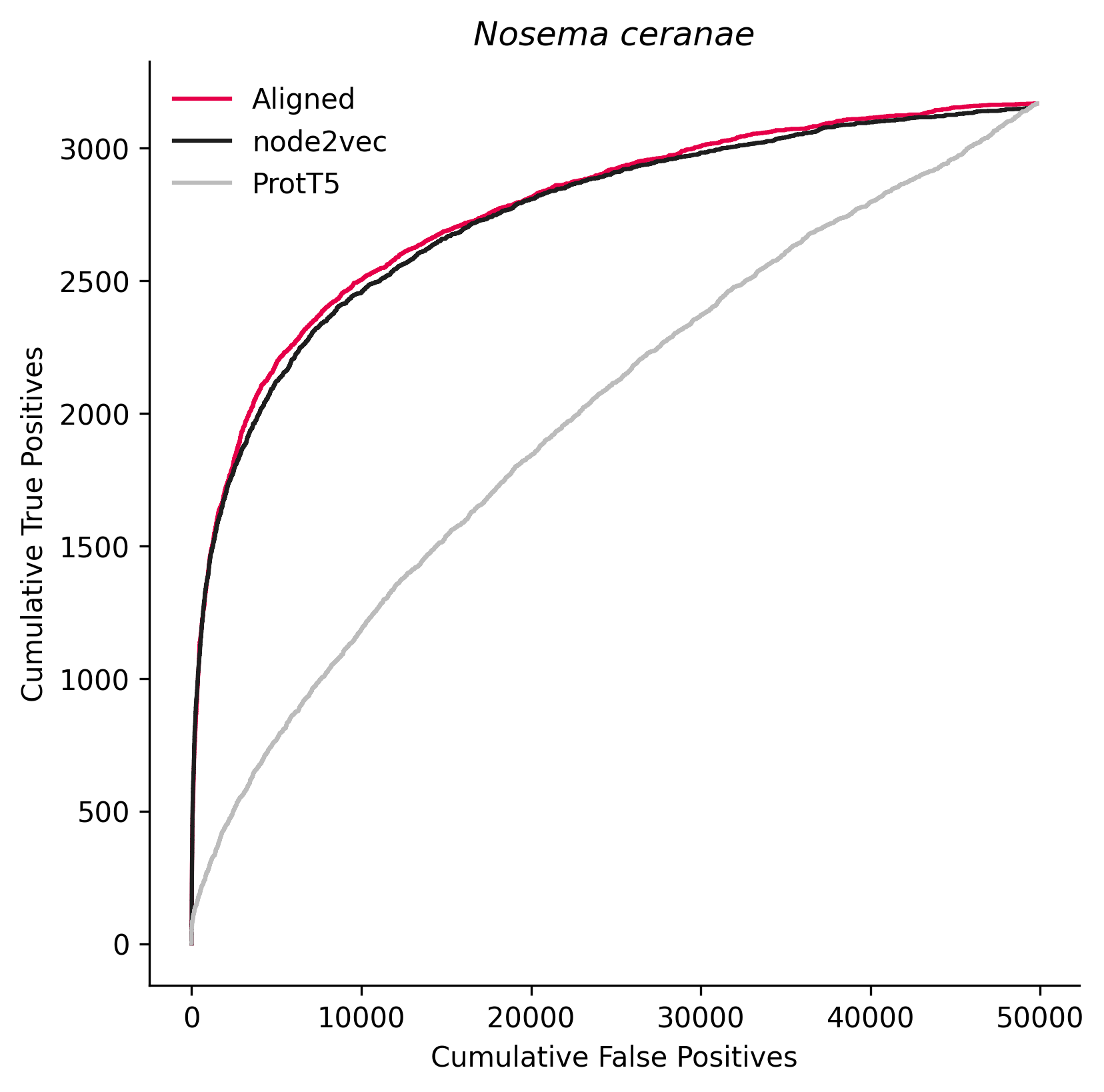

### 619300.png

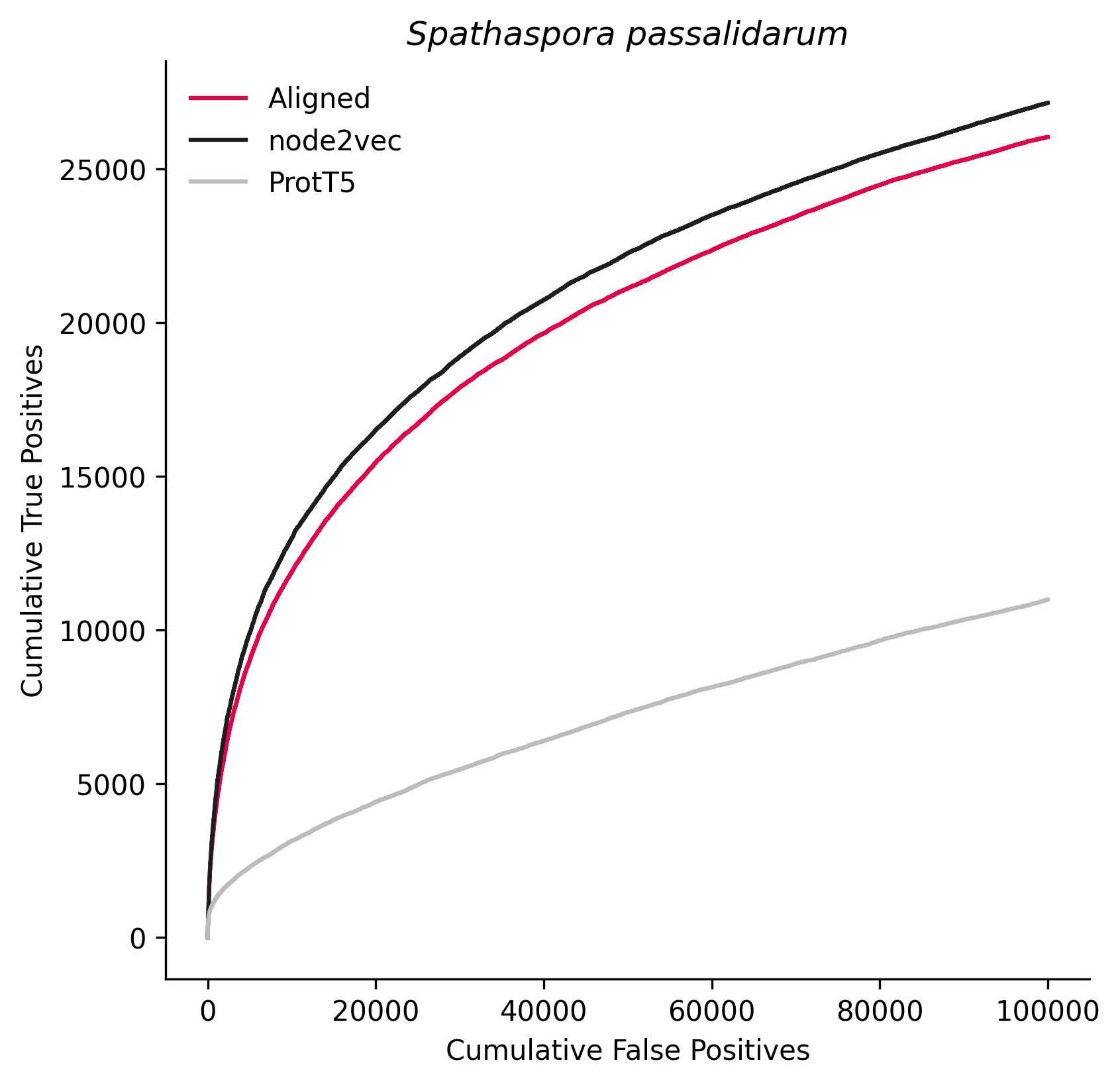

### 650164.png

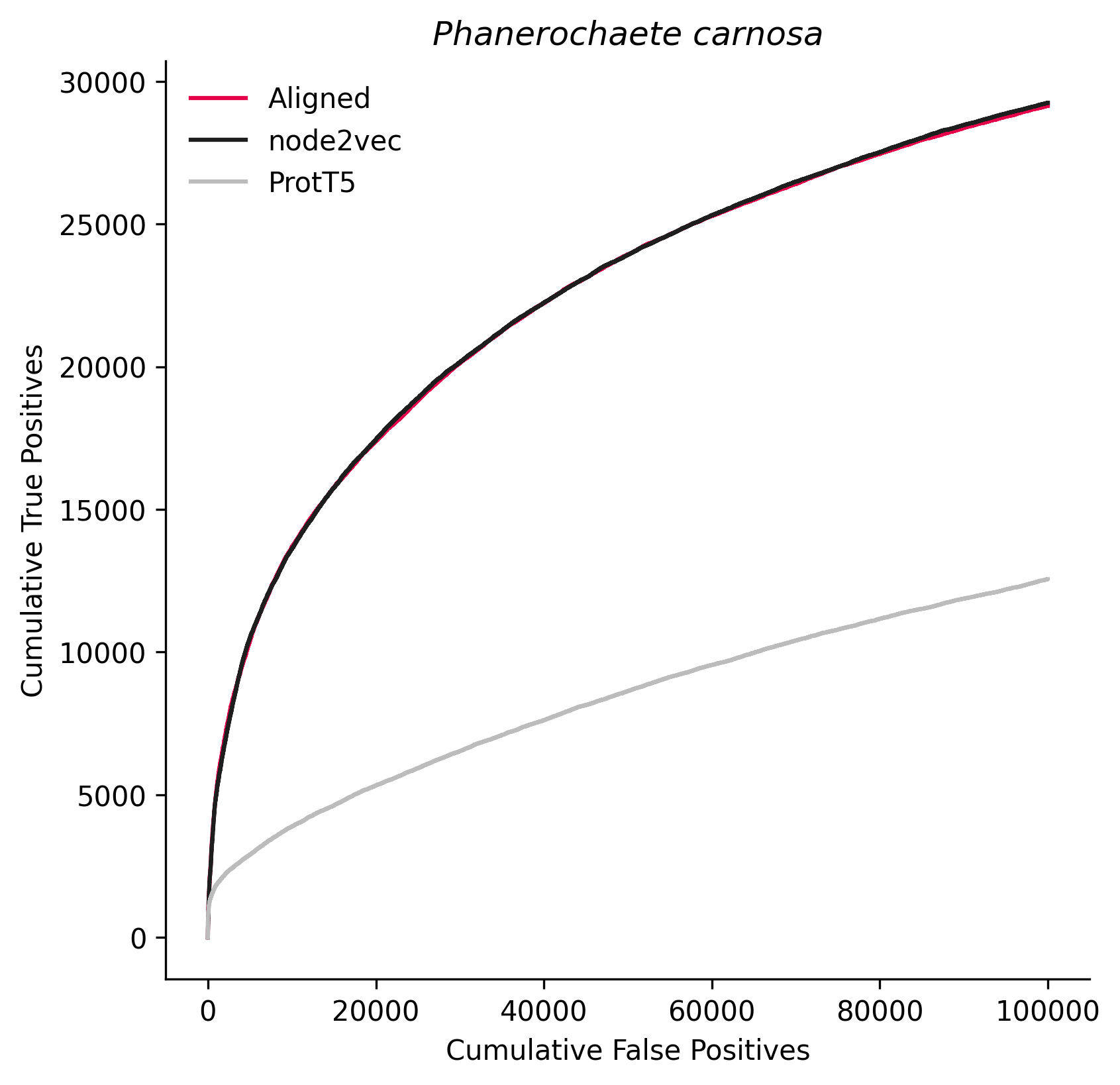

### 665079.png

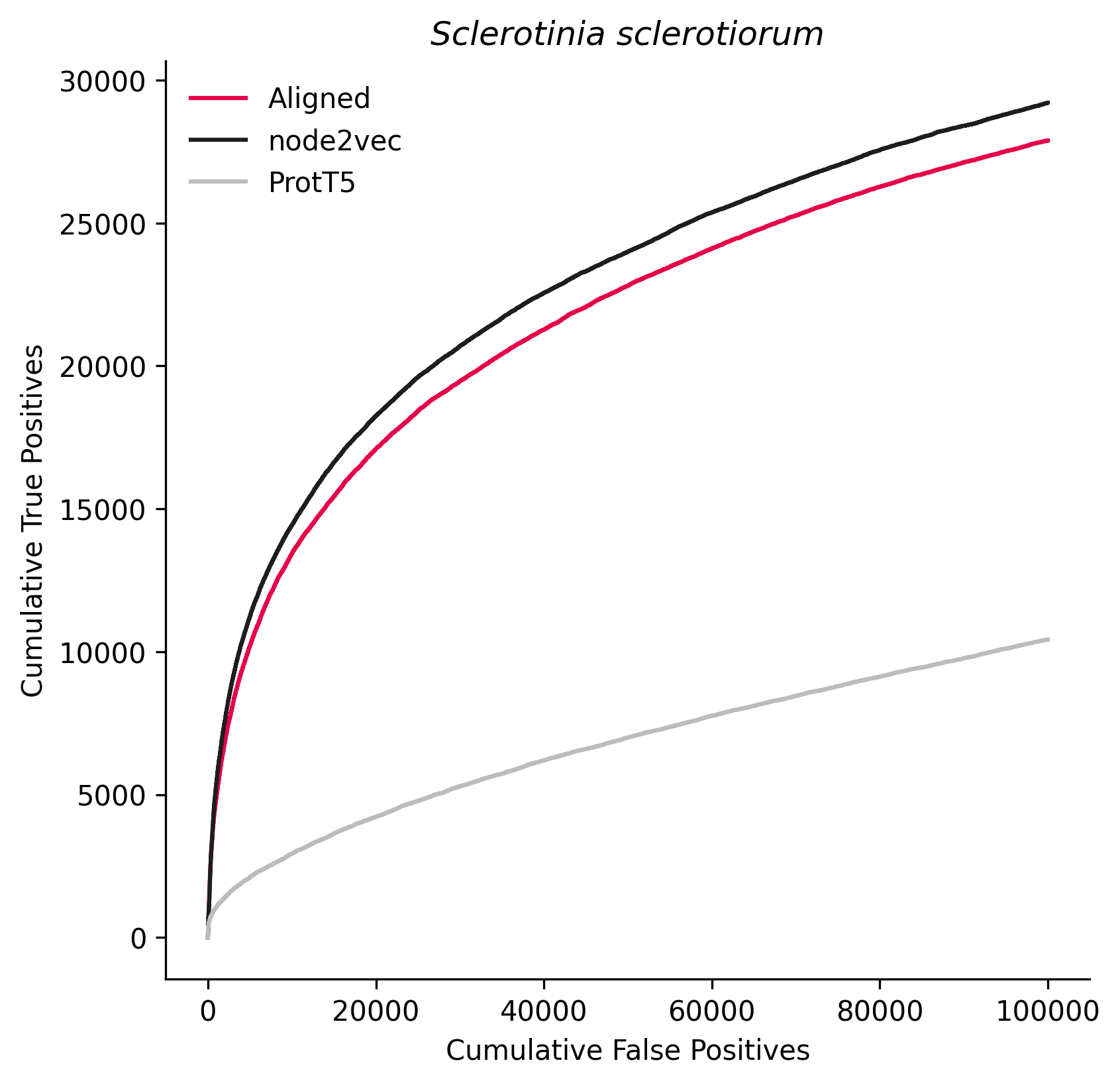

### 670483.png

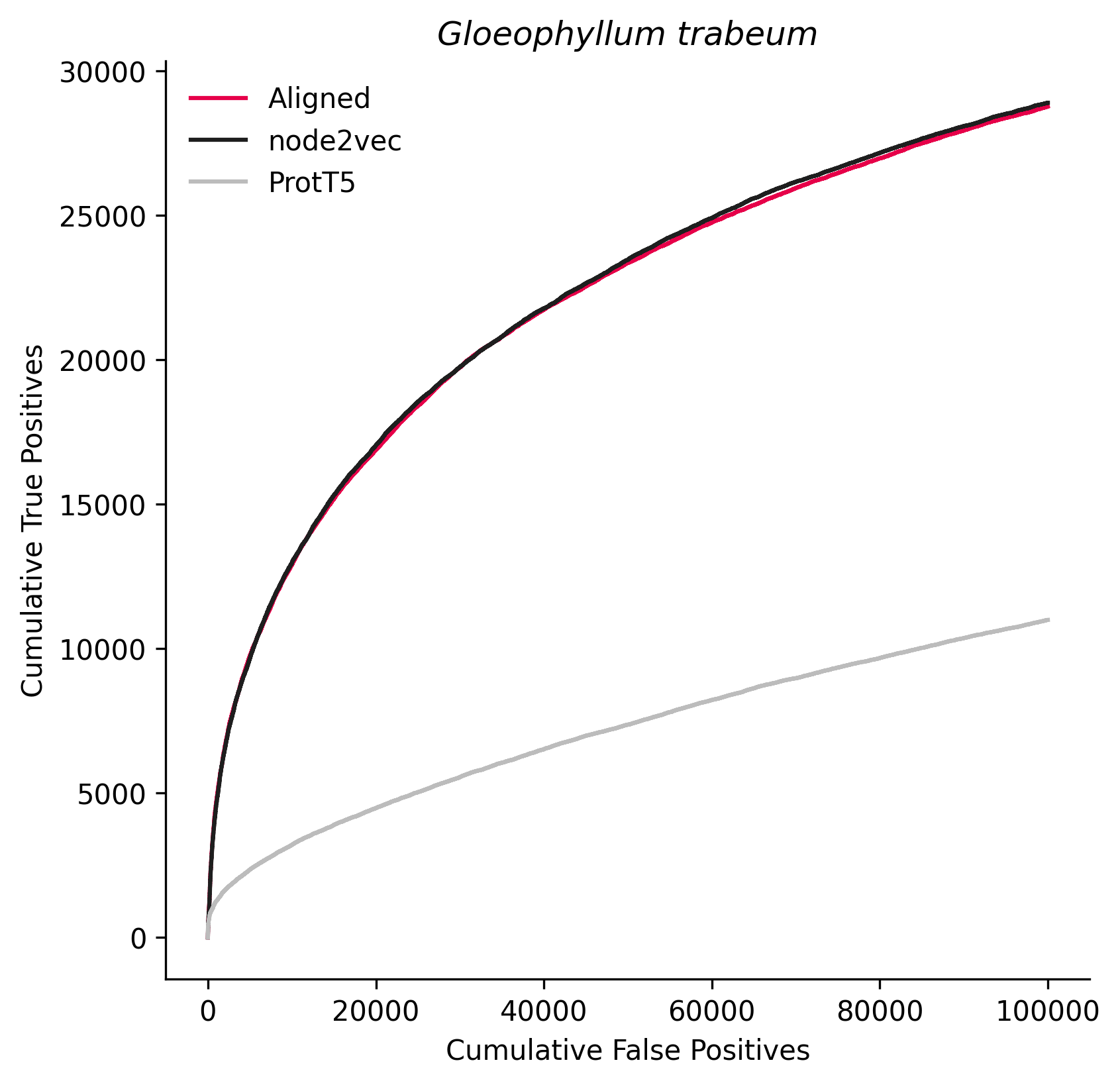

### 747525.png

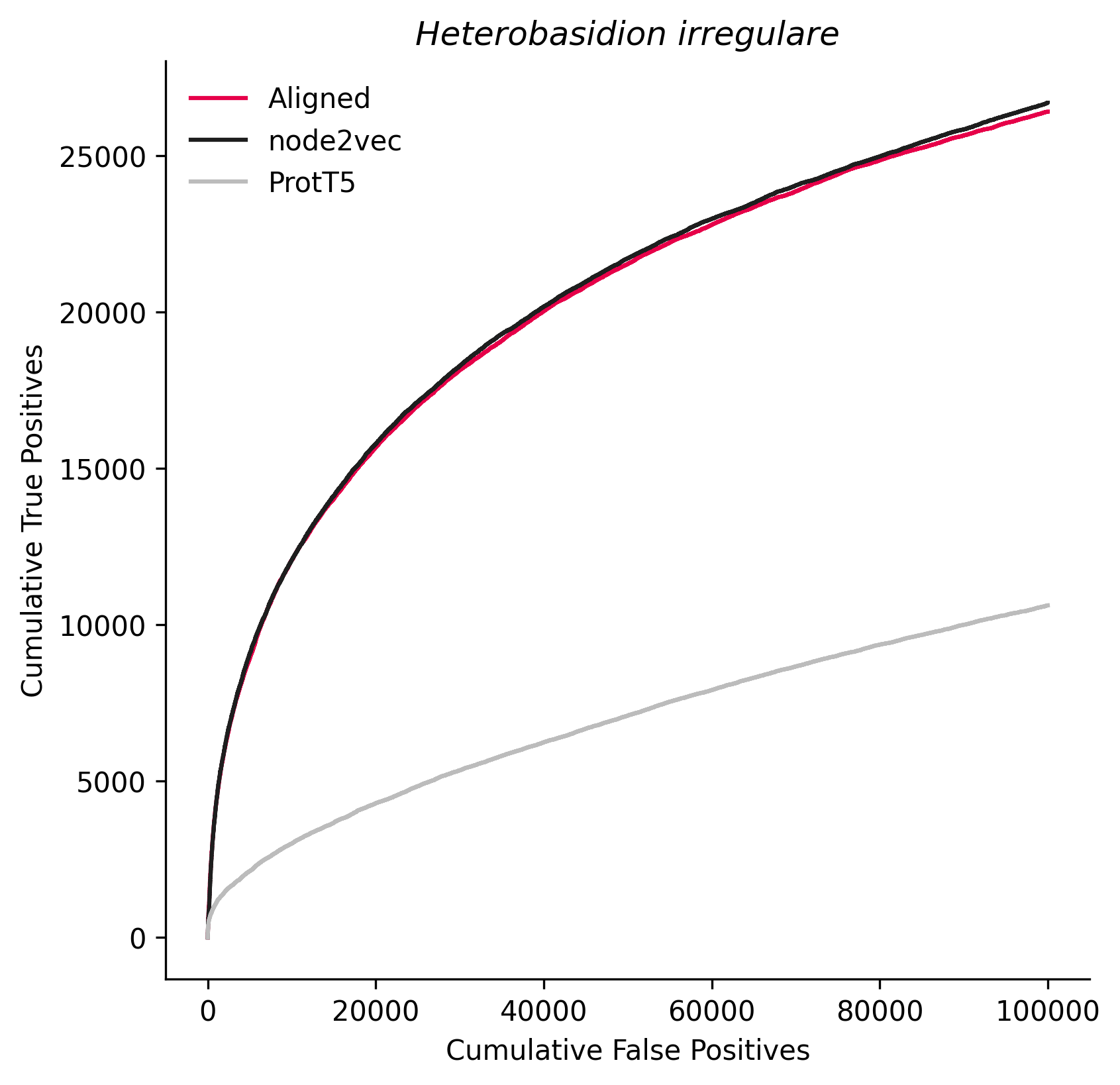

### 930090.png

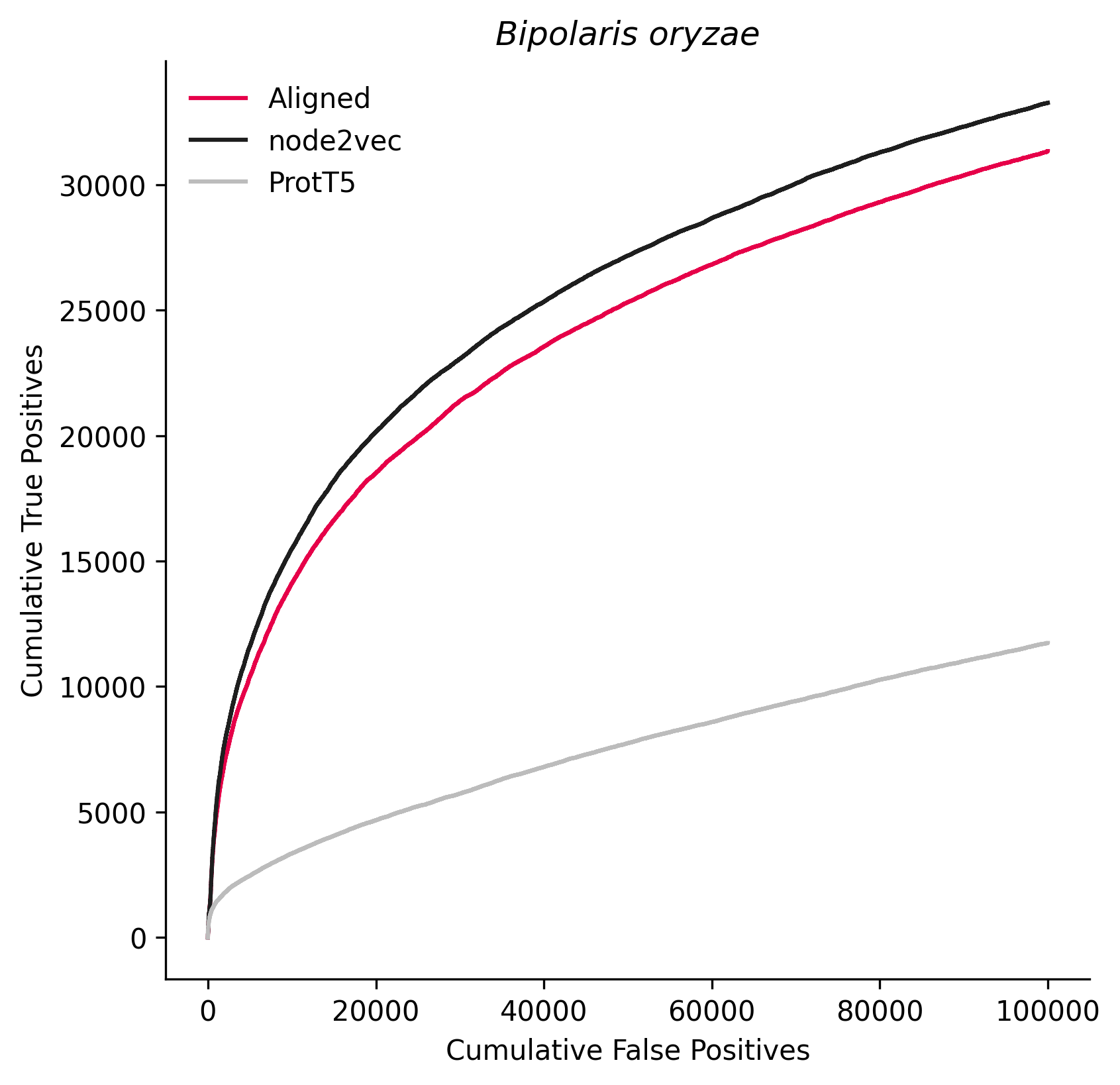

### 1194695.png

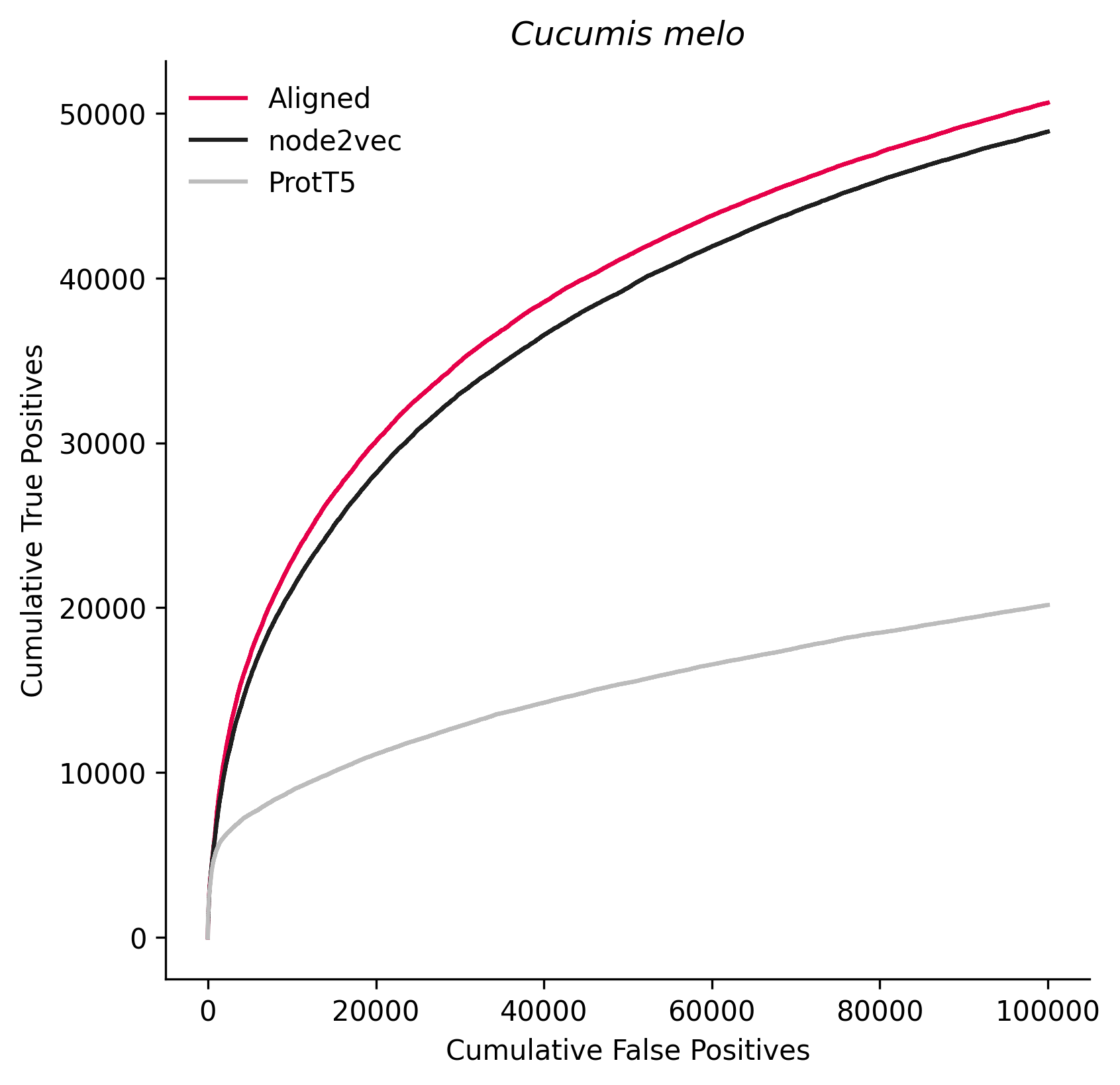

### 1229662.png

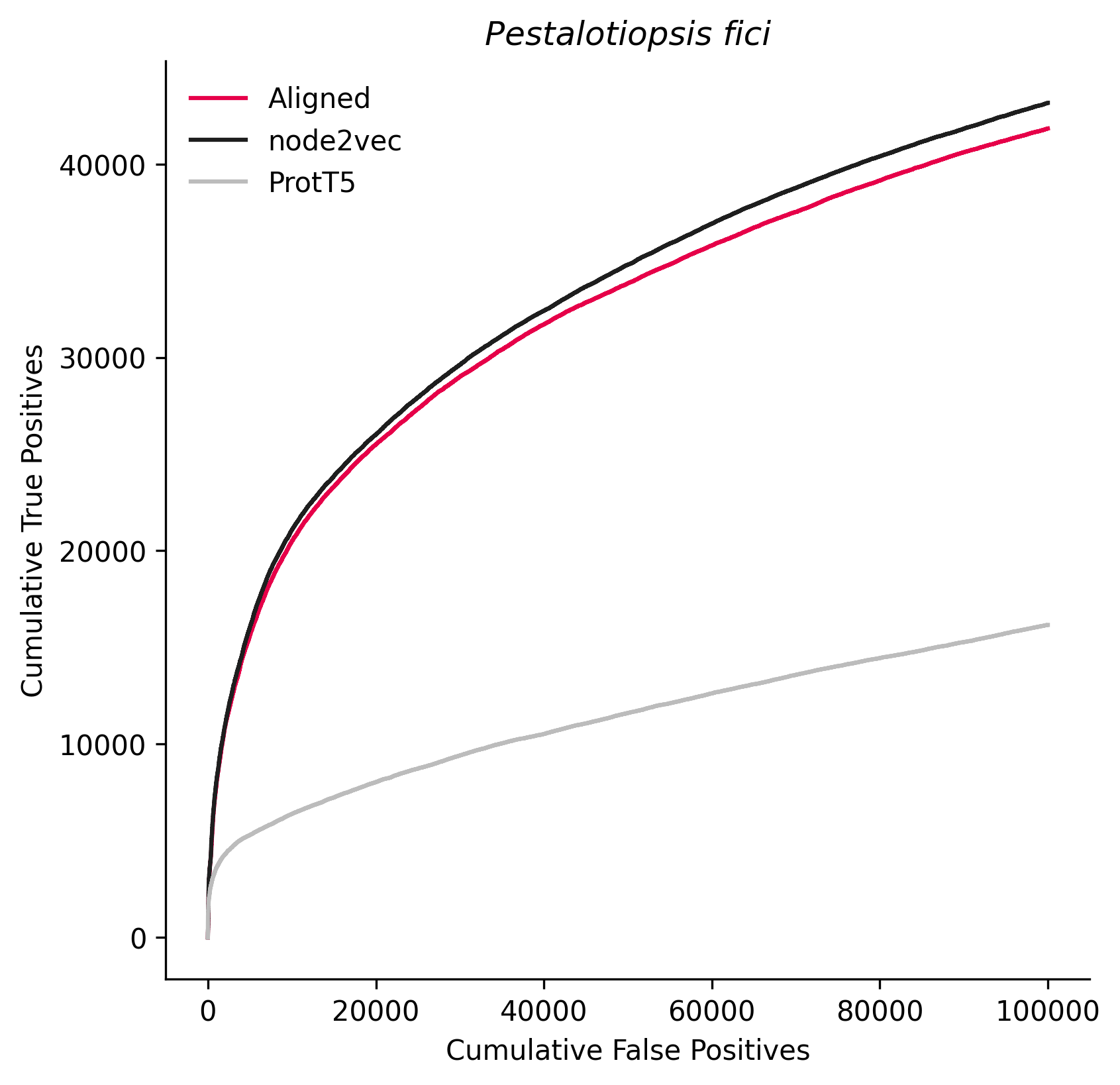

### 1676925.png

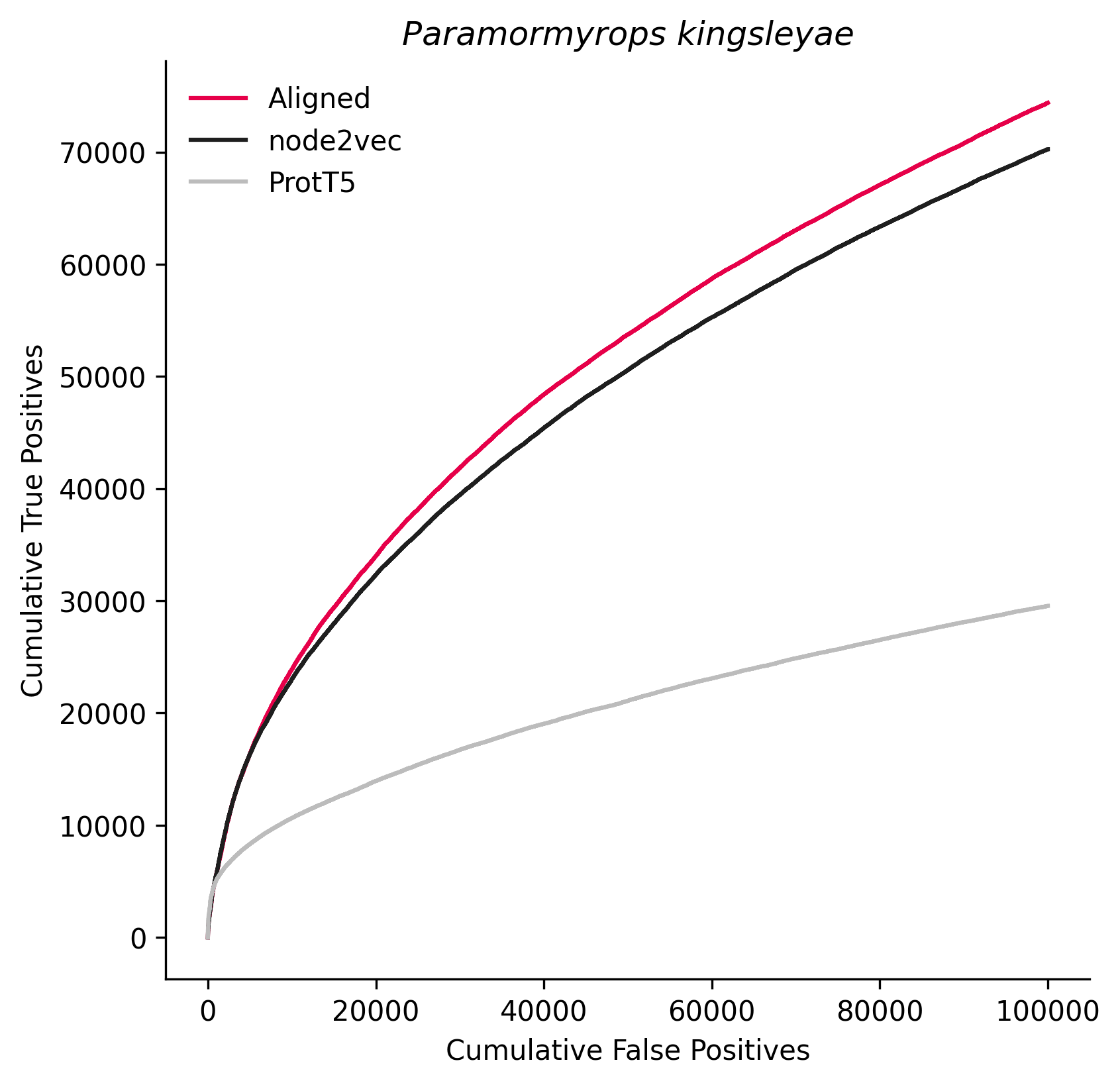

### 2015173.png

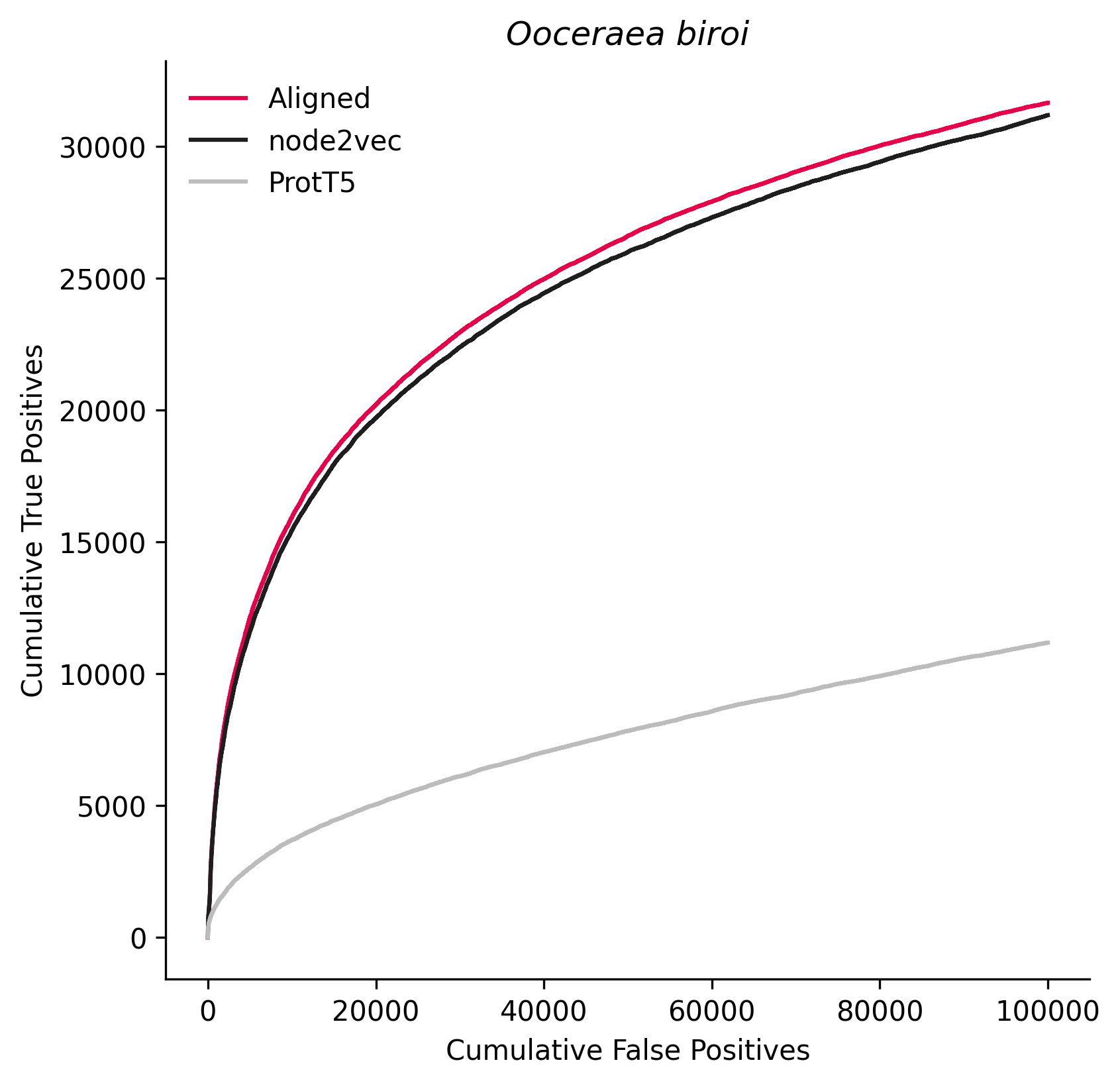
